# Complete sequencing of ape genomes

**DOI:** 10.1101/2024.07.31.605654

**Authors:** DongAhn Yoo, Arang Rhie, Prajna Hebbar, Francesca Antonacci, Glennis A. Logsdon, Steven J. Solar, Dmitry Antipov, Brandon D. Pickett, Yana Safonova, Francesco Montinaro, Yanting Luo, Joanna Malukiewicz, Jessica M. Storer, Jiadong Lin, Abigail N. Sequeira, Riley J. Mangan, Glenn Hickey, Graciela Monfort Anez, Parithi Balachandran, Anton Bankevich, Christine R. Beck, Arjun Biddanda, Matthew Borchers, Gerard G. Bouffard, Emry Brannan, Shelise Y. Brooks, Lucia Carbone, Laura Carrel, Agnes P. Chan, Juyun Crawford, Mark Diekhans, Eric Engelbrecht, Cedric Feschotte, Giulio Formenti, Gage H. Garcia, Luciana de Gennaro, David Gilbert, Richard E. Green, Andrea Guarracino, Ishaan Gupta, Diana Haddad, Junmin Han, Robert S. Harris, Gabrielle A. Hartley, William T. Harvey, Michael Hiller, Kendra Hoekzema, Marlys L. Houck, Hyeonsoo Jeong, Kaivan Kamali, Manolis Kellis, Bryce Kille, Chul Lee, Youngho Lee, William Lees, Alexandra P. Lewis, Qiuhui Li, Mark Loftus, Yong Hwee Eddie Loh, Hailey Loucks, Jian Ma, Yafei Mao, Juan F. I. Martinez, Patrick Masterson, Rajiv C. McCoy, Barbara McGrath, Sean McKinney, Britta S. Meyer, Karen H. Miga, Saswat K. Mohanty, Katherine M. Munson, Karol Pal, Matt Pennell, Pavel A. Pevzner, David Porubsky, Tamara Potapova, Francisca R. Ringeling, Joana L. Rocha, Oliver A. Ryder, Samuel Sacco, Swati Saha, Takayo Sasaki, Michael C. Schatz, Nicholas J. Schork, Cole Shanks, Linnéa Smeds, Dongmin R. Son, Cynthia Steiner, Alexander P. Sweeten, Michael G. Tassia, Françoise Thibaud-Nissen, Edmundo Torres-González, Mihir Trivedi, Wenjie Wei, Julie Wertz, Muyu Yang, Panpan Zhang, Shilong Zhang, Yang Zhang, Zhenmiao Zhang, Sarah A. Zhao, Yixin Zhu, Erich D. Jarvis, Jennifer L. Gerton, Iker Rivas-González, Benedict Paten, Zachary A. Szpiech, Christian D. Huber, Tobias L. Lenz, Miriam K. Konkel, Soojin V. Yi, Stefan Canzar, Corey T. Watson, Peter H. Sudmant, Erin Molloy, Erik Garrison, Craig B. Lowe, Mario Ventura, Rachel J. O’Neill, Sergey Koren, Kateryna D. Makova, Adam M. Phillippy, Evan E. Eichler

## Abstract

We present haplotype-resolved reference genomes and comparative analyses of six ape species, namely: chimpanzee, bonobo, gorilla, Bornean orangutan, Sumatran orangutan, and siamang. We achieve chromosome-level contiguity with unparalleled sequence accuracy (<1 error in 500,000 base pairs), completely sequencing 215 gapless chromosomes telomere-to-telomere. We resolve challenging regions, such as the major histocompatibility complex and immunoglobulin loci, providing more in-depth evolutionary insights. Comparative analyses, including human, allow us to investigate the evolution and diversity of regions previously uncharacterized or incompletely studied without bias from mapping to the human reference. This includes newly minted gene families within lineage-specific segmental duplications, centromeric DNA, acrocentric chromosomes, and subterminal heterochromatin. This resource should serve as a definitive baseline for all future evolutionary studies of humans and our closest living ape relatives.

## INTRODUCTION

High-quality sequencing of ape genomes has been a high priority of the human genetics and genomics community since the initial sequencing of the human genome in 2001^1,2^. Sequencing of these genomes is critical for reconstructing the evolutionary history of every base pair of the human genome—one of the grand challenges put forward to the genomics community after the release of the first draft of the Human Genome Project^3^. As a result, there have been numerous publications ranging from initial draft genomes to significant updates over the last two decades^4–7^. Due to the repetitive nature of ape genomes, however, complete assemblies have not been achieved. Current references lack sequence resolution of some of the most dynamic genomic regions, including regions corresponding to lineage-specific gene families.

Advances in long-read sequencing and new assembly algorithms were needed to overcome the challenge of repeats and finish the first complete, telomere-to-telomere (T2T) assembly of a human genome^8,9^. Using these same methods, we recently published six additional pairs of complete sex chromosomes from distinct branches of the ape phylogeny^10^. Although these initial projects targeted haploid chromosomes and required substantial manual curation, improved assembly methods now enable the complete assembly of diploid chromosomes^11,12^. Using these methods, we present here the complete, phased, diploid genomes of six ape species making all data and curated assemblies freely available to the scientific community. We organize the manuscript into three sections focused primarily on 1) finishing the genomes and the development of an ape pangenome, 2) the added value for standard evolutionary analyses, and 3) providing new evolutionary insights into the previously unassembled regions. Although the interior of the ribosomal DNA (rDNA) arrays as well as some small portions of the largest centromeres remain unresolved, these genomes represent an order of magnitude improvement in quality over the prior ape references and are of equivalent quality to the T2T-CHM13 human reference. We propose that these assemblies will serve as the definitive references for all future studies involving human/ape genome evolution.

## RESULTS

### Section I: Ape genome assembly and a pangenome resource

Unlike previous reference genomes that selected female individuals for improved representation of the X chromosome^4–7^, we focused on male samples (**Table 1**) in order to fully represent both sex chromosomes^10^ and provide a complete chromosomal complement for each species. Samples from two of the species, bonobo and gorilla, originated from parent–child trios (**Supplementary Note I**) facilitating phasing of parental haplotypes. For other samples where parental data were not available, deeper Hi-C datasets were used (**Table 1**) to achieve chromosome-scale phasing. For all samples, we prepared high-molecular-weight DNA and generated deep PacBio HiFi (high-fidelity; mean=90-fold sequence coverage) and ONT (Oxford Nanopore Technologies; mean=136.4-fold sequence coverage) sequence data (**Table Assembly S1**). For the latter, we specifically focused on producing at least 30-fold of ultra-long (UL > 100 kbp) ONT sequence data to scaffold assemblies across larger repetitive regions, including centromeres and segmental duplications (SDs). We applied Verkko^11^ (v. 2.0)—a hybrid assembler that leverages the accuracy of HiFi data for generating the backbone of the assembly (**Methods, Supplementary Note II**); UL-ONT sequencing for repeat resolution, local phasing, and scaffolding; and Hi-C or trio data for chromosome-scale phasing of haplotypes into a fully diploid assembly. To serve as a reference genome, a haploid “primary” assembly was selected from the diploid assembly of each species. For the trios, the most accurate and complete haplotype was selected as the primary assembly (maternal vs. paternal), and for the non-trios, the most accurate and complete chromosome was selected for each chromosome pair. With convention, both sex chromosomes and the mitochondrial genome were also included in the primary assembly.

**Table 1:** Summary of ape genome assemblies.

| Sample information |  |  |  | Assembly stats (v2.0)<br>Hap1 (Hap2) or mat (pat) |  |  |  |  |  |
| --- | --- | --- | --- | --- | --- | --- | --- | --- | --- |
| Common name | Scientific name | Tissue | Sex | Accession | Total bases (Gbp) | contig N50 (Mbp) | Number of T2T contigs | Number of non-rDNA gaps or missing telomeres | QV |
| Chimpanzee (PTR) | *Pan troglodytes* | Lymphoblastoid | M | PRJNA916736 | 3.136 (3.030) | 146.29 (140.84) | 19 (17) | 0 (2) | 66.0 |
| Bonobo (PPA)* | *Pan paniscus* | Fibroblast/<br>Lymphoblastoid | M | PRJNA942951 | 3.206 (3.072) | 147.03 (147.48) | 20 (19) | 0 (0) | 62.7 |
| Gorilla (GGO)* | *Gorilla gorilla* | Fibroblast | M | PRJNA942267 | 3.553 (3.350) | 151.43 (150.80) | 19 (22) | 4 (0) | 61.7 |
| Bornean orangutan (PPY) | *Pongo pygmaeus* | Fibroblast | M | PRJNA916742 | 3.164 (3.048) | 140.59 (137.91) | 15 (13) | 1 (2) | 65.8 |
| Sumatran orangutan (PAB) | *Pongo abelii* | Fibroblast | M | PRJNA916743 | 3.168 (3.077) | 146.20 (140.60) | 14 (13) | 2 (3) | 63.3 |
| Siamang (SSY) | *Symphalangus syndactylus* | Lymphoblastoid | M | PRJNA916729 | 3.235 (3.122) | 146.71 (144.67) | 24 (20) | 0 (5) | 66.4 |
| Average |  |  |  | - | 3.244 (3.117) | 146.38 (143.72) | 18.5 (17.3) | 1.2 (2) | 64.3 |

\*Sample with parental sequence data. QV represents the score from Illumina/HiFi hybrid-based approach.

Considering the diploid genomes of each species, 74% (215/290) of all chromosomes are T2T assembled (gapless with telomere on both ends) and at least 80.8% of chromosomes are T2T in at least one haplotype (**Fig. 1**, **Table 1 & AssemblyS2**). Overall, there are an average of six gaps or breaks in assembly contiguity per haplotype (range=1-12), typically localized to the rDNA array, reducing to an average of 1.6 gaps if those acrocentric chromosomes are excluded. All assemblies were curated to extend partially into each rDNA array from both sides, ensuring that no non-rDNA sequence was missed. In addition to gaps, we searched specifically for collapses and misassemblies using dedicated methods (**Table AssemblyS3**, **Methods**). We estimate, on average, 1–2 Mbp of collapse per haplotype assembly and a wider range of potential misassemblies with an average of 0.2 to 11 Mbp flagged per haplotype assembly (**Table AssemblyS3**). Comparison with Illumina data^13^ from the same samples provided a lower-bound accuracy of QV=49.3, limited by Illumina coverage loss in high-GC regions, while comparisons including HiFi data suggest even higher accuracy (QV=61.7; **Table AssemblyS1**). Overall, we estimate 99.2–99.9% of each genome is completely and accurately assembled, including heterochromatin. This is consistent with the T2T-CHM13v1.1 assembly, for which 0.3% of the genome was covered by known issues^13^. In short, these ape diploid genome assemblies represent an advance by at least one order of magnitude in terms of sequence accuracy and contiguity with respect to all prior ape genome assemblies^4–7^. For the first time, the centromeric regions, large blocks of SDs, and subterminal heterochromatin have been fully sequenced and assembled in both haplotypes as well as more subtle improvements genome-wide. For example, a comparison with previous genome assemblies for the same species shows an enrichment in sequence motifs capable of forming non-canonical (non-B) DNA (A-phased, direct, mirror, inverted, and short tandem repeats in particular) in newly gained regions of the new assemblies (**Table AssemblyS10; Supplementary Note III**); such motifs have been shown to be difficult sequencing targets^14^ but are resolved here. Each genome assembly was annotated by NCBI and has been adopted as the main reference in RefSeq, replacing the previous short- or long-read-based, less complete versions of the genomes and updating the sex chromosomes with the newly assembled and polished versions.

**Figure 1.**
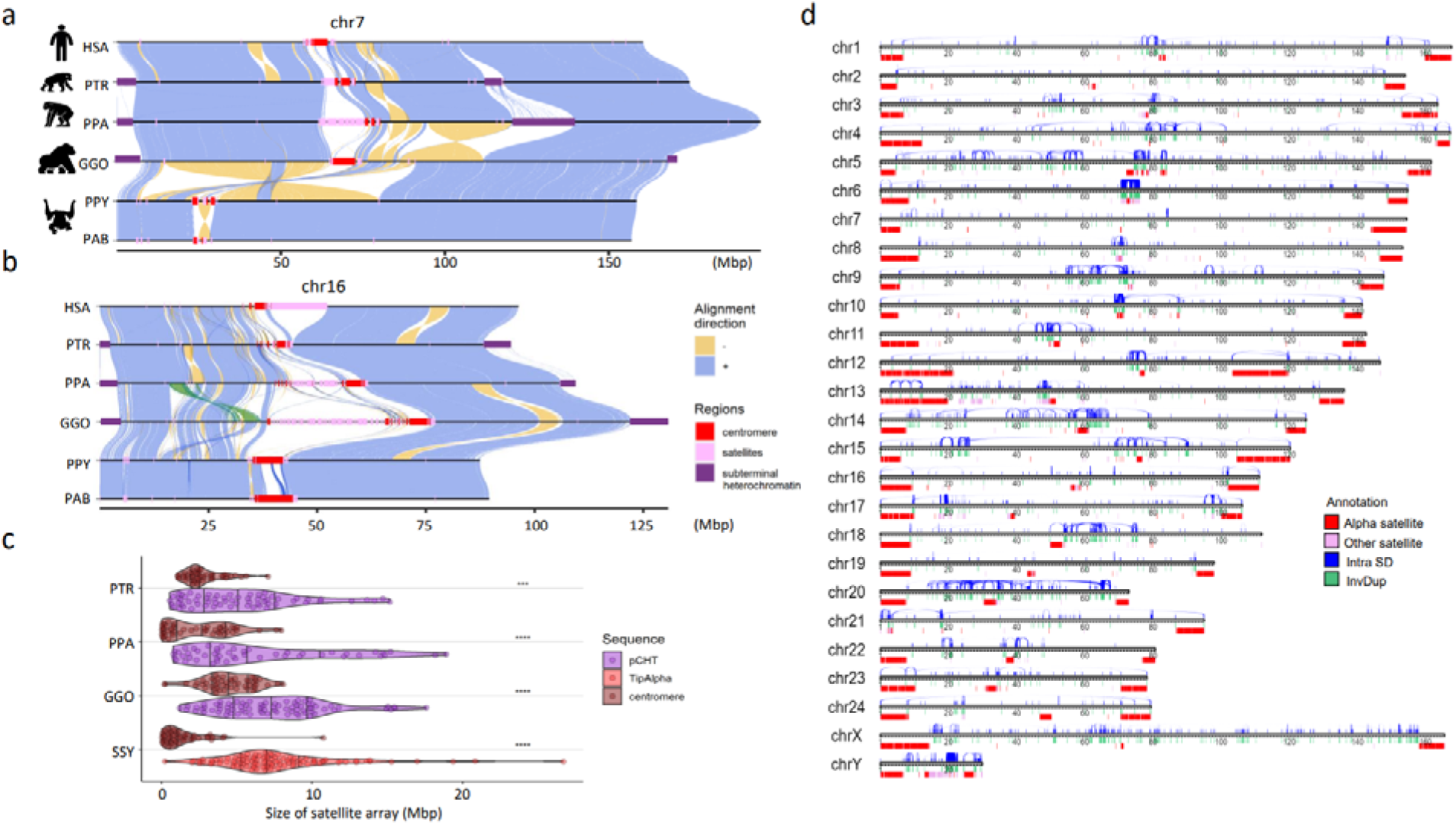
Chromosomal-level assembly of complete great ape genomes. **a)** A comparative ape alignment of human (HSA) chromosome 7 with chimpanzee (PTR), bonobo (PPA), gorilla (GGO), Bornean and Sumatran orangutans (PPY and PAB) shows a simple pericentric inversion in the *Pongo* lineage (PPY and PAB) and **b)** HSA chromosome 16 harboring complex inversions. Each chromosome is compared to the chromosome below in this stacked representation using the tool SVbyEye (https://github.com/daewoooo/SVbyEye). Regions of collinearity and synteny (+/blue) are contrasted with inverted regions (-/yellow) and regions beyond the sensitivity of minimap2 (homology gaps), including centromeres (red), subterminal/interstitial heterochromatin (purple), or other regions of satellite expansion (pink). A single transposition (green in panel b) relocates ∼4.8 Mbp of gene-rich sequence in gorilla from human chromosome 16p13.11 to human chromosome 16p11.2. **c)** Distribution of assembled satellite blocks for centromere (alpha) and subterminal heterochromatin including, African great ape’s pCht or siamang’s (SSY) α-satellite, shows that subterminal heterochromatin are significantly longer in ape species possessing both heterochromatin types (One-sided Wilcoxon ranked sum test; **** *p*< 0.0001; *** *p*< 0.001). **d)** Schematic of the T2T siamang genome highlighting segmental duplications (Intra SDs; blue), inverted duplications (InvDup; green), centromeric, subterminal and interstitial α-satellites (red), and other satellites (pink).

Human and nonhuman primate (NHP) genome assemblies are now comparable in quality, helping to mitigate reference biases in alignment and variant discovery. We employed Progressive Cactus^15^ to construct 7-way (six primary and T2T-CHM13) 8-way (six primary ape and two human haplotypes), and 16-way (diploid ape genomes including and four human haplotypes) reference-free multiple genome alignments (**Supplementary Note IV; Data Availability**). The more complete sequence and representation facilitates ancestral state reconstruction for more genomic regions. For example, we annotated the human–primate ancestral state of the GRCh38 reference genome by applying the parsimony-like method used by the 1000 Genomes Project and Ensembl^16^. We observed a genome-wide increase of 6.25% in the total ancestrally annotated base pairs over the existing Ensembl annotation (release 112), with the greatest autosomal increase for chromosome 19 (21.48%; **Fig. 2a**). We annotated over 18 million base pairs for chromosome Y, which is 4.67 times the annotated base pairs in the Ensembl annotation. Additionally, we find that the T2T annotation has more high-confidence bases in regions where the two annotations disagree most (**Fig.AncestralallelesS1**). We also constructed an interspecies 10-way pangenome representation of the ape genomes by Minigraph-Cactus^17^, using the ape and four human haplotypes (**Supplementary Note IV**). Compared to the recently released human pangenome from 47 individuals^18^, the resulting interspecies graph increases by ∼3-fold the number of edges and nodes, resulting in a 3.38 Gbp ape “pan”genome.

**Figure 2.**
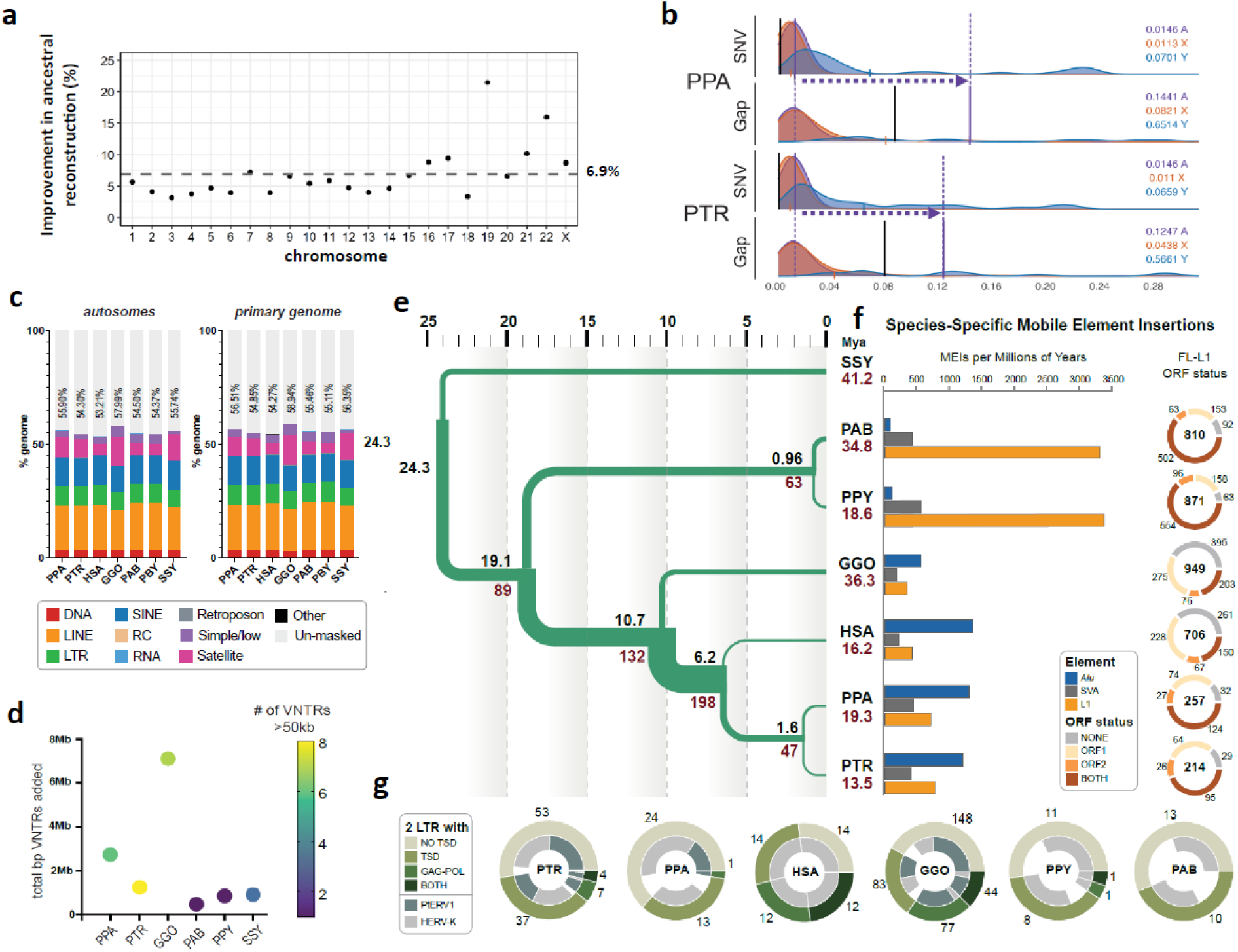
Genome resource improvements. **a)** Improvement in the ancestral allele inference by Cactus alignment over the Ensembl/EPO alignment of the T2T ape genomes. **b)** Genome-wide distribution of 1 Mbp single-nucleotide variant (SNV)/gap divergence between human and bonobo (PPA)/chimpanzee (PTR) genomes. The purple vertical lines represent the median divergence observed. The horizontal dotted arrows highlight the difference in SNV vs. gap divergence. The black vertical lines represent the median of allelic divergence within species. **c)** Total repeat content of ape autosomes and the primary genome including chrX and Y. **d)** Total base pairs of previously unannotated VNTR satellite annotations added per species. The color of each dot indicates the number of newly annotated satellites, out of 159, which account for more than 50 kbp in each assembly. (**Table Repeat S2**). **e)** Demographic inference. Black and red values refers to speciation times and effective population size (Ne), respectively. For Ne, values in inner branches refer to TRAILS estimates, while that of terminal nodes is predicted via msmc2, considering the harmonic mean of the effective population size after the last inferred split. **f)** (Left) Species-specific *Alu*, SVA and L1 MEI counts normalized by millions of years (using speciation times from (2e)). (Right) Species-specific Full-length (FL) L1 ORF status. The inner number within each circle represents the absolute count of species-specific FL L1s. **g)** Species-specific ERV comparison shows that the ERV increase in gorilla and chimpanzee lineages is due primarily to PTERV1 expansions.

As a second approach, we also applied pangenome graph builder (PGGB)^19^ to construct all-to-all pairwise alignments for all 12 human primate haplotypes along with three T2T human haplotypes (T2T-CHM13v2.0 and T2T-HG002v1.0). We used these pairwise alignment data to construct an implicit graph (**Methods**) of all six species and computed a conservation score for every base pair in the genome (**Fig. PanGenomeS1; Methods**). The approach is transitive without a reference bias and considers both assembled haplotypes for each genome, as well as unique and repetitive regions, identifying the most rapidly evolving regions in each primate lineage (**Fig. PanGenomeS1**). We highlight the performance of this implicit graph in some of the most structurally diverse and dynamic regions of our genome, including the major histocompatibility complex (MHC) and the chromosome 8p23.1 inversion (**Fig. PanGenomeS1**).

### Section II. Resource improvement highlights

#### Sequence divergence

The oft-quoted statistic of ∼99% sequence identity between chimpanzee and human holds for most of the genome when considering single-nucleotide variants (SNVs) (**Fig. 2b**). However, comparisons of T2T genomes suggest a much more nuanced estimate. Examining the distribution of 1 Mbp aligned windows shows that the tail of that distribution is much longer with 12.5–27.3% of the genome failing to align or inconsistent with a simple 1-to-1 alignment, especially within centromeres, telomeres, acrocentric regions, and SDs (**Figs. 1 & 2b**). We, therefore, considered SNV divergence separately from “gap” divergence, which considers poorly aligned sequences (**Methods**). Both parameters scale linearly with evolutionary time except for an inflated gorilla gap divergence (both between and within species comparisons) (**Fig. SeqDiv S1 & 2**). Gap divergence shows a 5- to 15-fold difference in the number of affected Mbp when compared to SNVs due to rapidly evolving and structural variant regions of the genome— most of which can now be fully accessed but not reliably aligned. As part of this effort, we also sequenced and assembled two pairs of closely related, congeneric ape species. For example, the Sumatran and Bornean orangutan species (the latter genome has not been sequenced previously) are the most closely related ape species, estimated to have diverged ∼0.5–2 million years ago (mya)^20–22^. The autosome sequence identity of alignable bases between these two closely related orangutan genomes was 99.5% while the gap divergence was ∼8.9% (autosomes). These numbers are highly consistent with analyses performed using alternative alignment approaches (**Table SeqDiv. S1 & S2, Table OrangSeqDivS3; Supplementary Note V**).

#### Speciation time and incomplete lineage sorting (ILS)

To jointly estimate speciation times (the minimum time at which two sequences can coalesce) and ancestral effective population sizes (Ne), we modeled ILS across the ape species tree (**Table.ILS.S1**)^23^. Among the great apes (human, chimpanzee, gorilla and orangutan), our analyses date the human–chimpanzee split at 5.5–6.3 mya, the African ape split at 10.6–10.9 mya, and the orangutan split at 18.2–19.6 mya (**Fig. 2e**). We infer ILS for an average of 39.5% of the autosomal genome and 24% of the X chromosome, representing an increase of approximately 7.5% compared to recent reports from less complete genomes^24^ in part due to inclusion of more repetitive DNA (**Fig.ILS.S1**). We estimate that the human–chimpanzee–bonobo ancestral population (average Ne=198,000) is larger than that of the human–chimpanzee–gorilla ancestor (Ne=132,000), suggesting an increase of the ancestral population 6–10 mya. In contrast, the effective population sizes of more terminal branches are estimated to be smaller. For example, we estimate it is much smaller (Ne=46,800) in the *Pan* ancestor at 1.7 mya and *Pongo* ancestor (Ne=63,000) at 0.93 mya, though these estimates should be taken with caution. For each terminal species branch, we infer the population size to range from 13,500 (*Pan troglodytes*) to 41,200 (*Symphalangus syndactylus*) (**Methods**). Compared to the autosomes, we find reduced X chromosome diversity for the African ape ancestor (Ne=115,600, X-to-A ratio of 0.87), and, more strongly, for the human–chimpanzee– bonobo ancestor (Ne=76,700, X-to-A ratio of 0.42). We additionally reconstruct a high-resolution, time-resolved ILS map (**Fig.ILS.S2**). T2T genomes support relatively high ILS estimates in previously inaccessible genomic regions, such as those encompassing the HLA genes (**Fig.ILS.S3**). Furthermore, multiple haplotypes for several species can also reveal cases of ancient polymorphism that have been sustained for thousands of years until present-day genomes, reflected in genomic regions with differential ILS patterns that depend on which combination of haplotypes are analyzed (**Fig.ILS.S3**).

#### Gene annotation

We applied two gene annotation pipelines (CAT and NCBI) to identify both protein-coding and noncoding RNA (ncRNA) genes for the primary assembly for each NHP. We complemented the annotation pipelines by direct mapping of Iso-Seq (50 Gbp of full-length non-chimeric [FLNC] cDNA) generated from each sample and searching for multi-exon transcripts. The number of protein-coding genes is very similar among different apes (Ne=22,114–23,735) with a little over a thousand genes predicted to be gained/duplicated or lost specifically per lineage (**Table. GeneS1**). Using the UCSC gene set, based on GENCODE^25^, we estimate that 99.0–99.6% of corresponding human genes are now represented with >90% of genes being full-length. We identify a fraction (3.3–6.4%) of protein-coding genes present in the NHP T2T genomes that are absent in the human annotation set used. This includes 770–1,482 novel gene copies corresponding to 315–528 families in the NHPs with ∼68.6% corresponding to lineage-specific SDs, all supported by Iso-Seq transcripts (**Table. GeneS1, S2**). In addition, 2.1%–5.2% of transcripts show novel NHP splice forms once again supported by Iso-Seq data (**Table. GeneS1**). We provide a unique resource in the form of a curated consensus protein-coding gene annotation set by integrating both the NCBI and CAT pipelines (**Methods**). Finally, we analyzed FLNC reads obtained from testis from a second individual^10^ to quantify the potential impact genome-wide on gene annotation and observed improvements in mappability, completeness, and accuracy (**Fig. Gene.S3 and Supplementary Note VII**). In gorilla, for example, we mapped 28,925 (0.7%) additional reads to the T2T assembly in contrast to only 171 additional reads to the previous long-read assembly^5^. Similarly, we observed 33,032 (0.7%) soft-clipped reads (>200 bp) in the gorilla T2T assembly in contrast to 89,498 (2%) soft-clipped reads when mapping to the previous assembly^5^. These improvements in mappability are non-uniformly distributed with loci at centromeric, telomeric, and SD regions, leading to increased copy number counts when compared to previous genome assemblies (**Fig. Gene.S3e-g**).

#### Repeat annotation and mobile element insertion (MEI) identification

Based on RepeatMasker annotations (Dfam 3.7) and extensive manual curation^26^ (**Methods; Supplementary Note VIII**), we generated a near-complete census of all high-copy repeats and their distribution across the ape genomes (**Table Repeat.S1; Extended data Table 2**). We now estimate that the autosomes of the ape genomes contain 53.21–57.99% detectable repeats, which include transposable elements (TEs), various classes of satellite DNA, variable number tandem repeats (VNTRs), and other repeats (**Fig. 2c**), significantly lower than the sex chromosomes (X [61.79–66.31%] and Y [71.14–85.94%])^10^. Gorilla, chimpanzee, bonobo, and siamang genomes show substantially higher satellite content driven in large part by the accumulation of subterminal heterochromatin through lineage-specific satellite and VNTR expansions (**Fig. 1**, **Fig. 2d, Extended data Table 2**). Satellites account for the largest repeat variation (**Extended data Table 2**), ranging from 4.94% satellite content in Bornean orangutan (159.2 Mbp total) to 13.04% in gorilla (462.50 Mbp total). Analyzing gaps in exon and repeat annotations led to the identification of 159 previously unknown satellite monomers (**Table Repeat S2-S9**), ranging from 0.474 to 7.1 Mbp in additional base pairs classified per genome (**Fig. 2d**). Of note, 3.8 Mbp of sequence in the gorilla genome consists of a ∼36 bp repeat, herein named VNTR_148, accounting for only 841.9 kbp and 55.9 kbp in bonobo and chimpanzee, respectively (**Fig. 2d**). This repeat displays a pattern of expansion similar to that of the unrelated repeat pCht/subterminal satellite (StSat)^10^, suggesting it may have undergone expansion via a similar mechanism.

We find that 40.74% (gorilla) to 45.81% (Bornean orangutan) of genomes correspond to TEs (**Extended data Table 2; Table Repeat S1**). Leveraging the unaligned sequences in a 7-way Cactus alignment, we define a comprehensive set of both truncated and full-length, species-specific LINE, *Alu*, ERV, and SVA insertions for each ape species (**Table Repeat S12**). Orangutans appear to have the highest L1 mobilization rate based both on absolute number of insertions and the number of full-length elements with intact open reading frames (ORFs), while the African apes (gorilla, chimpanzee, bonobo, and human) show a higher accumulation of Alu insertions (**Fig. 2f, Supp Fig. Species Specific MEI 1**). The number of L1s with intact ORFs varies by a factor of 5.83, with chimpanzee having the lowest (95) and orangutans having the highest with at least 2.5 times more L1s with intact ORFs (more than 500 in both orangutan species compared to 203 in gorilla). Humans and gorillas fall in between this spectrum. The overall number and high percentage of full-length L1 elements with intact ORFs in orangutans suggests recent high L1 activity. *Alu* activity is shown to be quiescent in the orangutans, consistent with previous reports^27^, suggesting a genome environment where L1s out-compete *Alu* retrotransposons. When considering only full-length ERV elements with both target site duplications, and *gag* (capsid) and *pol* (reverse transcriptase and integrase) coding domains, a striking difference is observed with higher full-length, species-specific ERV content in gorilla (44), followed by human with 12, and chimpanzee with only three. PtERV and HERVK account for the ERVs with both target site duplications and protein domains, along with more degraded ERVs in gorilla, human, chimpanzee, and bonobo (**Fig. 2g; Table Repeat S13**). In addition to MEIs, we also characterized the distribution of integrated NUMTs (nuclear sequences of mitochondrial DNA origin) in ape genomes (**Methods**). We observe a substantial gain in the number (3.7-10.5%) and total length of NUMTs (6.2-30%) (**Table Repeat S10**) over non-T2T assemblies, with the largest gain observed for bonobo; Sumatran and Bornean orangutan species differ in NUMT content by 73,990 bp despite their recent divergence.

**Extended data Table 2:** Overview of repeat content (Mbp and percentage) in the ape genomes.

|  | Human |  | Chimpanzee |  | Bonobo |  | Gorilla |  | Sumatran orangutan |  | Bornean orangutan |  | Siamang |  |
| --- | --- | --- | --- | --- | --- | --- | --- | --- | --- | --- | --- | --- | --- | --- |
|  | Mbp | % | Mbp | % | Mbp | % | Mbp | % | Mbp | % | Mbp | % | Mbp | % |
| **DNA** | 108.59 | 3.48 | 110.60 | 3.48 | 110.74 | 3.47 | 111.82 | 3.15 | 111.20 | 3.41 | 110.83 | 3.44 | 106.76 | 3.27 |
| **LINE** | 633.55 | 20.33 | 636.62 | 20.04 | 638.42 | 19.99 | 649.75 | 18.33 | 686.42 | 21.06 | 683.56 | 21.22 | 631.83 | 19.37 |
| **PLE** | 0.06 | 0.00 | 0.07 | 0.00 | 0.07 | 0.00 | 0.07 | 0.00 | 0.07 | 0.00 | 0.07 | 0.00 | 0.07 | 0.00 |
| **LTR** | 272.91 | 8.75 | 274.90 | 8.65 | 275.67 | 8.63 | 277.85 | 7.84 | 279.36 | 8.57 | 278.28 | 8.64 | 256.49 | 7.86 |
| **SINE** | 393.51 | 12.62 | 391.01 | 12.30 | 393.08 | 12.3 | 397.19 | 11.20 | 396.85 | 12.17 | 395.47 | 12.28 | 402.73 | 12.34 |
| **RC** | 0.45 | 0.01 | 0.46 | 0.01 | 0.46 | 0.01 | 0.46 | 0.01 | 0.46 | 0.01 | 0.46 | 0.01 | 0.44 | 0.01 |
| **Retroposon** | 4.31 | 0.14 | 4.78 | 0.15 | 4.90 | 0.15 | 5.21 | 0.15 | 3.87 | 0.12 | 4.39 | 0.14 | 6.66 | 0.20 |
| **Satellite** | 161.77 | 5.19 | 251.68 | 7.92 | 270.19 | 8.46 | 462.50 | 13.04 | 191.45 | 5.87 | 159.20 | 4.94 | 376.70 | 11.55 |
| **Simple/low** | 108.38 | 3.48 | 69.17 | 2.18 | 107.68 | 3.37 | 181.36 | 5.11 | 134.09 | 4.11 | 138.98 | 4.31 | 54.29 | 1.66 |
| **Other** | 5.34 | 0.17 | 2.21 | 0.07 | 2.61 | 0.08 | 2.15 | 0.06 | 2.23 | 0.07 | 2.35 | 0.07 | 1.34 | 0.04 |
| **RNA** | 2.78 | 0.09 | 1.55 | 0.05 | 1.53 | 0.05 | 1.38 | 0.04 | 1.89 | 0.06 | 1.59 | 0.05 | 1.36 | 0.04 |
| **Un-masked** | 1,434.70 | 46.02 | 1,434.70 | 45.15 | 1,389.24 | 43.49 | 1,456.09 | 41.06 | 1,451.97 | 44.54 | 1,445.75 | 44.89 | 1,424.20 | 43.65 |
| **Total masked** | 1,691.65 | 54.27 | 1,743.04 | 54.85 | 1,805.35 | 56.51 | 2,089.74 | 58.94 | 1,807.88 | 55.46 | 1,775.19 | 55.11 | 1,838.69 | 56.35 |
| **NUMT** | 0.64 | 0.021 | 0.79 | 0.025 | 0.87 | 0.027 | 0.63 | 0.018 | 0.76 | 0.024 | 0.84 | 0.026 | 0.52 | 0.016 |

#### Selection and diversity

Using short-read whole-genome sequencing data generated from the great ape genetic diversity project^28^ (**Supplementary Note IX**) and mapped to the T2T genomes, we searched for signatures of adaptation by identifying regions of hard^29^ and soft (partial)^30^ selective sweeps in 10 great ape subspecies (**Methods**). Across all taxa, we identify 143 and 86 candidate regions for hard and partial selective sweeps, respectively, with only two overlapping (**Table Selection S1**). Approximately 50% of hard (75/143) and 80% of partial selective (70/86) sweeps are novel and a total of 43 regions overlap with sweeps previously found in humans^31^. As expected, pathways related to diet (sensory perception for bitter taste, lipid metabolism, and iron transport), immune function (antigen/peptide processing, MHC-I binding–strongest signal for balancing selection), cellular activity, and oxidoreductase activity were enriched among bonobos, central and eastern chimpanzees, and western lowland gorillas. While some of these findings are confirmatory, the updated analysis provides remarkable precision. For example, among the well-documented bitter taste receptor targets of selection^32^, we detect significant enrichment in selection signals for such genes in bonobos (*TAS2R3, TAS2R4,* and *TAS2R5*) and western lowland gorillas (*TAS2R14, TAS2R20,* and *TAS2R50*), as well as identified a bitter taste receptor gene (*TAS2R42*) within a sweep region in eastern chimpanzees. Within the chimpanzee lineage, it is notable that hard sweep regions in both eastern and central chimpanzees show significantly greater differentiation (FST = 0.21 and 0.15, Mann-Whitney p < 0.001) when compared to the genome-wide average (FST = 0.09). One of these regions was enriched (**Table Selection S3**) for genes associated with epidermal differentiation (*KDF1* and *SFN*).

#### Immunoglobulin and major histocompatibility complex (MHC) loci

Complete ape genomes make it possible to investigate more thoroughly structurally complex regions known to have a high biomedical relevance, especially with respect to human disease. Importantly, four of the primate genomes sequenced and assembled here are derived from fibroblast (bonobo, gorilla and two orangutans as well as the human T2T reference) instead of lymphoblastoid cell lines. The latter, in particular, has been the most common source of most previous ape genome assemblies limiting characterization of loci subject to somatic rearrangement (e.g., VDJ genes)^33^. Thus, we specifically focused on nine regions associated with the immune response or antigen presentation that are subjected to complex mutational processes or selective forces.

##### Immunoglobulin and T-cell receptor loci

Antibodies, B-cell receptors, and T-cell receptors mediate interactions with both foreign and self-antigens and are encoded by large, expanded gene families that undergo rapid diversification both within and between species^34,35^. We conducted a comparative analyses of the immunoglobulin heavy chain (IGH), light chain kappa (IGK), and lambda (IGL) as well as T-cell receptor alpha (TRA), beta (TRB), gamma (TRG), and delta (TRD) loci in four ape species (**Supplementary Note X**) for which two complete intact haplotypes were constructed (**Fig. 3a, Fig.IG.S1a**). With respect to genes, we identify an average of 60 (IGHV), 36 (IGKV), 33 (IGLV), 46 (TRAV/TRDV), 54 (TRBV), and 8 (TRGV) putatively functional IG/TR V genes per parental haplotype per species across the seven loci (**Fig. 3a, Fig.IG.S1a**); and provide an expanded set of curated IG/TR V, D, and J sequences for each species, including ORF genes (**Table.IG.S1 and Table.IG.S2**). The ape IG genes cluster into phylogenetic subfamilies similar to human (**Fig. IG.S1b**) but there are large structural differences between haplotypes within and between species, accounting for as much as 33% of inter-haplotype length differences in IG and up to 10% in the TR loci (**Fig.IG.S2ab**). IG loci show the most pronounced differences, including large structural changes and a 1.4 Mbp inversion distinguishing the two IGL haplotypes of bonobo (**Fig. 3a; Fig. IG.S2cd**). These large-scale differences frequently correspond to ape-specific genes (those that comprise phylogenetically distinct clades exclusive of human genes) (**Fig. 3a; Fig. IG.S2e**). We observe the greatest number of ape-specific genes within IGH (**Fig. 3a; Fig.IG.S2f**), where we note a greater density of SDs longer than 10 kbp relative to the other six loci (**Fig. IG.S2.g**).

**Figure 3.**
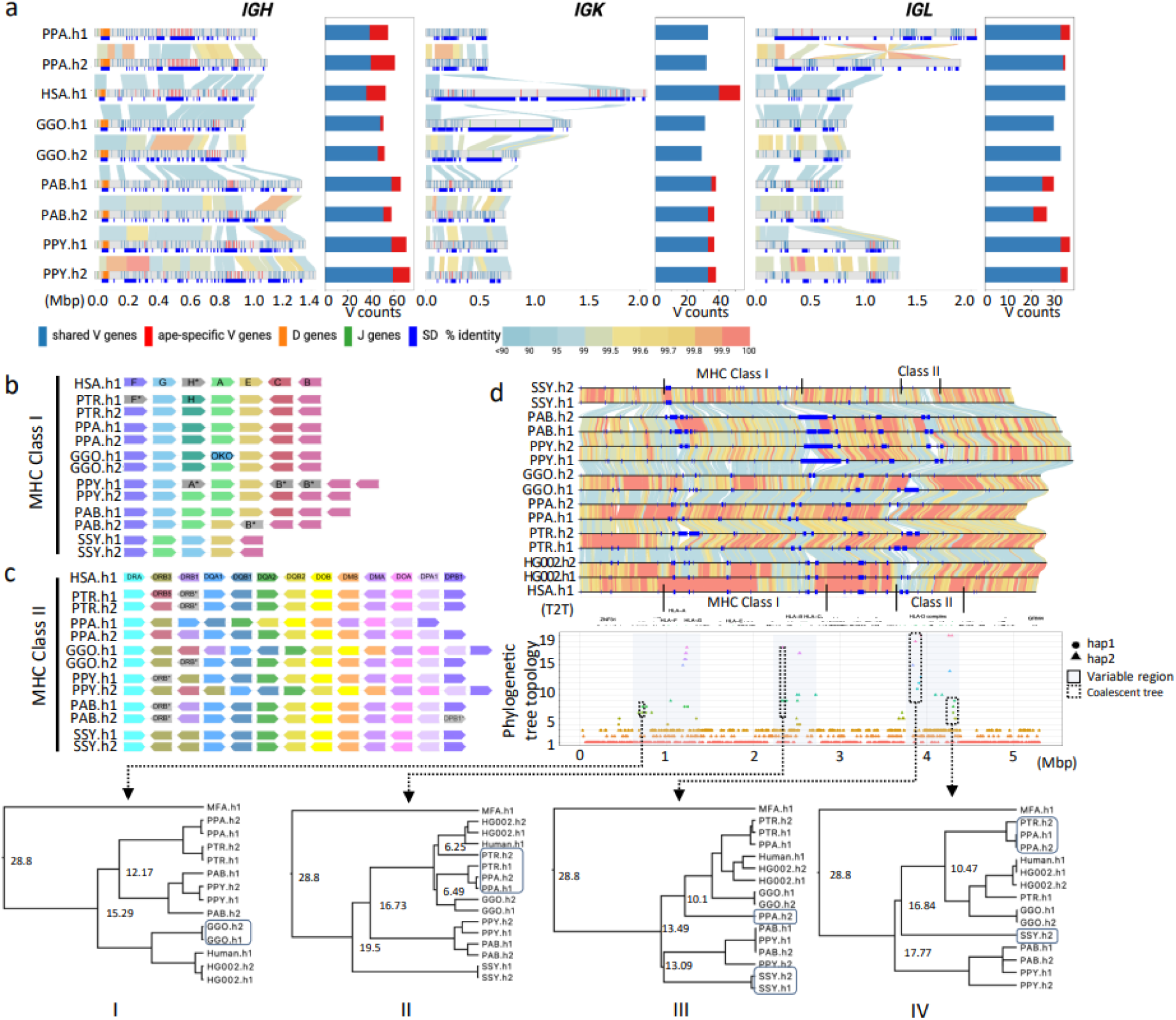
IG and MHC genome organization in apes. a) Annotated haplotypes of *IGH*, *IGK* and *IGL* loci across four primate species and one human haplotype (HSA.h1 or T2T-CHM13). Each haplotype is shown as a line in the genome diagram where the top part shows positions of shared V genes (blue), ape-specific V genes (red), D genes (orange), and J genes (green) and the bottom part shows segmental duplications (SDs) that were computed for a haplotype pair of the same species and depicted as dark blue rectangles. Human SDs were computed with respect to the GRCh38.p14 reference. Alignments between pairs in haplotypes are shown as links colored according to their percent identity values: from blue (<90%) through yellow (99.5%) to red (100%). The bar plot on the right from each genome diagram shows counts of shared and ape-specific V genes in each haplotype. b and c) show schematic representation of MHC locus organization for MHC-I and MHC-II genes, respectively, across the six ape haplotypes (PTR.h1/h2, PPA.h1/h2, GGO.h1/h2, PPY.h1/h2, PAB.h1/h2, SSY.h1/2) and human (HSA.h1). Only orthologs of functional human HLA genes are shown. Loci naming in apes follows human HLA gene names (HSA.h1), and orthologs are represented in unique colors across haplotypes and species. Orthologous genes that lack a functional coding sequence are grayed out and their name marked with an asterisk. One human HLA class I pseudogene (HLA-H) is shown, because functional orthologs of this gene were identified in some apes. **d)** Pairwise alignment of the 5.31 Mbp MHC region in the genome, with human gene annotations and MHC-I and MHC-II clusters. Below is the variation in phylogenetic tree topologies according to the position in the alignment. The x-axis is the relative coordinate for the MHC region and the y-axis shows topology categories for the trees constructed. The three prominent sub-regions with highly discordant topologies are shown through shaded boxes. Four sub-regions (1-4) used to calculate coalescence times are shown with dashed boxes.

##### MHC loci

We also completely assembled and annotated 12 ape haplotypes corresponding to the 4–5 Mbp MHC region (**Supplementary Note XI**). The loci encode diverse cell surface proteins crucial for antigen presentation and adaptive immunity^36^, are highly polymorphic among mammals^37^, and are strongly implicated in human disease via genome-wide association^38^. Comparative sequence analyses confirm extraordinary sequence divergence and structural variation (an average of 328 kbp deletions and 422 kbp insertions in apes compared to human), including duplications ranging from 99.3 kbp in siamang to 701 kbp in the Sumatran orangutan h2 (**Table MHC.S1-2**), as well as contractions and expansions associated with specific MHC genes (**Fig. 3b-c**). Overall, MHC class I genes show greater structural variation within and among the apes than MHC class II genes (**Fig. 3b-c**) with threefold greater average duplication sequences per haplotype (171 kbp vs. 62 kbp). Particularly high divergence in this region is seen in the siamang, which lacks a Sysy-C locus and exhibits an inversion between the MHC-G and MHC-A loci compared to the great apes (**Fig. 3b, Fig. MHC.S1-S8**). While MHC I gene content and organization is nearly identical in human, bonobo, and chimpanzee, other apes show much more variation, including additional genes such as Gogo-OKO, related but distinct from Gogo-A (**Fig. 3b**)^37^. We observe expansion of MHC-A and MHC-B genes in both orangutan species (**Fig. 3b, Fig. MHC.S6-S7**), with MHC-B being duplicated in both haplotypes of the two orangutan species while the MHC-A locus is only duplicated on one haplotype of each species. Similarly, both orangutan species show copy-number-variation of MHC-C, lacking on one haplotype but retaining it on the other (**Fig. 3b, Fig. MHC.S6-S7, Table MHC.S1**). All apes have a nearly identical set of MHC II loci with the exception of the *DRB* locus, which is known to exhibit copy-number-variation in humans^39^, and here shows the same pattern among the apes (**Fig. 3c, Fig. MHC.S1-S8, Table MHC.S2**). We also observe two cases where an MHC locus is present as a functional gene on one haplotype and as a pseudogene on the other haplotype (e.g., Gogo-DQA2 locus in gorilla and the Poab-DPB1 locus in Sumatran orangutan). Overall, this observed variation in MHC gene organization is consistent with long-term balancing selection^39^.

Given the deep coalescence of the HLA locus^40^, we performed a phylogenetic analysis with the complete ape sequences. We successfully constructed 1,906 trees encompassing 76% of the MHC region from the six ape species (**Fig. 3d**). We identify 19 distinct topologies (**Methods**) with three representing 96% (1,830/1,906) of the region and generally consistent with the species tree and predominant ILS patterns. The remaining 4% are discordant topologies that cluster within 200–500 kbp regions (**Table.MHC.S1**) corresponding to MHC I and II genes. We estimate coalescence times of these exceptional regions ranging from 10–24 mya (**Fig. 3d**). Finally, we performed genome-wide tests of selection as described above. We find that selection signatures and nucleotide diversity in the MHC region are among the top 0.1% genome-wide. These signatures confirm long-term balancing selection on MHC in multiple great ape lineages, including central and eastern chimpanzees, as well as at least two regions in MHC consistent with positive selection in bonobos and western chimpanzees^40^.

#### Epigenetic features

Using the T2T genomes, we also created a first-generation, multiscale epigenomic map of the apes, including DNA methylation, 3D chromatin organization, and DNA replication timing (**Supplementary Note XII-XIII**). The long-read sequencing data from individual ape species, for example, allowed us to construct a comparative map of 5-methylcytosine (5mC) DNA signatures for each ape genome sequenced here. We distinguish hypomethylated and hypermethylated promoters associated with gene expression and demonstrate that in each cell type, the majority (∼83%) of promoters are consistently methylated (8,174 orthologous ape genes assessed) (**Table. MET.S1-2**). Specifically, we identify 1,997 differentially methylated promoters (1,382 for fibroblast and 1,381 lymphoblast cell lines samples) as candidates for gene expression differences among the species (**Table. MET.S1-2**). Consistently methylated promoters were more lowly methylated, more highly expressed, and had a higher density of CpG sites compared to variably methylated promoters (P<10-16 two-sided Mann–Whitney U test, **Fig. MET.S1**). These results highlight the interactions between sequence evolution and DNA methylome evolution with consequences on gene expression in ape genomes^41,42^. Additionally, we mapped Repli-seq, including previously collected NHP datasets^43^, to investigate evolutionary patterns of replication timing. We identified 20 states with different patterns of replication timing. Overall, the replication timing program is largely conserved, with 53.1% of the genome showing conserved early and late replication timing across primates, while the remaining regions exhibit lineage-specific patterns (**Fig. RT.S1**) such as the very late replication pattern associated with heterochromatic caps in gorilla and chimpanzee. We inspected replication timing of SDs and found unique patterns for each type of lineage-specific SD, as shown in **Fig. RT.S2**.

#### Evolutionary rearrangements and serial ape inversions

Yunis and Prakash (1982)^44^ originally identified 26 large-scale chromosomal rearrangements distinguishing humans and other great ape karyotypes, including translocation of gorilla chromosomes 4 and 19, chromosome 2 fusion in human, and 24 peri- and paracentric inversions. We completely sequenced and analyzed 43 of the breakpoints associated with these chromosomal rearrangements, with variable length resolutions (average=350 kbp with a maximum of ∼700 kbp from previous cytogenetic mapping; **Table INV.S1 & Fig. INV.S1; Supplementary Note XIV**). These include six cases where the boundaries involving the centromere and/or the telomere are now fully resolved and additional cases where a more complex series of structural changes are suggested (**Fig. 4; Fig.INV.S1**). As an example, for human chromosome 3^45^ we discovered an additional evolutionary rearrangement and inferred the occurrence of an evolutionary new centromere in the orangutan lineage (**Fig. 4e**). This increases the number of new centromere seedings to 10, making this chromosome a hotspot for neocentromeres in humans (15 documented cases in humans^46,47^). The next highest is chromosome 11, which has only four such events^48^.

**Figure 4.**
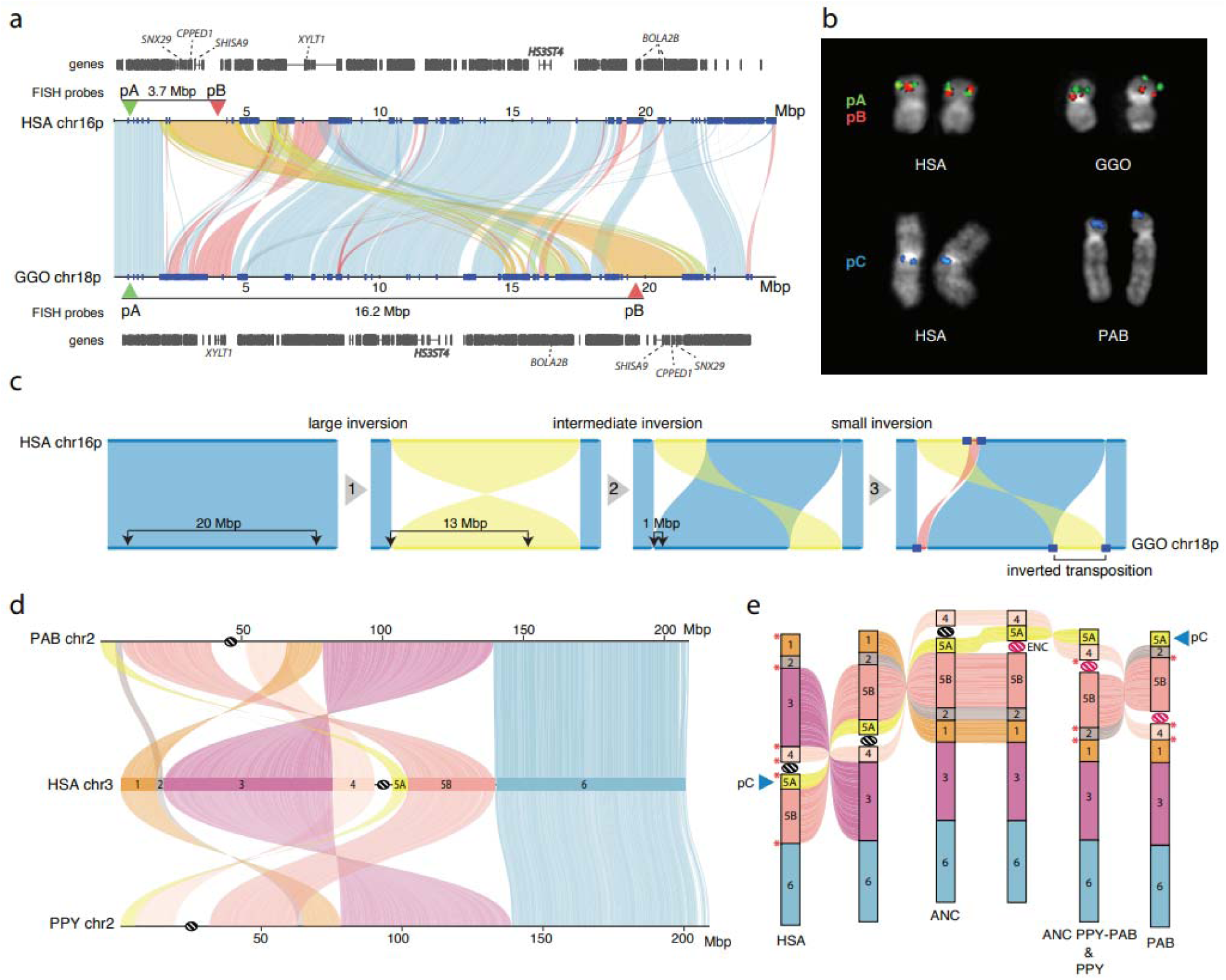
Great ape inversions and evolutionary rearrangements. **a)** Alignment plot of gorilla chr18p and human chr16p shows a 4.8 Mbp inverted transposition (yellow). SDs are shown with blue rectangles. **b)** Experimental validation of the gorilla chr18 inverted transposition using FISH with probes pA (CH276-36H14) and pB (CH276-520C10), which are overlapping in human metaphase chromosomes. The transposition moves the red pB probe further away from the green pA probe in gorilla, resulting in two distinct signals. FISH on metaphase chromosomes using probe pC (RP11-481M14) confirms the location of a novel inversion to the p-ter of PAB chr2. **c)** An evolutionary model for the generation of the inverted transposition by a series of inversions mediated by SDs. **d)** Alignment plot of orangutan chromosome 2 homologs to human chromosome 3 highlights a more complex organization than previously known by cytogenetics^45^: a novel inversion of block 5A is mapping at the p-ter of both chr2 in PAB and PPY. **e)** A model of serial inversions requires three inversions and one centromere repositioning event (evolutionary neocentromere; ENC) to create PPY chromosome 2, and four inversions and one ENC for PAB. Red asterisks show the location of SDs mapping at the seven out of eight inversion breakpoints.

During the finishing of the ape genomes, particularly the SDs flanking chromosomal evolutionary rearrangements, we noted several hundred smaller inversions and performed a detailed manual curation to catalog both homozygous and heterozygous events. Focusing on events larger than 10 kbp, we curate 1,140 interspecific inversions—522 are newly discovered^7,20,44,48–68^ (**Table INV.S2**); 632 of the events are homozygous (found in both the assembled ape haplotypes) with remainder present in only one of the two ape haplotypes and, thus, likely polymorphisms. We also refine the breakpoints of 85/618 known inversions and identify several events that appeared to be the result of serial inversion events. In particular, we identify a 4.8 Mbp fixed inverted transposition on chromosome 18 in gorilla (**Fig. 4a-c**) that was incorrectly classified as a simple inversion but more likely to be explained by three consecutive inversions specific to the gorilla lineage transposing this gene-rich segment to 12.5 Mbp downstream (**Fig. 4a-c**). Similarly, the complex organization of orangutan chr2 can be explained through a model of serial inversions requiring three inversions and one centromere repositioning event (evolutionary neocentromere; ENC) to create PPY chromosome 2, and four inversions and one ENC for PAB (**Fig. 4d,e**). SDs map to seven out of eight inversion breakpoints. In total, 416 inversions have an annotated gene mapping at least one of the breakpoints with 724 apparently devoid of protein-coding genes (**Table INV.S2**). Of these inversions, 63.5% (724/1140) have annotated human SDs at one or both ends of the inversion representing a significant 4.1-fold enrichment (*p*<0.001). The strongest predictable signal was for inverted SDs mapping to the breakpoints (6.2-fold; *p*<0.001) suggesting non-allelic homologous recombination driving many of these events. We also observed significant enrichment of novel transcripts (**Table Gene.S2**) at the breakpoints of the inversions of African great apes (*p* < 0.036). Finally, we assigned parsimoniously >64% of homozygous inversions to the ape phylogeny (**Fig. INV.S3**) with the remaining inversions predicted to be recurrent. The number of inversions generally correlates with evolutionary distance (r^2^=0.77) with the greatest number assigned to the siamang lineage (*n*=44). However, the human lineage shows fivefold less than that expected based on branch length and the trend still holds when using the Bornean orangutan as a reference instead of human.

#### Structurally divergent and accelerated regions of mutation

Previous studies have pinpointed rapidly evolving regions associated with genes under positive selection^69^ or cis-regulatory elements (CREs) undergoing functional changes^70^. We utilized three strategies to systematically assess regions of accelerated mutation. First, was a bottom-up mutation-counting approach that identifies windows of ancestor quickly evolved regions (AQERs) based on sequence divergence^70^ (**Methods; Supplementary Note XV**). We identified 14,210 AQER sites (**Table AQER**) across four primate lineages, including 3,268 on the human branch (i.e., HAQERs). Our analysis more than doubles the number of HAQERs identified from previous gapped primate assemblies (*n*=1,581) (**Fig. 5a, Fig. AQER.S1**). Such elements are highly enriched in repetitive DNA, though not universally. With respect to MEIs, AQERs are depleted in SINEs, but enriched within the VNTRs of hominin-specific SVA elements (**Fig. 5b**). Additionally, HAQERs also exhibit a significant enrichment for bivalent chromatin states (repressing and activating epigenetic marks) across diverse tissues, with the strongest enrichment being for the bivalent promoter state (*p*<1e-35) (**Fig. 5a**; **Table AQER.S1**)—a signal not observed among other apes likely due to the chromatin states being called from human cells and tissues (**Supplementary Note**). An example of a human-specific HAQER change includes an exon and a potential CRE in the gene *ADCYAP1*, in the layer 5 extratelencephalic neurons of primary motor cortex. This gene shows convergent downregulation in human speech motor cortex and the analogous songbird vocal learning layer 5 type extratelencephalic neurons necessary for speech and song production^71,72^. We find here downregulation in layer 5 neurons of humans relative to macaques (RNA-seq) and an associated unique human epigenetic signature (hyper methylation and decreased ATAC-Seq) in the middle HAQER region of the gene that is not observed in the same type of neurons of macaque, marmoset, or mouse (**Fig. 5c, Fig. AQER.S2-S3**).

**Figure 5.**
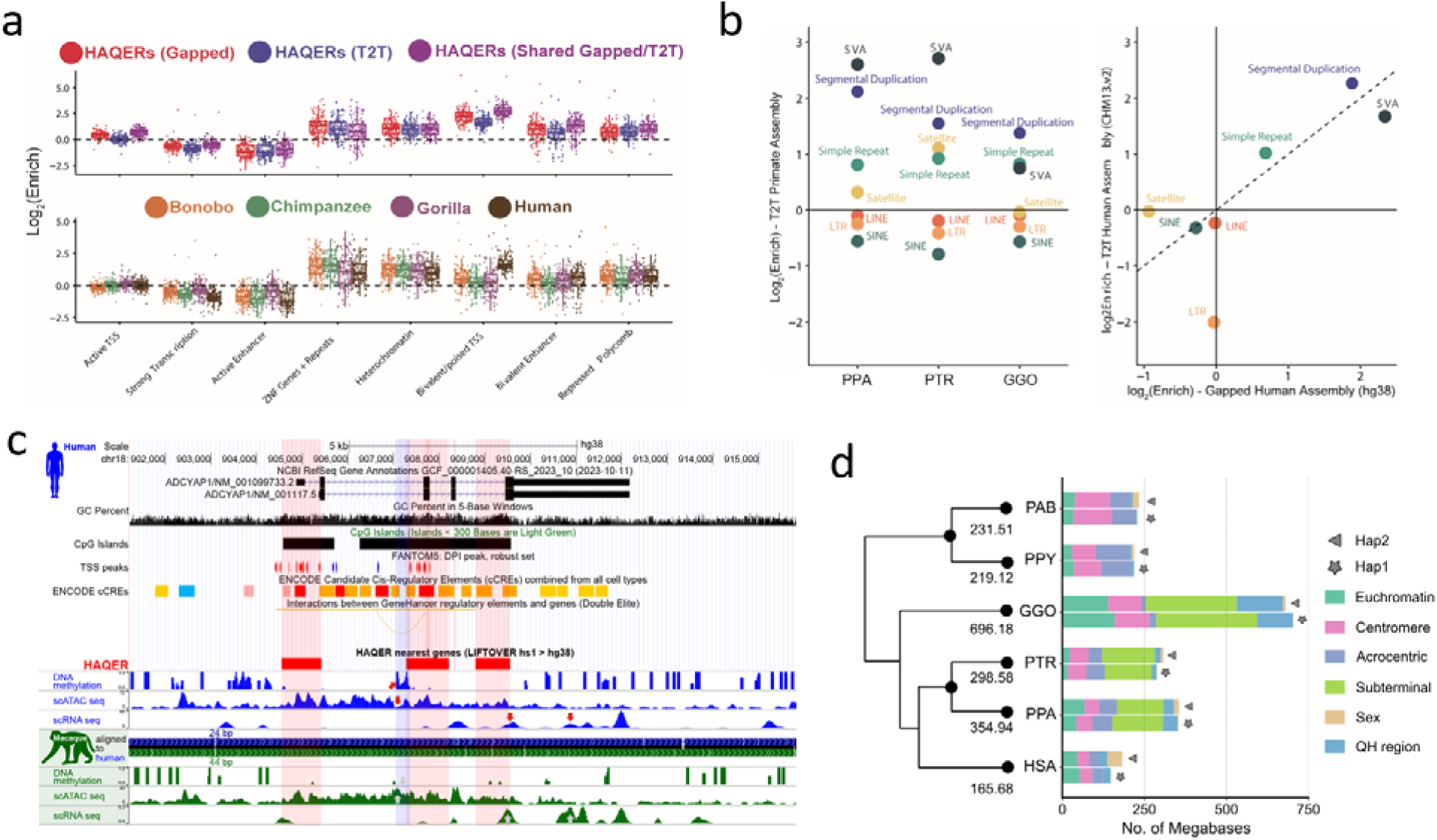
Divergent regions of the ape genomes. **a)** HAQER (human ancestor quickly evolved region) sets identified in gapped (GRCh38) and T2T assemblies show enrichments for bivalent gene regulatory elements across 127 cell types and tissues, with the strongest enrichment observed in the set of HAQERs shared between the two analyses (top). The tendency for HAQERs to occur in bivalent regulatory elements (defined using human cells and tissues) is not present in the sets of bonobo, chimpanzee, or gorilla AQERs (ancestor quickly evolved regions; bottom). **b)** AQERs are enriched in SVAs, simple repeats, and SDs, but not across the general classes of SINEs, LINEs, and LTRs (left). With T2T genomes, the set of HAQERs defined using gapped genome assemblies became even more enriched for simple repeats and SDs (right).**c)** HAQERs in a vocal learning-associated gene, *ADCYAP1* (adenylate cyclase activating polypeptide 1), are marked as containing alternative promoters (TSS peaks of the FANTOM5 CAGE analysis), candidate cis-regulatory elements (ENCODE), and enhancers (ATAC-Seq peaks). For the latter, humans have a unique methylated region in layer 5 extra-telencephalic neurons of the primary motor cortex. Tracks are modified from the UCSC Genome Browser^75^ above the HAQER annotations and the comparative epigenome browser^76^ below the HAQER annotations. **d)** Lineage-specific structurally divergent regions (SDRs). SDRs are detected on two haplotypes and classified by different genomic content. The average number of total bases was assigned to the phylogenetic tree.

The second approach applied a top-down method that leveraged primate genome-wide all-by-all alignments to identify larger structurally divergent regions (SDRs) flanked by syntenic regions (**Methods; Supplemental Note XVI**) (Mao et al, 2024). We identified an average of 327 Mbp of SDRs per great ape lineage (**Fig. 5d**). SDRs delineate known sites of rapid divergence, including centromeres and subterminal heterochromatic caps but also numerous gene-rich SD regions enriched at the breakpoints of large-scale rearrangements (**Fig. SDR.S1**). The third approach used a gene-based analysis (TOGA–Tool to infer Orthologs from Genome Alignments) that focuses on the loss or gain of orthologous sequences in the human lineage (**Supplementary Note XVII**)^73^. TOGA identified six candidate genes from a set of 19,244 primate genes as largely restricted to humans (absent in >80% of the other apes; **Table TOGA.S1**). Among the candidate genes is a processed gene, *FOXO3B*, (present in humans and gorillas) whose paralog, *FOXO3A*, has been implicated in human longevity^74^. While the *FOXO3B* is expressed, its study has been challenging because it is embedded in a large, highly identical SD mediating Smith-Magenis deletion syndrome (**Fig. TOGA.S1**). While extensive functional studies will be required to characterize the hundreds of candidates we identified, we generated an integrated genomic (**Table SDR.S1**) and genic (**Table SDR.S2**) callset of accelerated regions for future investigation.

### Section III. New genomic regions

In addition to these improved insights into genes, repeats, and diversity, the contiguity afforded by the complete genomes allowed regions typically excluded from both reference genomes and evolutionary analyses to be investigated more systematically. We highlight four of the most notable: acrocentric, centromeres, subterminal heterochromatic caps, and lineage-specific SDs.

#### Acrocentric chromosomes and nucleolar organizer regions

The human acrocentric chromosomes (13, 14, 15, 21, 22) are the home of nucleolar organizer regions (NORs) and encode ribosomal RNA (rRNA) components of the 60S and 40S subunits. The precise sequence of the human NORs and the surrounding heterochromatin of the short arms was only recently elucidated in the first T2T human genome^8^. Human acrocentric chromosomes typically contain a single NOR with a head-to-tail rDNA array that is uniformly transcribed in the direction of the centromere. Each NOR is preceded by a distal junction (DJ) region extending approximately 400 kbp upstream of the rDNA array and including a 230 kbp palindrome (**Fig. Acro S1; Supplemental Note XVIII**) that encodes a long ncRNA associated with nucleolar function^77^. A variable patchwork of satellites and SDs flank the NOR, where heterologous recombination is thought to occur, as well as within the rDNA array itself, to maintain NOR homology through the action of concerted evolution^78^.

One conspicuous observation confirmed by our assemblies is that the ape NORs exist on different chromosomes for each species (**Fig. 6b, Fig. Acro.S2**). For example, HSA15 is NOR-bearing (NOR+) in human but not in chimpanzee or bonobo (NOR-), while HSA18 is NOR+ in both chimpanzee and bonobo, but NOR- in human^79^ (**Fig. 6b, Fig. Acro.S2**). Among great apes, we find the total number of NORs per haploid genome varies from as few as two in gorilla to 10 in both orangutans, while the siamang maternal genome sequenced here harbors only a single NOR (**Fig. 6b, Fig. Acro.S2**). We also find NORs on both orangutan and siamang Y chromosomes^10^, and partial DJ fragments on the chimpanzee and bonobo chrY (**Fig. 6b, Fig. Acro.S2**), suggesting their ancestral chrY may have been NOR+. Except for rRNA genes, all ape NOR-bearing chromosome short arms appear to be satellite-rich and gene-poor (**Fig. 6c,d**), with the NORs restricted to the end of an autosomal short arm or the end of a Y chromosome long arm. We identify, however, multiple acrocentric chromosomes with heterochromatic sequence on their short arm, but without an NOR (e.g., gorilla HSA2A, HSA9, HSA13, HSA15, and HSA18). Unlike the NOR+ acrocentrics, these NOR-acrocentrics carry multiple predicted protein-coding genes on their short arms. Thus, short-arm heterochromatin is strongly associated with ape NORs though not always predictive of their presence.

**Figure 6.**
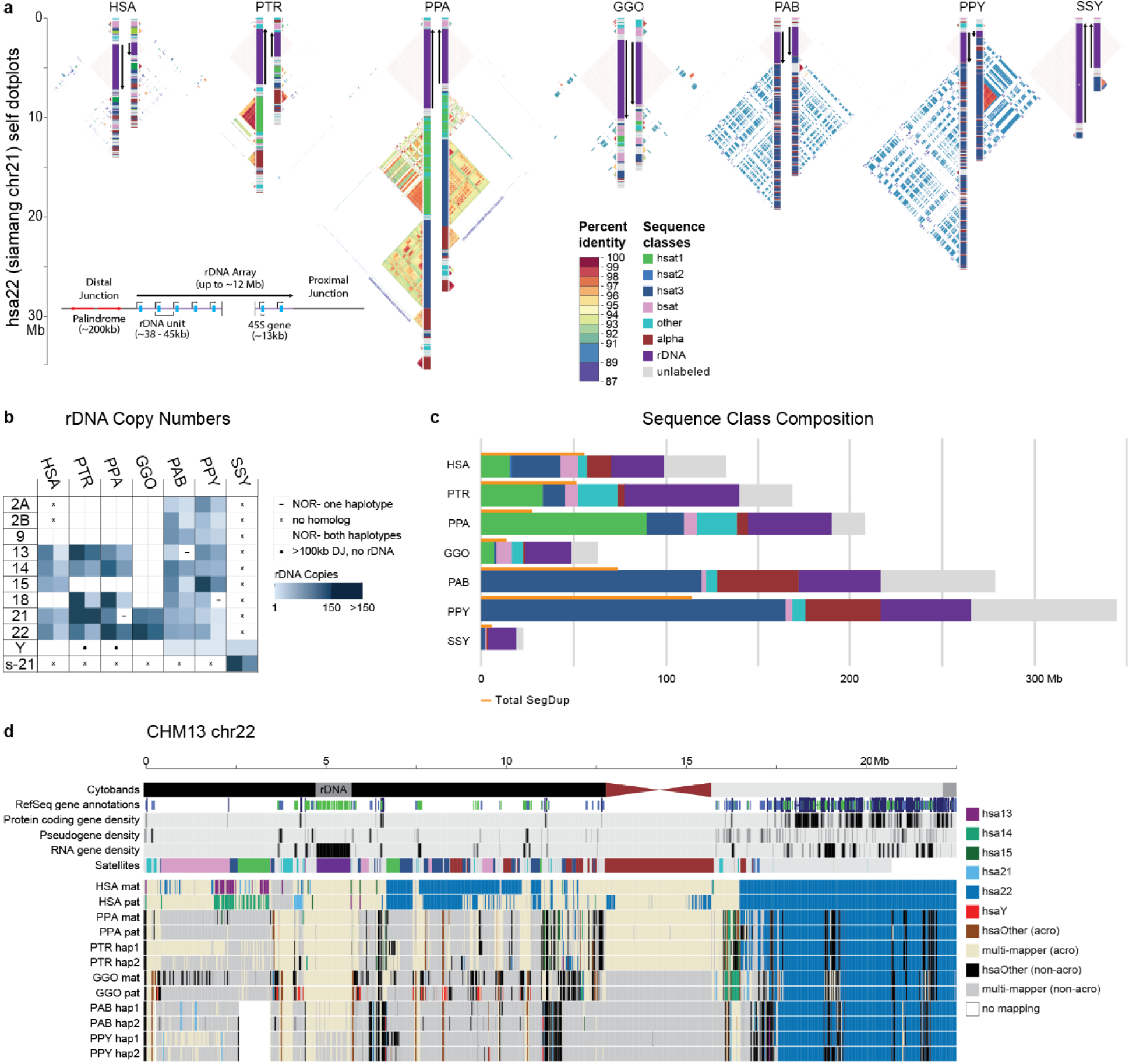
Organization and sequence composition of the ape acrocentric chromosomes. **a)** Sequence identity heatmaps and satellite annotations for the NOR+ short arms of both HSA22 haplotypes across all the great apes, and siamang chr21 (the only NOR+ chromosome in siamang) drawn with ModDotPlot^82^. The short arm telomere is oriented at the top of the plot, with the entirety of the short arm drawn to scale up to but not including the centromeric α-satellite. Heatmap colors indicate self-similarity within the chromosome, and large blocks indicate tandem repeat arrays (rDNA and satellites) with their corresponding annotations given in between. Human is represented by the diploid HG002 genome. **b)** Estimated number of rDNA units per haplotype (hap) for each species. HSA numbers are given in the first column, with the exception of “s-21” for siamang chr21, which is NOR+ but has no single human homolog. **c)** Sum of satellite and rDNA sequence across all NOR+ short arms in each species. “unlabeled” indicates sequences without a satellite annotation, which mostly comprise SDs. Total SD bases are given for comparison, with some overlap between regions annotated as SDs and satellites. **d)** Top tracks: chr22 in the T2T-CHM13v2.0 reference genome displaying various gene annotation metrics and the satellite annotation. Bottom tracks: For each primate haplotype, including the human HG002 genome, the chromosome that best matches each 10 kbp window of T2T-CHM13 chr22 is color coded, as determined by MashMap^83^. On the right side of the centromere (towards the long arm), HSA22 is syntenic across all species; however, on the short arm synteny quickly degrades, with very few regions mapping uniquely to a single chromosome, reflective of extensive duplication and recombination on the short arms. Even the human HG002 genome does not consistently align to T2T-CHM13 chr22 in the most distal (left-most) regions.

Estimated rDNA copy number for ape arrays varies from 1 on chrY of Bornean orangutan to 287 on HSA21 of chimpanzee; total diploid rDNA copy number similarly varies from 343 in siamang to 1,142 in chimpanzee (**Methods**, **Fig.6b, Fig. Acro.S2, Table Acro S3**), with total rDNA copy number varying widely between individual haplotypes of the same species, as expected^80^. Heterozygous NOR loss was observed in bonobo (HSA21), Sumatran orangutan (HSA13), and Bornean orangutan (HSA18), all of which were mediated by a truncation of the chromosome prior to the typical NOR location (**Fig. Acro.S2**). The structure and composition of both satellites and SDs varies considerably among the apes (**Fig.6a,c**). The orangutan acrocentrics are dominated by HSat3 and α-satellite, compared to the more balanced satellite composition of the other apes. Gorilla is notable for the presence of double NORs on both haplotypes of HSA22, with the additional NORs inverted relative to the first and including a complete DJ but only a single, inactive rDNA unit (**Fig. Acro.S3**).

At the chromosome level, the high level of synteny on the long arms of the NOR+ chromosomes quickly degrades when transitioning to the short arm, with almost no sequence aligning uniquely between different ape species (**Fig. 6d**). Even the haplotypes of a single human genome aligned best to different reference chromosomes on their distal ends, supporting prior observations of extensive heterologous recombination^78^. Despite their widespread structural variation, the ape NOR+ chromosomes share common features such as homogeneous rDNA arrays containing highly conserved rRNA genes. We extracted representative rDNA units from each assembly to serve as a reference for each species and confirmed a similar sequence structure, including the presence of a central microsatellite region within the intergenic spacer sequence for all species (**Fig. Acro.S4**), but with relatively high nucleotide substitution rates outside of the >99% identical 18S and 5.8S coding regions^81^. Despite its conserved co-linear structure, nucleotide identity of the intergenic spacer varied from 95.19% for human versus bonobo to just 80.60% for human versus siamang (considering only SNVs, **Table Acro S2**). The DJ sequence was found to be conserved across all great apes and present as a single copy per NOR, including the palindromic structure typical of the human DJ, with the exception of siamang, which contains only one half of the palindrome on each haplotype but in opposite orientations (**Fig. Acro.S5**). The transcriptional direction of all rDNA arrays is consistent within each species, with the chimpanzee and bonobo arrays inverted relative to human (**Fig. 6a**). This inversion includes the entire DJ sequence, confirming a prior FISH analysis that found the chimpanzee DJ had been relocated to the centromeric side of the rDNA array^77^. Our comparative analysis supports the DJ as a functional component of ape NORs that is consistently positioned upstream of rRNA gene transcription, rather than distally on the chromosome arm.

#### Centromere satellite evolution

The assembly of five nonhuman great ape genomes allowed us to assess the sequence, structure, and evolution of centromeric regions at base-pair resolution for the first time. Using these assemblies, we identify 227 contiguous centromeres out of a possible 230 centromeres across five NHPs, each of which were composed of tandemly repeating α-satellite DNA organized into higher-order repeats (HORs) belonging to one or more α-satellite suprachromosomal families (SFs) (**Fig. 7; Fig. CEN.S1; Supplemental Note XIX**). In specific primate lineages, different SFs have risen to high frequency, such as SF5 in the orangutan and SF3 in the gorilla. We carefully assessed the assembly of each of these centromeres, checking for collapses, false duplications and misjoins (**Methods**), and found that approximately 85% of bonobo, 69% of chimpanzee, 54% of gorilla, 63% of Bornean orangutan, but only 27% of Sumatran orangutan centromeres are complete and correctly assembled (**Fig. 7a; Fig. CEN.S1**). Most of the assembly errors are few (∼2 per centromere haplotype, on average) and typically involve a few 100 kbp of centromere satellite sequence that still need further work to resolve.

**Figure 7.**
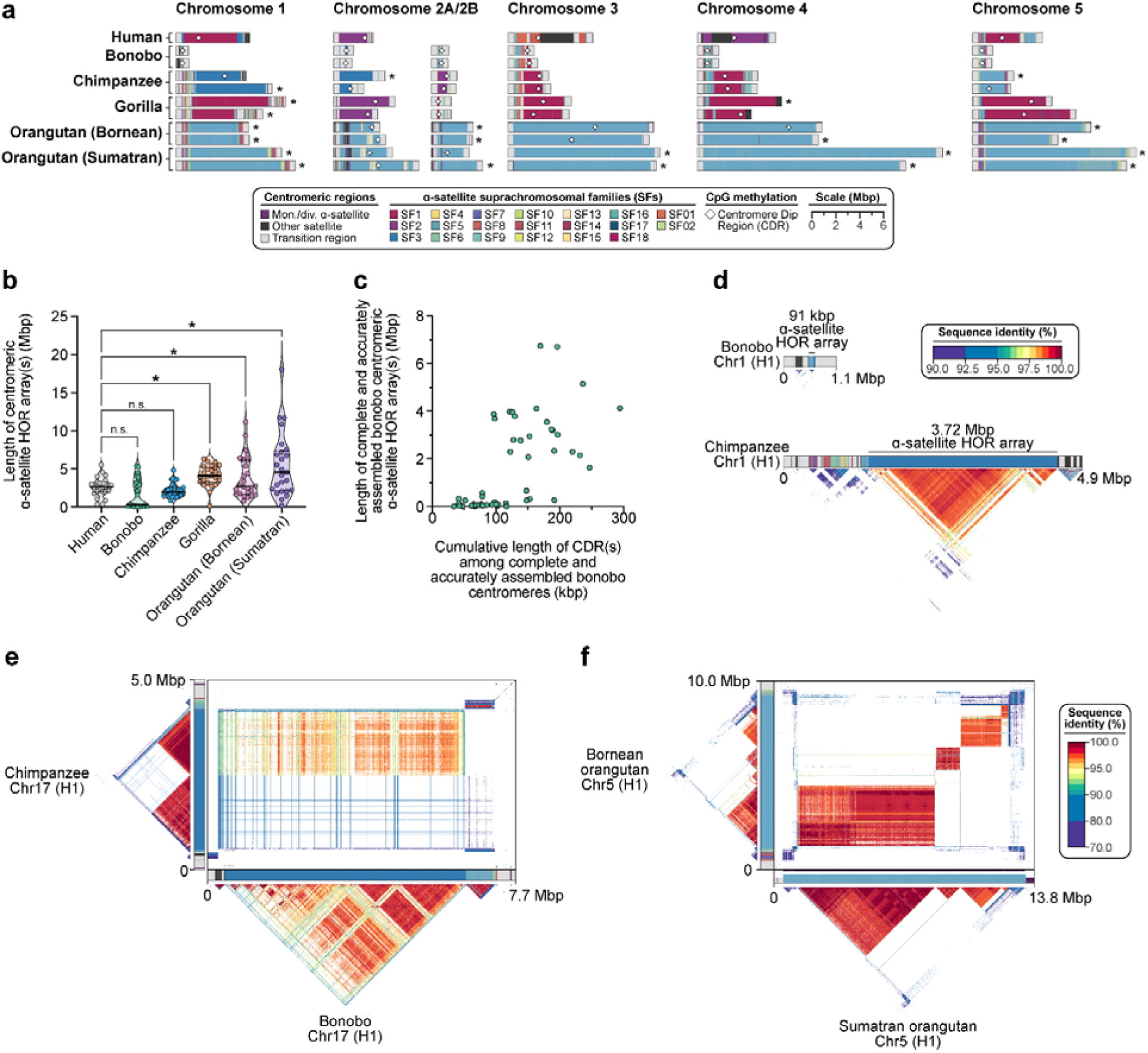
Assembly of 237 NHP centromeres reveals variation in α-satellite HOR array size, structure, and composition. **a)** Sequence and structure of α-satellite HOR arrays from the human (T2T-CHM13), bonobo, chimpanzee, gorilla, Bornean orangutan, and Sumatran orangutan chromosome 1–5 centromeres, with the α-satellite suprachromosomal family (SF) indicated for each centromere. The sequence and structure of all completely assembled centromeres is shown in **Fig. CENSATS1**. **b)** Variation in the length of the α-satellite HOR arrays for NHP centromeres. Bonobo centromeres have a bimodal length distribution, with 28 chromosomes showing “minicentromeres” (with α-satellite HOR arrays <700 kbp long). **c)** Correlation between the length of the bonobo active α-satellite HOR array and the length of the CDR for the same chromosome. **d)** Example showing that the bonobo and chimpanzee chromosome 1 centromeres are divergent in size despite being from orthologous chromosomes. **e)** Sequence identity heatmap between the chromosome 17 centromeres from bonobo and chimpanzee show a common origin of sequence as well as the birth of new α-satellite HORs in the chimpanzee lineage. **f)** Sequence identity heatmap between the chromosome 5 centromeres from the Bornean and Sumatran orangutans show highly similar sequence and structure, except for one pocket of α-satellite HORs that is only present in the Bornean orangutan. *, *p* < 0.05; n.s., not significant.

Focusing on the completely assembled centromeres, we identify several unique characteristics specific to each primate species. First, we find that the bonobo centromeric α-satellite HOR arrays are, on average, 0.65-fold the length of human centromeric α-satellite HOR array and 0.74-fold the length of its sister species, chimpanzee (**Fig. 7b**). A closer examination of bonobo α-satellite HOR array lengths reveals that they are bimodally distributed, with approximately half of the bonobo centromeres (27/48) having an α-satellite HOR array with a mean length of 110 kbp (range: 15–674 kbp) and the rest (21/48) having a mean length of 3.6 Mbp (range: 1.6–6.7 Mbp; **Fig. 7c**). The bimodal distribution persists in both sets of bonobo haplotypes. This >450-fold variation in bonobo α-satellite HOR array length has not yet been observed in any other primate species and implies a wide range of centromeric structures and sizes compatible with centromere function. Indeed, no “mini-centromere” arrays have been observed in the chimpanzee, despite its recent speciation from bonobo (∼1.7 mya; **Fig. 7d**).

As previously noted^84^, chimpanzee α-satellite HOR arrays are consistently smaller: 0.86-fold the length of their human counterparts (**Fig. 7a**). Additionally, the chimpanzee centromeres are typically composed of a single α-satellite HOR array flanked by short stretches of divergent α-satellite HORs and monomeric sequences, which are interspersed with TEs before extending into the p- and q-arms (**Fig. 7d**). In contrast, the gorilla α-satellite HOR arrays are, on average, 1.58-fold larger than human (**Fig. 7a-b**), and unlike bonobo and chimpanzee, they are composed of punctuated regions of α-satellite HORs, or regions of α-satellite HORs that have high sequence identity within them but much lower sequence identity with neighboring regions, flanked by larger transition zones to monomeric α-satellite sequence. The gorilla centromeres show a high degree of haplotypic variation, with many paternal and maternal centromeres varying in size, sequence, and structure. We find that 30.4% (7 out of 23) gorilla α-satellite HOR array pairs vary in size by >1.5-fold (especially HSA chromosomes 1, 2a, 4, 10, 15, 18 and 19), and 9 out of 23 pairs (∼39.1%) have α-satellite HOR arrays with >5% sequence divergence between homologs (HSA chromosomes 1, 4–6, 10–12, 15, and 19). Finally, the Bornean and Sumatran orangutan α-satellite HOR arrays are among the largest (1.52- and 2.11-fold larger, on average, than humans; **Fig. 7b**) and are characterized by multiple pockets of divergent α-satellite HORs. A typical Bornean or Sumatran orangutan centromere has three or four distinct pockets of α-satellite HORs, with up to nine distinct HOR arrays observed in a single centromere (Bornean chromosome 19).

Congeneric species of *Pan* and *Pongo* present an opportunity to assess the evolution of centromeric α-satellites over evolutionary periods of time. Comparison of the centromeric α-satellite HOR arrays from orthologous chromosomes across the bonobo and chimpanzee genomes reveals, for example, that 56% of them (14 out of 25 centromeres, including both X and Y) share a common identifiable ancestral sequence, such as that present in HSA chromosome 17 (**Fig. 7e**). On this chromosome, the entire bonobo α-satellite HOR array is ∼92–99% identical to one domain of α-satellite HORs present in the chimpanzee centromere. However, the chimpanzee centromere contains a second domain of α-satellite HORs that spans approximately half of the α-satellite HOR array. This domain is <70% identical to the bonobo α-satellite HORs, indicating the formation of a new α-satellite HOR array subregion acquired specifically in the chimpanzee lineage. Thus, over <2 mya, a new α-satellite HOR arises and expands to become the predominant HOR distinguishing two closely related species (**Fig. 7e**). Given the shorter speciation time of orangutan (0.9 mya), α-satellite HOR evolution is more tractable, with α-satellite HORs sharing >97% sequence identity, including domains with 1:1 correspondence. However, in about a fifth of orangutan centromeres, we identify stretches of α-satellite HORs present in Bornean but not Sumatran (or vice versa). The emergence of lineage-specific α-satellite HOR sequences occurred on five chromosomes (HSA chromosomes 4, 5, 10, 11, 16; **Fig. 7f**) and is marked by extremely high sequence identity (>99%) between α-satellite HOR arrays, suggesting rapid turnover and homogenization of newly formed orangutan α-satellite HORs.

We leveraged the new assemblies of these NHP centromeres to assess the location and distribution of the putative kinetochore—the large, proteinaceous structure that binds centromeric chromatin and mediates the segregation of chromosomes to daughter cells during mitosis and meiosis^85,86^. Previous studies in both humans^8^ and NHPs^84,87^ have shown that centromeres typically contain one kinetochore site, marked by one or more stretch of hypomethylated CpG dinucleotides termed the centromere dip region (CDR)^88^. We carefully assessed the CpG methylation status of all 237 primate centromeres and found that all contain at least one region of hypomethylation, consistent with a single kinetochore site. Focusing on the bonobo centromeres, where we find a bimodal distribution in α-satellite HOR array length (**Fig. 7b**), we show that CDR length and centromere length correlate (R^2^=0.41). In other words, the bonobo “minicentromeres” tend to associate with smaller CDRs when compared to larger centromeres (**Fig. 7c**). While much more in-depth functional studies need to be performed, this finding suggests the reduced α-satellite HOR arrays in bonobo are effectively limiting the distribution of the functional component of the centromere.

#### Subterminal heterochromatin

In addition to centromeres, we completely sequenced and assembled the subterminal heterochromatic caps of siamang, chimpanzee, bonobo, and gorilla (**Fig. 1c & Fig. 8a**). In total, these account for 1.05 Gbp of subterminal satellite sequences (642 Mbp or 18.2% of the siamang genome). These massive structures (up to 26 Mbp in length) are thought to be composed almost entirely of tandem repetitive DNA: a 32 bp AT-rich satellite sequence, termed pCht7 in *Pan* and gorilla, or a 171 bp α-satellite repeat present in a subset of gibbon species^89–91^. While their function is not known, these chromosomal regions have been implicated in nonhomologous chromosome exchange and unique features of telomeric RNA metabolism^92,93^. Our analysis indicates that we successfully sequenced 79 gapless subterminal caps in gorilla (average length=6.6 Mbp) and 57 and 46 caps in chimpanzee and bonobo, including both haplotypes (average lengths 4.8 and 5.2 Mbp, respectively) with less than 3.8% of pCht arrays flagged as potentially misassembled (**Fig. 8a**). Siamangs possess the largest (average length 6.7 Mbp) and most abundant subterminal satellite blocks (96 out of 100 chromosomal ends across the two haplotypes).

**Figure 8.**
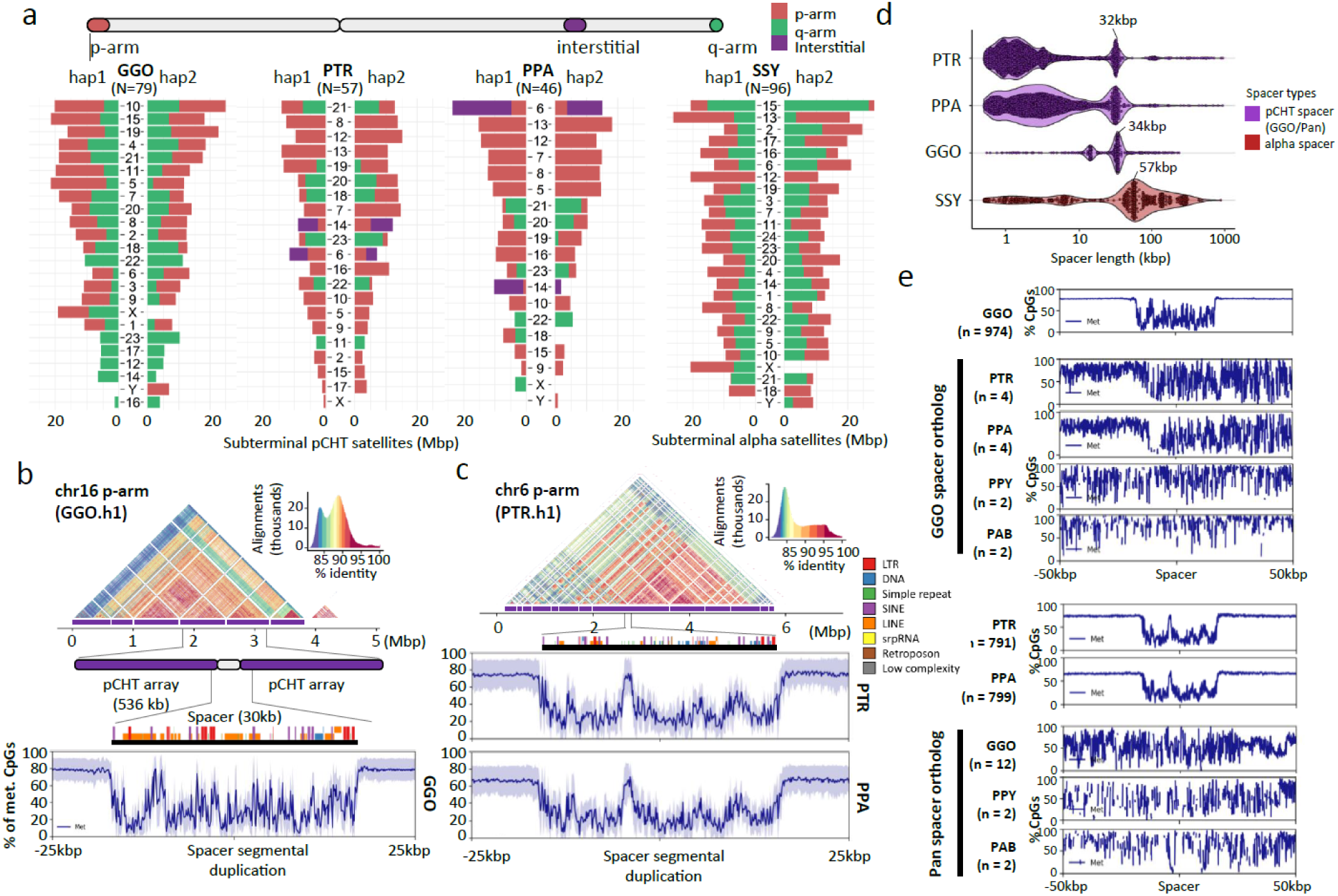
Subterminal heterochromatin analyses. **a)** Overall quantification of subterminal pCht/α-satellites in the African great ape and siamang genomes. The number of regions containing the satellite is indicated below the species name. The pChts of diploid genomes are quantified by Mbp, for ones located in p-arm, q-arm, and interstitial, indicated by orange, green, and purple. Organization of the subterminal satellite in **b)** gorilla and **c)** *pan* lineages. The top shows a StainedGlass alignment plot indicating pairwise identity between 2 kbp binned sequences, followed by the higher order structure of subterminal satellite unit, as well as the composition of the hyperexpanded spacer sequence and the methylation status across the 25 kbp up/downstream of the spacer midpoint. **d)** Size distribution of spacer sequences identified between subterminal satellite arrays. **e)** Methylation profile of the subterminal spacer SD sequences compared to the interstitial ortholog copy.

In gorilla and chimpanzees (*Pan*), the caps are organized into higher order structures where pCht subterminal satellites form tracts of average length of 335 to 536 kbp interrupted by spacer SD sequences of a modal length of 32 kbp (*Pan*) or 34 kbp (gorilla; **Fig. SubterminalS1**). The spacer sequences are each unique to the *Pan* and gorilla lineages but we confirm that each began originally as a euchromatic sequence that became duplicated interstitially in the common ancestor of human and apes. For example, the 34 kbp spacer in gorilla maps to a single copy sequence present in orangutans and human chromosome 10, which began to be duplicated interstitially on chromosome 7 in chimpanzee but only in gorilla became associated with pCht satellites expanding to over 477 haploid copies as part of the formation of the heterochromatic cap. Similarly, the ancestral sequence of pan lineage spacer maps syntenically to orangutan and human chr9. The ancestral sequence duplicated to multiple regions in gorilla (q-arms of chr4, 5, 8, X, and p-arms of chr2A and 2B), before being captured and hyperexpanded (>345 copies) to form the structure of subterminal satellites of chimpanzee and bonobo. Analyzing CpG methylation, we find that each spacer demarcates a pocket of hypomethylation flanked by hypermethylated pCht arrays within the cap (**Fig. 8b-d**). Of note, this characteristic hypomethylation pattern is not observed at the ancestral origin or interstitially duplicated locations (**Fig. 8e**), suggesting an epigenetic feature not determined solely by sequence but by its association with the subterminal heterochromatic caps. Similar to the great apes, we find evidence of a hypomethylated spacer sequence also present in the siamang subterminal cap; however, its modal length is much larger (57.2 kbp in length) and its periodicity is less uniform occurring every 750 kbp (**Fig. SuterminalS2**). Nevertheless, the fact that these similar epigenetic features of the spacer evolved independently may suggest a functional role with respect to subterminal heterochromatic caps.

#### Lineage-specific segmental duplications and gene families

Compared to previous read-depth-based approaches that simply estimated copy number of SDs^94,95^, T2T genomes increase SD content and resolve sequence structures allowing us to distinguish SDs that are novel by location and composition within each species (**Fig. SD.S1**). Nonhuman great ape genomes (excluding siamang) generally harbor more SDs (**Fig. 9a**) when compared to humans (∼192 vs. an average of 215 Mbp in the nonhuman; they are comparable when normalized by the genome size). We also find that great apes, on average, have the highest SD content (208 Mbp per haplotype) when compared to non-ape lineages: mouse lemur, gelada, marmoset, owl monkey, and macaque (68.8–161 Mbp) (**Fig. 9a & SD.S2**). In contrast to our previous analysis^96^, orangutans show the greatest amount of SDs (225.3 Mbp/hap) compared to African great apes (204.3 Mbp/hap), which also exhibit larger interspersion of intrachromosomal SDs (**Fig. 9b & SD.S3**). The increased SD content in orangutans is due to a greater number of acrocentric chromosomes (10 vs. 5 on average for other apes) and a preponderance of clustered duplications. Consistent with the expansion of Asian great ape SDs, we find the largest number of lineage-specific SDs in the *Pongo* lineage (93.3 Mbp, followed by gorilla- and human-specific SDs (75.1 and 60.6 Mbp, respectively). Many SDs (79.3 to 95.6 Mbp per haplotype) in orangutan constitute massive, Mbp-scale SD clusters, including a mixture of tandem and inverted duplications up to 21.5 Mbp in size; in other species, the total number of such clustered duplications accounts for only 30 to 40 Mbp per haplotype (except bonobo). In general, the number of SDs assigned to different lineages correlates with branch length (r^2^=0.80) (**Fig. SD.S4**) with the exception of siamang and some ancestral nodes reflecting the great ape expansion of SDs^96^.

**Figure 9.**
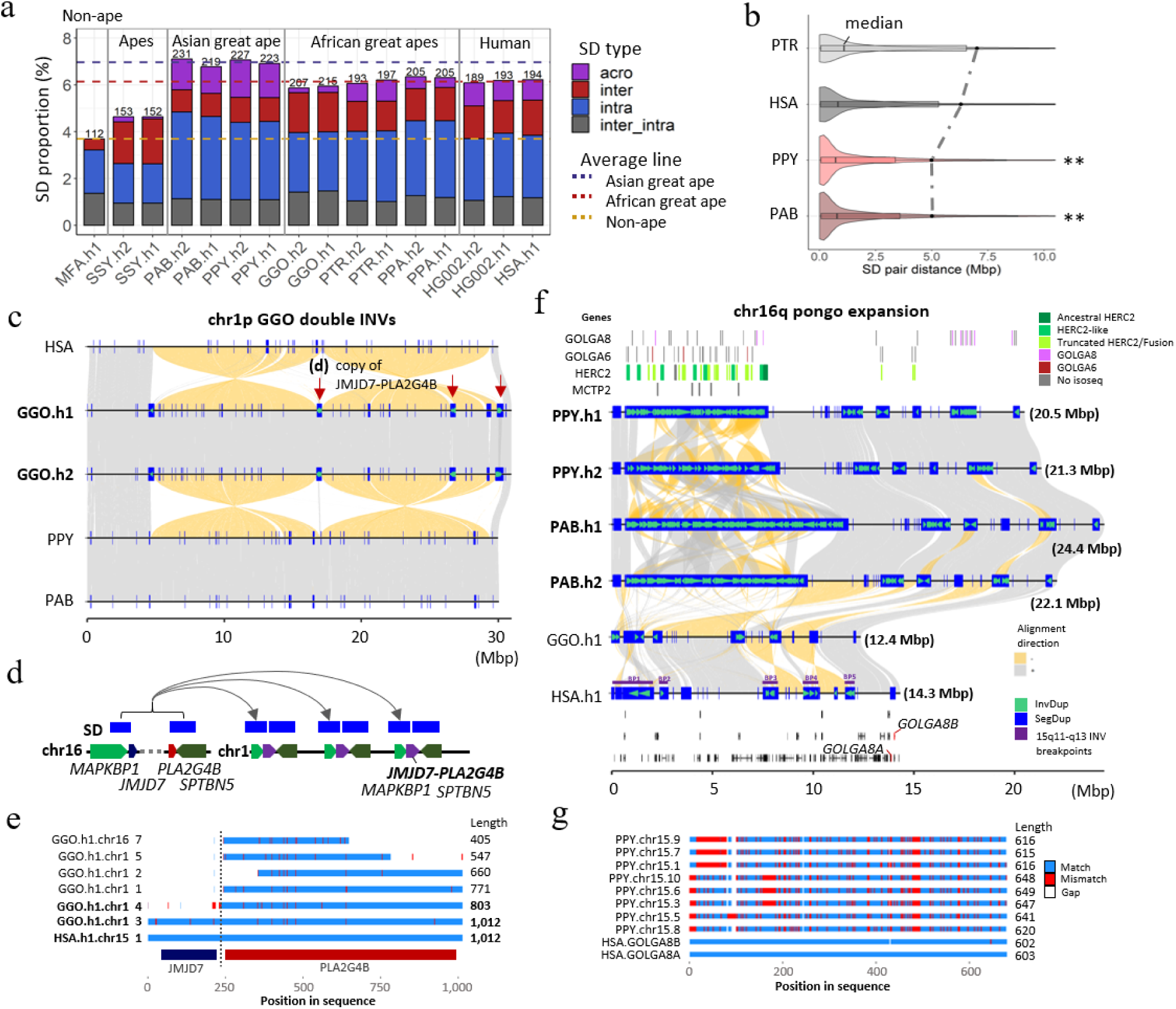
Ape SD content and new genes. **a)** Comparative analysis of primate SDs comparing the proportion of acrocentric (purple), interchromosomal (red), intrachromosomal (blue), and shared inter/intrachromosomal SDs (gray). The total SD Mbp per genome is indicated above each histogram with the colored dashed lines showing the average Asian, African great ape, and non-ape SD (MFA=*Macaca fascicularis*^78^; see **Fig.SD.S2** for additional non-ape species comparison). **b)** A violin plot distribution of pairwise SD distance to the closest paralog where the median (black line) and mean (dashed line) are compared for different apes (see **Fig. SD.S3** for all species and haplotype comparisons). An excess of interspersed duplications (*p*<0.001 one-sided Wilcoxon rank sum test) is observed for chimpanzee and human when compared to orangutan. **c)** Alignment view of chr1 double inversion. Alignment direction is indicated by + as gray and – as yellow. SDs as well as those with inverted orientations are indicated by blue rectangles and green arrowheads. The locations in which the *JMJD7-PLA2G4B* gene copy was found are indicated by the red arrows. **d)** duplication unit containing three genes including *JMJD7-PLA2G4B*. **e)** Multiple sequence alignment of the translated *JMJD7-PLA2G4B*. Match, mismatch and gaps are indicated by blue, red and white. Regions corresponding to each of *JMJD7* or *PLA2G4B* are indicated by the track below. **f)** Alignment view of chr16q. The expansion of *GOLGA6/8, HERC2,* and *MCTP2* genes are presented in the top track. 16q recurrent inversion breakpoints are indicated in the human genome. The track at the bottom indicates the gene track with *GOLGA8* human ortholog in red. **g)** Multiple sequence alignment of the translated *GOLGA8*.

Leveraging the increased sensitivity afforded by FLNC Iso-Seq, we annotated the transcriptional content of lineage-specific SDs identifying hundreds of potential genes, including gene family expansions often occurring in conjunction with chromosomal evolutionary rearrangements. We highlight two examples in more detail. First, at two of the breakpoints of a 30 Mbp double inversion of gorilla chromosome 1, we identify a gorilla-specific expansion of the genes *MAPKBP1* and *SPTBN5* as well as *PLA2G4B-JMJD7* (**Fig. 9c-e**) originating from an interchromosomal SD from ancestral loci mapping to HSA chromosome 15 (duplicated in other chromosomes in chimpanzee and bonobo; **Table Genes.S5**). We estimate these duplications occurred early after gorilla speciation, 6.1 mya (**Fig. SD.S5a**) followed by subsequent expansion resulting in the addition of eight copies (one truncated) mapping to two of the breakpoints of the double inversion. Investigating the Iso-Seq transcript model of this gene, revealed that five of the new gorilla copies are supported by multi-exon transcripts. Two of these additional copies possess valid start and stop codons spanning at least 70% of the homologous single-copy ortholog gene in humans (**Fig. 9e & SD.S5a**). Notably, the ancestral copy of this gene in gorilla (HSA chromosome 15q) is found to be highly truncated (40% of original protein) suggesting that the new chromosome 1 copies may have assumed and refined the function.

Second, in orangutan, we find a restructured 20-Mbp region corresponding to the Prader-Willi syndrome (PWS)^65^ and the 15q13 microdeletion syndrome^97^ region in humans. This includes a massive 6.8–10.8 Mbp expansion of clustered tandem and inverted duplications mapping distally to breakpoint 1 of PWS as well as smaller 200–550 kbp expansions of *GOLGA6/8* repeats distal to PWS BP3/4 (**Fig. 9f**). We estimate that the larger region, alone, is composed of 87–111 copies of fragments of *GOLGA6/8*, *HERC2* and *MCTP2*. We find Iso-Seq transcript support for 37–39 distinct orangutan copies. Using *GOLGA8* as a marker, we show that it has expanded to 10–12 copies (>70% of original length) in orangutan but exists as a single copy in gorilla and bonobo and in two copies (*GOLGA8A* and *B*) in human out of multiple *GOLGA8* genes, retaining at least 70% of sequence compared to orangutan sequence (**Fig. 9f-g, Fig. SD.S5b**). We estimate that the *Pongo* expansion of *GOLGA8* occurred 7.3 mya (**Fig. SD.S5b**), long before the species diverged. Alignment of the translated peptide sequence, we observe 17.1–23.7% divergence from the human copy (*GOLGA8A*; **Fig. 9g**). Based on studies of the African great ape genomes and humans, *GOLGA8* was among more than a dozen loci defined as “core duplicons” promoting the interspersion of SDs and genomic instability via palindromic repeat structures^98,99^. Our findings extend this recurrent genomic feature for the *GOLGA8* duplicons to the Asian ape genomes.

## DISCUSSION

The completion of the ape genomes significantly refines previous analyses providing a more definitive resource for all future evolutionary comparisons. These include an improved and more nuanced understanding of species divergence, human ancestral alleles, incomplete lineage sorting, gene annotation, repeat content, divergent regulatory DNA, and complex genic regions as well as species-specific epigenetic differences involving methylation. These preliminary analyses reveal hundreds of new candidate genes and regions to account for phenotypic differences among the apes. For example, we observed an excess of HAQERS corresponding to bivalent promoters thought to contain gene regulatory elements that exhibit precise spatiotemporal activity patterns in the context of development and environmental response^100^. Bivalent chromatin state enrichments have not yet been observed in fast-evolving regions from other great apes, which may reflect limited cross-species transferability of epigenomic annotations from human. The finding of a HAQER enriched gene, *ADCYAP1,* that is differentially regulated in speech circuits and methylated in the layer 5 projection neurons that make the more specialized direct projections to brainstem motor neurons in humans, shows the promise of T2T genomes to identify hard to sequence regions important for complex traits. Perhaps most importantly, we provide an evolutionary framework for understanding the ∼10%– 15% of highly divergent, previously inaccessible regions of ape genomes. In this regard, we highlight a few noteworthy findings.

### Orangutans show the greatest amount of recent segmental duplication

Comparative analyses suggest expansion of SDs in the common ancestor of the great ape lineage as opposed to the African great ape lineage as we originally proposed based on sequence read-depth analyses back to the human reference genome^95,96^. This discrepancy highlights the importance of *ab initio* sequence genome assembly of related lineages that are comparable in quality and contiguity. The assembly of the acrocentric chromosomes (of which orangutans have the maximum at 9/10) and the resolution of massive (10–20 Mbp) tandem SDs in the orangutan species account for the increase in SD content among the Asian great apes. The African great ape lineage still stands out for having the largest fraction of interspersed SDs—a genomic architectural feature that promotes recurrent rearrangements facilitating syndromic disease associated with autism and developmental delay in the human species^101^. Complete sequence resolution of NHP interspersed SDs provides a framework for understanding disease-causing copy number variants in these other NHP lineages^102^.

### Large-scale differences in acrocentric chromosomes

The short arms of NOR+ ape acrocentric chromosomes appear specialized to encode rRNA genes. On the autosomes, ape NORs exist exclusively on the acrocentric chromosomes, embedded within a gene-poor and satellite-rich short arm. On the Y chromosome, NORs occur occasionally toward the end of the chromosome and adjacent to satellites shared with other acrocentric chromosomes. Prior analysis of the human pangenome suggested heterologous recombination between chromosomes with NORs as a mechanism for concerted evolution of the rRNA genes^78,103^. Comparative analysis of ape genomes provides further support for this hypothesis. For example, the uniform direction of all rDNA arrays within a species would permit crossover recombination between heterologous chromosomes without substantial karyotypic consequence. However, rare translocations, mediated by the large SDs commonly surrounding the NORs, have occurred during ape evolution, resulting in a different complement of NOR+ acrocentric chromosomes and possibly creating reproductive barriers associated with speciation^104^.

### Lineage-specific gene family expansions/explosions and rearrangements

The number of lineage-specific duplications that encode transcripts and potential genes is now estimated at hundreds per ape lineage often occurring at sites of evolutionary chromosomal rearrangements that have been historically difficult to sequence resolve (**Table SD.S1**). Our analysis has uncovered hundreds of fixed inversions frequently associated with the formation of these lineage-specific duplications. These findings challenge the predominant paradigm that subtle changes in regulatory DNA^105^ are the major mechanism underlying ape species differentiation. Rather, the expansion, contraction, and restructuring of SDs lead to not only dosage differences but concurrent gene innovation and chromosomal structural changes^106^. Indeed, in the case of human, four such gene family expansions, namely *NOTCH2NL*^107^, *SRGAP2C*^108,109^, *ARHGAP11*^99^ and *TBC1D3*^110,111^, have been functionally implicated over the last decade in the frontal cortical expansion of the human brain^107–110^ as well as human-specific chromosomal changes^99^. Detailed characterization of the various lineage-specific expansions in NHPs will no doubt be more challenging yet it is clear that such SDRs are an underappreciated genic source of interspecific difference and potential gene neofunctionalization.

### Bonobo minicentromeres

We identified several idiosyncratic features of centromere organization and structure that characterize the different ape lineages, significantly extending earlier observations based on the characterization of five select centromeres^84^. Perhaps the most remarkable is the bimodal distribution of centromere HOR length in the bonobo lineage—19 of the 48 bonobo centromeres are, in fact, less than 100 kbp in size. Given the estimated divergence of the *Pan* lineage, such 300-fold reductions in size must have occurred very recently—in the last million years. These bonobo “minicentromeres” appear fully functional with a well-defined CDR (encompassing all of the α-satellite DNA). Thus, their discovery may provide a roadmap for the design of smaller, more streamlined artificial chromosomes for the delivery and stable transmission of new genetic information in human cells^112^.

### Epigenetic architecture of subterminal heterochromatin

Our analysis suggests that the subterminal chromosomal caps of chimpanzee, gorilla, and siamang have evolved independently to create multi-Mbp of heterochromatin in each species. In chimpanzee and gorilla, we define a common organization of a subterminal spacer (∼30 kbp in size) that is hypomethylated and flanked by hypermethylated heterochromatic satellite with a periodicity of one spacer every 335– 536 kbp of satellite sequence (pCht satellite in hominoids and α-satellite in hylobatids). In each case, the spacer sequence differs in its origin but has arisen as an ancestral SD^89^ that has become integrated and expanded within the subterminal heterochromatin. In contrast to the ancestral sequences located in euchromatin, the spacer sequences embedded within the subterminal caps acquire more pronounced hypomethylation signatures suggesting an epigenetic feature. This subterminal hypomethylation pocket is reminiscent of the CDRs identified in centromeres that define the sites of kinetochore attachment^113^ as well as methylation dip region observed among some acrocentric chromosomes^26^. It is tempting to speculate that the subterminal hypomethylation pocket may represent a site of protein binding or a “punctuation” mark perhaps facilitating ectopic exchange and concerted evolution driving persistent subtelomeric associations and meiotic exchanges between nonhomologous chromosomes^114^.

While the ape genomes sampled here are nearly complete, some limitations remain. Sequence gaps still exist in the acrocentric centromeres and a few other remaining complex regions where the largest and most identical tandem repeats reside. This is especially the case for the Sumatran orangutan centromeres where only 27% are completely assembled. The length (nearly double the size) and the complex compound organization of orangutan α-satellite HOR sequence will require specialized efforts to completely order and orientate^84^. Nevertheless, with the exception of these and other large tandem repeat arrays, we estimate that ∼99.5% of the content of each genome has been characterized and is correctly placed. Second, although we completed the genomes of a representative individual, we sequenced and assembled only two haplotypes from each species and more than 15 species/subspecies of apes remain^115^. Sampling more closely related species that diverged within the last million years will provide a unique opportunity to understand the evolutionary processes shaping the most dynamic regions of our genome. High-quality assemblies of all chimpanzee species^116^, as well as the numerous gibbon species^117^, will provide critical insight into selection, effective population size, and the rapid structural diversification of ape chromosomes at different time points. Finally, while high-quality genomes help eliminate reference bias, they do not eliminate annotation biases that favor the human. This will be especially critical for both genes and regulatory DNA that have rapidly diverged between the species.

## DATA AVAILABILITY

The raw genome sequencing data generated by this study are available under NCBI BioProjects, PRJNA602326, PRJNA976699–PRJNA976702, and PRJNA986878–PRJNA986879 and transcriptome data are deposited under BioProjects, PRJNA902025 (UW Iso-Seq) and PRJNA1016395 (UW and PSU Iso-Seq and short-read RNA-seq). The genome assemblies are available from GenBank under accessions: GCA_028858775.2, GCA_028878055.2, GCA_028885625.2, GCA_028885655.2, GCA_029281585.2 and GCA_029289425.2. Genome assemblies can be downloaded via NCBI (https://www.ncbi.nlm.nih.gov/datasets/genome/?accession=GCF_028858775.2,GCF_029281585.2,GCF_028885625.2,GCF_028878055.2,GCF_028885655.2,GCF_029289425.2). Convenience links to the assemblies and raw data are available on GitHub (https://github.com/marbl/Primates) along with a UCSC Browser hub (https://github.com/marbl/T2T-Browser). The UCSC Browser hub includes genome-wide alignments, CAT annotations, methylation, and various other annotation and analysis tracks used in this study. The T2T-CHM13v2.0 and HG002v1.0 assemblies used here are also available via the same browser hub, and from GenBank via accessions GCA_009914755.4 (T2T-CHM13), GCA_018852605.1 (HG002 paternal), and GCA_018852615.1 (HG002 maternal). The alignments are publicly available to download or browse in HAL118 MAF and UCSC Chains formats (https://cglgenomics.ucsc.edu/february-2024-t2t-apes).

## CODE AVAILABILITY

All code used for the reported analyses is available from our project’s GitHub repository: https://github.com/marbl/Primates

## ACRONYMS & ABBREVIATIONS

AQER: ancestor quickly evolved region
cDNA: complementary deoxyribonucleic acid
CDR: centromere dip region
CRE: cis-regulatory element
DJ: distal junction [region]
ENC: evolutionary neocentromere
ERV: endogenous retrovirus
FLNC: full-length non-chimeric
GGO: gorilla
HAQER: human ancestor quickly evolved region [human branch]
HAS: human
HiFi: high-fidelity
HOR: higher-order repeat
ILS: incomplete lineage sorting
LINE: long interspersed nuclear element
LTR: long terminal repeats
MEI: mobile element insertion
MHC: major histocompatibility complex
mya: million years ago
ncRNA: noncoding RNA
Ne: effective population sizes
NHP: nonhuman primate
NOR: nucleolar organizer region
NUMT: nuclear sequence of mitochondrial DNA origin
ONT: Oxford Nanopore Technologies
ORF: open reading frame
PAB: Sumatran orangutan
PacBio: Pacific Biosciences Inc.
PGGB: pangenome graph builder
PLE: Penelope-Like Retroelements
PPA: bonobo
PPY: Bornean orangutan
PTR: chimpanzee
PWS: Prader-Willi syndrome
RC: rolling circle repeats
rDNA/rRNA: ribosomal deoxyribonucleic/ribonucleic acid
SD: segmental duplication
SDR: structurally divergent region
SF: suprachromosomal family
SINE: short interspersed nuclear element
SNV: single-nucleotide variant
SVA: SINE-VNTR-Alu element
T2T: telomere-to-telomere
TE: transposable element
TOGA: Tool to infer Orthologs from Genome Alignments
UL: ultra-long
VNTR: variable number tandem repeat

## ACKNOWLEDGMENTS

We thank Richard Buggs for useful suggestions and Tonia Brown, Agostinho Antunes and Mohamed Emam for editing the manuscript and supplementary note. This research was supported, in part, by the Intramural Research Program of the National Human Genome Research Institute, National Institutes of Health (NIH), and extramural NIH grants R35GM151945 (to K.D.M.), R01 HG002385, R01 HG010169, U24 HG007497 (to E.E.E.), R35 GM146926 (to Z.A.S.). R35 GM146886 (to C.D.H.), R35GM142916 (to P.H.S.), R01HG012416 (to P.H.S. and E.G.), R35HG011332 (to C.B.L.), R01GM123312 (to R.J.O.), U24 HG010263 (to M.C.S), R35GM133747 (to R.C.M.), U41HG007234 (to P.H., M.D., B.P.), R01HG010329 (to P.H., M.D.), R01MH120295 (to M.D.), 1P20GM139769 (to M.K.K. and M.L.), R35GM133600 (to C.R.B and P.B.), and 1U19 AG056169-01A1, UH3 AG064706, and U19 AG023122 (to N.J.S.) as well as a Vallee Scholars Award to P.H.S. We also acknowledge financial support by the Deutsche Forschungsgemeinschaft (DFG, German Research Foundation) – 437857095, 444810852 (to T.L.L.), by Verne M. Willaman Endowment Professorship (to K.D.M.), and by the John and Donna Krenicki Endowment Professorship (to R.J.O.). Sequencing was partially supported by the NIH Intramural Sequencing Center. This work used the computational resources of the NIH HPC Biowulf cluster (https://hpc.nih.gov), the HPC in the Computational Biology Core in the Institute for Systems Genomics at UConn, Clemson University’s HPC Palmetto Cluster, and the HPC in the Genomics Institute at UCSC. RNA-seq sequencing was performed at PSU Genomics Core facility, and Hi-C sequencing was performed at the Genome Sciences core facility at the Penn State College of Medicine. This work was supported by the National Library of Medicine Training Program in Biomedical Informatics and Data Science (T15LM007093 to B.K.) and in part by the National Institute of Allergy and Infectious Diseases (P01-AI152999 to B.K). We acknowledge financial support under the National Recovery and Resilience Plan (NRRP), Mission 4, Component 2, Investment 1.1, Call for tender No. 104 published on 2.2.2022 by the Italian Ministry of University and Research (MUR), funded by the European Union – NextGenerationEU– Project Title Telomere-to-telomere sequencing: the new era of Centromere and neocentromere eVolution (CenVolution) – CUP H53D23003260006 - Grant Assignment Decree No. 1015 adopted on 07/07/2023 by the Italian MUR. – Project Title SUDWAY: Substance Use Disorders through Whole genome, psychological and neuro-endophenotypes AnalYsis CUP H53D2300331-Grant Assignment Decree No. 1015 adopted on 7 July 2023 by the Italian MUR. F.M. was supported by Fondazione con il Sud (2018-PDR-01136). We are grateful to the Frozen Zoo® at San Diego Zoological Society for providing the fibroblast cell lines. This work was supported by the Italian Ministry of University and Research (MUR) grant PRIN 2020 (project code 2020J84FAM, CUP H93C20000040001) to F.A. We thank the Genome in a Bottle Consortium for sharing preliminary RNA-seq data for HG002. E.E.E. and E.D.J. are investigators of the Howard Hughes Medical Institute. The work of DH, PM and FTN was supported by the National Center for Biotechnology Information of the National Library of Medicine (NLM), National Institutes of Health.

This article is subject to HHMI’s Open Access to Publications policy. HHMI lab heads have previously granted a nonexclusive CC BY 4.0 license to the public and a sublicensable license to HHMI in their research articles.

## AUTHOR CONTRIBUTIONS

Individual analysis leads are indicated with an asterisk. Lu.C., La.C., O.A.R., Cy.S., Ma.H., B.M., and K.D.M. managed sampling. B.M. and A.P.L. performed transcriptome data generation. K.H., G.G.B., S.Y.B., J.C. generated ONT long-read data. J.C., R.E.G. and Sa.S. provided Illumina sequencing data. G.H.G., K.M.M., P.H.S. and J.L.R. generated HiFi sequencing data. R.E.G. and Sa.S. made Hi-C libraries that were later sequenced by B.M. *B.D.P. and A.R. managed data, processed data submissions, and coordinated administrative tasks. W.T.H., J.W., A.R., and B.D.P. performed assembly QC. *D.A. and S.K. assembled the genomes. *A.R. performed polishing and created the genome browsers. Assembly generation was supervised by S.K. S.M. performed chromosome recognition and M.V. led definition of chromosome nomenclature. L.S., K.K. and K.D.M. analyzed non-B DNA. B.K., W.W., A.G., E.M., E.G., G.F. and P.H.S. created pangenome graph alignment. *G.H. generated the Cactus alignments. P.H.S., R.S.H., S.K.M., B.K., W.W., A.G., E.M., E.G., G.F. and K.D.M. performed divergence analysis. Co.S. and B.P. performed ancestral allele analysis. B.P., P.H., M.D., D.H., J.F.M., P.M., F.R.R., F.T., S.C. K.P., and K.D.M. analyzed and annotated genes. *P.H. annotated lineage-specific genes, integrated gene annotations and managed sharing of the annotation data. B.P. supervised gene annotation analysis. *F.M. and I.R. performed ILS analysis and predicted speciation times. *J.M.S., P.B., C.R.B., C.F., P.Z., G.A.H. and R.J.O. analyzed repeat content. E.T. and K.D.M. investigated NUMTs. M.L. and M.K.K. investigated specifically for species-specific MEIs. P.B. and C.R.B. performed ORF analysis on species-specific FL-L1s. R.J.O. supervised and integrated the results. *A.N.S., Ar.B., Q.L., M.C.S., M.G.T., Z.A.S., C.D.H., R.C.M., and K.D.M. analyzed population data and investigated selective sweeps. *Y.S., An.B., E.E., I.G., W.L., M.P., P.A.P., Sw.S., Z.Z., Yi.Z. and C.T.W. analyzed immunoglobulin loci. Y.S. and C.T.W. led the analysis and integrated the results. Jo.M., B.S.M., and T.L.L. performed annotation of MHC genes. P.H. supported and validated the MHC annotation. Mi.T., performed phylogenetic tree analysis across MHC loci. Y.E.L., D.R.S. and S.V.Y. analyzed epigenetic data focusing on methylation and gene expression. *J.M., M.L.Y., Y.Z. and T.S. analyzed replication timing. D.G. and T.S. generated Repli-seq data. *F.A., M.V. L.G., and D.Y. analyzed inversions and large-scale chromosome rearrangements. F.A., D.Y. and D.P. visualized the data. *J.L., J.H., S.Z. and Y.M. performed SDR analysis. A.P.C., Mi.H. and N.J.S. performed TOGA analysis. A.P.C. integrated the results. *Ya.L., *R.J.M., M.K., S.A.Z., C.B.L. analyzed divergent regions of the genome by predicting AQERs. C.B.L. supervised the analysis and summarized the results. C.L., Yo.L. and E.D.J. investigated further into candidate genes using AQER regions. *S.J.S., A.P.S., J.L.G., T.P., G.M.A. and M.B. analyzed acrocentric regions. T.P. and G.M.A. performed NOR chromosome imaging and quantification. S.J.S. and M.B. analyzed rDNA. A.P.S. generated dot plots. J.L.G., M.V. and A.M.P. supervised the analysis. *G.A.L., K.H.M. and H.L. investigated centromeres. G.A.L. integrated the section. *D.Y. and E.E.E. investigated subterminal heterochromatin. D.Y. performed the analyses and E.E.E. supervised the analysis. *D.Y. and E.E.E. analyzed SD. E.E.E. supervised the SD analysis. H.J. optimized the pipeline and D.P. and D.Y. visualized the data. P.H. analyzed novel genes and curated gene annotation across the region. E.E.E., D.Y., and A.M.P. wrote and edited the manuscript with input from all authors. E.E.E., A.M.P., and K.D.M. initiated and supervised the project, acquired the funding along with other senior authors. E.E.E. and A.M.P. coordinated the study.

