## Supplementary Note for "Complete sequencing of ape genomes"

#### Supplementary notes

### Table of Contents


| | |
|---|---|
| **I. Sequencing and samples** | **2** |
| **II. Genome assemblies** | **6** |
| **III. Non-B DNA annotations** | **17** |
| **IV. Genome alignment and sequence divergence** | **18** |
| **V. Sumatran vs. Bornean orangutan divergence** | **37** |
| **VI. Incomplete lineage sorting (ILS) and speciation times** | **39** |
| **VII. Gene annotation** | **47** |
| **VIII. Repeat annotation** | **56** |
| **IX. Selection analyses within NHP lineages** | **60** |
| **X. Immunoglobulin annotation and analysis** | **73** |
| **XI. MHC I and MHC II analyses** | **80** |
| **XII. Methylation** | **95** |
| **XIII. Replication timing** | **99** |
| **XIV. Evolutionary rearrangements and inversion characterization** | **104** |
| **XV. AQER** | **108** |
| **XVI. Structurally divergent regions** | **115** |
| **XVII. TOGA analysis** | **117** |
| **XVIII. Acrocentric region analysis** | **120** |
| **XIX. Centromere analyses** | **132** |
| **XX. Subterminal satellite** | **135** |
| **XXI. Segmental duplications** | **137** |

### I. Sequencing and samples

Contributing authors:

Katherine M. Munson, Kendra Hoekzema, Richard E. Green, Samuel Sacco, Gage H. Garcia, Gerard G. Bouffard, Shelise Y. Brooks, Juyun Crawford, David Gilbert, Takayo Sasaki, Lucia Carbone, Laura Carrel, Marlys Houck, Oliver A. Ryder, Cynthia Steiner, Alexandra P. Lewis, Barbara McGrath, Joana L. Rocha, Kateryna D. Makova

#### Methods

Sample selection and sequencing data were mostly reported in Makova et al, 2024[1]; briefly, including the whole-genome sequencing data (PacBio HiFi [high-fidelity] and ONT [Oxford Nanopore Technologies] long-read/Illumina short-read) derived from fibroblast cell lines of bonobo, gorilla, Bornean/Sumatran orangutans, and that of chimpanzee and siamang derived from lymphoblast cell lines, used directly in genome assembly. Moreover, with the parent–child trio available samples, including bonobo and gorilla, parental Illumina data were used for haplotype phasing. For the remaining samples, including chimpanzee, two orangutans and siamang, Hi-C data were used to perform haplotype phasing. Iso-Seq data from testes were used for gene annotation. On top of these data, additional data were also generated to assist in assembly and annotation of the autosomes, including 1) additional Iso-Seq/short-read RNA-seq data to assist annotation and 2) additional HiFi (bonobo lymphoblast cell line) and ONT (gorilla fibroblast, and bonobo lymphoblast cell lines) to improve genome assembly (**Table 1 & Table AssemblyS1**).

**PacBio Iso-Seq and RNA-seq Sequencing at Penn State University:** All male-derived cell lines were cultured, pelleted, and stored as described in Makova 2024[1]. Total RNA was isolated from approximately 5 million cells using the RNeasy Mini Kit (Qiagen) with on-column DNase digestion according to the manufacturer's protocol. RNA was eluted in nuclease-free water, snap-frozen in liquid nitrogen, and stored at -80°C while awaiting downstream analyses.

Uniquely indexed, short- and long-read SMRTbell templates were prepared for Iso-Seq (PacBio) transcriptome sequencing by the Huck Genomics Core Facility with the goal of achieving 3-4 million reads per SMRT cell. Size selection (~2 kbp and >3 kbp) was achieved by altering the volume of ProNex Beads after cDNA amplification. Samples were pooled in an equimolar pair-wise fashion and loaded onto SMRT cells for sequencing on the core facility's Sequel IIe System. Samples were pooled such that highly related species were not sequenced on the same flow cells.

For RNA-seq, uniquely indexed Illumina transcript libraries were prepared from cell line total RNA using the Illumina Stranded mRNA Prep Kit. An equimolar pool of all of the libraries was sequenced with a NextSeq 2000 P3; to achieve ~150 million pairs of 150 bp reads per sample (150 x 150 paired-end).

**PacBio Iso-Seq and Kinnex Sequencing at the University of Washington (UW):** Aliquots of the lymphoblast and fibroblast cell line RNA extracted by the Makova lab were shipped to the UW and prepared for PacBio Iso-Seq full-length transcriptome sequencing using the Iso-Seq Express Kit according to the manufacturer's protocol (PacBio, Preparing Iso-Seq Libraries using SMRTbell prep kit 3.0) with sample barcodes added during the cDNA PCR amplification step as in the protocol, and SMRTbell barcoded adapter plate 3.0 (PacBio P/N 102-009-200) used during library preparation with the SMRTbell prep kit 3.0 (PacBio P/N 102-182-700). Libraries were pooled in an equimolar fashion before sequencing on one SMRT Cell 8M on a PacBio Sequel II instrument using chemistry P3.1/C2.0 (PacBio P/Ns 102-333-400, 101-849-000).

Next, 5 ng of each cDNA were re-amplified for five cycles using barcoded primers compatible with a beta version of the Kinnex full-length RNA kit (PacBio P/N 103-072-000), then pooled in an equimolar fashion before Kinnex PCR and the remainder of the Kinnex full-length RNA protocol. The final library was sequenced on two SMRT Cell 25Ms on a PacBio Revio instrument using chemistry v1 and SMRT Link v12 (PacBio P/N 102-817-900).

After sequencing, data were analyzed using the Read Segmentation (Kinnex only) and Iso-Seq Analysis pipelines (Iso-Seq and Kinnex) in SMRT Link v12 to generate demultiplexed flnc.bam files.

**Table SequencingS1. Iso-Seq barcode**

| Sample Name | Iso-Seq cDNA barcode | Iso-Seq SPK3 ligation barcode | Kinnex cDNA barcode |
|---|---|---|---|
| PR00251_PPA_bonobo | bc1006 | bc2073 | IsoSeqX_bc01_5p |
| Jim_GGO_gorilla | bc1008 | bc2074 | IsoSeqX_bc02_5p |
| AG05252_PPY_Borang | bc1012 | bc2075 | IsoSeqX_bc03_5p |
| AG06213_PAB_Sorang | bc1018 | bc2076 | IsoSeqX_bc04_5p |
| AG18354_PTR_chimp | bc1019 | bc2077 | IsoSeqX_bc05_5p |
| Jambi_SSY_siamang | bc1020 | bc2078 | IsoSeqX_bc06_5p |

**Table SequencingS2. Summary of Iso-Seq data**

| Sample | Iso-seq | | RNA-seq | Total (Gbp) |
|---|---|---|---|---|
| | fibro/ lympho (Gbp) | Testes (Gbp) | fibro/ lympho (Gbp) | |
| Chimpanzee (PTR) | **35.7** (L) * | 10.86 | **48.25** (L) * | 94.81 |
| Bonobo (PPA) | **33.19** (F) * | 9.99 | **40.54** (F) * | 83.72 |
| Gorilla (GGO) | **35.6** (F) * | 12.44 | **40.76** (F) * | 88.8 |
| Bornean orangutan (PPY) | **34.31** (F) * | 7.38 | **45.83** (F) * | 87.52 |
| Sumatran orangutan (PAB) | **35.65** (F) * | 7.77 | **44.76** (F) * | 88.18 |
| Siamang (SSY) | **36.84** (L) * | - | **49.93** (L) * | 86.77 |
| Average | **35.21** | 9.69 | **45.01** | 89.92 |

*Newly generated sequencing data; (L) Lymphoblastoid cell line; (F) Fibroblast cell line.

**PacBio HiFi at University of California, Berkeley:** Lymphoblastoid cell lines available at the Coriell Institute for Medical Research were used for HiFi sequencing and expanded to a total culture size of 3×10e6 cells. The cell line expansions were derived from the original expansion culture to reduce the number of passages and minimize culturing time. Cells were washed in PBS and flash-frozen as dry cell pellets of 10e6 cells per vial. High-molecular weight (HMW) DNA was extracted on December 1-3, 2021, using the Circulomics CBB kit (102-573-600) from a frozen cell pellet (10e6 cells). DNA quantity, purity, and integrity were checked at different steps and at the end of the extraction protocol. DNA quantity was checked on a Qubit Fluorometer I with a dsDNA HS Assay kit (ThermoFisher) and sizes examined on a FEMTO pulse (Agilent Technologies) using a Genomic DNA 165 kb kit. Purity ratios were assessed with NanoDrop. A total of 54.8 micrograms of DNA (274 ng/uL in 200 uL volume, over 50 kbp length, and purity ratio 260/280: 1.82, 260/230:2.0) was used as input for library preparation.

A starting amount of 4-5 ug HMW gDNA was sheared to a target size of 20-30 kbp using a Megaruptor 3 instrument (Diagenode). The sheared DNA underwent size selection using a Pippin HT instrument (Sage Science) to target a size range of 15-22 kbp. Following size selection, the DNA was used for CCS (circular consensus sequencing) library preparation using the SMRTbell Express Template Prep Kit 2.0 and Enzyme Cleanup Kit 1.0 (PacBio). Each library was barcoded using PacBio Barcoded Overhang Adapters. Post-library preparation, the concentration of the DNA stock was measured using the DNA-HS Qubit assay, and the DNA size was estimated using the Fragment Analyzer or Femto Pulse. Sequencing was conducted on a PacBio Sequel IIe instrument, using version 2.0 sequencing reagents and operating on control software version 10.1.0.119549, with a movie collection time of 30 hours per 8M SMRT Cell.

**UL-ONT Sequencing at the University of Washington (PR00251_PPA & Jim_GGO):** Ultra-long-(UL-)ONT data were generated from the PR00251_PPA lymphoblast cell line and Jim_GGO fibroblast cell line according to a previously published protocol (Logsdon, protocols.io, 2020). Briefly, 3-5 x $10^7$ cells were lysed in a buffer containing 10 mM Tris-Cl (pH 8.0), 0.1 M EDTA (pH 8.0), 0.5% w/v SDS, and 20 mg/mL RNase A (Qiagen, 19101) for 1 hour at 37°C. 200 ug/mL Proteinase K (Qiagen, 19131) was added, and the solution was incubated at 50°C for 2 hours. DNA was purified via two rounds of 25:24:1 phenol-chloroform-isoamyl alcohol extraction followed by ethanol precipitation. Precipitated DNA was solubilized in 10 mM Tris (pH 8.0) containing 0.02% Triton X-100 at 4°C for two days. Libraries were constructed using the Ultra-Long DNA Sequencing Kit (ONT, SQK-ULK001) with modifications to the manufacturer's protocol. Specifically, ~40 ug of DNA was mixed with FRA enzyme and FDB buffer as described in the protocol and incubated for 5 minutes at RT, followed by a 5-minute heat-inactivation at 75°C. RAP enzyme was mixed with the DNA solution and incubated at RT for 1 hour before the clean-up step. Clean-up was performed using the Nanobind UL Library Prep Kit (Circulomics, NB-900-601-01) and eluted in 225 uL EB. 75 uL of library was loaded onto a primed FLO-PRO002 R9.4.1 flow cell for sequencing on the PromethION, with two nuclease washes and reloads after 24 and 48 hours of sequencing.

### II. Genome assemblies

Contributing authors:

Sergey Koren, Brandon Pickett, Arang Rhie, Dmitry Antipov, Julie Wertz, William T. Harvey, Sean McKinney, Mario Ventura, Adam M. Phillippy

##### Methods

The complete, haplotype-resolved assemblies were generated using a combination of Verkko[2] and expert manual curation. Parental-specific markers were generated using Merqury[3] with the commands:

```
cd maternal
ls *.fastq.gz > input.fofn
sh _submit_build.sh -c 30 input.fofn maternal
cd ../paternal
ls *.fastq.gz > input.fofn
sh _submit_build.sh -c 30 input.fofn paternal
cd ..
sh trio/hapmers.sh maternal/maternal.k30.meryl paternal/paternal.k30.meryl
```

Verkko v1.4.1 was run with the parameters --screen human and --trio maternal.hapmer.meryl paternal.hapmer.meryl for trios (bonobo and gorilla) using the *k*-mer databases built above or --hic1 *R1*fastq.gz --hic2 *R2*fastq.gz in the absence of trios (chimpanzee, orangutans, and siamang). Haplotype-consistent contigs and scaffolds were automatically extracted from the labeled Verkko graph, with unresolved gap sizes estimated directly from the graph structure (see Rautiainen et al.[2] for more details).

After the assembly was generated, manual interventions were employed to complete the Assembly. The ONT reads were re-aligned to the final graph using GraphAligner v1.0.17[4] with the command:

```
GraphAligner -t <cores> -g unitig-unrolled-unitig-unrolled-popped-unitig-normal-connected-tip.gfa -f split/ont<jobid>.fasta.gz -a aligned<jobid>.gaf --seeds-mxm-windowsize 5000 --seeds-mxm-length 30 --seeds-mem-count 10000 --bandwidth 15 --multimap-score-fraction 0.99 --precise-clipping 0.85 --min-alignment-score 5000 --hpc-collapse-reads --discard-cigar --clip-ambiguous-ends 100 --overlap-incompatible-cutoff 0.15 --max-trace-count 5 --mem-index-no-wavelet-tree
```

Using available information, such as parent-specific *k*-mer counts, depth of coverage, and node lengths, some artifactual edges could be removed and simple nonlinear structures resolved. For more complex cases, ONT reads aligned through the graph were used to select candidate resolutions consistent with the majority of alignments. In cases of unclear parental inheritance, nodes were arbitrarily assigned to a haplotype to avoid introducing gaps. Gaps filled by fewer candidate ONT sequences than Verkko's default of three were also patched at this stage after confirming the orientation and association of nodes via Hi-C links. The gap-filling sequences were added to the assembly graph as nodes using the same insert_aln_gaps.py script used internally by Verkko with parameters (minimum read support, distance from contig end) adjusted as appropriate to ensure a fill for each gap. A separate process was used to include assembly of the regions close to ribosomal DNA (rDNA) arrays. Initially, for each path ending at an rDNA sequence, the last reliable node present in only one path was selected. This was done manually, with the help of length, coverage, and graph structure. Then, Verkko's paths were extended with the help of improve_gaps_ont.py script. This script starts with unique nodes and extracts a set of ONT read alignments *S* that contain this node. Then, it iteratively adds the node most supported by alignments from *S*, if it is supported by 1.6-fold more reads than the second best, and the total number of supporting alignments is at least four. Graph structures which could not be resolved using the above methods were left as gaps in the assembly.

Once the paths for each chromosome were complete, Verkko was re-run to generate a new consensus with the commands:

```
cp  ../asm/6-layoutContigs/combined-alignments.gaf ./
cat ../asm/8-resolve/toalign_combined.gaf >> combined-alignments.gaf

cp  ../asm/6-layoutContigs/combined-edges.gfa ./
cat ../asm/8-resolve/gapfill_combined.gfa | grep '^L' >> combined-edges.gfa

ln -s ../asm/6-layoutContigs/combined-nodemap.txt

cp ../asm/6-layoutContigs/nodelens.txt ./
cat  ../asm/8-resolve/gapfill_combined.gfa | awk 'BEGIN \
     {
       FS="[ \t]+"; OFS="\t"; \
     } \
     ($1 == "S") && ($3 != "*") \
     { \
       print $2, length($3); \
     }' >> nodelens.txt

cat ../asm/8-resolve/gapfill_combined.gaf |awk '{print $1}' > ont.ids
```

```
grep -w -v -f ../asm/7-consensus/ont_subset.id tmp > ont.ids.extra
cp ../asm/7-consensus/ont_subset.fasta.gz ./
cat `ls ../asm/8-resolve/gapfill*fasta|grep -v hpc` | seqtk subseq - ont.ids.extra |pigz -c >> ont_subset.fasta.gz

<path to verkko>/src/scripts/get_layout_from_mbg.py combined-nodemap.txt combined-edges.gfa combined-alignments.gaf rukki.paths_with_short_arms.gaf nodelens.txt unitig-popped.layout unitig-popped.layout.scfmap

<path to verkko>/lib/verkko/bin/layoutToPackage -layout unassigned-unitig.layout -output packages/part###.cnspack -idmap  packages -partition 0.8 1.5 0.04 -reads ont_subset.fasta.gz <hifi reads>  > packages.report touch packages.finished

# for each package in parallel
<path to verkko>/lib/verkko/bin/utgcns -V -V -V -import packages/part<jobid>.cnspack -A packages/part<jobid>.fasta -C 2 -norealign -maxcoverage 50 -EM $MAXONT -e 0.05 -em 0.20 -l 3000 -threads 32 -edlib

<path to verkko>/lib/verkko/scripts/fasta_combine.py combine combined.fasta packages.tigName_to_ID.map  unitig-popped.layout.scfmap  packages/part*.fasta
```

Lastly, short sequences, EBV, and mitochondrial sequences were identified using human reference and the Verkko screen-assembly.pl script with the identity threshold set to 90%. EBV was identified in bonobo, chimpanzee, and siamang.

Short arms of acrocentric chromosomes were scaffolded to the remaining chromosomes with Hi-C reads. Initially, potential short and long arms were discovered using length and proximity to rDNA containing nodes. Then, for each short arm *A*, we selected the corresponding long arm as the arm with the highest total number of Hi-C links connecting its nodes to *A*. Details of the algorithm are described in the Verkko 2.0 paper (Antipov et al.). The assembly was versioned as v1.4.1r and was subject for polishing and curation.

The initial assemblies were polished by adapting the process previously described in Mc Cartney et al.[5] and Rhie et al.[6] Briefly, short nucleotide variation (SNV) like errors were called from short and long reads with DeepVariant and filtered for correcting consensus and phasing errors using BCFtools and Merfin[7]. Pre- and post-polished assemblies were evaluated with *k*-mers from HiFi and Illumina reads with Merqury, along with read-coverage-based analysis using scripts from Mc Cartney et al.[5] Systematic errors were found close to the rDNA gap flanking sequences and were manually patched afterwards.

**Read mapping**

Unlike the X and Y chromosomes, the autosomes share more homology, making it difficult to uniquely map reads to the proper haplotype. This becomes challenging when phasing error persists in the underlying consensus, especially when collapsed haplotype errors exist in long stretches of a nearly homozygous region. Therefore, we added one more error type to target, in addition to the classic consensus errors—the phasing error.

First, HiFi and ONT read sets were aligned to the diploid genome (all-to-dip) as well as to each haploid genome (all-to-hap) with Winnowmap v2.03 using the pipeline from T2T-Polish (https://github.com/arangrhie/T2T-Polish/tree/master/winnowmap). The X was included to the paternal and Y to the maternal haploid genome. In brief, the top 0.02% repetitive 15-mers were collected using Meryl:

```
meryl count k=15 $ref output merylDB
meryl print greater-than distinct=0.9998 merylDB > repetitive_k15.txt
```

and mapped with Winnowmap and sorted, filtered for primary alignments with option -ax map-pb for HiFi and -ax map-ont for ONT reads:

```
winnowmap --MD -W repetitive_k15.txt -ax $map -I12g -t$cpus $ref $reads >
$tmp/$out.sam
samtools sort -@$cpus -m2G -T $tmp/$out.tmp -O bam -o $out.sort.bam $tmp/$out.sam
samtools view -F0x104 -@$cpus -hb $out.sort.bam > $out.pri.bam
```

The three reference versions (all-to-dip and two haplotypes for all-to-hap) were indexed for Illumina read mapping with BWA v0.7.17 (https://github.com/arangrhie/T2T-Polish/tree/master/bwa).

```
bwa index $ref
```

Each paired set of fastq files were provided for alignment, with duplicates removed with fixmate and SAMtools v1.17.

```
bwa mem -t $cpu $ref $r1 $r2 > $tmp/$out.sam
samtools fixmate -m -@$cpu $tmp/$out.sam $tmp/$out.fix.bam
samtools sort -@$cpu -O bam -o $out.bam -T $tmp/$out.tmp $tmp/$out.fix.bam
samtools index $out.bam
samtools markdup -r -@$cpu $out.bam $out.dedup.bam
samtools index $out.dedup.bam
```

The bam files were merged at the end for variant calling.

```
samtools merge -@ $cpu -O bam -b $lst $out.bam
samtools index $out.bam
```

**Variant calling**

Once the read alignment finished, Illumina and HiFi read alignments were used for variant calling with DeepVariant Hybrid mode (https://github.com/arangrhie/T2T-Polish/blob/master/deepvariant/_submit_mrg_hybrid_dv.sh). DeepVariant v1.5.0 was used in default mode.

```
samtools merge -@$cpu -O bam -o $BAM_HYBR $BAM_HIFI $BAM_ILMN
samtools index $BAM_HYBR
```

DeepVariant step 1 - make examples. For the all-to-dip alignment, MQ filter was lowered to 1 to include read alignments in the more homozygous region.

```
# for all-to-hap alignments
extra_args=""
# for all-to-dip alignments
extra_args="--min_mapping_quality 1"

seq 0 $((N_SHARD-1)) \
  | parallel -j ${SLURM_CPUS_PER_TASK} --eta --halt 2 \
  --joblog "logs/log" --res "logs" \
  make_examples    \
    --mode calling \
    --ref "${REF}" \
    --reads "${BAM}" \
    --examples $OUT/tfrecord@${N_SHARD}.gz $extra_args \
    --sample_name "${SAMPLE}" \
    --task {}
```

step 2 - call variants (this step was run on gpu nodes).

```
call_variants \
  --outfile "${CALL_VARIANTS_OUTPUT}" \
  --examples "$OUT/examples/tfrecord@${N_SHARD}.gz" \
  --checkpoint /opt/models/hybrid_pacbio_illumina/model.ckpt
```

step 3 - post process variants

```
postprocess_variants \
  --ref "${REF}" \
  --infile "${CALL_VARIANTS_OUTPUT}" \
  --outfile "$OUT/$OUT.vcf.gz"
```

For the ONT read alignments, PEPPER-MARGIN-DeepVariant v0.8 was used, as DeepVariant v1.5.0 was not available for R9 data. Similar to the hybrid mode, all-to-dip alignments were processed with mapping quality options --pepper_min_mapq 1 --dv_min_mapping_quality 1. For faster processing, this step was performed on each chromosome and the resulting VCF file was merged at the end with BCFtools v1.17.

```
# Per-chromosome
run_pepper_margin_deepvariant call_variant \
  -b $BAM -f $REF -o $OUTPUT_DIR \
  $MQ_OPT \
  -t $THREADS -r $REGION --ont_r9_guppy5_sup --gpu

# At the end, merge all per-chr VCFs
ls ${in}*/PEPPER_MARGIN_DEEPVARIANT_FINAL_OUTPUT.vcf.gz >
r9_files_to_mrg.list
bcftools concat -D -a --threads $cpus --no-version -Oz -o $out.vcf.gz -f
r9_files_to_mrg.list
bcftools index $out.vcf.gz
vcf_stats_report --input_vcf $out.vcf.gz --outfile_base $out
```

**SNV correction**

Variants were filtered with BCFtools v1.17 and Merfin v1.1 using *k*-mers from Illumina and HiFi. The overall diagram of the filtering is in **Fig. AssemblyS1**. A custom script was made to run this filtering (https://github.com/arangrhie/T2T-Polish/blob/master/variant_call/snv_candidates.sh) and applied to each genome.

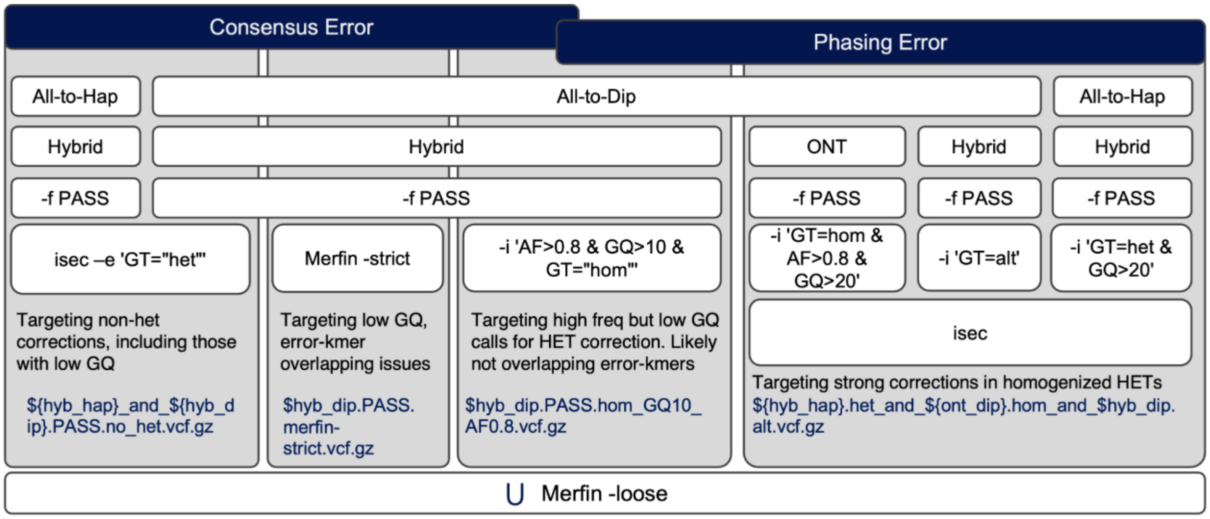


**Figure AssemblyS1. SNV-like error filtering criteria.** All-to-Hap: Reads to each haplotype alignment. All-to-Dip: Reads to diploid (both haplotype) alignments. Hybrid: Merged HiFi and Illumina read alignments. U: Union. The rest are BCFtools and Merfin options to specify filtering conditions.

The final VCF file, snv_candidates.merfin-loose.vcf.gz, was used to make an internal 20231003 release (v1.4.2) with BCFtools consensus:

```
bcftools consensus -H1 --chain v1.4.1r_to_v1.4.2r.chain -f ${sp}_v1.4.1r.analysis-dip.fa
snv_candidates.merfin-loose.vcf.gz > ${sp}_v1.4.2r.analysis-dip.fa
```

#### Structural variant (SV) correction

While evaluating corrections, we identified regions flanking rDNA or gaps that contained clusters of needed corrections, which is difficult to properly correct with variant calling based polishing approaches. For correcting these errors, we revisited the assembly graph and manually created patch consensus to replace those regions. We primarily targeted regions around the rDNA, subtelomeric regions missing telomeres, and a few other regions flagged by the coverage-based analysis (described in the evaluation section).

Approximate regions containing the rDNA were identified in v1.4.1r assembly using a canonical human version of the 45S sequence[8] with MashMap3 v.3.1.1:

```
mashmap  -t $cpu --noSplit -q human_45S.fa \
  -r $sp_ver.analysis-dip.fa -s 13332 --pi 85 -f none -o 45S_to_$asm.mashmap.out

cat 45S_to_$asm.mashmap.out |\
  awk -v OFS='\t' '{print $6, $8, $9, $1, $(NF-1), $5}' |\
  awk -F ":" '{print $1"\t"$3}' |\
  awk -v OFS='\t' '{print $1, $2,$3, "45S", (100*$6), $7}' \
```

```
> 45S_to_$asm.mashmap.bed
```

We identified an issue with Verkko consensus mis-assigning in regions with high-coverage repeats represented as gaps in the assembly. Once fixed, the paths were regenerated as described above to generate patch sequences. The patch sequences were trimmed down to contain a maximum of 500 kbp sequence and aligned to the target chromosome in the v1.4.1r version with wfmash. Alignments were compared to the SNV correction candidates and error *k*-mers from the hybrid *k*-mer database (described in the evaluation section). Patch sequences were further narrowed down manually to only contain target regions with less or no *k*-mers flagged as errors. Gaps flanking with rDNA have been resized to 1 Mbp in the new patch sequences.

wfmash v0.10.4 was run as follows:

```
wfmash --no-split -ad -t24 ${sp}_${ver}.$chr_hap.fa \
    patch.fa -s50000 -p95 > $out.sam
```

Once the target region to patch in v1.4.1 coordinates were identified, the corresponding query sequence was determined from the wfmash alignments. If the patch sequences had to be split at the gap, for later adjusting the gap size, both sides were considered to make one patch sequence (clip) and grouped. A custom script (https://github.com/arangrhie/T2T-Polish/blob/master/patch/samRefPos2QryPos.jar) was used to retrieve the corresponding query position from a given reference position:

```
if [[ $telo == "clip" ]]; then
  target1=`echo $target | awk -F "|" '{print $1}'`
  samtools view ${chr_hap}_fix_to_$ver.bam |\
    java -jar -Xmx1g samRefPos2QryPos.jar - $target1 > $chr_hap.patch.bed
  target2=`echo $target | awk -F "|" '{print $2}'`
  samtools view ${chr_hap}_fix_to_$ver.bam |\
    java -jar -Xmx1g samRefPos2QryPos.jar - $target2 >> $chr_hap.patch.bed
else
  samtools view ${chr_hap}_fix_to_$ver.bam |\
    java -jar -Xmx1g samRefPos2QryPos.jar - $target > $chr_hap.patch.bed
fi
```

Patch replacements were created in VCF format to contain target sequence in the REF field and the new patch sequence in ALT field:

```
## Per $chr_hap vcf
chr=`head -n1 $chr_hap.patch.bed | awk '{print $1}'`
pos=`head -n1 $chr_hap.patch.bed | awk '{print ($2+1)}'`
ref=`cat ref.seq`
alt=`cat qry.seq`
cat $sp_ver.header.vcf > $sp_ver.$chr_hap.rDNA_patch.vcf
echo -e "$chr\t$pos\t.\t$ref\t$alt\t1\t.\t.\tGT\t1/1" >> $sp_ver.$chr_hap.rDNA_patch.vcf
```

Missing telomeric sequences at the end of chromosomes were found using the 'telo -d 10000' function of seqtk v1.3, and patch sequences were made in the same way as the rDNA locus patches.

Lastly, SNV correction candidates overlapping the target region for patching were removed, and merged with the patch sequences. The resulting VCF was re-applied to the v1.4.1r version with BCFtools consensus as follows:

```
bcftools consensus -c $sp.${OLD}_to_$NEW.chain \
  -f ../$sp/$sp_ver.analysis-dip.fa -HA \
  $sp_ver.snv_sv_edits.vcf.gz > $sp_new.analysis-dip.fa
```

Scripts used for rDNA patch and telomere patch VCFs are available on https://github.com/arangrhie/T2T-Polish/tree/master/patch, make_rDNA_patch.sh and make_telo_patch.sh. Script for excluding SV edits and merging the SNV edits to make the final consensus is available as merge_snv.sh.

##### Haplotype assignment, chromosome orientation, and numbering

Chromosome numbers and orientation were identified using prior markers established for each species. Sequences were renamed to contain the species-specific chromosome number and human ortholog number and were reversed accordingly to have the p-arms at the beginning. For the non-trio species, haplotype numbers were reassigned to keep the rDNA containing haplotype as haplotype 1 if the partnering chromosome had no rDNA, or the more continuous (telomeres found on both ends), less gaps, less errors assigned as haplotype 1. For sex chromosomes, chrX was always assigned to haplotype 1, and chrY as haplotype 2, respectively. Haplotype 1 assemblies and the Y chromosome were regrouped as the primary assembly set for convenience if a linear representation of the species was needed. For the gorilla and chimpanzee, which had the parental information available, haplotypes were assigned as mat or pat for maternal or paternal, respectively. Using the same criteria for choosing haplotype 1 and the primary haplotype in the non-trios, the haplotypes were reassigned to keep the more continuous, accurate haplotype in the primary assembly set.

##### Assessment of the genome assembly

Alignments were generated by mapping reads onto each assembly using the assembly_eval pipeline (https://github.com/EichlerLab/assembly_eval). Minimap2 (v2.26) was used for the alignment with standard parameters.

**Flagger** (v0.3.3) (https://github.com/mobinasri/flagger) was run using the alignment file containing HiFi reads mapped back to each assembly. The pipeline was run using flagger_end_to_end_with_mapping.wdl file deposited in the repository.

**NucFreq** was run using HiFi reads as a part of the assembly_eval pipeline (https://github.com/EichlerLab/assembly_eval). Collapses are defined as locations where the second most common base had a depth of coverage greater than 5, and Duplicated/HiFi-deplete are defined as regions of the assembly with 0 or decreased read coverage by HiFi reads.

QV was estimated using Meryl v1.4.1 and Merqury commit 01a39a6 using a hybrid database of 31-mers collected from Illumina and HiFi reads. Hybrid databases were made as described in https://github.com/arangrhie/T2T-Polish/tree/master/merqury. Switch error rate was estimated using parental Illumina 31-mers, if available.

#### Chromosome nomenclature

To accurately determine proper nomenclature for chromosomes based on the centromeric position, we employed three approaches: a ratio-based approach comparing the lengths of the long (l) and short (s) arms in base pairs (r=l/s), the centromeric index-based approach comparing short arm to the total chromosome length (c) (I=100s*c) and the centromeric index-based approach using the total chromosome length without the centromeric sequence length (c') (I'=100s*c') as per the classical definitions outlined by the Denver Study Group in 1960[9] and by Levan et al. in 1964[10]. To integrate new sequence data with traditional cytogenetic information to best define the arms lengths, the centromeric regions had to be excluded from the total chromosome length calculations for both the long and short arms. Traditionally, the centromere was not included in size measurements due to its over-condensed nature; rather, it was solely identified along the chromosomes. Consequently, we focused solely on the lengths of the q and p arms, thereby circumventing also the highly variable centromeric sequences present in each chromosome.

Additionally, we used the above-mentioned methods on human chromosomes and find the ratio criteria was the most reliable using the ranges: $0,7<r<1.58$ for metacentric (M), $1.58<r<3$ for submetacentric (S) and $r>3$ for acrocentric (A).

Employing this methodology (**Fig. AssemblyS2** and **Table AssemblyS4-9**), we successfully categorized each chromosome and identified the acrocentric chromosomes in chimpanzees, bonobos, gorillas, and orangutans (red arrows).

To avoid confusion with nomenclature, we referred to great ape chromosomes considering the human synteny instead of species-specific nomenclature. For example, PTR chromosome 2, being homologous to human chromosome 3, was assigned as chromosome 3 (III in Roman numberals in **Fig. AssemblyS2**). The only exception to this is chromosome 2 where we used 2A and 2B for 2p-2q and 2q chromosome syntenic blocks, respectively.

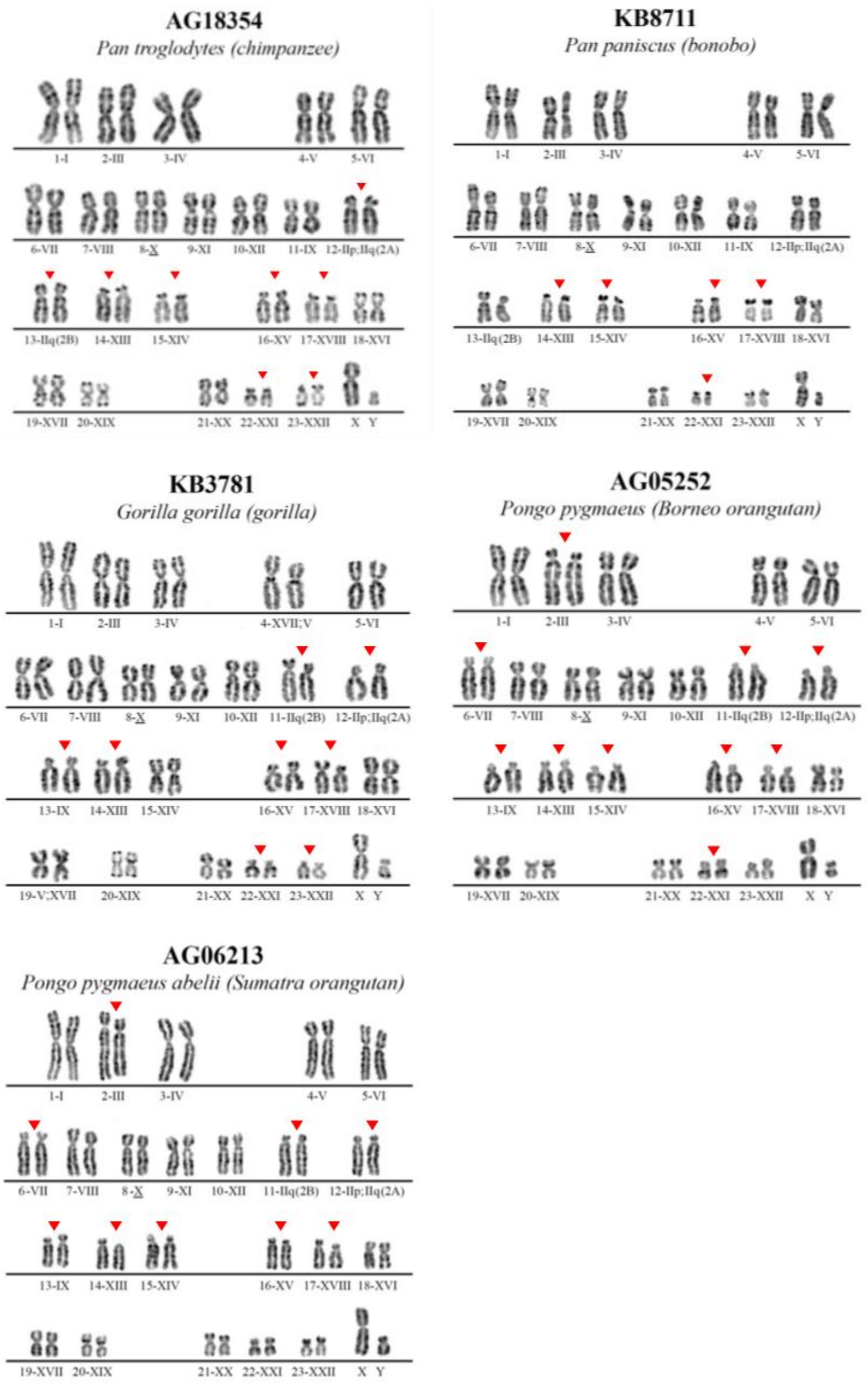


**Figure AssemblyS2. Karyotyping of the ape tissues.**

### III. Non-B DNA annotations

Contributing authors:

Kaivan Kamali, Linnéa Smeds, Edmundo Torres-González, Kateryna Makova

##### Methods

Non-B DNA motifs were annotated in each species using gfa (https://github.com/abcsFrederick/non-B_gfa). We used default settings, except the flag --*skipGQ*, to annotate A-phased repeats, direct repeats, inverted repeats, mirror repeats, short tandem repeats, and Z-DNA. G-quadruplexes were annotated using Quadron[11]. For each motif type, the output was converted to bed format, and mergeBed from BEDTools[12] was used to merge any overlapping annotations.

##### Alignments to old assembly versions

For each species that had a corresponding older non-T2T assembly, the new T2T assembly was aligned to its predecessor (panPan3 for bonobo, panTro6 for chimpanzee, gorGor6 for gorilla and ponAbe3 for Sumatran orangutan; Bornean orangutan and siamang were excluded from this analysis because they were sequenced here for the first time and thus their previous assemblies are unavailable). Each chromosome pair (new vs. old) was aligned separately with LASTZ[13], using the flags --*format=general:name1,size1,zstart1,end1,name2,strand2,zstart2+,end2+,nmatch,nmismatch,cgap,score,id%,blastid%,con%  --progress --ambiguous=iupac*. To assess regions in the T2T assemblies that were previously unassembled, we merged all the aligned sequences, and then extracted everything that did *not* align to the previous assembly using BEDTools complement. This is from now on referred to as the 'new' sequence.

##### Enrichment of non-B DNA in new sequence

To investigate the density of non-B motif annotations in new versus previously assembled sequence, we intersected the non-B annotations of each motif type separately with the new regions defined above, as well as to the previously assembled regions, using BEDTools intersect and the flag -wao that outputs the number of overlapping base pairs. The overlaps with new and previous sequence were summed up for each motif type separately, and the fold enrichment was calculated as non-B density in the new sequence divided by non-B density in previously assembled sequence.

### IV. Genome alignment and sequence divergence

Contributing authors:

Peter H. Sudmant, Giulio Formenti, Erin K. Molloy, Wenjie Wei, Andrea Guarracino, Bryce Kille, Erik Garrison, Wenjie Wei, Cole Shanks, Prajna Hebbar, Glenn Hickey, Benedict Paten

#### Methods

##### Pangenome alignment

To perform an all-vs-all alignment across the entire set of primate genomes, we applied an iterative all-vs-one approach using wfmash (https://github.com/waveygang/wfmash). We mapped all genomes against a single target genome at a time, repeating this process for each genome in the dataset as the target. This strategy breaks down the computationally intensive all-vs-all alignment into more manageable all-vs-one steps, enabling efficient parallel processing. For the mapping step, we used MashMap3 (https://github.com/marbl/MashMap), integrated into wfmash. Each iteration works by splitting each query genome into overlapping 5 kbp segments and mapping each segment to the current target. The mappings are then filtered to keep only those with >70% identity, merging those closer than 20 kbp. Next, we used the mappings as input to guide the alignment process in wfmash. This results in aligned PAF files for each target sequence that include base-level alignment as CIGAR strings (**Fig. PanGenomeS1a**). Finally, we used paf2chain (https://github.com/AndreaGuarracino/paf2chain) to convert the aligned PAF files into CHAIN format. We repeated this procedure to map and align the HPRC year 1 human genome assemblies against each primate target genome. This allows comparative analyses between the primate genomes and the diverse set of human haplotypes.

##### Implicit pangenome graph (impg)

Pangenome graphs condense many-way sequence relationships (similarities) into a graphical model that avoids redundancy. These graphs imply alignments: in principle you can extract alignments back out of the graph, which exactly encompass the relationships between individual genomes seen in the graph. And the reverse is true as well: a set of alignments between sequences imply a graph.

A common technique in algorithms is to avoid a hard problem by simulating its result. For instance, interval trees are data structures used to index and query range overlaps. They are used in BEDTools and alignment algorithms. But they are very heavy to instantiate literally. In practice, it is always better to use an implicit interval tree[14]. We build on this same data structure to create an implicit pangenome graph out of a set of base-level alignments between all pairs of haplotypes in the T2T-primates collection.

The implicit pangenome graph lets us produce specific products and query the pangenome without instantiating it. These include:

- Graph subset: using a compressed index of the alignments, the impg tool lets us extract all subsequences of genomes that match any locus on any genome in the pangenome. This matching is transitive and exactly equivalent to what would be obtained from subset operations on a full graph build. (Here we show the MHC and 8p23.1 inversion extracted from the implicit graph using this technique.)
- Reference-relative multiple alignments in a "pseudo-MAF" format usable by many downstream comparative genomics tools.
- Divergence: also from MAFs / flip side of conservation.
- Conservation: from the MAFs we build a track of conservation relative to each genome in the cohort.

Our alignment and pangenome analysis approach has some advantages over current standards (e.g., Cactus) in being tree-free, repeat-masker free, capable of considering all haplotypes, and easy to parallelize. The approach is fundamentally pangenomic in that we make results in any frame of reference.

**Subgraph analysis from implicit graph**

To efficiently extract subgraphs and analyze the primate pangenome without constructing a full graph, we used impg (implicit pangenome graph). impg is a tool that projects sequence ranges through many-way pairwise alignments, such as those built by wfmash and minimap2, to identify homologous loci across multiple genomes.

impg uses coitrees (implicit interval trees) to provide efficient range lookup over the input alignments. CIGAR strings are converted to a compact delta encoding, enabling fast and memory-efficient projection of sequence ranges through alignments. The output is provided in BED, BEDPE, and PAF formats, making it straightforward to extract FASTA sequences for downstream use in multiple sequence alignment or pangenome graph building.

To build the impg index, we first generated a compressed index of the wfmash alignments as a tiny auxiliary file that indexes a bgzipped PAF file. We then used impg to extract sequences corresponding to specific loci of interest, such as the MHC and 8p23.1 inversion regions (**Fig. PanGenomeS1**), from the primate genomes.

The resulting sequences were then used for downstream analyses, such as building local subgraphs with pggb and visualizing them with odgi, as described in the previous sections. This approach allowed us to efficiently analyze specific regions of interest in the primate pangenome without the need to build and manipulate a full pangenome variation graph. Future work will enable distributed construction of large comparative genomic type pangenome graphs but is here applied in an exploratory way.

By using impg to extract homologous sequences and build local subgraphs, we were able to perform detailed comparative analyses of specific genomic regions across the primate genomes in a computationally efficient manner. This approach demonstrates the utility of impg for managing and analyzing large collections of genomes in a pangenomic context.

##### 8p23.1 subgraph generation

Human 8p23.1 is interesting due to its defensin gene content and high polymorphism in humans and across the primate lineage. To generate a subgraph of the 8p23.1 locus across multiple primate genomes, we first extracted the locus with flanking sequence from the human GRCh38 reference genome using coordinates grch38#1#chr8:5748405-13676927. We then used a wfmash whole-genome alignment of primate genomes to identify syntenic regions in the other primate genomes. The wfmash alignments were processed into an implicit graph (impg) data structure. We used impg to collect the sequences corresponding to the 8p23.1 locus from each primate genome:

```
impg -p primates16.20231205.paf.gz -t 64 -r grch38#1#chr8:5748405-13676927 -x | bedtools sort | bedtools merge -d 1000000 | awk '$3 - $2 > 2000000 { print $1":"$2"-"$3 }' > primates.8p23.1.merged-1m.gt2m.regions
```

```
samtools faidx -r primates.8p23.1.merged-1m.gt2m.regions ../primates16.20231205.fa.gz | bgzip -@24 -l9 >../primates16.20231205.8p23.1.merged-1m.gt2m.fa.gz
```

These sequences were then built into a local sequence graph using pggb with the following parameters: -c 10 -t 96 -p 90 -s 10k -k 19. -c 10 forces a high rate of multi-mapping between homologous sequences, which we found necessary to build a compact representation of the repetitive locus that modulates the recurrent polymorphic inversion.

To aid visualization in 1D, the graph was sorted using odgi sort, taking the chm13 sequence path as the reference order, with the following parameters: -t 96 -p Y -H <(odgi paths -i ${graph} -L | grep chm13) -x 500 -P -K 0.5 -v 10000000000000 -g 0.0000000000000001

The sorted graph was visualized using odgi viz (**Fig. PanGenomeS1b**) to generate a rendering showing the graph topology colored by path inversion status relative to the chm13 order and a rendering showing the graph topology colored by local graph depth.

##### MHC locus subgraph generation

To generate a subgraph of the MHC locus across multiple primate genomes, we first extracted the locus with flanking sequence from the chm13 reference genome using coordinates chm13#1#chr6:28385000-33300000. We then used the wfmash whole-genome alignment of primate genomes to identify syntenic regions in the other primate genomes. The wfmash alignments were processed into an implicit graph (impg) data structure using the following command:

```
impg -p primates16.20231205_wfmash-v0.12.5/chm13\#1.aln.paf -q chm13#1#chr6:28385000-33300000 | tee mhc.bed
```

The resulting sequences were then built into a local sequence graph using pggb with the following parameters:

> -i /lizardfs/erikg/primates/primates16.20231205.mhc.impg.merge-500kb.fa.gz -o mhc.primates.'$i' -t 96 -p 90 -s 10k -k 47

The resulting graph was visualized using odgi (**Fig. PanGenomeS1i-l**). The graph was sorted and visualized as described for the 8p23.1 locus above to generate renderings showing the graph topology colored by path inversion status and by local graph depth.

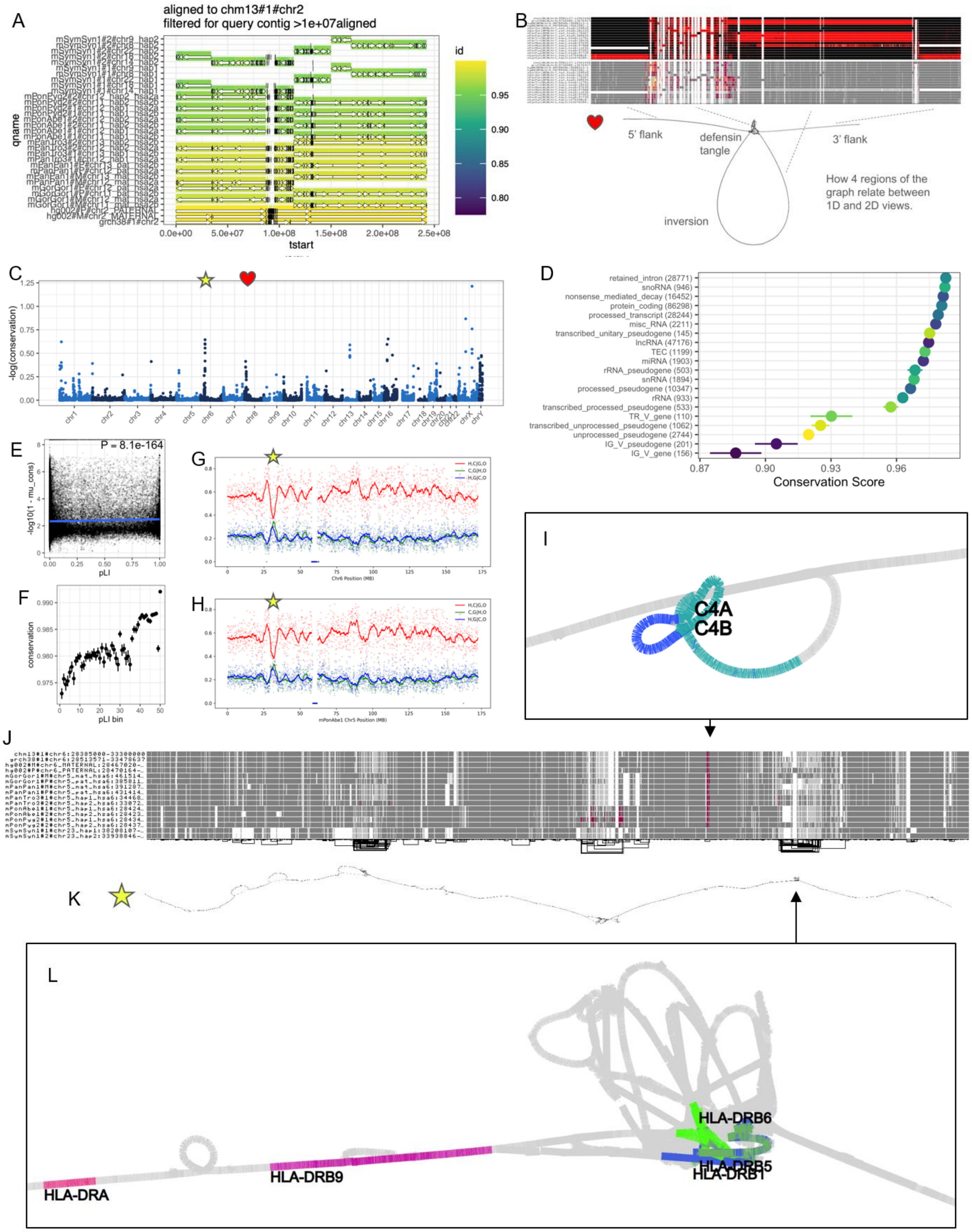


**Figure PanGenomeS1. Analyses on the implicit pangenome graph.** Analyses are based on a 16-way alignment of all T2T assemblies in the present study, CHM13, GRCh38, and HG002 to all others. Each genome serves as a reference, allowing for universal annotation of conservation and across all assemblies. (A) Alignments show the fusion of acrocentric chromosomes to form chromosome 2 in the human lineage. (B) Extraction of the pangenome graph around an inversion located on human 8p23.1. The inversion is polymorphic across great ape clade, as shown by visualizations with odgi viz (top, with black and red indicating relative orientation in the graph). (C) Conservation scores relative to CHM13, showing a drop in conservation at the MHC (star) and also in the 8p23.1 locus (heart). Higher values indicate lower conservation. (D) Conservation scores relative to gene classes shows protein-coding related sequences as the most conserved, while lncRNA, rRNA, immune, and pseudogenes are the among least conserved classes. (E) Conservation scores plotted versus probability of loss-of-function intolerance (pLI) scores. All genes are plotted. (F) Genes are binned into 50 bins by pLI score. (G-H) Quartet based analysis of phylogenies relative to 100 kbp bins of either human chromosome 6 from CHM13 (G) or chromosome 5 (hsa6) of mPonAbe1 (H) show that, broadly, human clusters with chimpanzee (H,C|G,O), while a peak of the (H,G|C,O) phylogeny indicates that our assemblies support a deep coalescence of the MHC. (I-L) Diverse views into an extraction of the MHC from the implicit pangenome graph. *(J) A 1-D odgi viz rendering shows expected pattern of diversity in MHC, with structures mostly conserved with the exception of large deletions seen in gibbon. (K) A rendering of the entire region with odgi draw, demonstrating its collinearity across the clade. (I) A subgraph around the C4A/C4B locus suggests that there are distinct versions of the inserted endogenous retroviral sequence (nested loop). (L) MHC class II cell surface receptor genes show rapid structural evolution and diversity across the clade, as shown by the tangled structure around HLA-DRB1, HLA-DRB5, and HLA-DRB6.

##### wgatools pseudo-MAF (pMAF)

We generated all-to-all alignments using wfmash (https://github.com/waveygang/wfmash), producing one PAF with each assembly as a target/reference. Next, we used wgatools (github.com/wjwei-handsome/wgatools) to filter the PAF (16 PAFs) for min alignment len >10 Mbp and then convert PAF to MAF (384 MAFs).

##### SNP vs. gap divergence

We computed SNV and gap divergence from each pairwise alignment, estimated with wfmash v.0.13, for 1 Mbp segments running across the target haplotype. We report the mean and median SNV and gap divergence. We also binned segments based on SNV and gap divergence to create density plots (**Fig. SeqDiv S1** & **S2**). SNV divergence is defined as the fraction of positions in the target haplotype where the two haplotypes are in different nucleotide states. Gap divergence is defined as the fraction of positions in the target haplotype that are not aligned to the other haplotype, which could be due to biological processes (e.g., gene loss/gain and insertions/deletions), missing data, or technical problems (e.g., alignment failure due to SVs, repetitive elements, etc.).

Autosome SNV divergence between human and NHPs was lowest for human-chimpanzee and human-bonobo (0.15-0.16%), then human-gorilla (0.19-0.20%) and lastly human-orangutan (0.36%). However, autosome gap divergence showed different trends, as it was highest for human-gorilla (17.9-27.3%). The within-species autosome gap divergence was also highest for gorilla (13.8%), which could be due to the assembly (larger number of contigs), SVs, or mobile elements. Within-species autosome SNV divergence was lowest for human (0.16%), followed by chimpanzee (0.27%), orangutan (0.35%), bonobo (0.36%), and lastly gorilla (0.58%). Trends are similar for X and Y chromosomes, with divergence values being lower for X and higher for Y.

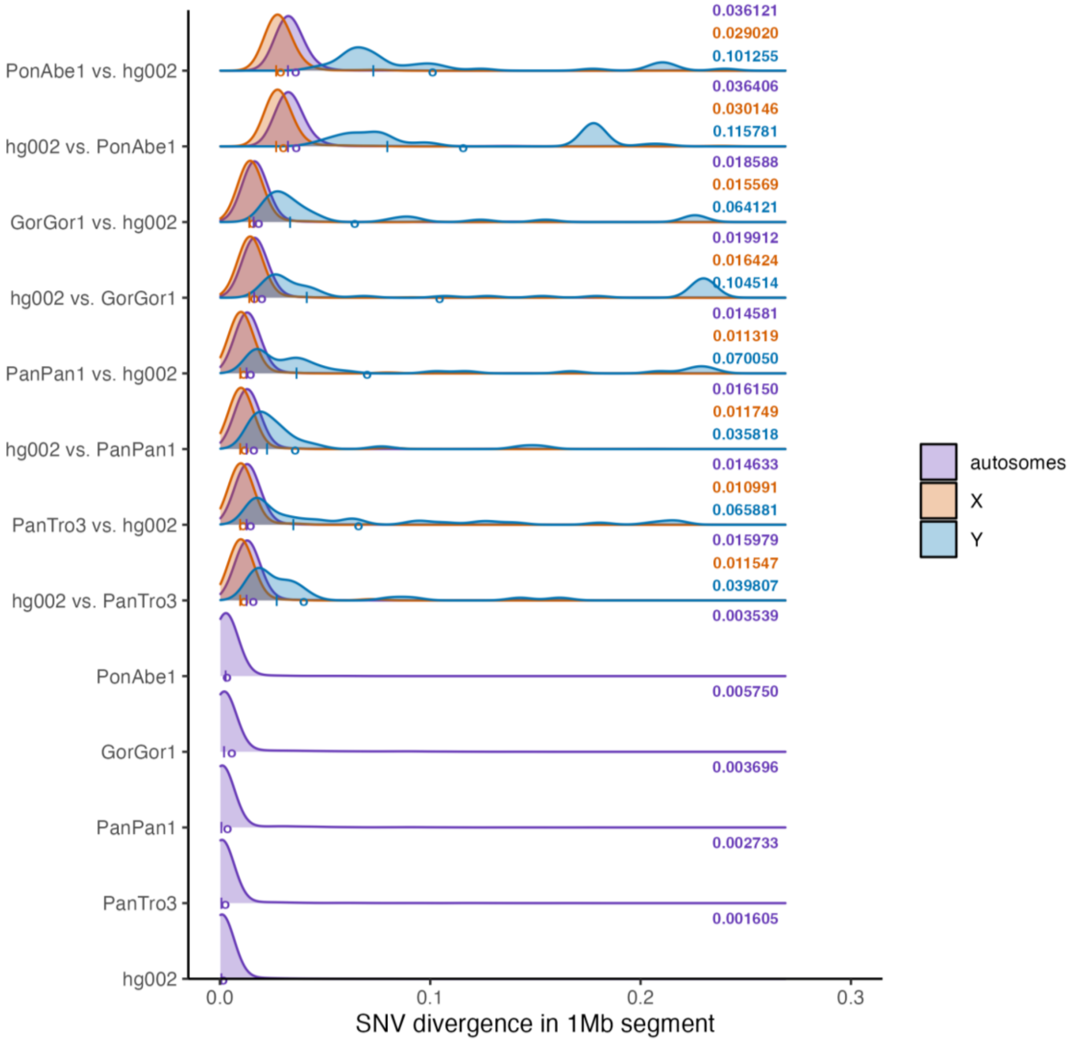


**Figure SeqDiv S1.** Plots show 1 Mbp segments binned by SNV divergence for each pairwise alignment (note that density, rather than counts, are shown). The maternal or primary haplotype is used for comparisons between human and NHP haplotypes. The second haplotype listed was aligned to the first/target haplotype (note that "A vs. B" and "B vs. A" are different pairwise alignments because the former includes the entire A haplotype with no gaps and the latter includes the entire B haplotype with no gaps). Density plots are broken down according to whether segments come from an autosome, the X chromosome, or the Y chromosome. Mean SNV divergence is reported for these three cases (numeric values and circles); medians are also shown (| characters).

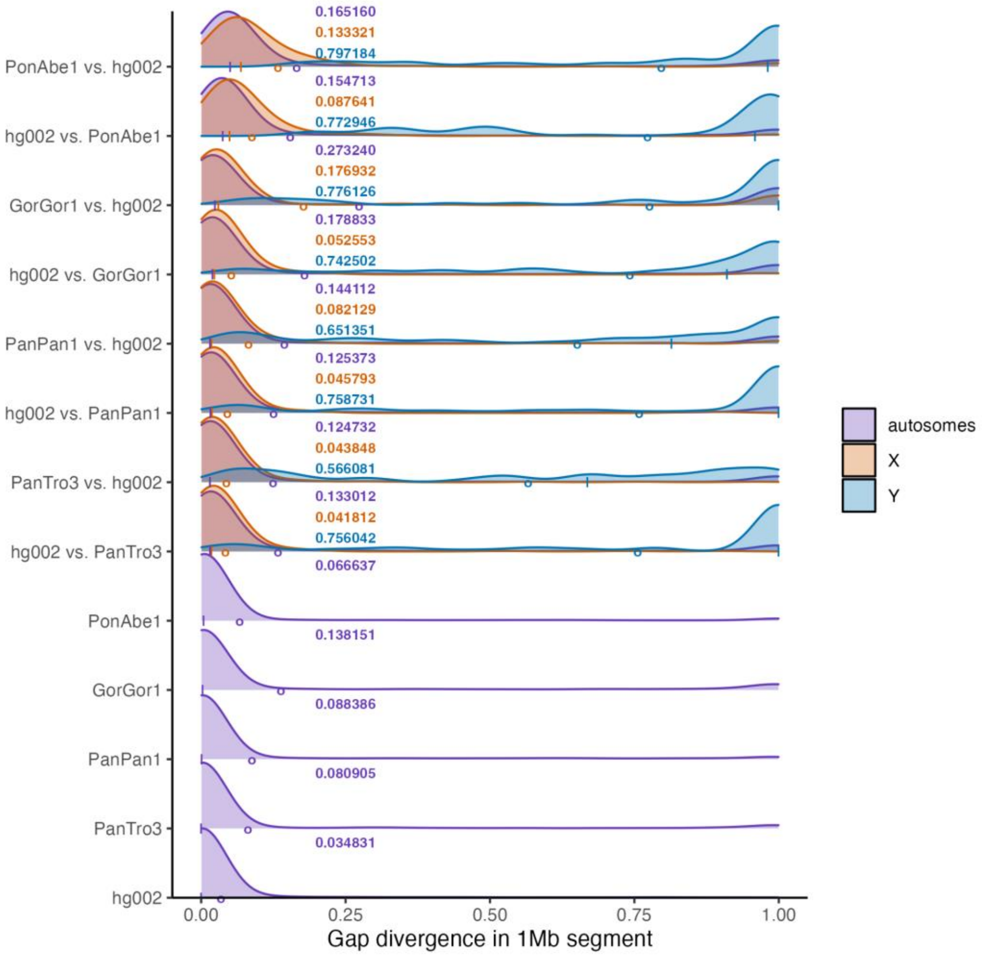


**Figure SeqDiv S2.** Plots show 1 Mbp segments binned by gap divergence for each pairwise alignment.

#### Conservation analyses

We generated per-base conservation scores and conserved elements tracks using PhastCons[15] with two approaches, following the PhastCons HOWTO (http://compgen.cshl.edu/phast/phastCons-HOWTO.html) as described in Secomandi et al.[16]. In the first approach, we used default parameters (unsupervised EM learning), splitting alignments into 1 Mbp chunks and combining predictions. For the second approach, we applied a fourfold degenerate site model, using entire alignments for all chromosomes except for chromosome 2, which was split into 1 Mbp chunks. In this approach, we fine-tuned the 4d model parameters --target-coverage and --expected-length to maximize enrichment (Jaccard index) of CHM13 exon sequences. The initial model was estimated with maximum likelihood and a tree from TimeTree (https://timetree.org).

We analyzed the conservation scores generated from the fourfold degenerate-sites-based model in several different ways. First, we assessed the distribution of conservation scores binned into 1000 base-pair windows (**Fig. SeqDiv S3**). Conservation scores were strongly skewed towards 1 (mean 0.964), as expected for such closely related species. We next assessed conservation over coding and noncoding genes using the Comparative Annotation Toolkit (CAT) liftoff track from UCSC from T2T-CHM13 of genecode genes (**Fig. SeqDiv S4-S5**). Gene conservation was distributed as expected with retained introns, snoRNAs, nmd-associated transcripts, and protein-coding genes exhibiting the highest conservation and various classes of pseudogene exhibiting the lowest. We plotted these conservation scores genome-wide revealing several hotspots of reduced conservation, including the rapidly evolving MHC locus. Hotspots additionally tended to correspond to the subtelomeric ends of chromosomes and the Y chromosome additionally showed reduced constraint. Finally, we compared conservation scores to pLI, which is a metric of constraint from human population genetics (**Fig. SeqDiv S1E-F**). pLI was strongly correlated with conservation (P=8e-164) indicating that genes under constraint in the human population are also conserved in apes. Nevertheless, genes that are off the trend line indicate those that exhibit increased constraint between species and decreased constraint within the human population (bottom right corner) and those that exhibit increased constraint within the human population and reduced constraint across species (top left corner).

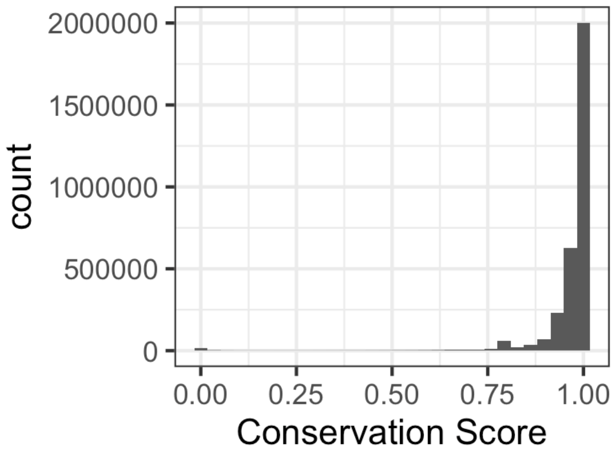


**Figure SeqDiv S3. Conservation scores binned in 1000 base-pair windows.**

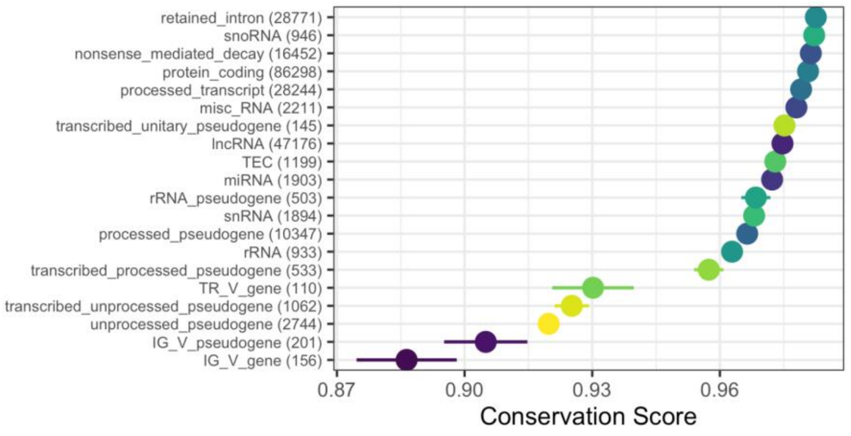


**Figure SeqDiv S4. Average conservation scores as a function of genomic feature.**

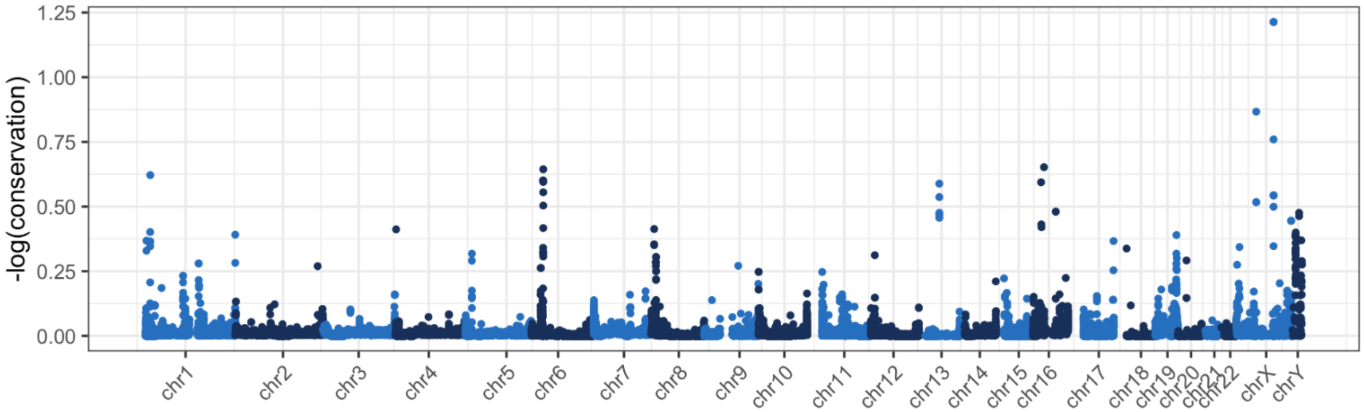


**Figure SeqDiv S5. Conservation scores per gene plotted as a Manhattan plot. Higher values indicate lower conservation.**

#### Minigraph-Cactus pangenome graphs

We constructed two pangenome graphs using Minigraph-Cactus[17]. The first of 10 African ape haplotypes, includes diploid assemblies for chimpanzee, bonobo, gorilla and T2T-HG002, as well as hg38 and hs1 on which it is referenced. The second graph was constructed from diploid Bornean and Sumatran orangutan genomes (4 haplotypes total) and is referenced on the primary Bornean orangutan assembly. In both cases, the graphs were constructed all at once, rather than being split by reference chromosome, as was previously done for the HPRC[18], in order to better account inter-chromosomal alignments.

The graphs were constructed on a Slurm cluster using Cactus v2.7.1 and the following commands.

```
TOIL_SLURM_ARGS="--partition=long --time=8000" cactus-
pangenome ./js-pg ./10-t2t-apes-mc-2023v2.seqfile --outDir
10-t2t-apes-mc-2023v2 --outName 10-t2t-apes-mc-2023v2 --
reference hs1 hg38 --noSplit --gbz clip full --gfa clip
full --xg clip full --odgi --vcf --giraffe clip --haplo
clip --vcfReference hs1 hg38 --logFile 10-t2t-apes-mc-
2023v2.log --batchSystem slurm --coordinationDir /data/tmp
--caching false --batchLogsDir ./batch-logs --consMemory
1500Gi --indexMemory 1500Gi --mgMemory 500Gi --mgCores 72 -
-mapCores 8 --consCores 128 --indexCores 72 --giraffe clip


TOIL_SLURM_ARGS="--partition=long --time=8000" cactus-
pangenome ./js-pg ./4-t2t-orangs-mc-2023v2.seqfile --outDir
4-t2t-orangs-mc-2023v2 --outName 4-t2t-orangs-mc-2023v2 --
reference mPonAbe1_pri mPonAbe1_alt --noSplit --gbz clip
full --gfa clip full --xg clip full --odgi --vcf --giraffe
clip --haplo clip --vcfReference mPonAbe1_pri mPonAbe1_alt
--logFile 4-t2t-orangs-mc-2023v2.log --batchSystem slurm --
coordinationDir /data/tmp --batchLogsDir ./batch-logs --
consMemory 1500Gi --indexMemory 1500Gi --mgMemory 500Gi --
mgCores 72 --mapCores 8 --consCores 128 --indexCores 72 --
giraffe clip
```

Note that the input, output including UCSC Genome Browser track hubs, and all steps to reproduce Minigraph-Cactus pangenomes can be found at https://cglgenomics.ucsc.edu/february-2024-t2t-apes/.

The statistics of these graphs, alongside the HPRC v1.1 Minigraph-Cactus graph are as follows:

**Table PangenomeGraphS1:** Pangenome graph statistics. “Length” refers to the total length of all nodes in the graph and “Avg. Clipped” is the amount of sequence (bp), on average, clipped out of the graph for each non-reference genome.

| Pangenome | Haplotypes | Reference | Ref. Length | Nodes | Edges | Length | Avg. Clipped |
|---|---|---|---|---|---|---|---|
| HPRC v1.1 | 90 | T2T-CHM13 | 3117292070 | 93165628 | 128451813 | 3338032439 | 166761108 |
| African Apes | 10 | T2T-CHM13 | 3117292070 | 264554139 | 361346441 | 3383639539 | 380296289 |
| Orangutan | 4 | B. Orang (primary) | 3259853530 | 69685551 | 94270798 | 3350249901 | 193691200 |

Minigraph-Cactus produces VCF output alongside the graph representations. We used `bcftools norm -f`, `vcf-bub -l 0 -a 100000` then `vcfwave -I 1000` from ghcr.io/pangenome/pggb:202402032147026ffe7f for normalization, then Truvari v4.2.2 to merge similar SVs: `truvari collapse -r 500 -p 0.95 -P 0.95 -s 50 -S 100000`. Note that multiallelic sites were split with `bcftools norm -m -any | bcftools sort` before `truvari` and remerged with `bcftools norm -m +any | bcftools sort` after `truvari`. Note that these VCFs exclude sites with variants >100 kbp. The number of variants in each VCF, again including HPRC v1.1, for comparison is as follows:

**Table PangenomeGraphS2:** The number of variant sites in pangenome VCFs. SVs include sites with an allele of length >50 bp and ≤100 kbp.

| Pangenome | SNPs | MNPs | Indels | SVs |
|---|---|---|---|---|
| HPRC v1.1 | 22237802 | 799337 | 5694302 | 231751 |
| African Apes | 66573699 | 3541028 | 8589725 | 310406 |
| Orangutan | 18381520 | 1374958 | 3161356 | 107778 |

Multiple Alignment Format (MAF) files were exported from the pangenome graphs using `cactus-hal2maf`.

```
for i in hs1 hg38 ; do TOIL_SLURM_ARGS="--partition=long --
time=8000" cactus-hal2maf ./js ./10-t2t-apes-mc-
2023v2.full.hal ./10-t2t-apes-mc-2023v2.${i}.maf.gz --
filterGapCausingDupes --refGenome $i --chunkSize 500000 --
batchCores 64 --noAncestors --batchCount 16 --batchSystem
slurm --caching false --logFile ./10-t2t-apes-mc-
2023v2.${i}.maf.gz.log --batchLogsDir batch-logs-16apes --
coordinationDir /data/tmp ;done
```

```
for i in GCA_028885655.2 GCA_028885685.2 ; do
TOIL_SLURM_ARGS="--partition=long --time=8000" cactus-
hal2maf ./js ./4-t2t-orangs-mc-2023v2.full.hal ./4-t2t-
orangs-mc-2023v2.${i}.maf.gz --filterGapCausingDupes --
refGenome $i --chunkSize 500000 --batchCores 64 --
noAncestors --batchCount 16 --batchSystem slurm --caching
false --logFile ./4-t2t-orangs-mc-2023v2.${i}.maf.gz.log --
batchLogsDir batch-logs-16apes --coordinationDir
/data/tmp ; done
```

We computed coverage statistics for these alignments using `taffy coverage` (as included in Cactus v2.7.1) and aggregated them into the following table:

**Table PangenomeGraphS3:** Alignment coverage of T2T-CHM13 (hs1) in the African ape pangenome.

| Region | Length | Query | Aligned (pct) | Identical (pct) | Aligned 1:1 (pct) | Identical 1:1 (pct) |
|---|---|---|---|---|---|---|
| Autosomes | 2900572475 | GRCh38 | 92.34 | 92.24 | 92.15 | 92.04 |
| Autosomes | 5801128381 | HG002 | 93.59 | 93.48 | 93.17 | 93.07 |
| Autosomes | 5801111812 | Chimp | 89.35 | 88.17 | 88.96 | 87.78 |
| Autosomes | 5801111812 | Bonobo | 89.3 | 88.12 | 88.88 | 87.71 |
| Autosomes | 5801111812 | Gorilla | 88.29 | 86.81 | 87.62 | 86.14 |
| X | 154259566 | GRCh38 | 96.34 | 96.28 | 96.33 | 96.27 |
| X | 154259566 | HG002 Mat | 97.72 | 97.67 | 97.71 | 97.66 |
| X | 154259566 | Chimp Pri | 94.85 | 93.86 | 94.79 | 93.81 |
| X | 154259566 | Bonobo Mat | 94.94 | 93.95 | 94.83 | 93.84 |
| X | 154259566 | Gorilla Mat | 94.31 | 92.91 | 94.16 | 92.76 |
| Y | 62460029 | GRCh38 | 34.7 | 34.67 | 34.64 | 34.6 |
| Y | 62460029 | HG002 Pat | 99.94 | 99.94 | 99.94 | 99.94 |
| Y | 62460029 | Chimp Pri | 19.86 | 15.41 | 16.83 | 12.59 |
| Y | 62460029 | Bonobo Pat | 18.17 | 14.26 | 17.61 | 13.8 |
| Y | 62460029 | Gorilla Pat | 12.81 | 7.09 | 11.04 | 6.26 |

**Table PangenomeGraphS4:** Alignment coverage of Bornean orangutan primary in the orangutan pangenome.

| Region | Length | Query | Aligned (pct) | Identical (pct) | Aligned 1:1 (pct) | Identical 1:1 (pct) |
|---|---|---|---|---|---|---|
| Autosomes | 3028670501 | B.Orang | 91.2 | 90.92 | 90.48 | 90.2 |
| Autosomes | 6057341002 | S.Orang | 90.07 | 89.68 | 89.1 | 88.72 |
| X | 162586321 | S.Orang Pri | 94.1 | 93.8 | 94.1 | 93.8 |
| Y | 67827326 | B.Orang Alt | 0.91 | 0.89 | 0.91 | 0.89 |
| Y | 67827326 | S.Orang Alt | 1.14 | 1.13 | 0.16 | 0.15 |
| Y | 67827326 | S.Orang Pri | 41.72 | 39.14 | 41.55 | 38.98 |

**Progressive Cactus genome alignment**

We used Progressive Cactus[19] to construct two genome alignments: An 8-way primary alignment of the six T2T apes plus hg38 and hg31, as well as a 16-way diploid alignment of the same samples, but also including HG002. We used MashTree v1.4.6[19] with default arguments to compute guide trees for the alignment, restricting to autosomes in the case of the diploid genome assemblies. The resulting guide tree for the 8-way primary alignment was:

> (((GCA_028885655.2:0.0017500000000000016,GCA_028885625.2:0.001729999999999999):0.014950000000000001,(GCA_029281585.2:0.00877,((GCA_029289425.2:0.0019999999999999983,GCA_028858775.2:0.0022900000000000004):0.0043300000000000005,(hs1:5.000000000000004E-4,hg38:5.000000000000004E-4):0.005989999999999999):0.0014300000000000007):0.007359999999999985):0.011345,GCA_028878055.2:0.011345000000000003);

and the Cactus (v2.7.1) commands used to construct the alignments were:

```
TOIL_SLURM_ARGS="--partition=long --time=8000" cactus ./js-8apes ./8-t2t-apes-2023v2.seqfile ./8-t2t-apes-2023v2.hal --batchSystem slurm --caching false --consCores 64 --configFile ./config-slurm.xml --logFile 8-t2t-apes-2023v2.hal.log --batchLogsDir batch-logs-8apes --coordinationDir /data/tmp

TOIL_SLURM_ARGS="--partition=long --time=8000" cactus ./js-16apes ./16-t2t-apes-2023v2.seqfile ./16-t2t-apes-
```

```
2023v2.hal --batchSystem slurm --caching false --consCores
64 --configFile ./config-slurm.xml --maxOutgroups 3 --
chromInfo 16-t2t-apes-2023v2.chroms --logFile 16-t2t-apes-
2023v2.hal.log --batchLogsDir batch-logs-16apes --
coordinationDir /data/tmp
```

Note that the input, output including UCSC Genome Browser track hubs, and all steps to reproduce Progressive Cactus alignments can be found at https://cglgenomics.ucsc.edu/february-2024-t2t-apes/.

We extracted an MAF version of each alignment referenced on each species and computed the coverage, all as described above for the pangenome graphs. The coverage on T2T-CHM13/hs1 is described in these tables.

**Table PangenomeGraphS5:** Alignment coverage of T2T-CHM13 (hs1) in the 8-way primary Progressive Cactus alignment.

| Region | Length | Query | Aligned (pct) | Identical (pct) | Aligned 1:1 (pct) | Identical 1:1 (pct) |
|---|---|---|---|---|---|---|
| Autosomes | 2900555906 | GRCh38 | 93.81 | 93.67 | 87.08 | 86.96 |
| Autosomes | 2900555906 | Chimp | 91.47 | 90.22 | 86.1 | 84.95 |
| Autosomes | 2900555906 | Bonobo | 91.48 | 90.22 | 85.98 | 84.83 |
| Autosomes | 2900555906 | Gorilla | 90.89 | 89.33 | 85.44 | 83.99 |
| Autosomes | 2900555906 | B.Orang | 88.49 | 85.51 | 83.41 | 80.62 |
| Autosomes | 2900555906 | S.Orang | 88.47 | 85.49 | 83.4 | 80.61 |
| Autosomes | 2900555906 | Siamang | 84.73 | 81.36 | 79.91 | 76.78 |
| X | 154259566 | GRCh38 | 97.66 | 97.56 | 88.9 | 88.82 |
| X | 154259566 | Chimp Pri | 95.41 | 94.37 | 86.48 | 85.57 |
| X | 154259566 | Bonobo Pri | 95.44 | 94.4 | 86.49 | 85.59 |
| X | 154259566 | Gorilla Pri | 94.53 | 93.08 | 85.62 | 84.34 |
| X | 154259566 | B.Orang Pri | 90.69 | 88.05 | 81.78 | 79.44 |

| | | | | | | |
|---|---|---|---|---|---|---|
| X | 154259566 | S.Orang Pri | 90.59 | 87.95 | 81.64 | 79.31 |
| X | 154259566 | Siamang Pri | 84.09 | 81.22 | 75.71 | 73.21 |
| Y | 62460029 | GRCh38 | 40.22 | 40.12 | 12.21 | 12.14 |
| Y | 62460029 | Chimp Pri | 33.93 | 32.97 | 8.01 | 7.75 |
| Y | 62460029 | Bonobo Pri | 34.12 | 33.16 | 8.3 | 8.02 |
| Y | 62460029 | Gorilla Pri | 31.33 | 30.28 | 6.64 | 6.46 |
| Y | 62460029 | B.Orang Pri | 26.96 | 25.35 | 6.1 | 5.71 |
| Y | 62460029 | S.Orang Pri | 26.99 | 25.37 | 6.09 | 5.7 |
| Y | 62460029 | Siamang Pri | 21.18 | 19.65 | 3.91 | 3.66 |

**Table PangenomeGraphS6:** Alignment coverage of T2T-CHM13 (hs1) in the 16-way diploid Progressive Cactus alignment.

| Region | Length | Query | Aligned (pct) | Identical (pct) | Aligned 1:1 (pct) | Identical 1:1 (pct) |
|---|---|---|---|---|---|---|
| Autosomes | 2900572475 | GRCh38 | 93.57 | 93.42 | 87.99 | 87.87 |
| Autosomes | 5801144950 | HG002 | 95.07 | 94.83 | 88.99 | 88.8 |
| Autosomes | 5801128381 | Chimp | 91.07 | 89.84 | 85.02 | 83.89 |
| Autosomes | 5801111812 | Bonobo | 91.08 | 89.84 | 84.92 | 83.79 |
| Autosomes | 5801128381 | Gorilla | 90.62 | 89.08 | 84.5 | 83.08 |
| Autosomes | 5801144950 | B.Orang | 88.39 | 85.42 | 82.55 | 79.79 |
| Autosomes | 5801144950 | S.Orang | 88.4 | 85.42 | 82.58 | 79.82 |
| Autosomes | 5801111812 | Siamang | 84.87 | 81.5 | 79.05 | 75.96 |
| X | 1542595 | GRCh38 | 97.26 | 97.11 | 86.19 | 86.13 |

| | | | | | | |
|---|---|---|---|---|---|---|
| | 66 | | | | | |
| X | 154259566 | HG002 Mat | 96.99 | 96.91 | 85.39 | 85.34 |
| X | 154259566 | HG002 Pat | 7.48 | 6.96 | 0.12 | 0.09 |
| X | 154259566 | Chimp Alt | 1.67 | 1.42 | 0.24 | 0.19 |
| X | 154259566 | Chimp Pri | 91.66 | 90.5 | 80.71 | 79.88 |
| X | 154259566 | Bonobo Mat | 91.03 | 90 | 80.31 | 79.5 |
| X | 154259566 | Bonobo Pat | 4.93 | 4.39 | 0.38 | 0.32 |
| X | 154259566 | Gorilla Mat | 90.21 | 88.78 | 79.55 | 78.39 |
| X | 154259566 | Gorilla Pat | 4.36 | 3.9 | 0.32 | 0.28 |
| X | 154259566 | B.Orang Alt | 1.47 | 1.24 | 0.21 | 0.17 |
| X | 154259566 | B.Orang Pri | 86.95 | 84.36 | 75.91 | 73.76 |
| X | 154259566 | S.Orang Alt | 1.49 | 1.26 | 0.21 | 0.17 |
| X | 154259566 | S.Orang Pri | 86.14 | 83.54 | 75.2 | 73.07 |
| X | 154259566 | Siamang Alt | 1.81 | 1.6 | 0.68 | 0.64 |
| X | 154259566 | Siamang Pri | 79.55 | 76.76 | 69.73 | 67.46 |
| Y | 62460029 | GRCh38 | 38.96 | 38.67 | 8.9 | 8.87 |
| Y | 62460029 | HG002 Mat | 20.66 | 19.39 | 0.3 | 0.26 |
| Y | 62460029 | HG002 Pat | 58.41 | 58.06 | 27.15 | 27.02 |
| Y | 62460029 | Chimp Alt | 5.25 | 4.82 | 0.19 | 0.16 |
| Y | 62460029 | Chimp Pri | 27.95 | 27.1 | 4.4 | 4.32 |
| Y | 62460029 | Bonobo Mat | 18.92 | 17.66 | 0.38 | 0.33 |

| | | | | | | |
|---|---|---|---|---|---|---|
| Y | 62460029 | Bonobo Pat | 21.98 | 21.24 | 4.53 | 4.43 |
| Y | 62460029 | Gorilla Mat | 18.03 | 16.83 | 0.29 | 0.25 |
| Y | 62460029 | Gorilla Pat | 20.46 | 19.75 | 4.04 | 3.93 |
| Y | 62460029 | B.Orang Alt | 4.22 | 3.85 | 0.08 | 0.07 |
| Y | 62460029 | B.Orang Pri | 24.41 | 22.92 | 3.51 | 3.28 |
| Y | 62460029 | S.Orang Alt | 4.23 | 3.86 | 0.1 | 0.08 |
| Y | 62460029 | S.Orang Pri | 24.49 | 23.01 | 3.55 | 3.32 |
| Y | 62460029 | Siamang Alt | 3.64 | 3.31 | 0.05 | 0.04 |
| Y | 62460029 | Siamang Pri | 19.52 | 18.1 | 2.2 | 2.05 |

**Annotation of the human-primate ancestral allele**

We used the parsimony-like method used by the 1000 Genomes Project and Ensembl[20,21] with the following tree for this annotation, ((Gorilla,((Bonobo,Chimp)b,Human)a)c), where a, b, and c refer to the inferred ancestral sequences. Instead of using the EPO pipeline used by Ensembl, we used an 8-way alignment produced by Progressive Cactus available here https://cgl.gi.ucsc.edu/data/cactus/t2t-apes/8-t2t-apes-2023v2/. The Ensembl human-primate ancestor based in GRCh38 was downloaded from https://ftp.ensembl.org/pub/release-112/fasta/ancestral_alleles/.

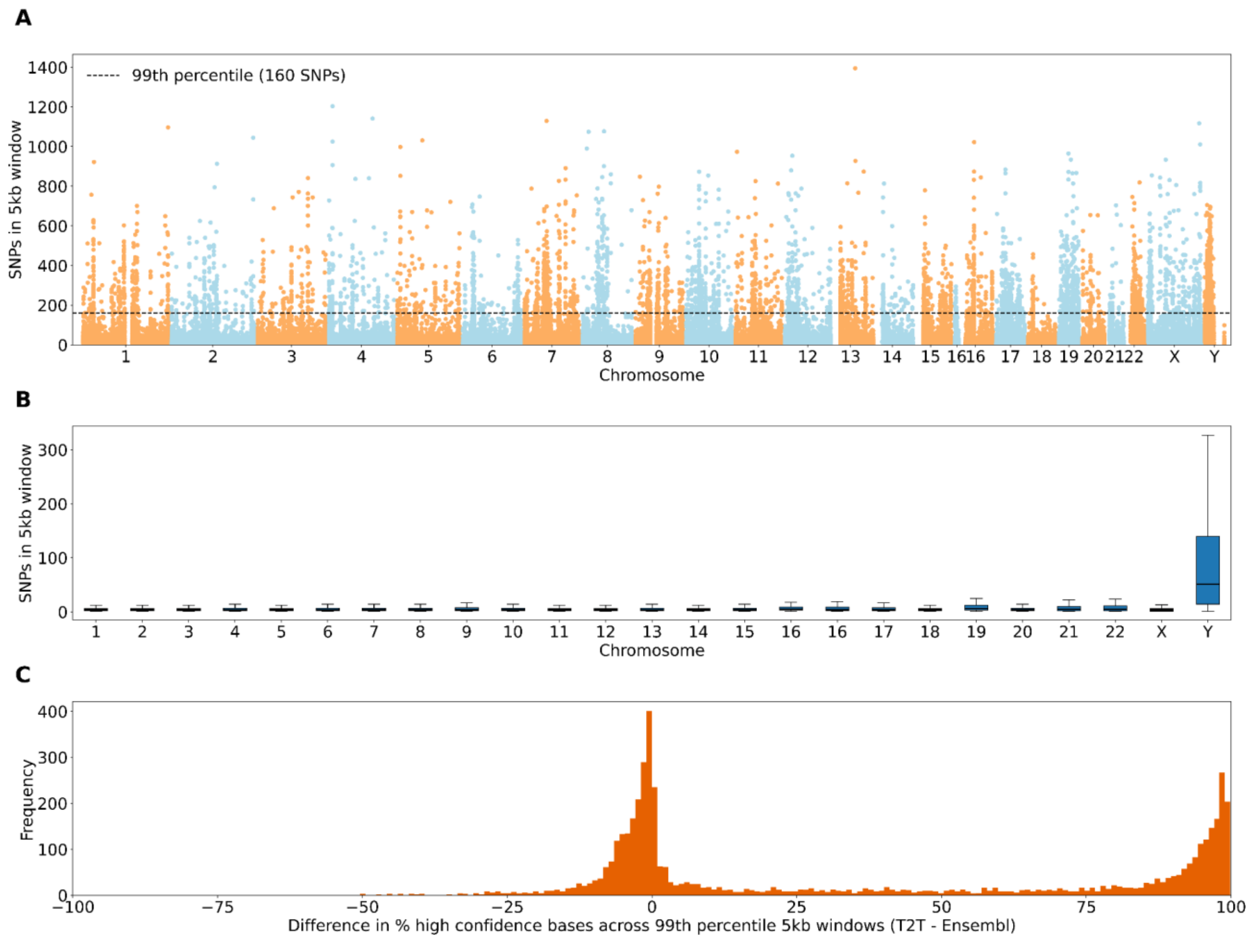


**Figure AncestralAlleles S1. Annotation of ancestral allele.** (A) SNPs per 5 kbp window between the T2T annotation and the Ensembl annotation of the human-primate ancestor, both based on GRCh38. (B) Boxplots showing the distribution of SNPs in 5 kbp windows for autosomes. (C) Difference in the percentage of high-confidence bases in 5 kbp windows (n=4840, shown in panel A exceeding the 99$^{th}$ percentile in SNPs, between the T2T annotation and Ensembl annotation. The ancestral base is recorded in high confidence if all three ancestral sequences agree on the base, otherwise it is low confidence indicating partial agreement.

### V. Sumatran vs. Bornean orangutan divergence

Contributing authors:

Robert S. Harris, Saswat Mohanty, and Kateryna D. Makova (Penn State University)

##### Methods

Pairwise alignments between the two orangutan genomes were extracted from the 8-way cactus[19] alignment (8-t2t-apes-2023v2.hal):

> hal file → hal2maf → maf_filter_to_species_set → mafDuplicateFilter → pairwise alignments

Sequence identity statistics were collected from these alignments:

> pairwise alignments → maf_to_plain_pairwise_identity → stats.

In particular, we computed sequence identity over alignable bases, as well as blast identity over alignment length (**Table OrangDivS1-3**) over each chromosome as well as weighted by alignment length average across the autosomes:

> identity m/(m+mm) = 99.63%

and

> blast identity m/(m+mm+i+d) = 99.38%

where m = match, mm = mismatch, i = insertion, d = deletion.

Separately, we aligned the two orangutan genomes using LASTZ[13] and computed the same metrics (m, mm, i, d, identity, and blast identity) for these alignments. The following parameters were used:

> --notransition
>
> --scores=scoring/human_chimp.v2_scoring
>
> --allocate:traceback=1.5G

We computed sequence identity over alignable bases, as well as blast identity over alignment length over each chromosome (**Table OrangDivS2**) as well as weighted by alignment length average across the autosomes:

> identity m/(m+mm) =97.17%

and

> blast identity m/(m+mm+i+d) =96.34%

#### Summary of results

Here, for the first time, we sequenced and assembled the genome of Bornean orangutan, significantly (i.e., to the T2T level) improved the genome of Sumatran orangutan, and performed their detailed comparison. The two species diverged very recently, only approximately 0.5-2 mya[22-24] and are the most closely related species in our dataset. The sequence identity of alignable bases between the two orangutan genomes was 99.63% from 8-way alignments (considering autosomes only, **Table OrangDivS1**; sequence identities for the sex chromosomes are reported in Makova et al.[1] and 97.17% from 2-way alignments (again, considering autosomes only; both autosomal and sex chromosome values are reported in **Table OrangDivS2**).

### VI. Incomplete lineage sorting (ILS) and speciation times

Contributing authors:

Francesco Montinaro, Iker Rivas-González

##### Methods

Divergence time represents the average coalescent time between two sequences and can vary significantly across the genome. In contrast, speciation time refers to the minimum time at which two sequences can coalesce, reflecting when species become reproductively isolated.

We estimated ILS among different primates using TRAILS, which integrates a Hidden Markov Model of the ILS signal with a time discretization approach for the unbiased inference of demographic parameters, allowing the posterior decoding of both topology and coalescent times.

We performed the ILS estimation on the following four-species (ABCD) phylogenies (**Table ILS.S1**):

- *Homo sapiens; Pan troglodytes; Gorilla gorilla, Pongo abelii* (HCGO)
- *Pan troglodytes; Pan paniscus; Homo sapiens; Pongo abelii* (CBHO)
- *Homo sapiens, Pan troglodytes, Pongo abelii, Symphalangus syndactylus* (HCOS)
- *Homo sapiens, Gorilla gorilla, Pongo abelii, Symphalangus syndactylus* (HGOS)
- *Pongo Abelii, Pongo Pygmaeus, Homo sapiens, Symphalangus syndactylus* (OOHS)

We also harnessed msmc2[32] to infer the population size of single species; we started from the diploid multi-alignment, removed duplicates and created the multihet-step msmc2 input file using the MsmcOutput flag. Next, we estimated the between haplotype coalescence rates across time using default parameters and converted the inferred metrics to effective population size as in Schiffels[32] and Wang et al.[33] We used the same mutation rate of TRAILS analysis and the following generation times:

1. Chimp: 24 years
2. Bonobo: 24 years
3. Human: 25 years
4. Gorilla: 19 years
5. Borneo and Sumatra Orangutan: 25 years
6. Gibbon: 15 years

##### Summary of results

We started from the haploid or diploid multiz-alignment of the eight primate species and extracted the four relevant species for each phylogeny. We then merged syntenic regions separated by less than 200 bp and retained only blocks longer than 2 kbp. Every analysis was repeated three times, alternatively including the maternal/primary or paternal/alternative haplotype of the analyzed individuals. For Homo sapiens, we also included the t2t hs1 T2T-CHM13 haplotype as detailed in **Table ILS.S2**. For all the analyses we used a mutation rate of $1.25 \times 10^{-8}$ and the average generation time across the species for specific node:

1. Chimp-Bonobo: 24 years.
2. Homo-Chimp-Bonobo: 24.5 years.
3. Homo-Chimp-Gorilla: 21.75 years.
4. Homo-Chimp-Gorilla-Orangutan: 23.375 years.
5. Homo-Chimp-Gorilla-Orangutan-Gibbon: 19.2 years.
6. Orangutan Borneo-Orangutan Sumatra: 25 years.

**Table ILS.S2**: Details of the GenBank accession numbers for the harnessed haplotypes for each analyzed phylogeny in TRAILS.

| Name phylogeny | A | B | C | D |
|---|---|---|---|---|
| HCGO | hs1 | GCA_028858775.2 | GCA_029281585.2 | GCA_028885655.2 |
| HCGOm | hg002_mat | GCA_028858775.2 | GCA_028885495.2 | GCA_028885655.2 |
| HCGOp | hg002_pat | GCA_028858805.2 | GCA_028885475.2 | GCA_028885685.2 |
| CBHO | GCA_028858775.2 | GCA_029289425.2 | hs1 | GCA_028885655.2 |
| CBHOm | GCA_028858775.2 | GCA_028858845.2 | hg002_mat | GCA_028885655.2 |
| CBHOp | GCA_028858805.2 | GCA_028858825.2 | hg002_pat | GCA_028885685.2 |
| HCOS | hs1 | GCA_028858775.2 | GCA_028885655.2 | GCA_028878055.2 |
| HCOSm | hg002_mat | GCA_028858775.2 | GCA_028885655.2 | GCA_028878055.2 |
| HCOSp | hg002_pat | GCA_028858805.2 | GCA_028885685.2 | GCA_028878085.2 |
| HGOS | hs1 | GCA_029281585.2 | hs1 | GCA_028878055.2 |
| HGOSm | hg002_mat | GCA_028885495.2 | hg002_mat | GCA_028878055.2 |
| HGOSp | hg002_pat | GCA_028885475.2 | hg002_pat | GCA_028878085.2 |
| OOHS | GCA_028885655.2 | GCA_028885625.2 | hs1 | GCA_028878055.2 |
| OOHSm | GCA_028885655.2 | GCA_028885625.2 | hg002_mat | GCA_028878055.2 |
| OOHSp | GCA_028885685.2 | GCA_028885525.2 | hg002_pat | GCA_028878085.2 |

For each of the 15 trees, we performed two optimization steps for the parameter and posterior probability estimations using the starting values as in Rivas-Gonzales et al.{Rivas-González, 2023 #61} The first and second optimization steps were carried out by setting three discrete time intervals for both AB and ABC species.

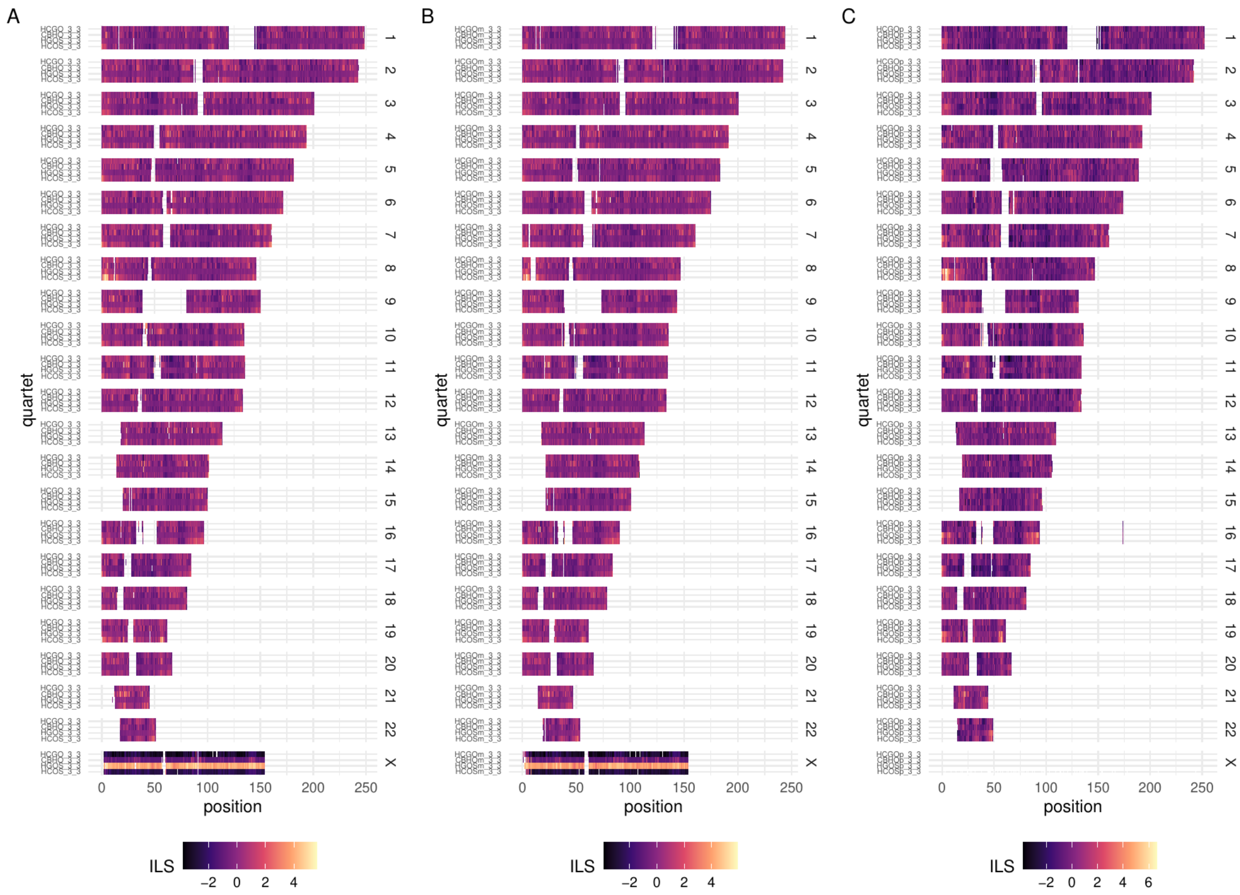


**Fig ILS.S1. Inference of ILS and demographic parameters.** Inference of ILS proportion for 500 kbp windows among four phylogenies using chm/primary (A), maternal/primary (B), and paternal/alternative (C) haplotypes.

The parameter inference results across the five species' trees are summarized in **Table ILS.S1**. The estimated time and population size parameters across the three replicates of each phylogeny are consistent. This is also confirmed when the correlation is assessed across genes for phylogenies using primary, maternal, or paternal haplotypes (**Fig. ILS.S2**). In fact, for HCGO, considering the ILS proportion in windows that overlap with genes, a high correlation across replicates has been observed ($R^2$ HCGO vs. HCGOm = 0.88; $R^2$ HCGO vs. HCGOp = 0.89).

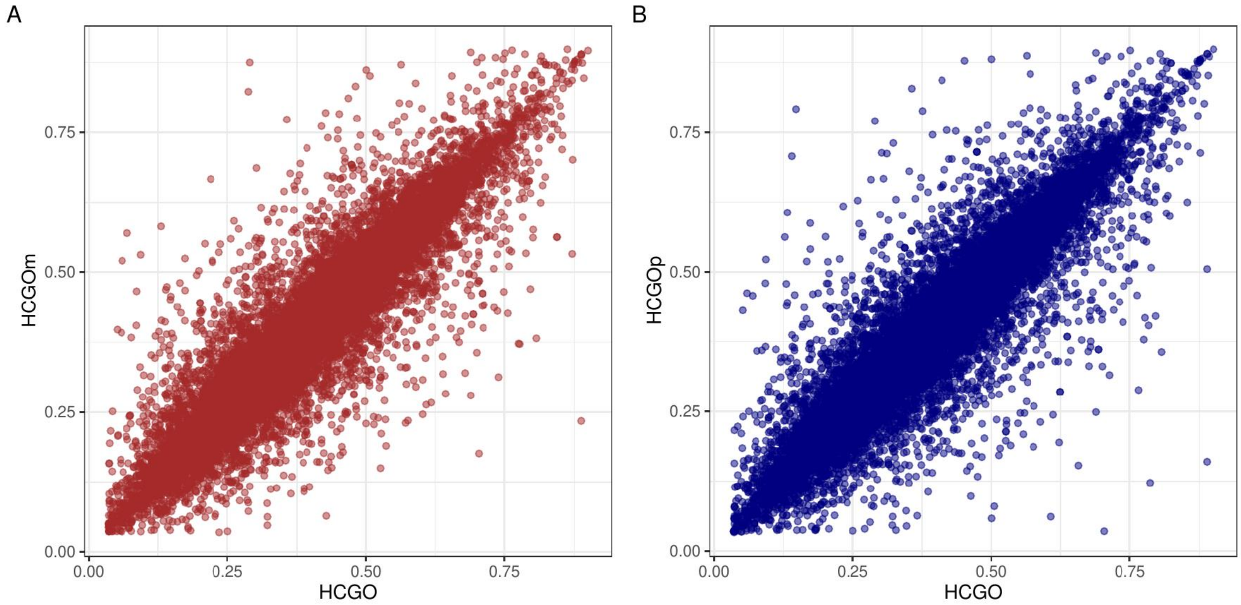


**Figure ILS.S2. Comparison of genic ILS proportion for the HCGO phylogeny across different replicates.** A) HCGO vs. HCGOm. B) HCGO vs. HCGOp.

For the HCGO tree, we inferred the presence of 39.5% of autosomal genome and 24% of chromosome X in ILS, with an increase of approximately 7.5% compared to previous estimates[34]. Human and chimpanzee speciation time from the ancestral species has been estimated between 5.6 and 6.3 mya, respectively, in line with previous research. The speciation time from gorilla to the HGO ancestral species was estimated to be 10.6-10.9 mya, and the orangutan speciation to 18.2-19.6 mya. The population size of the ancestral population of human and chimpanzee (N=198,000) is larger than that estimated for human, chimpanzee, and gorilla (Ne=132,000), suggesting an increase of the population size between 6 and 12 mya. Moreover, we confirm the substantially reduced diversity for chromosome X for the HC ancestral population (Ne=76,700) but not for the population ancestral to HCG (N=115,600). This pattern can be explained by multiple population dynamics, such as strong selective sweeps on the X chromosome or, alternatively, reduced size for the female founder population.

For the CBHO phylogeny, we inferred approximately 5.8% of ILS, with an X chromosome estimate of 3.4%. The speciation time of the split is 1.58 mya, in line with previous research.[35] We estimated a speciation time between human and CB to approximately 6.8 mya and the O to CBHO 17.7 mya, confirming the robustness of the inference irrespectively of the phylogeny analyzed. The ancestral population size for the CB population (N=46,800) is about a third of that estimated for CBH (N= 115,600). We inferred 0.7% of ILS for the HCOS topology across the autosomal genome and 0.5% on the X chromosome. For HGOS, we inferred approximately 1.8% and 1.4% ILS across autosomal and X chromosomes, respectively.

Compared to previously reconstructed maps, the T2T assemblies allow to compute ILS in previously inaccessible genomic regions such as that encompassing the HLA genes (**Fig. ILS.S3**), which shows relatively high levels of ILS for the HCGO phylogeny. In fact, many HLA genes show an ILS proportion higher than 0.6, with high concordance across replicates (**Table ILS.S3**).

**Table ILS.S3. ILS of HLA genes.**

| Gene | ILS proportion | | |
|---|---|---|---|
| | HCGO | HCGOm | HCGOp |
| HLA-DMB | 0.8063075084 | 0.7542187326 | 0.7887304234 |
| HLA-DQA1 | 0.7867638855 | 0.889432011 | 0.1220549421 |
| HLA-DMA | 0.765377122 | 0.7646002827 | 0.7353741486 |
| HLA-E | 0.7520348678 | 0.7475375258 | 0.7395590469 |
| HLA-DPB2 | 0.6873574726 | 0.4777737681 | 0.6505982899 |
| HLA-DOB | 0.6154815119 | 0.5663079763 | 0.6512983486 |
| HLA-F-AS1 | 0.5094971594 | 0.32544819 | 0.2353133172 |
| HLA-DOA | 0.4851129494 | 0.5051618998 | 0.4735974102 |
| HLA-DPA1 | 0.4705243042 | 0.457028394 | 0.4924196097 |
| HLA-DPB1 | 0.3886423843 | 0.3876098463 | 0.4407424665 |
| HLA-DRA | 0.3776579652 | 0.454397155 | 0.4601328216 |
| HLA-L | 0.3064009895 | 0.2705134076 | 0.3276100972 |

To further explore the locus, we analyzed the chr6:25Mbp-40Mbp region, both at window and base-pair posterior decoding level. When 100 kbp windows are analyzed, the region harboring HLA genes shows an increased ILS proportion between 30 and 33 Mbp, with a very similar pattern in all the considered phylogenies (**Fig. ILS.S3**).

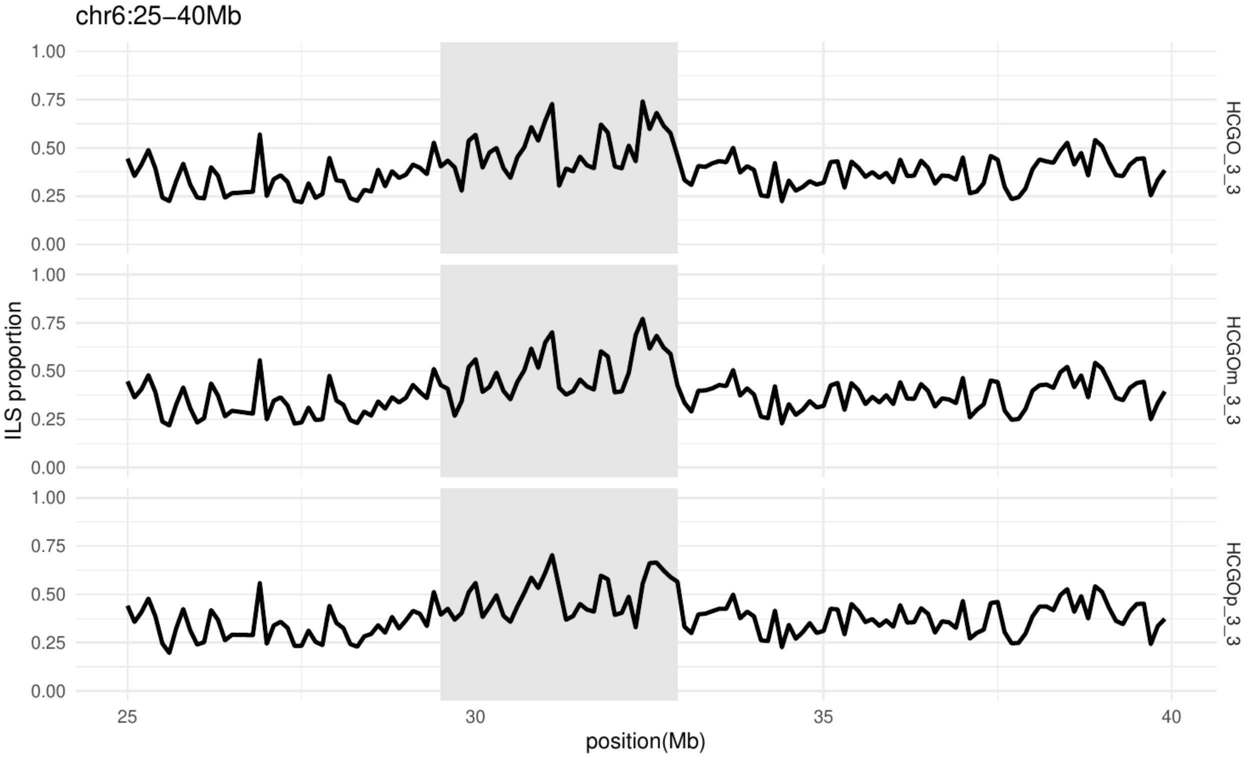


**Figure ILS.S3. ILS proportion in the chr6:25-40Mbp for HCGO phylogeny considering 100 kbp windows.**

The posterior decoding gives per-base-pair posterior probabilities of observing the hidden states and, thus, we can build an ILS map at the highest resolution. **Fig. ILS.S4** reveals that there are stretches of the genome that favor one of the two alternative topologies that do not follow the canonical species tree. For example, at around chr6:32.85Mbp, there is a region favoring the chimpanzee-gorilla topology, while chr6:29.30Mbp favors a human-gorilla topology.

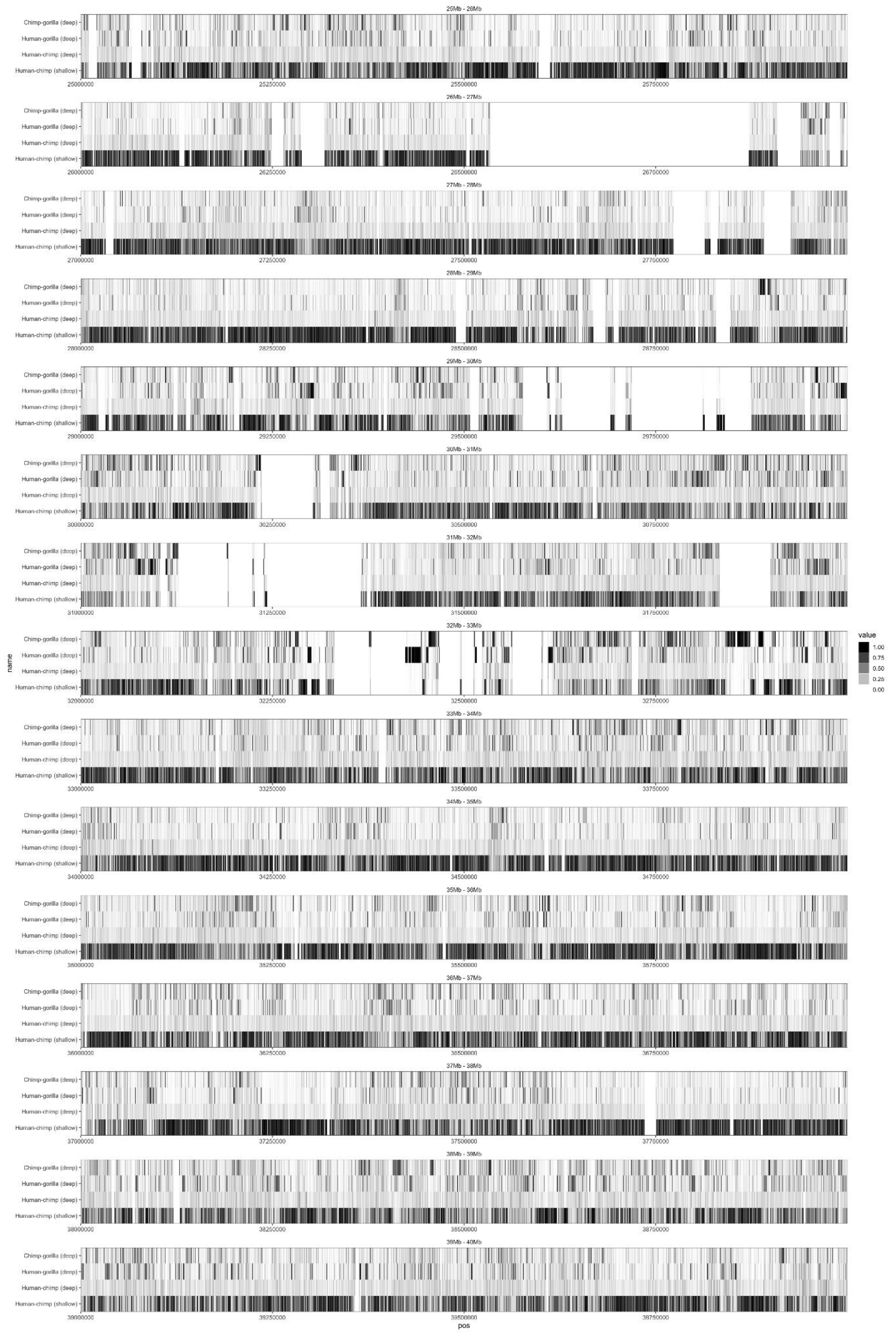


**Figure ILS.S4. ILS proportion in the chr6:25-40Mbp for HCGO phylogeny at base-pair level.**

The effective population time through time is shown in **Fig. ILS.S5**

We confirm a general decline of Pan genus population size starting approximately 5 mya, and a subsequent increase between 1 and 2 mya. For gorilla, we inferred a relatively low constant sample size, possibly reflecting an inbreeding history[36]. For orangutans, we observe a similar trend until 1 mya, confirming the estimates obtained with TRAILS, followed by a steep decline in the Bornean orangutan. Notably, the effective population size for the Sumatran orangutan is the historically highest before 100,000-200,000 years ago, as reflected by the broad geographic distribution of Pleistocene fossil remains in mainland Asia.

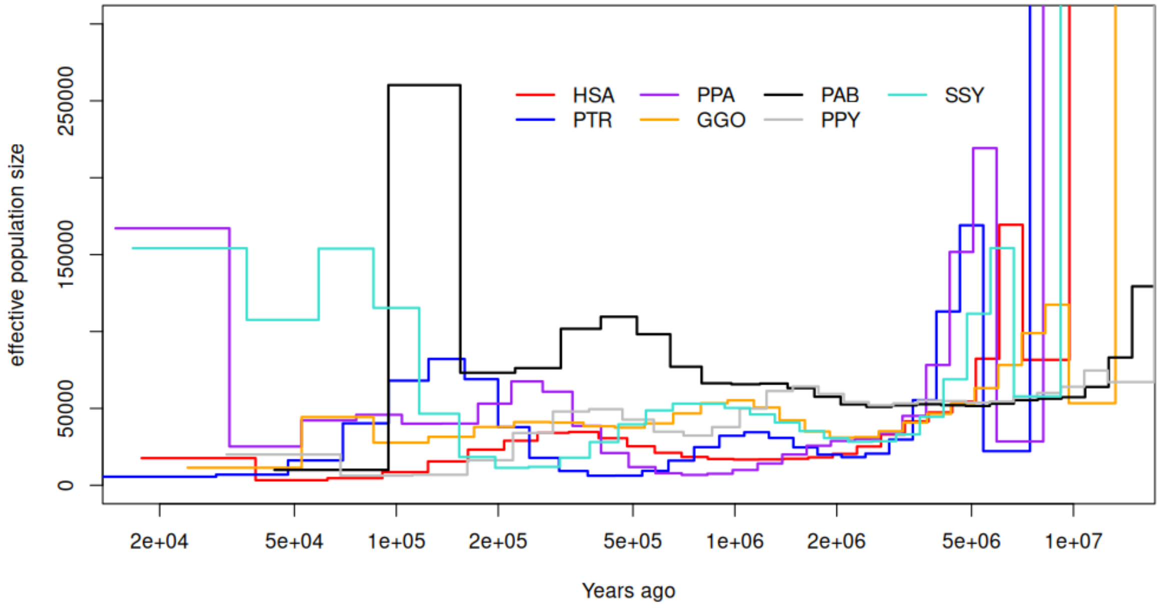


**Figure ILS.S5. Effective population size trajectories through time as inferred by msmc2.**

### VII. Gene annotation

Contributing authors:

Prajna Hebbar, Francisca R. Ringeling, Françoise Thibaud-Nissen, Diana Haddad, Patrick Masterson, Karol Pal, Juan F. I. Martinez, Mark Diekhans, Stefan Canzar, Kateryna D. Makova, Benedict Paten

#### Methods

##### RefSeq annotation

The *de novo* gene annotations of the six primate assemblies were performed by the NCBI Eukaryotic Genome Annotation Pipeline (version 10.2) between Feb 27 and Mar 13, 2024, as previously described[1]. Protein-coding and long noncoding genes were predicted based on the alignments to the genome of same-species PacBio Iso-Seq and RNA-seq reads queried from the Sequence Read Archive (SRA), RefSeq human transcripts and proteins, and GenBank Primate proteins. The number of Iso-Seq reads used for the annotation ranged from 16.57 million (*Symphalangus syndactylus*) to 21.99 million (*Pan paniscus*) while the number of RNA-seq reads ranged from 1.34 billion (*Pongo pygmaeus*) to 6.96 billion (*Pan troglodytes*).

##### CAT gene annotation

Genome annotation was performed using CAT. First, whole-genome alignments between the ape (gorilla, chimpanzee, bonobo, Sumatran orangutan, Bornean orangutan, and siamang) and human GRCh38, and T2T-CHM13v2.0) genomes were generated using Cactus (8-way primary alignment described above). CAT then used the whole-genome alignments to project the UCSC GENCODEv35 CAT/Liftoff v2 annotation set from T2T-CHM13v2 to the primates. CAT was run with transMap, AUGUSTUS, Liftoff[25], AUGUSTUS-PB, and miniprot[26] modes. transMap lifts over gene annotations from the reference onto all the genomes in the cactus alignment. Liftoff lifts over gene annotations from a reference onto a minimap2 alignment between the reference and target genome. The miniprot mode uses protein homology information to improve gene annotations. CAT was given Iso-Seq FLNC data to provide extrinsic hints to the Augustus PB (PacBio) module of CAT, which performs *ab initio* prediction of coding isoforms. CAT then combined these *ab initio* prediction sets with the various human gene projection sets to produce the final gene sets and UCSC assembly hubs used in this project.

RNA-seq reads were aligned using minimap2[27] using the following command:

```
minimap2 -a -x sr --sam-hit-only --secondary=no --eqx -t 4 mmdb/0.mmi
rnaseq_data/0_0.fasta
```

Iso-Seq reads were aligned using minimap2 using the following command:

```
minimap2 -ax splice:hq -uf --sam-hit-only --secondary=no --eqx -t 4 mmdb/0.mmi
isoseq_data/0_0.fasta
```

CAT[26] (https://github.com/ComparativeGenomicsToolkit/Comparative-Annotation-Toolkit) was run using the following command:

```
luigi --module cat RunCat --hal=8-t2t-apes-2023v2.hal --ref-genome=hs1 -- workers=10 -
-config=t2t.apes.config --work-dir t2t_apes_2023v2/cat_work --out-dir
t2t_apes_2023v2/cat_output --local-scheduler --binary-mode local --augustus --augustus-
pb --liftoff –miniprot --maxCores 45 --assembly-hub >& log_t2t_apes_2023v2_CAT.txt
```

with the T2T-CHM13v2 annotation from UCSC GENCODEv35 CAT/Liftoff v2 as input along with a config file with locations to reference gff3, RNA-seq, and Iso-Seq BAM files.

**Novel gene annotation and curation of the integrated protein-coding gene annotation set**

The annotations generated by CAT were first compared with the gene annotations generated by the NCBI RefSeq pipeline. For protein-coding genes, the two sets displayed a high concordance, with an average Jaccard similarity score of over 0.9. Upon inclusion of pseudogenes and other noncoding genes, the similarity scores drop to 0.78 due to differences in biotype assignment methods between the pipelines. To resolve these differences, we provide a unique and useful gene annotation resource in the form of a consensus gene annotation between the two pipelines. To generate this, a reliable orthologous gene set was first generated. The orthologous genes were identified using the transMap method of CAT, which uses the Cactus alignment to map the orthologs. The cases where genes were mapped to a completely different neighborhood than the one in human were flagged and resolved using mappings from the liftoff mode. Then loci of all the protein-coding genes from this set were compared to the orthologous loci assigned by the NCBI RefSeq pipeline. For genes that were mapped to two completely different loci, the transcripts from both were mapped against human. Depending on the percentage identity of the generated protein to human (>50%), the transcripts were either discarded or assigned as ortholog/novel paralog. The novel gene loci that were annotated by either pipeline were collected and filtered on three levels: length of the protein generated >200 AA, identity with human protein >50%, and Iso-Seq transcript support. These were then merged into the consensus gene set.

**Lineage-specific gene family analysis**

Using the CAT gene annotations, we find 99.0%-99.63% of human genes annotated on the primates with at least 90% completeness. There are about 185 gene families that have fully intact protein-coding copies present only in humans. We also identified a fraction (2.0%-3.4%) of putative protein-coding genes present in the T2T genomes of the NHPs that were absent in the human annotation set used. In addition to this, 2.1%-5.0% of transcripts annotated exhibited Iso-Seq-supported splice junctions that are unique to the NHPs. The novel protein-coding genes and exons, which have been identified as gained or lost between humans and NHPs, are documented in **Table GeneS1**. There have been a number of gene family expansions in the NHPs, with between 1394-2056 novel gene copies found across the 184-258 families. Around 50% of these overlap with lineage-specific segmental duplication (SD) regions. A few of these occur in regions of interest.

##### Segmental duplication (*MAPKBP1, JMJD7-PLA2G4B, SPTBN5*) in chr1 in gorilla

*MAPKBP1* codes for a protein that plays a regulatory role in the JNK and NOD2 pathways. This is a gene present on chromosome 15 in humans. The ortholog of this gene is present in chromosome 16 (homologous to 15) in gorillas. However, there is a significant expansion of this gene in chromosome 1, in tandem with the *JMJD7-PLA2G4B* and *SPTBN5* genes (**Fig. 6**). The three genes form a unit that is repeated eight times in a region spanning 13.5 Mbp near the breakpoint of double tandem inversion specifically found in the gorilla genome. It is important to note that each copy of *MAPKBP1* and *JMJD7-PLA2G4B* are supported by Iso-Seq transcripts and have valid open-reading frames (ORFs). An alignment of the proteins is provided in **Fig. GeneS1**. There are also six copies of *PLA2G4B-JMJD7* spanning at least 80% of the homologous human sequence in chimpanzee and three copies in bonobo, all supported by multiple Iso-Seq transcripts (**Table Gene S5**).

##### Expansion (*HERC2, GOLGA6/8, MCTP2*) in chr16 in Pongo

*HERC2* is a duplicon that has been associated with the common breakpoint regions of Prader-Willi and Angelman syndrome deletions along with *GOLGA6* and *GOLGA8*. In genomic regions spanning over 20 Mbp in both orangutans, multiple copies of these genes, along with copies of other medically important genes, such as *MCTP2*, have been identified. For the Bornean orangutan, 25 copies of *GOLGA6*, 33 copies of *GOLGA8*, 23 copies of *HERC2*, 5 putative *HERC2-GOLGA* fusion gene copies, and 5 copies of *MCTP2* were located. In Sumatran orangutan, there are 34 copies of *GOLGA6*, 42 of *GOLGA8*, 21 copies of *HERC2*, 11 putative *HERC2-GOLGA* fusion gene copies, and 4 *MCTP2* copies. Every *HERC2* copy annotated in the **Table GeneS5** has Iso-Seq support. However, not all *GOLGA* copies have Iso-Seq support.

##### *LRPAP1* expansion in gorilla

*LRPAP1* encodes for a protein that interacts with the LDL-receptor protein and is present in chromosome 4 in humans. The ortholog of this gene is present in chromosome 3 (homologous to 4) in gorillas. This gene undergoes copious expansion across chromosomes, all of these falling in lineage-specific SD regions. This gene also is not duplicated alone. It forms a unit with the genes *DOK7* and *HGFAC*, which are expanded by valid ORFs. However, only *LRPAP1* gene copies have Iso-Seq transcript support. Apart from the orthologous copy in chromosome 3, four additional copies in chr12, two in chr14, and one each in chr16, chr22, and chrY were identified. It is also to be noted that two copies, one in chr12 and one in chr14, do not co-occur with *DOK7* and *HGFAC*. These gene loci are mentioned in **Table GeneS5**.

##### *PSMA5* expansion in Pan and Gorilla

*PSMA5* gene duplication with SORT1-like pseudogene in chimpanzee, bonobo, and gorilla *PSMA5* is duplicated twice in chimpanzee and bonobo and thrice in gorilla in conjunction with the *SORT1*-like pseudogene right downstream of it in all cases. This is seen in both haplotypes. These duplicated copies occur upstream of the ancestral copy in all of the cases and all copies have Iso-Seq support and valid ORFs. The duplicated *PSMA5* copies are all truncated in the same manner, and the accompanying *SORT1* copies are all pseudogenized. All of these regions overlap with lineage-specific SD regions in the species as well.

In addition, genes that specifically duplicated in orangutans with Iso-Seq support as well as valid ORF were found as follows:

*HTATIP2*: 2 copies in each haplotype
*PRMT3*: 2 copies in each haplotype
*HEATR6*: 2 copies in each haplotype
*FAHD2A*: 2 copies in each haplotype
*COG7*: 3 copies in each haplotype
*RFX8*: 5 copies in each haplotype
*RBIS*: 5 copies in each haplotype
*PIGW*: 3 copies in each haplotype
*HSFX2*: 2 copies in each haplotype
*NUTM2B*: 2 copies in each haplotype

##### Analysis of human Pongo and Pan-specific genes

Novel gene copies that were present only in the specific genus were collected and analyzed for gene ontology enrichment; 185 human-, 212 Pan-, and 234 Pongo-specific expanded genes families, which qualify by having novel copies with Iso-Seq support and valid ORFs, were used. Gene ontology analysis suggests enrichment of metabolic process in pan genus, as well as signaling and neurogenesis/nervous system development functions in pongo (**Table GeneS6**).

##### Improved Iso-Seq transcript mapping

All long-read samples were aligned using minimap2 v2.24-r1122[27] with parameters recommended for PacBio Iso-Seq cDNA (-ax splice:hq -uf) allowing up to 15 alignments per read (-N 15). Mismatch and indel rates were computed based on primary alignments only using Perbase v0.9.0 (Stadick 2023 *https://github.com/sstadick/perbase*). Mismatch rates are defined as the sum of all bases covered by at least one read with a base different from the reference, divided by all covered bases. Indel rates correspond to the sum of all bases overlapping an insertion or deletion, divided by all covered bases. Reads for which the total number of soft-clipped bases exceeded 200 bp were counted using a custom awk script.

Transcripts were assembled from aligned reads using StringTie2 v2.2.1[28] with default parameters. We did not provide a reference annotation file to be able to attribute differences in transcript assembly to the quality of the reference sequence alone. To compare transcripts assembled from reads mapped to the T2T reference and from those mapped to previous assemblies, we lifted gene models inferred by StringTie2 for T2T genomes over to previous assemblies using Liftoff v1.6.3[25]. Based on the new coordinates, we then compared transcripts assembled based on T2T and previous genome versions using Gffcompare v0.12.9[29], which defines two transcripts as equal (class code "=") if the coordinates of all donor and acceptor sites match, that is, if they contain the same set of introns.

StringTie2 and most other existing algorithms infer transcripts locus by locus. If not relying on a gene annotation, loci are identified by sets of reads, i.e., read bundles, that together span a (almost) contiguous genomic region. We ran StringTie2 with option -v and parsed the output with custom scripts to collect read bundles that allow for gaps, i.e., genomic regions that are not covered by any reads, of length at most 50 bp. The similarity of bundles formed by reads mapped to T2T and to previous assemblies was measured by the Jaccard index. The Jaccard index, or Jaccard similarity, between two bundles $RB_{T2T}$ and $RB_{prev}$ takes values between 0 and 1 and is defined as $| RB_{T2T} \cap RB_{prev} | / | RB_{T2T} \cup RB_{prev} |$.

To find all bundles in the previous assembly that have a Jaccard similarity to any of the T2T bundles above a threshold of 0.1, we performed an all-pairs similarity search based on an index proposed in Bayardo, Ma, & Srikant[30] and implemented in Python library SetSimilaritySearch (https://github.com/ekzhu/SetSimilaritySearch). For each species, we compiled all genome-wide bundle similarities in a bipartite graph using Python package NetworkX (https://github.com/networkx/networkx), where the two node sets correspond to bundles found in the two compared genome assemblies, and edge weights equal the Jaccard similarities.

Then for a given gene in T2T, either protein-coding (**Fig. Gene.S3f**) or member of a multicopy gene family (**Fig. Gene.S3g**), we find the read bundle spanning it by querying an interval tree (https://github.com/chaimleib/intervaltree) that stores all T2T bundles, with the start and end coordinates of the gene. Traversing nodes adjacent to that read bundle in the above similarity graph then yields all similar bundles in the previous assembly. All chromosome map figures depicting bundle similarities were plotted with the aid of the R package chromoMap v4.1.1[31].

To detect multicopy gene families, we used blastp to pairwise compare protein sequences from all protein-coding genes (longest isoform per gene as annotated by NCBI). Homology was defined based on a cutoff of 50% sequence identity and 35% protein[31]. Each set of pairwise homologous genes (under transitive closure) then forms a multicopy gene family. We then represented the gene family by all distinct bundles spanning any of its members. In a second step, we extended families by paralogous loci that might have been missing in the annotation or that were pseudogenized. To this end, we found T2T bundles with Jaccard similarity >0.8 to any of the original family members. To compare copy numbers supported by RNA reads to previous assemblies, we found all bundles in previous assemblies that were similar to any of the T2T family members using the graph-based approach described above.

**Table Gene.S4: Assembly accessions used in this study.**

| Species | T2T | | non-T2T | | |
|---|---|---|---|---|---|
| | Name | Accession | Name | Accession | Y amended |
| Gorilla | mGorGor1/Jim_GGO | GCF_029281585.2 | Kamilah_GGO | GCF_008122165.1 | GCA_015021865.1 |
| Chimpanzee | mPanTro3/AG18354_PTR | GCF_028858775.2 | Clint_PTRv2 | GCF_002880755.1 | * |
| Bonobo | mPanPan1/PR00251_PPA | GCF_029289425.2 | panpan1.1 | GCF_000258655.2 | GCA_015021855.1 |
| Sumatran Orangutan | mPonAbe1/AG06213_PAB | GCF_028885655.2 | Susie_PABv2 | GCF_002880775.1 | GCA_015021835.1 |

*Chimpanzee's previous assembly already included a Y chromosome.

#### Summary of results

We used PacBio Iso-Seq long reads from testis RNA samples[1] of four great apes (chimpanzee, Sumatran orangutan, gorilla, and bonobo) to quantify the potential impact of T2T genome assemblies on read mapping and observed improvements in mappability, soft-clipping, and error rates. Iso-Seq reads were mapped with minimap2 to both T2T assemblies and previous assemblies (**Table Gene.S4**). We found 1,075 (0.07%), 1,353 (0.09%), 28,925 (0.7%) and 3,361 (0.1%) reads that were unmapped to previous assemblies but mapped to T2T assemblies in chimpanzee, bonobo, gorilla, and Sumatran orangutan, respectively (**Fig. Gene.S3a**). Conversely, only a few reads that could not be mapped to T2T mapped to the previous assembly (90, 24, 171 and 769 for chimpanzee, bonobo, gorilla and Sumatran orangutan, respectively). Soft-clipping allows for the mapping of Iso-Seq reads that do not align to the genome end to end. Large soft-clipping events may indicate a missing sequence in the reference genome or a rearrangement in the sequenced individual. We found many more reads with large-scale (>200 bp) soft-clipping when they were aligned to the previous assembly compared to T2T: 29,507 (2%) versus 23,358 (1.5%) in chimpanzee; 67,428 (4.3%) versus 24,192 (1.5%) in bonobo, 89,498 (2%) versus 33,032 (0.7%) in gorilla, and 30,043 (1%) versus 27,250 (0.9%) in Sumatran orangutan (**Fig. Gene.S3b**). We then looked at mismatch and indel rates and found that Iso-Seq reads mapped to T2T had consistently lower error rates compared to the previous assembly, even though the differences are modest (**Fig. Gene.S3b**).

We hypothesized that differences in read mapping might lead to discrepancies in transcript assembly and, indeed, we found thousands of transcripts uniquely assembled from reads mapped to T2T or to previous assemblies. While the majority of transcripts assembled with StringTie2 in all analyzed species were identical in T2T and previous assemblies, we found that all species had between 4,930 (in 1,929 loci) and 10,365 (in 3,816 loci) transcripts uniquely assembled from reads mapped to one or the other genome (**Fig. Gene.S3c**). Differences in mapping statistics (soft-clipping, mismatch and indel rates) were more pronounced in loci with differently assembled transcripts compared to genomic regions where all transcripts agreed (**Fig. Gene.S3d**), suggesting a larger fraction of correctly inferred transcripts among transcripts unique to T2T than among those unique to previous assemblies.

To identify genomic regions where T2T assemblies have improved mappability of Iso-Seq reads, we searched for regions where different sets of reads mapped in T2T assemblies compared to previous assemblies. Transcript assembly algorithms such as StringTie2 group reads that map to the same locus into so-called 'bundles'. We used the bundles generated by StringTie2 to identify these discrepant regions without relying on a specific algorithm to process them, and without relying on a high-quality gene annotation of previous assemblies. Measuring the similarity of bundles as the fraction of shared reads (Jaccard similarity, **Methods**), we found that most expressed protein-coding genes in T2T assemblies had a highly similar bundle mapped to the previous assembly (**Fig. Gene.S3f**). At the same time, we observed many regions in the T2T genome for which no similar bundles of reads mapped to the previous assembly. These regions are spread across all chromosomes, but are more prevalent around centromeric and telomeric regions, highlighting the improved resolution of repetitive regions in the T2T assembly. We also observed that low bundle similarity regions (Jaccard similarity < 0.2) overlap SDs more often than regions with a highly similar read bundle (Jaccard similarity > 0.8) (**Fig. Gene.S3e**).

We then leveraged the analysis of bundle similarities in T2T and previous assemblies to show how T2T assemblies improve the resolution of multi-copy gene families. Multi-copy gene families are prevalent in great apes and are highly relevant to the study of gene duplication and evolution. Starting from a curated list of gene families (**Methods**) and their corresponding read bundles in T2T assemblies, we looked for all similar read bundles mapping to previous assemblies and compared their numbers. For the vast majority of gene families, we found more gene copies in T2T assemblies compared to previous assemblies (**Fig. Gene.S3g**).

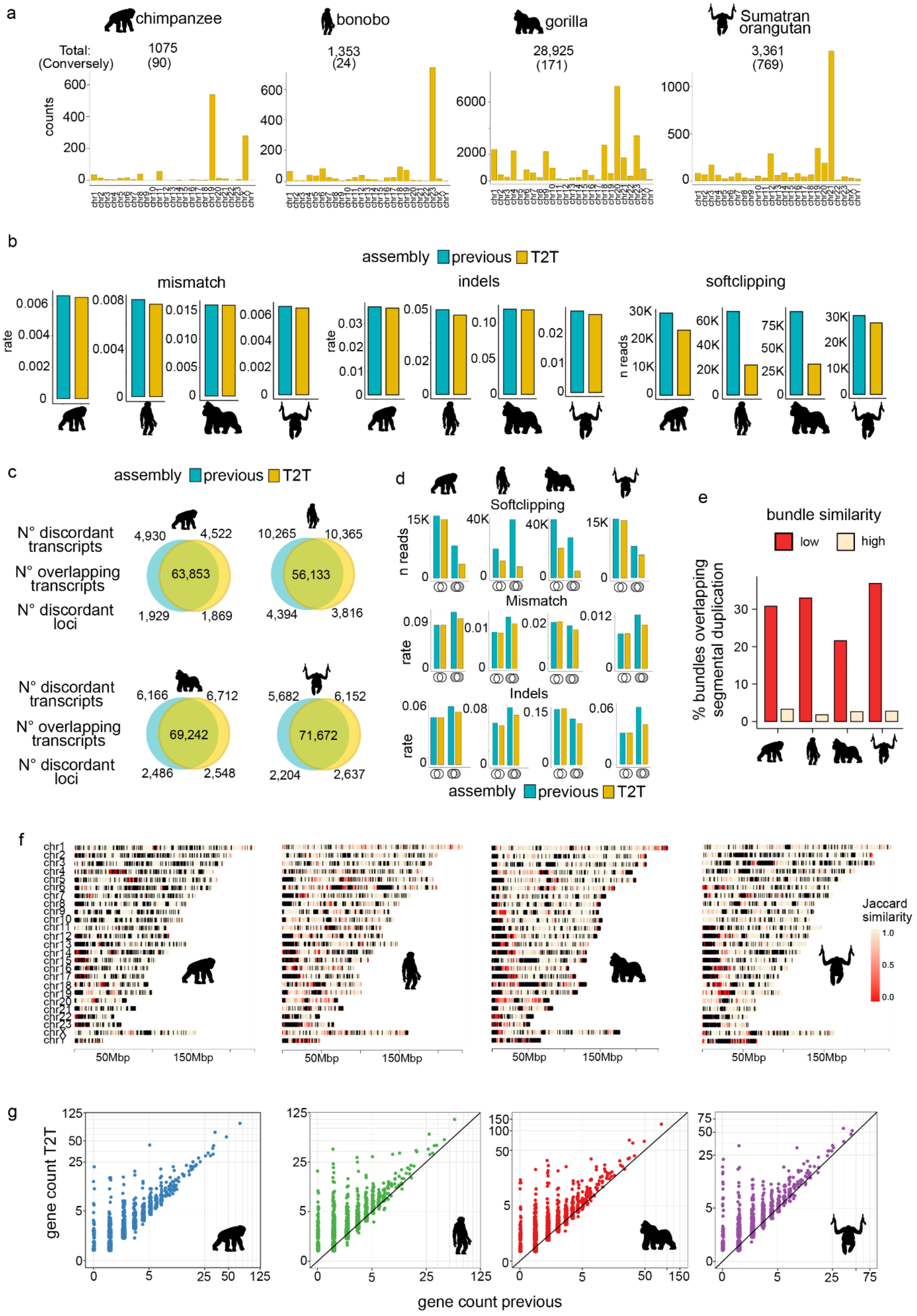
a
chimpanzee
bonobo
gorilla
Sumatran orangutan
Total:
(Conversely)
1075
(90)
1,353
(24)
28,925
(171)
3,361
(769)
counts
b
assembly previous T2T
mismatch
indels
softclipping
rate
n reads
c
assembly previous T2T
N° discordant transcripts
N° overlapping transcripts
N° discordant loci
4,930
4,522
63,853
1,929
1,869
10,265
10,365
56,133
4,394
3,816
6,166
6,712
69,242
2,486
2,548
5,682
6,152
71,672
2,204
2,637
d
Softclipping
Mismatch
Indels
n reads
rate
assembly previous T2T
e
bundle similarity
low
high
% bundles overlapping segmental duplication
f
Jaccard similarity
50Mbp
150Mbp
g
gene count T2T
gene count previous

**Figure Gene.S3. T2T assemblies improve transcript inference.** (a) Number of reads that remain unmapped when aligned to previous assemblies but map to T2T assemblies. Each plot shows the number of such reads per chromosome. Total number is shown above each plot and in parenthesis the converse number, i.e., number of reads unmapped in T2T mapped to previous assemblies. (b) Mapping statistics for Iso-Seq reads aligned to previous or T2T assemblies. Mismatch rate was calculated as the sum of all bases covered by at least one read with a base different from the reference, divided by all covered bases. Indel rate was calculated as the sum of all bases overlapping an insertion or deletion, divided by all covered bases. For soft-clipping plots, reads for which the total number of soft-clipped bases exceeded 200 bp were counted. (c) Comparison of transcripts assembled from Iso-Seq reads aligned to previous or T2T assemblies. (d) Mapping statistics for genomic regions where transcript predictions from reads aligned to previous assemblies are equal (matching intron chains) to those from reads mapped to T2T assemblies versus regions with non-equal transcripts. (e) Overlap of SDs and genomic regions with low bundle similarity (Jaccard similarity < 0.2) versus regions with high bundle similarity (Jaccard similarity > 0.8). Barplots show the percentage of regions that overlap in more than 80% of their length with SDs. (f) Ideograms showing T2T chromosomes colored by Jaccard similarities between bundles of reads mapped to T2T assemblies and read bundles on previous assemblies. (g) Scatterplots show number of gene copies per gene family on previous and T2T assemblies.

### VIII. Repeat annotation

Contributing authors:

Jessica M. Storer, Gabrielle A. Hartley, Mark Loftus, Parithi Balachandran, Panpan Zhang, Edmundo Torres-González, Hailey Loucks, Karen H. Miga, Kateryna D. Makova, Cedric Feschotte, Christine R. Beck, Miriam K. Konkel, Rachel J. O'Neill

#### Methods

##### Satellite and repeat annotations

We produced comprehensive repeat annotations across the ape lineage by integrating a combination of known repeats and models identified in human T2T-CHM13[37], T2T-Y[6], ape X/Y chromosomes[1], and *de novo* repeat curation (**Table Repeat.S1**). To identify canonical and novel repeats, we utilized the previously described pipeline[37], with modifications to include both the Dfam 3.691 and Repbase (v20181026)[38] libraries for each species during RepeatMasker[39] annotation. An initial RepeatMasker run identified canonical repeats, while a subsequent RepeatMasker run was completed to include repeat models first identified in the analysis of T2T-CHM13, T2T-Y, ape X/Y chromosomes (**Table Repeat.S2**) and newly defined satellites and 17 variants of pCht/StSat derived from Cechova, M. et al.[40] and the resulting annotations were merged. Because we previously discovered that prior taxonomic labeling for repeats was once considered lineage-specific (e.g., PtERV with a current taxonomic label in the repeat library of *Pan troglodytes*, therefore missed in searches of the gorilla and bonobo genomes) were excluded from the bonobo and gorilla genomes[1], an additional RepeatMasker run between the first search for canonical repeats and a subsequent search for novel repeats was performed. All of the results were combined as described previously[37].

To identify and curate previously undefined satellites, we utilized additional TRF[41] and ULTRA[42] screening of annotation gaps >10 kbp in length. Potential gaps were identified via BEDTools v2.29.0[12] by subtracting both the repeat and gene annotations for each ape reference sequence. To identify potential redundancy, satellite consensus sequences generated from gaps identified in each species were compared using crossmatch and were used as a RepeatMasker library to search for overlap in the other five analyzed primate species. Consensus sequences were considered redundant if there was a significant annotation overlap in the RepeatMasker output. Repeat consensus sequences were manually curated using RepeatMasker searches to ensure accuracy and identify additional variants.

##### Species-specific DNA

Species-specific insertions/expansions were characterized by identifying unaligned regions from Cactus[19] alignments of the seven primate X and Y chromosomes with halAlignExtract[43]. Unaligned regions were filtered by length and for tandem repeats using TRF and ULTRA.

RepeatMasker was used to identify the content of the lineage-specific insertions/expansions using the approach described above.

**Segmental duplication overlap**

Overlap between lineage-specific DNA and SDs was determined by utilizing BEDTools intersect using the default settings. The overlapping bp for the species-specific dataset and the SD dataset were calculated.

**Species-specific mobile element insertions (MEIs)**

Species-specific (SS) non-long terminal repeat retrotransposon (i.e., non-LTR: *Alu*, LINE/L1, and SVA) MEIs were characterized from the unaligned regions of Cactus alignments of the six great ape assemblies (PonPyg, PonAbe, GorGor, PanTro, PanPan, and hs1) with the siamang (SymSyn) gibbon assembly used as an outgroup. A local RepeatMasker (v4.1.6) installation with a standard library (Dfam v3.8)[44] was first used to identify the repetitive element content of the SS sequences. All SS sequences less than 15 kbp in length and annotated as having a non-zero percentage of repetitive sequence (any kind of repetitive sequence) by RepeatMasker were selected as potential candidates for downstream analyses. For each candidate SS locus, 500 bases of the flanking sequence up and downstream of the insertion coordinate was retrieved and fused. This fused sequence was provided to a local BLAT[45] installation (Standalone BLAT version: 36x2, parameters: -minScore=650, gap size +/- 20% SS sequence length) to query for homologous flanking segments containing similar sequence to the SS locus within all seven genomes. The sequence for each BLAT hit was pulled using SAMtools (version: 1.10) and then cut into *k*-mers (k=14), along with the original SS locus+flanking. Their *k*-mer-profile dissimilarity[46] was calculated to quantify the dissimilarity between the SS locus+flanking and each BLAT hit. Using a conservative approach, the BLAT hit was deemed a match if the *k*-mer-profile dissimilarity was ≤0.5. A candidate SS locus was filtered from the dataset if the SS sequence+flanking was identified in any other species. If a candidate SS locus was deemed unique to a species but duplicated, it was noted and all duplications were counted as only one potential insertion. The remaining 'high-confidence' SS loci were then screened for non-LTR mobile elements (SS sequence element percentage: LINEs/SVA: ≥20% to allow for transductions, *Alu* elements: ≥80%). These putative SS MEIs were subsequently put through a stringent filtering process screening for A-tails (minimum tail length ≥6 bp), percent divergence (LINEs/SVAs: <15% divergence, *Alu*: <6%), subfamily (LINEs: L1HS/PA4/PA3/PA2/PA1, *Alu* elements: AluY and derivatives), element length (LINEs/SVA: ≤10 kbp, Alu: ≤500 bp), and then randomly sampled and manually spot checked for quality control.

Candidate ERVs were identified in each reference genome assembly using RetroTector with scores of ≥300[47] and a Perl script (http://doua.prabi.fr/software/one-code-to-find-them-all). The resulting ERV loci were merged and further filtered based on the presence of two flanking LTRs and a table of ERVs and their associated LTRs, as described by Kojima et al.[48] To examine species-specific ERVs, we downloaded liftover files from the human genome to the genomes of five apes and repeated the process for each of the other apes. In total, we used 30 liftover files from the Genome Archive collection of UCSC Genome Browser[49]. Reciprocal liftover analysis was conducted to infer the presence/absence of each two LTR ERVs across six primate genomes, using the parameter -minMatch=0.1 (minimum ratio of bases that must remap). Lifted ERVs shared at orthologous genomic positions were deemed ancestral and likely fixed within each species[50] and, thus, were filtered out. These sequences were further analyzed for target site duplications on each side of the ERVs using BLAT. To generate a comprehensive catalog of *gag* (capsid domain) and *pol* (reverse transcriptase and integrase) domains, we performed a six-ORF translation of two LTR ERVs using ORFfinder with the parameters -ml 300 (minimum ORF size 100 codons) and -s 1 (use the standard genetic code but allowing noncanonical start codons)[47]. Short, encoded proteins were concatenated with "N" connection based on coordinates. For the capsid CA domain homology within the *gag*, we utilized hmmscan following a previously described method[51]. We employed HMMER with default parameters, a bit-score threshold of 25, and a length threshold of 125 amino acids[52]. For reverse transcriptase and integrase domain homology within the *pol*, the domains of all collected proteins were predicted using InterProScan[53]. Additionally, we collected all ERV proteins annotated in Repbase and the Repeat Protein Database and performed InterProScan and CD-BLAST on previously characterized consensus sequences of ERVs.

To determine intact ORFs across lineage-specific full-length L1s, we followed a previously described method[54] to detect intact ORF1p and ORF2p. To account for the millions of years of divergence between species, we lowered the amino acid similarity cut off from 5% to 15%. Once we obtained ORF calls, to curate the lower 15% threshold, we compared intact ORFs from 6-15% divergence across human assemblies and found evidence for downstream ORF2p usage, but these putative proteins retained the function of catalytic domains of ORF2p. Therefore, the thresholds we set are appropriate for proper identification of protein-coding ORFs that may be retrotransposition competent.

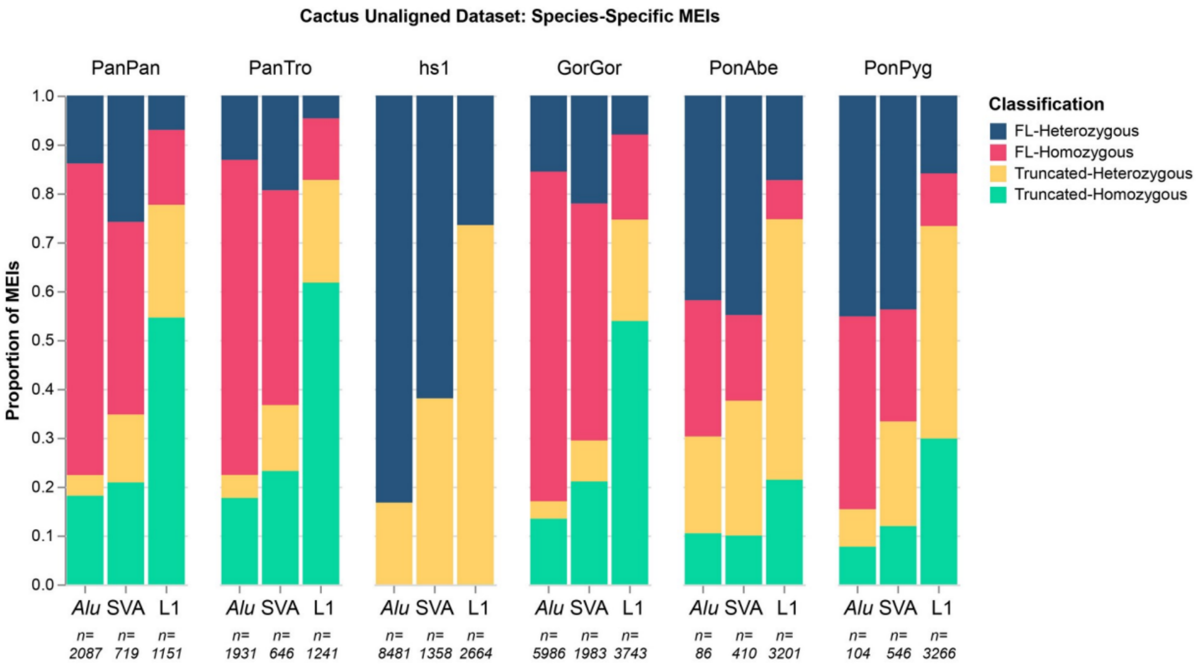


**Figure Species-Specific MEI Classifications.** A stacked bar graph showing the proportion of species-specific MEIs that are both full length (FL) and homozygous (magenta), FL and heterozygous (dark blue), truncated and homozygous (green), and truncated and heterozygous (yellow). MEIs were not checked for homozygosity within humans, only FL or truncated, as hs1 is a haploid genome. The PonAbe and PonPyg genomes show a larger proportion of heterozygous LINE/L1s compared to the African great apes suggesting an elevated retrotransposition rate of L1s in the Pongo lineages.

##### Nuclear sequences of mitochondrial DNA origin (NUMT)

NUMTs are not included in the RepeatMasker annotation; therefore, we used a different approach to identify them. We identified NUMTs by aligning the T2T mitochondrial assemblies to nuclear assemblies for each species, including T2T and non-T2T assemblies. We used BLASTN aligner[55] with parameters previously used to study NUMTs in the human T2T-CHM13 assembly (-evalue 0.0001 -gapopen 5 -gapextend 2 -penalty -3 -reward 2 -task blastn)[56]. Overlapping alignments and those within 1 kbp of each other were merged using BEDTools, as they likely represent the same NUMT event. The following non-T2T assemblies were used for comparison: panPan3[35], GRCh38[57], gorGor6, panTro6, and ponAbe3[58]. The panPan3, gorGor6, and ponAbe3 assemblies of female subjects were supplemented with a Y chromosome (NCBI accessions GCA_015021855.1, GCA_015021865.1, and GCA_015021835.1, respectively)[59].

### IX. Selection analyses within NHP lineages

Contributing authors:

Abigail N. Sequeira, Qiuhui Li, Arjun Biddanda, Rajiv McCoy, Michael Schatz, Michael Tassia, Zachary A. Szpiech, Christian D. Huber, Kateryna D. Makova

#### Methods

##### Read mapping and variant calling

We performed read alignment and variant calling using 129 samples described in the T2T ape sex chromosome study[1]. For the T2T references, we generated karyotype-specific references following a published masking method[6] to improve variant representation in sex chromosome pseudoautosomal regions. Reads from XX and XY samples were aligned to their respective masked T2T reference genomes. Variant identification followed the T2T-chrXY ape paper method using GATK v4.4.0.0 HaplotypeCaller[60] for initial calling, GenotypeGVCFs for joint genotyping, and applying standard quality control to SNPs and indels.

##### Haplotype phasing and curation

Haplotype phasing was performed across all primate T2T autosomal genomes using BEAGLE v4.0[61] (*impute=false nthreads=8 burnin=4 iterations=12 seed=42*). In all cases the effective population size (*Ne*) and error parameters were estimated on a per-taxa and per-chromosome level prior to phasing. No reference panel was used during the phasing process (https://github.com/aabiddanda/haplotype-phasing). We filtered the phased VCFs for bi-allelic sites that fell within high-confidence regions, resulting in the removal of less than 1% of called SNPs for each species (**Table SelectionS1**). We ran two different selective sweep detection methods, SweepFinder2[62] and saltiLASSI[63], for 10 great ape taxa: bonobo, Bornean orangutan, central chimpanzee, eastern chimpanzee, eastern lowland gorilla, mountain gorilla, Nigerian chimpanzee, Sumatran orangutan, western chimpanzee, and western lowland gorilla. We excluded the Cross River gorilla from the subsequent analyses as it had a sample size of one.

##### SweepFinder2

We generated non-reference allele frequency files for each population, excluding positions that were monomorphic or did not contain the non-reference allele from the filtered VCFs. We removed between 32,675 and 2,337,860 variants and retained between 4,467,345 and 24,868,785 bi-allelic SNPs across all 10 taxa (**Table SelectionS2**). We used SweepFinder2 to first calculate the whole-genome site frequency spectrum (SFS) for each population using the whole-genome allele frequency file (*SweepFinder2 -f WG.freq.file SFS*). Then using this pre-calculated SFS, we ran SweepFinder2 to calculate a likelihood ratio score along a 1 kbp grid for each autosome for each population (*SweepFinder2 -lg 1000 Chr.freq.file SFS Out.file*).

**saltiLASSI**

We computed the saltiLASSI statistic with lassip (v1.2.0) for each taxon in three steps. The first step created window-based spectra files for each autosome following these parameters: a 201 SNP window size, a 100 SNP step size, a *–k* of five or ten to estimate the haplotype frequency spectrum (HFS), *–salti*, and *–unphased*. Because *–k* cannot be larger than the sample size (*n*), if *n* > 10, *k* was set to ten and if *n* < 10, *k* was set to five. The second step calculates the mean genome-wide HFS using the window-based spectra files generated for each autosome. The final step calculates the saltiLASSI likelihood ratio score for each autosome using the genome-wide HFS and the corresponding window-based spectra file. Together, the code was written as such:

> lassip –vcf taxon.chr.vcf –pop taxon.IDs.txt –winsize 201 –winstep 100 -k 10 –calc-spec –hapstats –salti –unphased –out taxon.chr.spectra
>
> lassip --spectra taxon.chr*.spectra --avg-spec --out taxon.avg.spec.
>
> lassip --spectra taxon.chr.spectra --salti --null-spec taxon.avg.spec --out taxon.chr.final.out

**Determining candidate sweeps**

For the raw SweepFinder2 and saltiLASSI results, we filtered out positions for which a likelihood ratio was calculated but fell outside an accessibility mask generated for each reference species. Then, we downloaded gene annotation files for each reference from NCBI **(**GCF_029281585.2, GCF_029289425.2, GCF_028858775.2, GCF_028885655.2, GCF_028885625.2) and filtered the annotation files for the protein-coding biotype and for entries listed as “gene”. For each gene, we added a 50 kbp flank to the start and end position to capture signals in potential regulatory sequence of each gene. Finally, we paired each position for which a likelihood ratio score was calculated with the corresponding gene and found the maximum likelihood score for each gene so that every gene has a single representative score.

To determine significant sweep regions from SweepFinder2, we normalized the gene-specific score distribution according to a procedure described in Souilmi et al[64]. We first log transformed the maximum likelihood statistic for each gene. Next, we binned these scores based on gene length and performed a robust Z-transformation. Finally, we calculated the p-values for each Z-score, assuming a standard normal distribution. We estimated the false discovery rate (FDR) and corresponding q-values using the R package qvalue[65] (v2.34.0). Next, we defined sweep regions by combining genes that had a q-value of 0.1 or smaller and were within 1 Mbp of each other, to take into account that single sweep signals often span multiple genes. Lastly, we filtered out any sweep region that did not contain at least one gene with a q-value of ≤0.01.

We took an outlier approach for identifying significant sweeps with saltiLASSI. However, because the likelihood statistic in the MHC region for each taxon was substantially higher than other likelihood statistics and was potentially caused by strong balancing selection and not positive selection, we took the top 0.1 percentile of the likelihood statistic before and after filtering out the MHC region. Finally, we combined the two to have one single dataset for each species. We again concatenated genes that were within 1 Mbp of each other to determine sweep regions.

**Fst**

Fst outlier peaks across the genome often reflect regions evolving under local adaptation. Therefore, we took the top 0.1% of Fst values between central and eastern chimpanzees (Fst = 0.09). We did not further investigate other taxa pairs because they were either fairly diverged (Fst > 0.21) or there were no clear Fst peaks across the genome-wide distribution. We assigned genes to the Fst peak regions and compared these regions to sweep regions identified by SweepFinder2 and saltiLASSI.

**Gene enrichment**

We performed a gene enrichment analysis with GOWINDA[66] on each taxon that had candidate sweeps called for SweepFinder2, saltiLASSI, and the top 0.1% Fst regions for central and eastern chimpanzees. GOWINDA requires four files: a whole-genome annotation as a .gtf, a file with the total number of SNPs, a file of candidate SNPs, and a gene set file. However, instead of providing SNP files, we provided the positions of the calculated likelihood ratio scores or the center of the Fst and saltiLASSI windows. Using the python scripts provided by the GOWINDA package, we converted each annotation file from a .gff to a .gtf. We downloaded a human gene set file from FuncAssociate 3.0[67] as recommended by GOWINDA. The gene set file contained the HGNC IDs for each gene associated with their respective GO category. For the total SNP file, we used the raw genome-wide positions (chromosome and position) from SweepFinder2, saltiLASSI, and Fst calculations. The candidate SNP files consisted of the positions for which the likelihood ratio or Fst window midpoint fell within the candidate sweep regions or was a top 0.1% of Fst value. We ran GOWINDA using *–mode gene* and *–gene-definition updownstream50000*. The full command was as follows: *java -Xmx4g Gowinda-1.12.jar –snp-file sel.scan.pos.txt –candidate-snp-file candidate.SNP.txt –gene-set-file funcassociate.go.txt –annotation-file annotation.gtf –simulations 100000 –min-significance 1 –mode gene –min-genes 1 –gene-defintion updownstream50000 –threads 8 –output-file out.GO.enrichment.txt*. We considered GO terms that had an FDR of ≤0.1 to be significantly enriched.

**Table SelectionS1: Number of positions removed and retained after filtering for bi-allelic SNPs.**

| Taxa | Number of Positions Removed | Final Number of Positions |
|---|---|---|
| Bonobo | 3,045,253 | 8,196,463 |
| Central Chimpanzee | 5,790,571 | 25,929,241 |
| Eastern Chimpanzee | 5,430,524 | 17,413,277 |
| Nigerian Chimpanzee | 5,145,851 | 10,418,097 |
| Western Chimpanzee | 4,176,231 | 5,325,379 |
| Eastern Lowland Gorilla | 3,244,557 | 6,805,205 |
| Mountain Gorilla | 3,697,135 | 7,439,936 |
| Western Lowland Gorilla | 4,618,357 | 16,338,062 |
| Bornean Orangutan | 3,881,414 | 9,512,242 |
| Sumatran Orangutan | 5,420,219 | 1,4677,131 |

**Table SelectionS2: Breakdown of the number of positions removed and kept for the allele frequency files generated for SweepFinder2.**

| Taxa | Number of Positions Removed | Final Number of Positions |
|---|---|---|
| Bonobo | 89,667 | 8,106,796 |
| Central Chimpanzee | 1,060,456 | 24,868,785 |
| Eastern Chimpanzee | 1,188,738 | 16,224,539 |
| Nigerian Chimpanzee | 825,808 | 9,592,289 |
| Western Chimpanzee | 68,734 | 5,256,645 |
| Eastern Lowland Gorilla | 2,337,860 | 4,467,345 |
| Mountain Gorilla | 2,117,885 | 5,322,051 |
| Western Lowland Gorilla | 32,675 | 16,305,387 |
| Bornean Orangutan | 308,627 | 9,203,615 |
| Sumatran Orangutan | 503,784 | 14,173,347 |

**Summary of results**

To identify regions harboring population genetic signatures of adaptation in 10 great ape taxa, we used two complementary methods. SweepFinder2 scans for regions exhibiting distorted allele frequency patterns characteristic of a fixed hard sweep (i.e., an excess of low- and high-frequency alleles), whereas saltiLASSI scans for distorted haplotype frequency patterns indicative of a soft or partial sweep. Across all taxa, we identified 143 and 86 candidate regions using SweepFinder2 and saltiLASSI, respectively. Only two candidate regions overlapped between the two methods (**Table SelectionS4**), consistent with their sensitivities for detecting distinct modes of positive selection.

We next performed a gene set enrichment analysis for GO terms in the sweep regions. We found significant enrichment for genes involved in pathways related to diet (sensory perception for bitter taste, lipid metabolism, and iron transport), immune function (antigen/peptide processing, MHC-I binding), cellular activity, and oxidoreductase activity in bonobos, central and eastern chimpanzees, and western lowland gorillas.

Selection signatures were strongest in the MHC region, a gene-rich locus previously described as a target of strong selection, especially balancing selection[68]. Earlier studies, however, suggested that an ancient MHC-I sweep in the bonobo and central-eastern chimpanzee ancestor results from an adaptation to simian immunodeficiency virus (SIV)-like retroviruses[69-71]. We found evidence of long-term balancing selection on MHC in multiple great ape lineages, including central and eastern chimpanzees, as well as at least two regions in MHC consistent with positive selection in bonobos and western chimpanzees.

Genes encoding bitter taste receptors in primates have been well documented to have undergone species-specific adaptation, especially in chimpanzees[72,73] and gorillas[74]. In agreement with this, we detected significant enrichment in selection signals for such genes in bonobos (*TAS2R3*, *TAS2R4*, *TAS2R5*) and western lowland gorillas (*TAS2R14*, *TAS2R20*, *TAS2R50*), as well as identified a bitter taste receptor gene (*TAS2R42*) within a sweep region in eastern chimpanzees.

To assess the impact of selective sweeps on genome-wide genetic differentiation, we examined $F_{ST}$ values within and outside of SweepFinder2 sweep regions in recently diverged eastern and central chimpanzee subspecies. Notably, sweep regions in both subspecies exhibited significantly higher differentiation ($F_{ST}$ = 0.21 and 0.15, Mann-Whitney $p < 0.001$) compared to the genome-wide average ($F_{ST}$ = 0.09). No increased differentiation was observed within saltiLASSI sweep regions (Mann-Whitney $p > 0.05$). These findings suggest that hard selective sweeps play an important role in shaping genomic variation across eastern and central chimpanzees.

**Selection scans**

To call selective sweeps, we ran two sweep detection methods: SweepFinder2 and saltiLASSI. SweepFinder2 is an SFS-based tool that can detect hard sweeps, whereas saltiLASSI is based on an HFS and can detect hard, soft, and partial sweeps. For determining SweepFinder2 sweep regions, we used a q-value approach in which we concatenated genes with q-value < 0.1 that are within 1 Mbp of each other and called sweep regions significant if they contained at least one gene with a q-value of 0.01 or lower. SweepFinder2 detected sweeps in five out of 10 analyzed great ape taxa: bonobo, central, eastern, and western chimpanzee, and western lowland gorilla (30, 22, 62, 11, and 18 sweeps, respectively; **Table SelectionS3**). Unsurprisingly, these taxa had larger sample sizes (*n* = 13, 18, 19, 11, and 27, respectively). We took an outlier approach for determining sweeps with the saltiLASSI method and identified 4-18 selective sweeps across all 10 taxa (**Table SelectionS3**). We found minimal overlap between the sweep regions called by SweepFinder2 and saltiLASSI. However, we found one sweep region called by both SweepFinder2 and saltiLASSI on chromosome 10 in western lowland gorillas, one partially overlapping sweep called by both methods on chromosome 5 in bonobos, and one sweep region called on chromosome 3 in central and eastern chimpanzees (**Table SelectionS4**). Each method also identified at least one sweep region occurring in between the various chimpanzee subspecies; SweepFinder2 classified two sweeps in central and eastern chimpanzees and saltiLASSI classified a sweep in central and Nigerian chimpanzees (**Table SelectionS4**). We compared our sweep regions to those from previous literature by identifying at least one common candidate gene and corroborated sweep signals in bonobos[75], central, eastern, and western chimpanzees[75,76], and western lowland gorillas[74,75]. We identified 75 and 70 novel sweep regions via SweepFinder2 and saltiLASSI, respectively, as well as a total of 43 regions that were previously found in humans[75,77].

Beyond these two selection scan methods, we also assessed the top 0.1% of Fst values computed between central and eastern chimpanzees (**Table SelectionS4**). There were no overlaps between the top 0.1% Fst values and saltiLASSI sweeps, but we found overlaps for five SweepFinder2 sweeps in eastern chimpanzees. These sweeps were located on chromosomes 1, 5, 10, and 18 (**Table SelectionS4**). Furthermore, four out of five of the sweeps contained genes in significantly enriched pathways (see below for more detail).

The MHC, in particular, showcased a complex selection signature (**Fig. SelectionS1-3**). The MHC is a gene-dense region that is subject to heavy selective pressure, especially for balancing selection[68]. Overall, we observed strong saltiLASSI peaks in either the MHC class I or II region in eight out of 10 taxa (central, eastern, and western chimpanzees, all three gorilla subspecies, bonobos, and Bornean orangutans), with six (central, eastern, and western chimpanzees, eastern and western lowland gorilla, and bonobos) being significant and ranking among the highest for peak strength (**Fig. SelectionS1-3**). Moreover, we observed overlapping positive peaks of Tajima's *D* and nucleotide diversity for bonobos, central, eastern, and western chimpanzees, western lowland gorillas, and Bornean orangutans (**Fig. SelectionS1-3**). However, in eastern lowland gorillas we observed a negative Tajima's *D* and increased nucleotide diversity and in mountain gorillas we observed a positive Tajima's *D* and a decreased nucleotide diversity, giving conflicting signatures of selection (**Fig. SelectionS2**). Given the results from Tajima's *D* and the nucleotide diversity estimates, it is likely that saltiLASSI was picking up balancing selection signals rather than positive selection in all taxa except for eastern lowland and mountain gorillas. In the case of SweepFinder2, we observed evidence for hard sweeps in bonobos and western chimpanzees (**Fig. SelectionS1D and S3A**). Broadly, we find that there is a complex selection signature occurring in the MHC region across the various great ape species where clear balancing selection is observed in many of the great ape lineages and at least two instances consistent with positive selection, similar to previous findings[75,78].

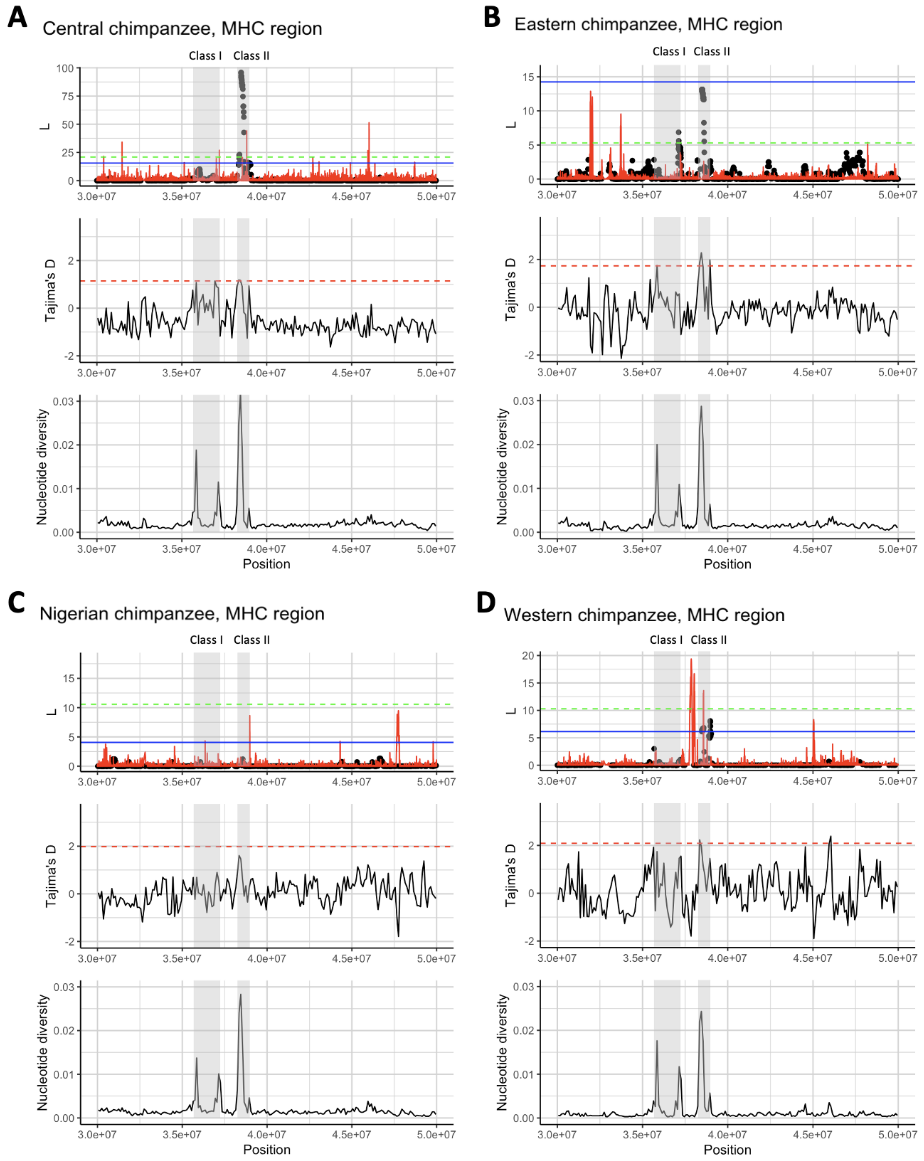


**Figure SelectionS1. Selection scans, Tajima's *D*, and nucleotide diversity for the chimpanzee MHC region.** (A-D) Gray boxes represent the locations of MHC class I and II genes. Top panel: SweepFinder2 (red solid line) and saltiLASSI (black dots) likelihood ratio scores are plotted along genomic position. SweepFinder2 results for eastern, Nigerian, and western chimpanzees were scaled down by a factor of 10. Central chimpanzee SweepFinder2 results were not scaled down. The blue solid line marks the top 0.1% of saltiLASSI results and the green dotted line marks the top 0.1% of SweepFinder2 peaks, scaled down for eastern, Nigerian, and western chimpanzees. saltiLASSI peaks that are above the blue line were counted as candidate sweeps. Middle panel: Tajima's *D* calculated across 100 kbp windows. The red dotted line represents the top 0.1% of values. Bottom panel: Nucleotide diversity calculated across 100 kbp windows.

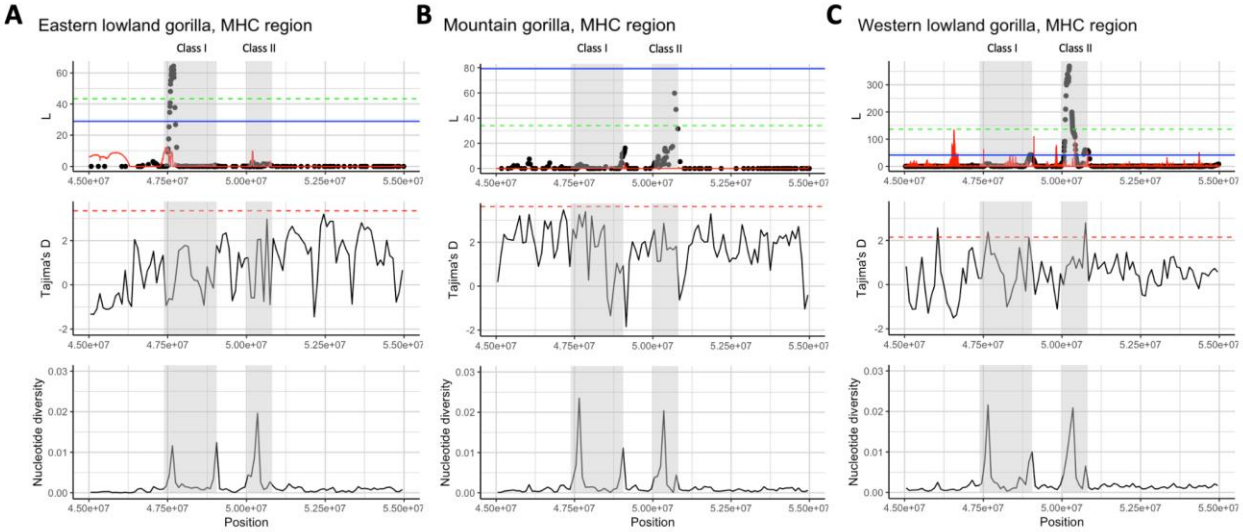


**Figure SelectionS2. Selection scans, Tajima's *D*, and nucleotide diversity for the gorilla MHC region.** (A-C) Gray boxes represent the locations of MHC class I and II genes. Top panel: SweepFinder2 (red solid line) and saltiLASSI (black dots) likelihood ratio scores are plotted along genomic position. SweepFinder2 results for eastern lowland gorilla and mountain gorilla were scaled down by a factor of 100. Western lowland gorilla SweepFinder2 results were not scaled down. The blue solid line marks the top 0.1% of saltiLASSI results and the green dotted line marks the top 0.1% of SweepFinder2 peaks, scaled down for eastern lowland gorilla and mountain gorilla. saltiLASSI peaks that are above the blue line were counted as candidate sweeps. Middle panel: Tajima's *D* calculated across 100 kbp windows. The red dotted line represents the top 0.1% of values. Bottom panel: Nucleotide diversity calculated across 100 kbp windows.

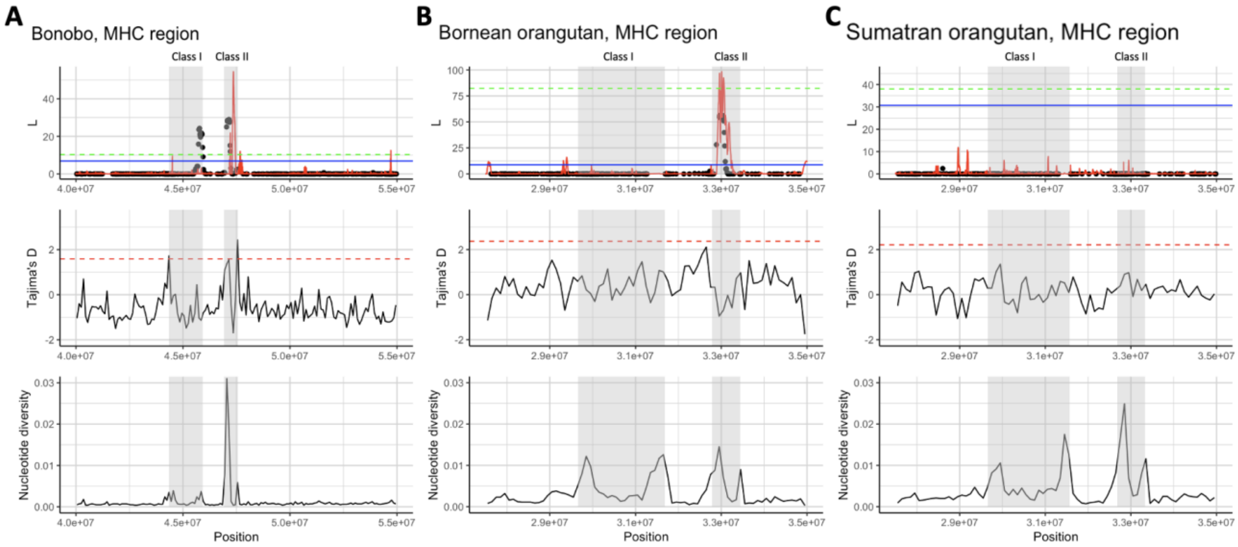


**Figure SelectionS3. Selection scans, Tajima's *D*, and nucleotide diversity for the bonobo and orangutan MHC regions.** (A-C) Gray boxes represent the locations of MHC class I and II genes. Top panel: SweepFinder2 (red solid line) and saltiLASSI (black dots) likelihood ratio scores are plotted along genomic position. SweepFinder2 results for bonobo, Bornean and Sumatran orangutans were scaled down by a factor of 10. The blue solid line marks the top 0.1% of saltiLASSI results and the green dotted line marks the top 0.1% of SweepFinder2 scaled peaks. saltiLASSI peaks that are above the blue line were counted as candidate sweeps. Middle panel: Tajima's *D* calculated across 100 kbp windows. The red dotted line represents the top 0.1% of values. Bottom panel: Nucleotide diversity calculated across 100 kbp windows.

Following the selection scans, we ran a gene set enrichment analysis via GOWINDA to test if genes with certain GO terms are enriched in the sweep regions. We considered any GO term that had an FDR of 0.1 or smaller to be significantly enriched. Out of the 10 taxa, only bonobos, central and eastern chimpanzees, and western lowland gorillas had significantly enriched GO terms. Sweep regions detected by SweepFinder2 were enriched in bonobos, eastern chimpanzees, and western lowland gorillas while regions identified by saltiLASSI were found to be enriched in central chimpanzees (**Table SelectionS5**). Of the top 0.1% Fst peaks between central and eastern chimpanzees, we find enrichment for genes related to the regulation of epidermal cell division (**Table SelectionS5**). Of note, the two genes in this gene set both fall in the same sweep region detected by SweepFinder2 in eastern chimpanzees, suggesting species-specific differentiation in these two genes between central and eastern chimpanzees. Across all three selection tests, enrichment was found for genes involved in diet (sensory perception for bitter taste, lipid metabolism, and iron transport), immune function (antigen/peptide processing, MHC class I binding), cellular activity, and oxidoreductase activity (**Table SelectionS5**).

#### Diet-related function

Among the enriched GO terms, we identified several pathways related to diet. Pathways involving lipid metabolism, oxidoreductase activity, and iron transport were found for genes within eastern chimpanzee sweeps (**Table SelectionS5**). Chimpanzees are highly frugivorous omnivores[79-81]. Moreover, several studies, particularly in eastern chimpanzees, have reported them to engage in geophagy (intentional eating of soils) of termite mounds and clay-infused water[82-85]. It is hypothesized that these are adaptive behaviors to either provide protection against plant toxins and parasites or to supplement essential elements such as iron[86]. While these behaviors are not specific to eastern chimpanzees—indeed geophagy is found across a wide variety of taxa among and outside of primates[86,87]—the iron transport pathway was only significantly enriched in eastern chimpanzees, pointing to recent adaptation to dietary iron availability in this subspecies.

The oxidoreductase pathway encompasses a wide array of enzymes that are crucial for maintaining cellular homeostasis, energy production, biosynthesis, detoxification, and signaling. Notably, all but one of the 12 candidate genes that are in the oxidoreductase activity pathway are strongly connected to diet or diet-related disease. These 11 genes are involved with androgen metabolism, aldehyde oxidation, breaking down fatty acid chains, and vitamin metabolism. Furthermore, these have been associated with nonalcoholic fatty liver disease[88,89] and obesity[90,91].

Genes encoding for bitter taste receptors (TAS2Rs or T2Rs) have an interesting evolutionary history. In primates, T2Rs have been well documented to have undergone species-specific modes of selection, especially in chimpanzees[72,73] and gorillas[74]. Upon examining the sweep identified by both SweepFinder2 and saltiLASSI on chromosome 10 in western lowland gorillas, we find three bitter taste receptor genes (*TAS2R14*, *TAS2R20*, and *TAS2R50*) within the sweep region, corroborating previous findings[74]. In chimpanzees, Hayakawa et al.[72] conclude that balancing selection was the main driver for western chimpanzee taste receptor evolution and purifying selection in the human TAS2R cluster in eastern chimpanzees. Notably, our results find a selective sweep pattern in eastern chimpanzees harboring *TAS2R42* (**Table SelectionS3**), indicating that positive selection in bitter taste reception might also play a role in chimpanzees. We also identified one significant sweep region in bonobos that contain TAS2R genes (**Table SelectionS3**). In sum, our results suggest local adaptation at taste receptor genes for western lowland gorillas, eastern chimpanzees, and bonobos.

#### MHC region and immune-related function

As stated above, we observed complex selection patterns in the MHC region. In the case of bonobos and chimpanzees, previous hypotheses have postulated that an ancient sweep occurred in the ancestor of bonobos and chimpanzees in MHC-I driven by an adaptation to better combat a SIV-like retrovirus[69-71]. However, Pawar et al.[92] did not find evidence for an ancient sweep in the central-eastern chimpanzee ancestor (possibly due to lack of power), instead suggesting recent balancing selection in the defense against SIV infection in central and eastern chimpanzees[92,93] as the more likely explanation for MHC genetic patterns. Our results show evidence for both positive and balancing selection in bonobos in the MHC class I and II regions (**Table SelectionS1; Fig. SelectionS1A**), but only evidence for balancing selection (see above discussion) for central and eastern chimpanzees (**Fig. SelectionS1A-B**). Similar to Pawar et al.[92], we do not see evidence of positive selection in central and eastern chimpanzees in the MHC-I region but instead two large peaks of diversity at the left and right ends of the region (**Fig. SelctionS1**). However, we do see positive selection signatures in western chimpanzees between the class I and class II gene regions (**Table SelectionS1; Fig. SelectionS1D**).

Gene sets related to MHC protein-binding and antigen processing were significantly enriched sweep regions in bonobos (**Table SelectionS5**). While there were several different genes included in the enriched pathways, *TAP1* and *TAP2* were found in every MHC-related pathway for bonobos (**Table SelectionS5**). *TAP1* and *TAP2* are viral interacting proteins (VIPs) harboring variation associated with risk of respiratory infection (e.g., by influenza)[94] and interact with herpes and pox viruses[95] in humans. In a previous study, Enard et al.[96] reported increased rates of adaptation in VIPs in mammal lineages. This sentiment is echoed by Pawar et al.[92] who found selection on VIPs involved with herpes and influenza, among other viruses, in gorillas. Our results indicate local adaptation at VIPs in bonobos, adding support to the hypothesis that viruses are a major driver of protein adaptation in mammals.

**Fst increased within SweepFinder2 sweeps, but not within saltiLASSI sweeps**

We compared Fst values outside and within eastern chimpanzee sweep regions and found an increase in Fst within sweep regions (meanout = 0.09, meanwithin = 0.21, Mann-Whitney, z = -14.35, p < 2.2e-16; **Fig. SelectionS4**). We see a similar significant increase in Fst within central chimpanzee sweeps (meanout = 0.09, meanwithin = 0.15, Mann-Whitney, z = -5.44, p = 5.19e-8; **Fig. SelectionS4**), but the increase is not as high. However, eastern chimpanzees have significantly higher Fst values within sweeps compared to central chimpanzees (Mann-Whitney, z = -3.40, p < 0.001). Moreover, the maximum Fst value within central chimpanzee sweeps was 0.42 whereas the maximum Fst value for eastern chimpanzees was 0.67. For saltiLASSI sweeps, we observe the opposite pattern with Fst decreasing within eastern chimpanzee sweep regions (meanout = 0.09, meanwithin = 0.07, Mann-Whitney, p = 0.047; **Fig. SelectionS5**), but no difference for central chimpanzee sweep regions (meanout = 0.09, meanwithin = 0.09, Mann-Whitney, p = 0.63; **Fig. SelectionS5**). Collectively, this suggests that local adaptation plays a large role in subspecies differentiation between central and eastern chimpanzees.

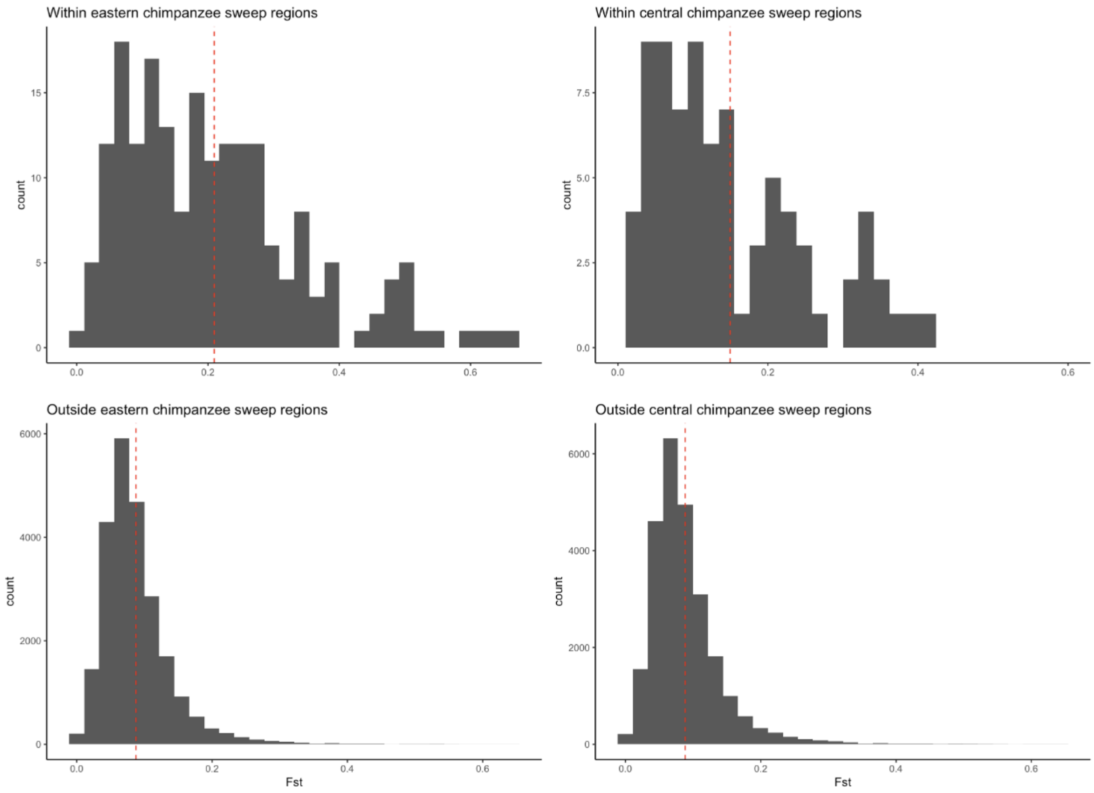


**Figure SelectionS4. Central and eastern chimpanzee Fst distribution within and outside of SweepFinder2 sweep regions.** Histograms comparing the *Fst* distributions within and outside of central and eastern chimpanzees. The red dashed lines mark the mean *Fst* values. The mean *Fst* for outside sweep regions for both subspecies was the same as the genome-wide average (*Fst* = 0.09). The mean *Fst* within sweeps for central and eastern chimpanzees (*Fst* = 0.15 and 0.21, respectively, $p < 0.001$) were significantly larger than outside sweep regions.

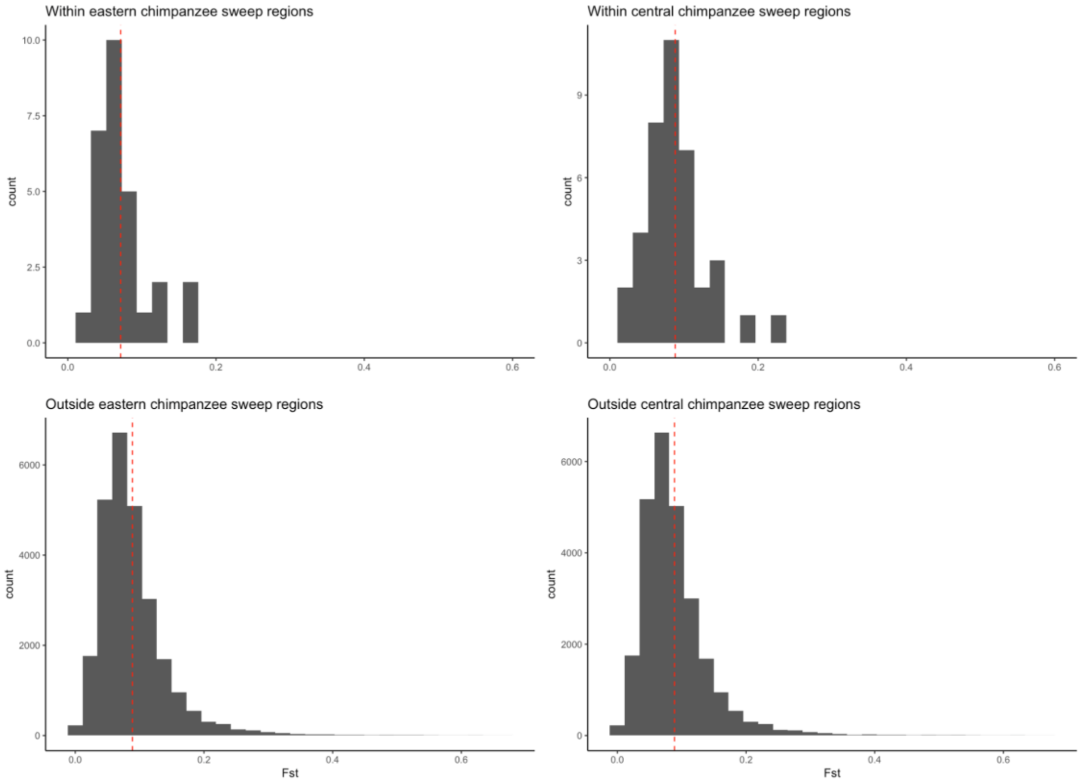


**Figure SelectionS5. Central and eastern chimpanzee Fst distribution within and outside of saltiLASSI sweep regions.** Histograms comparing the *Fst* distributions within and outside of central and eastern chimpanzees. The red dashed lines mark the mean *Fst* values. The mean *Fst* for outside sweep regions for both subspecies was the same as the genome-wide average (*Fst* = 0.09). The mean *Fst* within central chimpanzee sweep regions was not significantly different (*Fst* = 0.09) and the mean *Fst* within eastern chimpanzee sweeps was significantly smaller (*Fst* = 0.07, $p = 0.047$).

### X. Immunoglobulin annotation and analysis

Contributing authors:

Yana Safonova, Corey T. Watson, Anton Bankevich, Matt Pennell, Yixin Zhu, Swati Saha, William Lees, Eric Engelbrecht, Pavel Pevzner, Ishaan Gupta, Zhenmiao Zhang

#### Methods

Analyses of the immunoglobulin (IG) heavy (IGH), light chain kappa (IGK) and lambda (IGL), T-cell receptor (TR) beta (TRB), TR alpha/delta (TRA/D), and gamma (TRG) loci in four ape species (bonobo, gorilla, Bornean orangutan, and Sumatran orangutan) for which two complete intact haplotypes were constructed were conducted. Genomes of two species (chimpanzee, siamang gibbon) derived from lymphoblastoid cell lines contain somatic rearrangements driven by V(D)J recombination and were excluded from the analysis.

##### Annotating germline IG/TR genes and loci

Germline immunoglobulin (IG) and T-cell receptor (TR) variable (V), diversity (D), and joining (J) genes were predicted using the IgDetective[97] and Digger[98] tools. Digger was run using human recombination signal sequences for the IG and TR genes, with output from IgDetective as the starting germline database. Boundaries of IG and TR loci were defined according to the leftmost and rightmost IG genes. We assessed per base read support of the assemblies spanning the IG and TR loci. To ensure base-level accuracy in the genomic assemblies and read support of the predicted germline IG and TR genes, PacBio HiFi reads were remapped to each T2T genome using minimap2[27], followed by analysis of per-base read support using SAMtools[99]. In each of the four species (Bornean orangutan, bonobo, gorilla, and Sumatran orangutan), we found that the mean coverage of remapped HiFi reads to each respective haplotype assembly ranged from 28 to 75. Additionally, 99.9% of assembly bases for the IG and TR loci were supported by at least 80% of the mapped reads, with 100% of IG/TR gene-coding bases supported at this read support level (**Table IG.S3**). For resulting V gene annotations and presented analyses, subfamilies were assigned according to the closest human V gene. Given the patterns of divergence and haplotype complexity in the IG/TR loci observed among species, the grouping and assignment of homologous genes among haplotypes and species is nontrivial. We determined that this will require more detailed phylogenetic and comparative genomic analysis of additional haplotypes in each species. Consequently, we determined that the use of standard gene identifiers based on position and cross-species orthology was not warranted. Assignments of permanent identifiers to each unique germline sequence in each species is currently under review by the International Union of Immunological Societies TR-IG Nomenclature Review Committee (https://iuis.org/committees/nom/nomenclature-sub-committees/immunoglobulins-ig-t-cell-receptors-tr-and-major-histocompatibility-nomenclature-sc/).

##### Finding SD blocks

We decomposed IG/TR loci into the alphabet of duplication subunits[100] using a modification of the Sibelia tool[101] that uses iterative de Bruijn graphs for analyzing synteny blocks and SDs. This modification enabled analysis of highly repetitive IG/TR loci that result in particularly complex de Bruijn graphs where the original Sibelia algorithm has limitations. The constructed block decompositions enabled comparison of segmental/tandem duplications in IG/TR loci between different haplotypes of the same species.

##### Finding units of *IGHV3-30* and *IGHV4-59* tandem units

To reveal units of tandem duplications containing *IGHV3-30*-like and *IGHV4-30*-like V genes in the IGH loci across species, SDs found in the bonobo IGH loci and the human IGH T2T locus were identified and aligned to all IGH loci in the other three species; the procedure was repeated until all units were detected. The detected units were numbered according to their order in the corresponding locus/haplotype. The phylogenetic tree of the units was computed using the ClustalW2 tool[102] and visualized using the Iroki tool[103]. The same procedure was applied to tandem duplications containing *IGHV1-58*-like and *IGHV4-59*-like V genes in IGH loci of the Sumatran and Bornean orangutans.

##### Comparative analysis of IG/TR loci and genes

Pairs of IG/TR germline sequences (using the IgDetective gene sets) of all loci were aligned using YASS[104] where the longest nonoverlapping alignments were selected to visualize the alignment blocks (as shown in **Fig. 3a** and **Fig.IG.S1a**). The repetitiveness of a locus was computed as the fraction of bases among the total bases in that haplotype that were spanned by repetitive sequence of length at least 10 kbp. To define “ape-specific” V genes within a locus, the V genes were combined with known human V genes from the same locus and the nucleotide distances between all pairs of genes were computed. The human set used included all *01 alleles for every curated human functional, ORF, and in-frame pseudogene IG and TR gene in the International IMmunoGeneTics Information System (IMGT) database[105] (imgt.org; date downloaded: July 1, 2024). The hierarchical clustering maximizing the number of clusters consisting of at least three nonhuman genes was applied, and genes corresponding to these clusters were reported as ape-specific. Similarly, human-specific genes were computed in comparison with bonobo V genes and were identified in clusters consisting of at least two human genes. To estimate the allelic diversity of curated IG/TR genes within a species and locus, for each V gene, the closest V gene (by sequence alignment) from the alternative haplotype of the same locus and the same species was found. Distances between identified pairs of genes were collected across all six loci and four species. The pairs of genes with zero distance were referred to as identical.

#### Summary of results

##### Gene annotation

We compiled complete IG/TR gene annotation sets for each species, including annotations for functional and ORF V, D, and J genes/alleles. Annotation sets from IgDetective and Digger are provided in **Table.IG.S1** and **Table.IG.S2**, respectively, including positions within each haplotype, and in the case of Digger annotations, predicted upstream regulatory sequences, leader sequences, introns, and recombination signal sequences. Consolidated sets for each species and locus will be made available via OGRDB[106].

##### Locus architecture and gene divergence of the ape IG and TR loci

As shown in **Fig. 3a** for the IG loci, we computed within and between species haplotype alignments for each of the TR loci (**Fig.IG.S1.a**). These initial comparisons revealed a greater degree of synteny between species, and less structural variation within and between species relative to that observed for the IG loci (see below). To quantify this, we computed within haplotype SD blocks (see methods above) and calculated the percent of bases covered by SD in each locus and haplotype. On average, we found that across species, the IG loci had a consistently high fraction of bases spanned by SDs. In contrast, while both TRB and TRG loci had comparable levels of SD, the fraction of bases covered by SD in the TRA/D locus was lowest among all species (**Fig.IG.S2.a**). IGK showed the greatest degree of variation in this computed fraction, and also the greatest degree of inter-haplotype length variation mainly associated with long insertions (**Fig.IG.S2.b**). Additionally, several tandem duplication regions were associated with length differences between haplotypes both within and between species in IG loci. As an example, we conducted inter-haplotype and inter-species analyses of two regions in the IGH locus that harbor expanded tandem duplication blocks. In human, these two distinct regions harbor the genes *IGHV3-30* (and related paralogs) and *IGHV1-58/IGHV4-59*, respectively (**Fig.IG.S2.ce**); the *IGHV3-30* region is known to be highly diverse with respect to structural variation in the human population[107]. Of note, these regions exhibited extensive expansion and contraction between ape species (**Fig.IG.S2.ce**).

In addition to tandem duplication expansions and contractions, we also noted large structure variants, particularly in the IG loci, including insertions and inversions between species and haplotypes. Comparison of two haplotypes of the bonobo IGL locus revealed a 1.4 Mbp inversion (denoted as INV) flanked by a mosaic tandem repeat consisting of directed and inverted units of two types denoted as A (~58 kbp) and B (~27 kbp) with variable counts of repeat copies (**Fig. IG.S2.d**). The maternal haplotype can be represented as $A_1B_1A_2B_2A_2$ + INV + $A_3$’$B_3$’$A_4$’ and the paternal haplotype can be represented as $A_5B_4A_5$ + INV’ + $A_6$’, where ‘ refers to the inverted orientation of the block. These observations raised a concern about the assembly accuracy as the complex repeat structure marks hotspots for possible structural assembly errors. To verify the assembly quality, HiFi reads were assembled using the LJA genome assembler based on construction of highly accurate de Bruijn graphs[108]. Analysis of the de Bruijn graph revealed that the flanking repeat copies A and B from distinct haplotypes sufficiently diverged from each other to make assembly error practically impossible. Contigs generated by LJA also supported the presence of the inversion within the bonobo IGL locus.

As expected, these large SVs were associated with the presence of ape-specific genes (defined in Methods), including examples of IGHV genes residing within tandem SD/repeat expansions and contractions (**Fig 3A, Fig IG.S1.e**). Across all species, the greatest number of species-specific genes were observed in IGH (**Fig. IG.S2.f**), which positively correlated with a greater density of long repeats (≥10 kbp) in IGH relative to the other five loci ($r=0.51$, $P=6.95\times10^{-5}$; **Fig. IG.S2.g**). In addition, within species, the IG loci were characterized by higher V gene distances between haplotypes ($P=0.013$, Kruskal-Wallis test) and lower fractions of identical V gene sequences between haplotypes ($P=3.03\times10^{-13}$, Kruskal-Wallis test) compared to TR loci (**Fig. IG.S2.hi**). Together these data suggested that the IG loci undergo more rapid divergence than the TR loci, potentially due to evolutionary and functional constraints placed on TR loci by required interactions with MHC. While these haplotypes provide clear evidence for rapid divergence in these critical immune loci, it is imperative to note that further sampling of haplotypes for these species will be necessary for characterizing additional SVs and more complete sets of IG/TR genes and alleles for each of these species. Even in humans, IG and TR genes continue to be discovered[109].

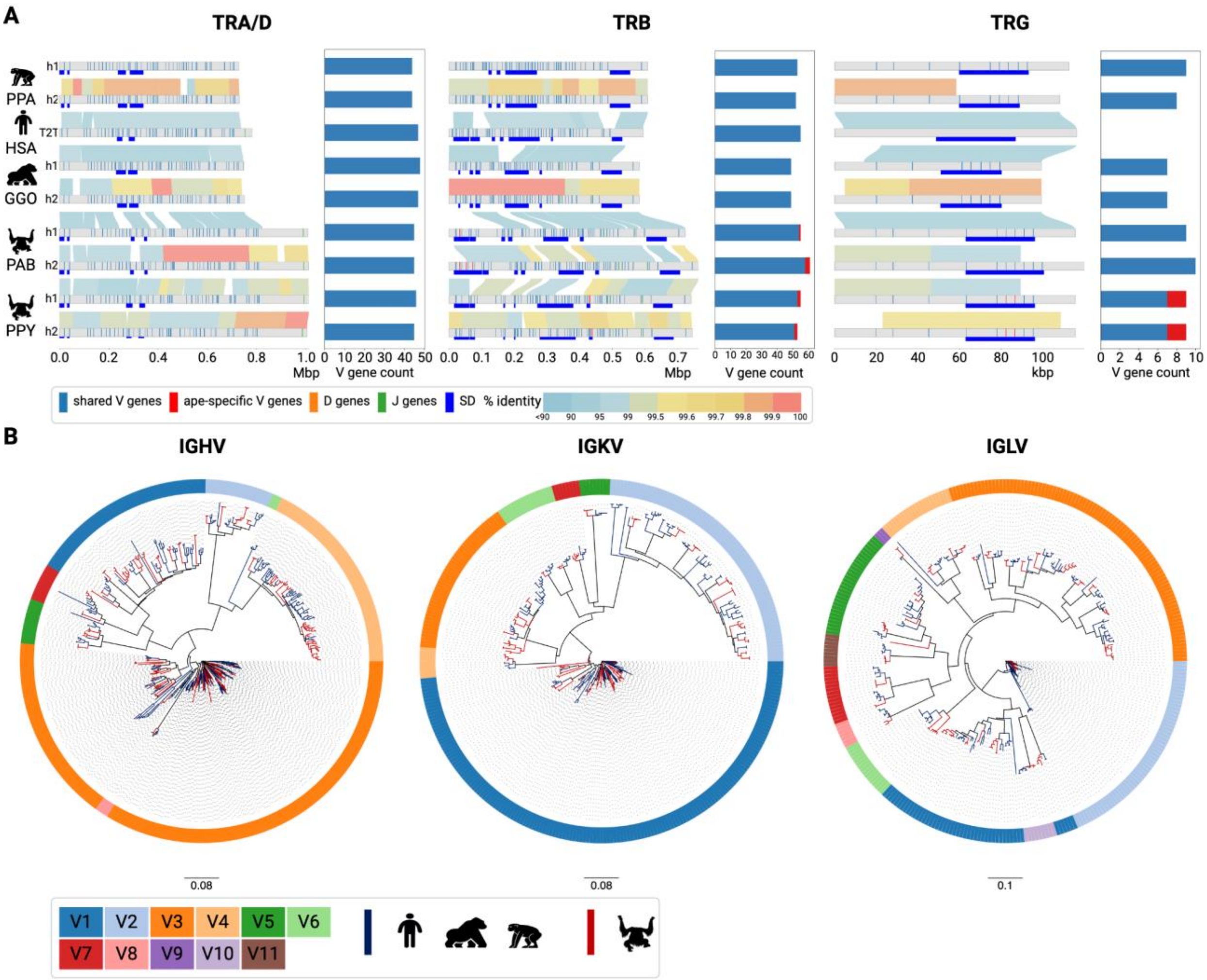


**Figure IG.S1. Comparative analysis of IG/TR loci across four great ape species.**
**(A)** Annotated haplotypes of TRA/D, TRB, and TRG loci. The legend is consistent with **Fig. 3a**.
**(B)** Phylogenetic trees of IGHV, IGKV, and IGLV genes collected across four ape species (PPA, GGO, PAB, PPY) and combined with known human genes. The outer circle in each gene tree shows gene subfamilies. The branches corresponding to individual genes were colored in dark blue if they correspond to the human, gorilla, or bonobo and red if they correspond to orangutan species.

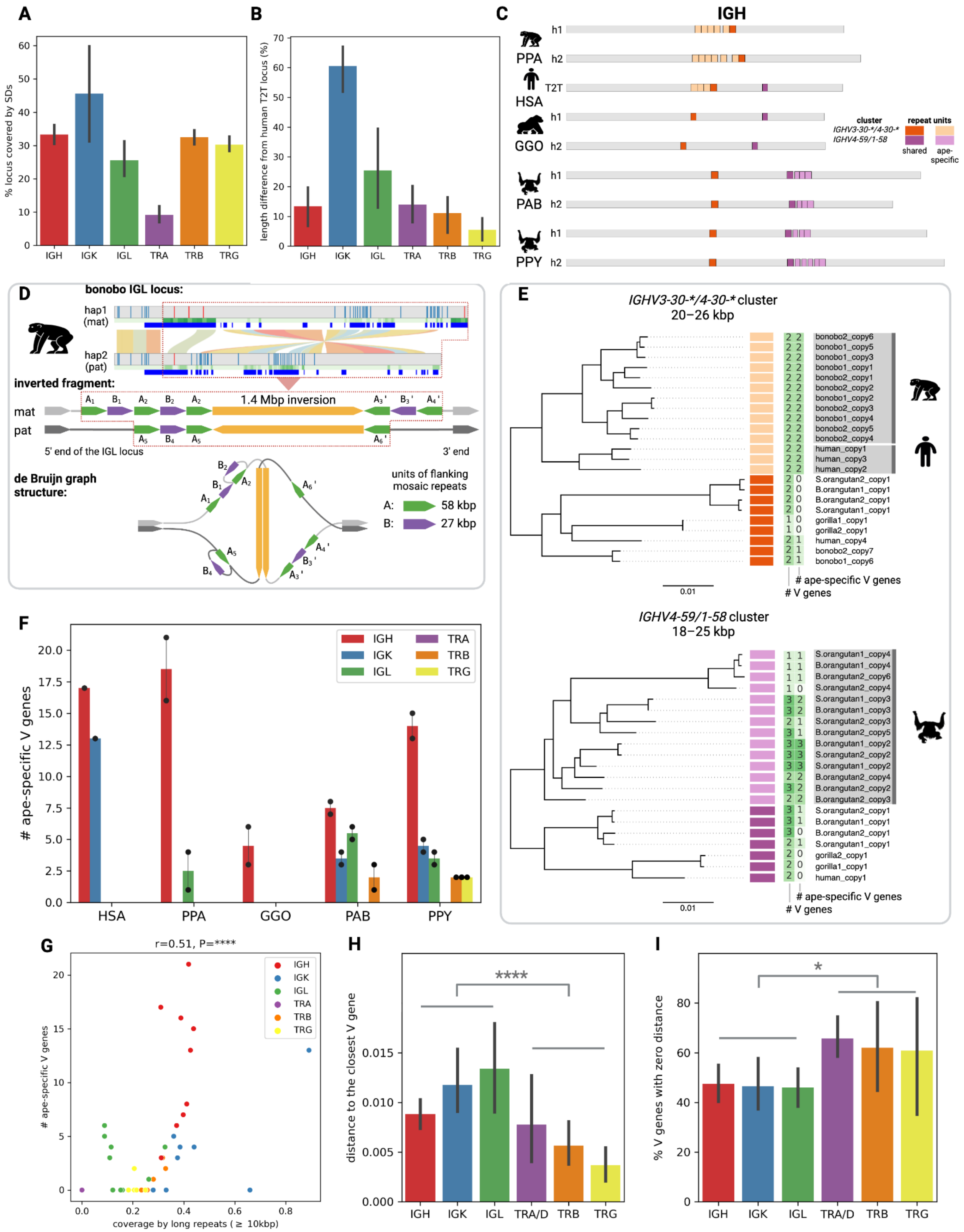


**Figure IG.S2. The diversity analysis of IG/TR loci and genes across five great ape species (PPA, GGO, PAB, PPY, HSA). (A)** Percentage of the IG/TR locus lengths covered by SD

blocks computed between haplotypes of the same species and collected across five ape species. **(B)** Length differences (%) computed for all ape IG/TR loci with respect to the corresponding human T2T locus. **(C)** A diagram showing positions of two tandem repeat units containing *IGHV3-30-*/4-30-** genes (orange) and *IGHV4-59/1-58* genes (purple) in IGH loci of five ape species. Shared and ape-specific units are colored in dark and pale colors, respectively.
**(D)** Genomic analysis of the inversion structure in the bonobo IGL locus. The top part shows an alignment of two haplotypes of the bonobo IGL locus with the inverted fragment shown as a red dashed rectangle. The middle part shows the genomic structure of the inverted fragment that includes a 1.4 Mbp inversion (shown in yellow) and mosaic repeats flanking it (shown in green and purple). The bottom part shows the simplified structure of the de Bruijn graph corresponding to the inverted fragment. For visualization purposes, bulges representing divergence between haplomes were not shown in the graph. Lengths of genomic blocks and de Bruijn graph edges are not up to scale. **(E)** Phylogenetic trees of units of the *IGHV3-30-*/4-30-** and *IGHV4-59/1-58* clusters. Colors of shared and ape-specific units are consistent with panel B. Counts of all V genes and ape-specific V genes in each repeat unit are shown in green. Subtrees corresponding to bonobo (the left tree), the human (left), and both orangutan species (right) are highlighted in gray. **(F)** Counts of ape-specific V genes across IG/TR loci and five great ape species.
**(G)** Counts of ape-specific V genes vs. repetitiveness of the corresponding locus across IG/TR loci and five great ape species. **(H)** The distances between closest pairs of V genes from different haplotypes within the same locus and species collected across IG/TR loci and four ape species (PPA, GGO, PAB, PPY). The distance is computed as the fraction of nonmatching positions in the alignment. **(I)** The fractions of V genes from different haplotypes within the same locus and species with identical gene sequences collected across IG/TR loci and four ape species (PPA, GGO, PAB, PPY).

### XI. MHC I and MHC II analyses

Contributing authors:

Joanna Malukiewicz, Britta S. Meyer, Mihir Trivedi, Prajna Hebbar, Tobias L. Lenz

##### Methods

We defined the MHC genomic region as all loci located between *GABBR1* and *KIFC1*, as previously determined by Shiina et al.[110] This region is located on chromosome six in humans[110]. We first identified the chromosomal location of the MHC genomic region in the ape T2T assemblies of bonobo (*Pan paniscus*; NCBI assembly NHGRI_mPanPan1-v2.0_pri), chimpanzee (*Pan troglodytes*; NCBI assembly NHGRI_mPanTro3-v2.0_pri), western lowland gorilla (*Gorilla gorilla gorilla*; NCBI assembly NHGRI_mGorGor1-v2.0_pr), Bornean orangutan (*Pongo pygmaeus*; NCBI assembly NHGRI_mPonPyg2-v2.0_pri), Sumatran orangutan (*Pongo abelii*; NCBI assembly NHGRI_mPonAbe1-v2.0_pri), and siamang (*Symphalangus syndactylus*; NCBI assembly NHGRI_mSymSyn1-v2.0_pri). The chromosomal locations of each ape MHC genomic region were located by first aligning each respective haplotype of each T2T ape assembly to the human T2T assembly (NCBI assembly T2T-CHM13v2.0) as a reference with minimap2 v2.28[27]. SAMtools[111] was then used to filter resulting SAM files for mapped reads with the "-F4" flag with the view subcommand. Then filtered SAM files were sorted with SAMtools sort subcommand and simultaneously converted to BAM file format. BAM files were indexed with SAMtools index subcommand. Finally, the SAMtools view subcommand was used to subset BAM files from the regions of each ape assembly haplotype that mapped to human chromosome 6 between coordinates 2937649-3331258. These coordinates flank the MHC region of human T2T assembly T2T-CHM13v2.0.

To annotate putative classical and nonclassical MHC class I and class II genes within the two individual haplotypes of each ape T2T genomic assembly, we used two complementary approaches. First, we used EXONERATE 2.4[112] with the "est2genome" mapping model (https://www.ebi.ac.uk/about/vertebrate-genomics/software/exonerate). EXONERATE was run recursively with functional human HLA gene and CDS annotations from T2T-CHM13v2.0 and the IPD-MHC database (Release 3.12.0.0 build 211; https://www.ebi.ac.uk/ipd/mhc/) gene and CDS annotations for chimpanzee, bonobo, western lowland gorilla, and both orangutan species. For all ape species except siamang, chromosome 5 was the query and for siamang chromosome 23 served as the query and EXONERATE results were filtered to matches of greater than 95%. Second, we mapped human HLA gene annotations to chromosome 23 of siamang and chromosome 5 for all remaining ape species with MINIMAP2. Results between EXONERATE and minimap2 were compared for concordance. Then annotations were manually verified and curated with ALIVIEW 2.8[113] to retain only a single gene annotation per locus, and all gene annotations were individually compared against all available human and ape species orthologs to confirm proper annotation. Due to the absence of previous gene and CDS annotations for the siamang, we assigned MHC gene annotations for presumed start codon to stop codon based on corresponding human HLA gene and CDS annotations. MHC class I and class II gene names were assigned according to human and ape species orthologs.

Phylogenetic trees were produced for MHC class I loci and MHC class II DRB loci to confirm the identity of MHC genes exhibiting copy number variation. First, separate MHC class I and MHC-DRB multiple sequence alignments were produced with the online version of MAFFT (https://mafft.cbrc.jp/alignment/server/index.html). Both alignments consisted of genomic sequence of annotated MHC genes from ape T2T assemblies, human coding sequence (CDS) and genomic sequence of genes from T2T-CHM13v2.0. Genomic and coding sequences for MHC class II DRB genes (n=71) and MHC class I loci (n=406) were obtained from the IPD-MHC database (https://www.ebi.ac.uk/ipd/mhc/, Release 3.12.0.0 (2024-01) build 211). Multiple sequence alignments were created using MAFFT[114], and intronic sequence was removed from final alignments.

To generate phylogenetic trees from coding sequence from all MHC class I and the MHC class II DRB genes, respectively, the IQ-TREE platform (http://iqtree.cibiv.univie.ac.at) was used with default settings[115]. For the MHC class II DRB genes, a codon partition file was provided to enhance the accuracy of the analysis. IQ-TREE automatically selected the best-fitting substitution models and calculated node support for the phylogenetic trees with Ultrafast Bootstrap[116]. Phylogenetic trees were plotted using RStudio (version 2024.04.2+764) with the ggtree[117] and ape[118] packages, retaining bootstrap values above 70.

Dot plots were used to highlight structural variations between the haplotypes of each primate species. Two haplotypes from each species were aligned using NUCMER and show-coords from the MUMMER package v4.0.0rc1[119] to generate delta and coordinates files from the FASTA sequences. These coordinates, along with BED file annotations for coding genes and pseudogenes within the MHC regions, were used to generate and customize the dot plots with the SVbyEye package (https://github.com/daewoooo/SVbyEye) in R, along with ggplot2[120].

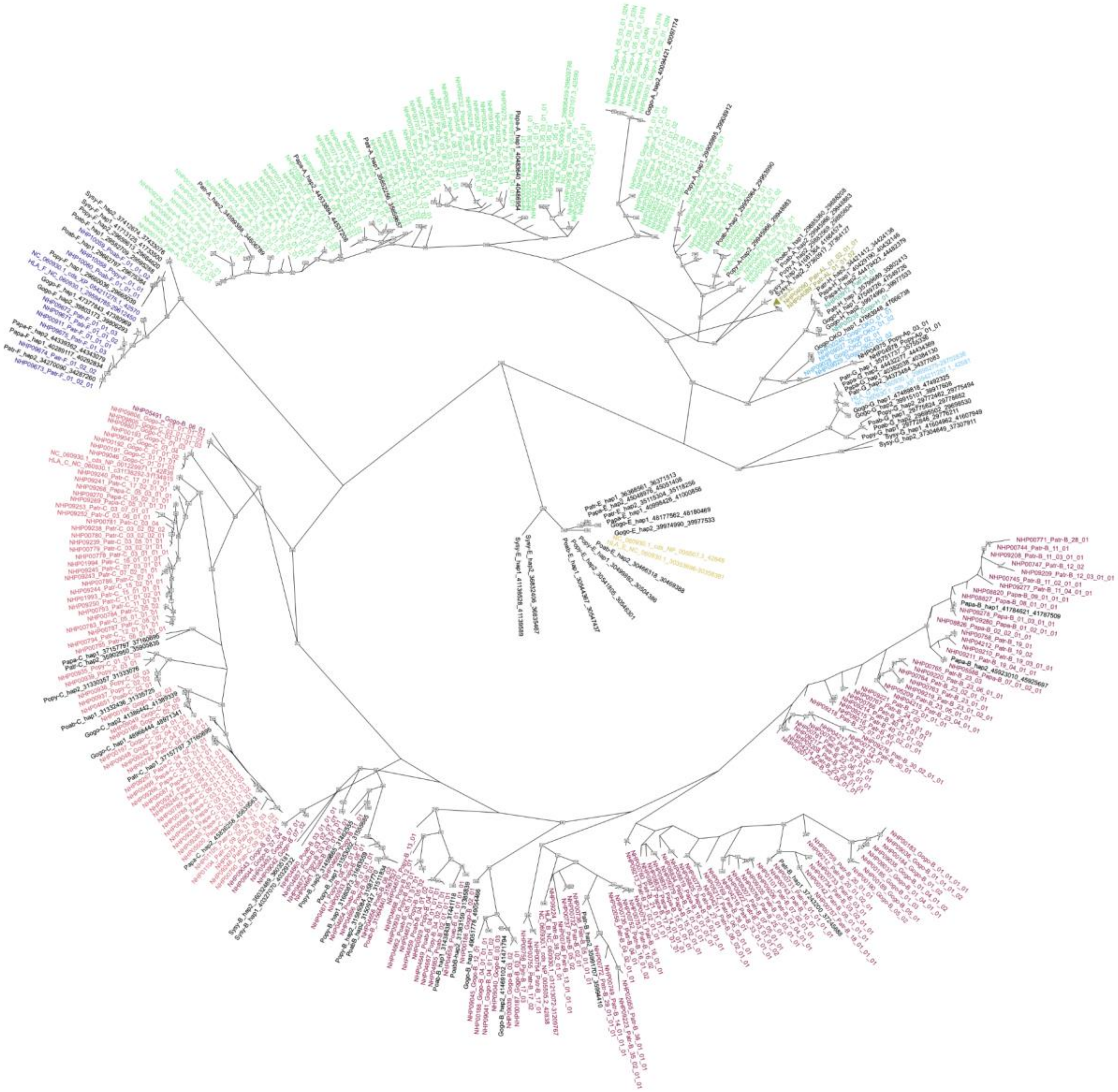

**Figure MHC.S1. Unrooted maximum likelihood phylogeny of MHC class I coding genes.** Phylogeny of coding sequence of human ape MHC class I loci from the six T2T assemblies, genomic sequence of species MHC class I genes obtained from the IMGT MHC database, and annotated MHC class I genes from the human T2T-CHM13v2.0 genomic assembly. NHP gene names are abbreviated according to species (Patr-*Pan troglodytes*, Papa-Pan paniscus, Gogo-*Gorilla gorilla*, Popy-*Pongo pygmaeus*, Poab- *Pongo abelii*, Sysy-*Symphalangus syndactylus*). Clusters of orthologous loci are represented by unique colors that match those shown in **Fig. 3b** for a given MHC I coding gene or pseudogene. Newly annotated MHC I genes from this study are shown in black. IQ-TREE chose GTR+F+I+G4 as the best fitting model for this tree following the BIC criterion.

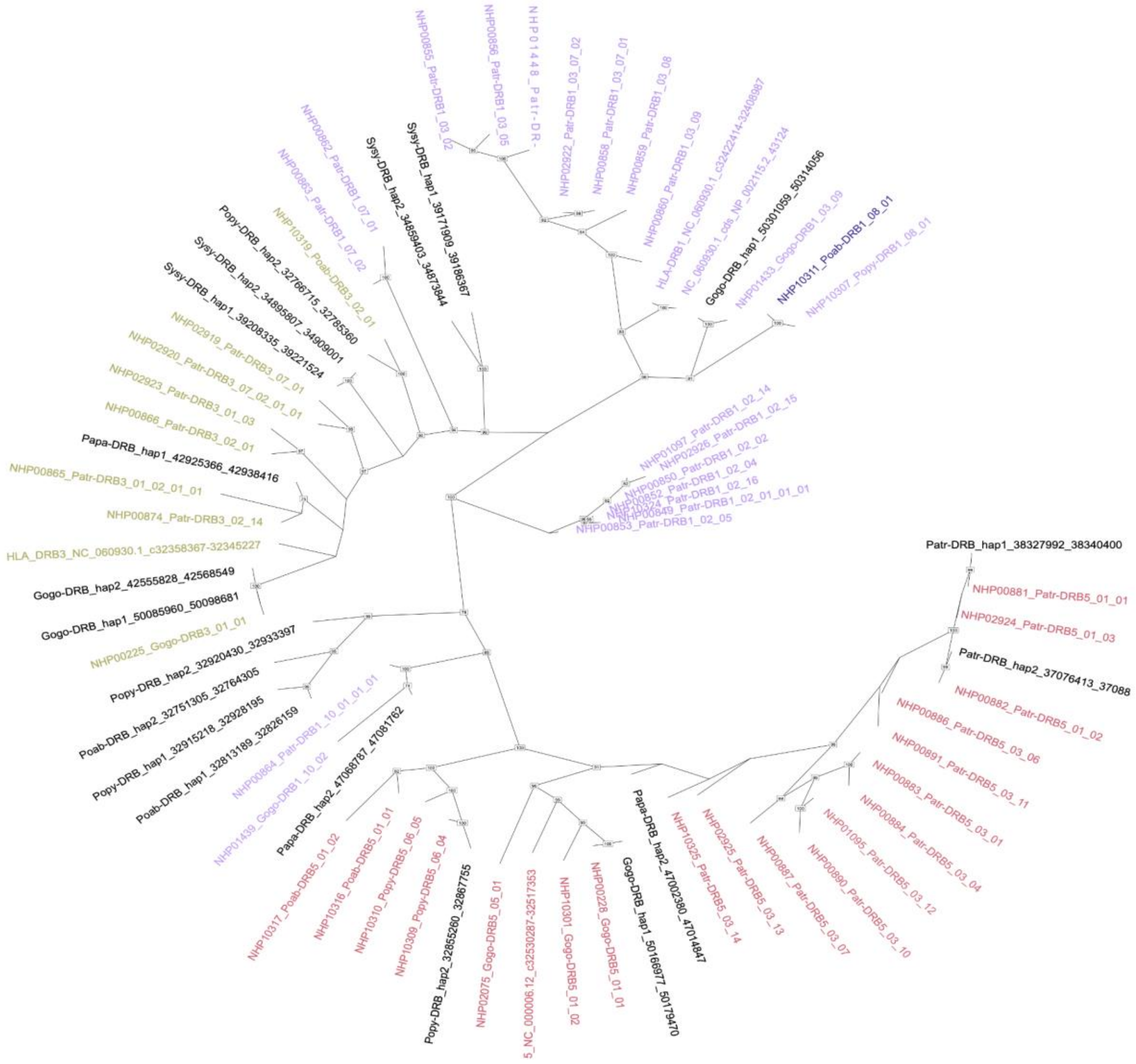


**Figure MHC.S2. Unrooted maximum likelihood phylogeny of MHC class II DRB coding genes.** Phylogeny of coding sequence of human and ape MHC class II DRB loci from the six T2T assemblies, genomic sequence of species MHC class II DRB genes obtained from the IMGT MHC database, and annotated MHC class II genes from the human T2T-CHM13v2.0 genomic assembly. NHP gene names are abbreviated according to species (Patr-*Pan troglodytes*, Papa-*Pan paniscus*, Gogo-*Gorilla gorilla*, Popy-*Pongo pygmaeus*, Poab-*Pongo abelii*, Sysy-*Symphalangus syndactylus*). Clusters of orthologous loci are represented by unique colors that match those shown in **Fig. 3c** for a given MHC II DRB coding gene. Newly annotated MHC II genes from this study are shown in black. IQ-TREE chose K2P+I+G4 as the best fitting model for codons 1 and 2, the HKY+F+I+G4 model for codon 3 following the BIC criterion.

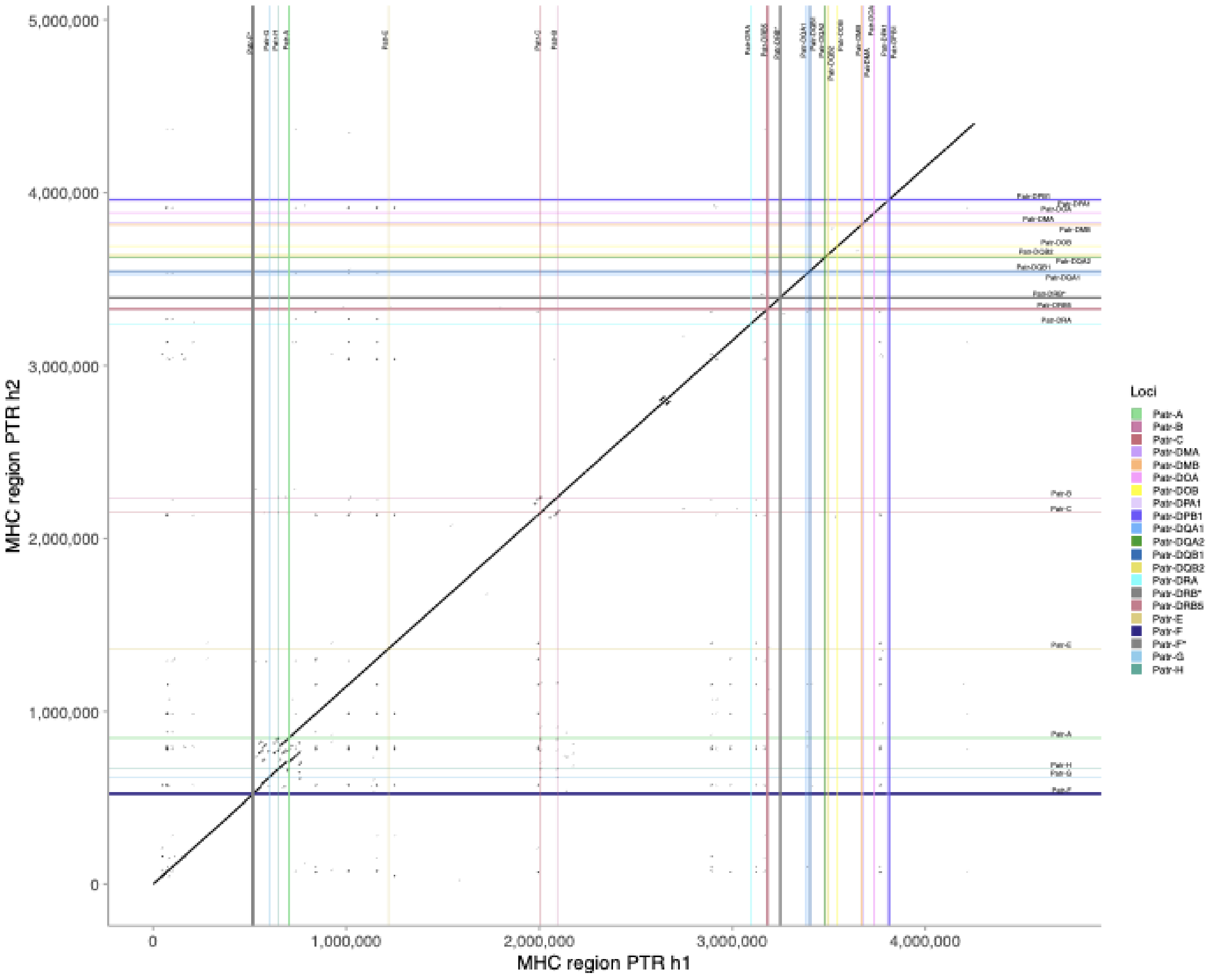


**Figure MHC.S3. Dot plot of PTR.h1/h2.** Dot plots were made for the two MHC region haplotypes of *Pan troglodytes*. Locations of MHC coding genes and pseudogenes (labeled with *) for haplotypes 1 and 2 are labeled on the dot plot and also represented by horizontal lines (haplotype 1) and vertical (haplotype 2) lines. Unique colors representing each locus match those shown in **Fig. 3b-c** for a given MHC locus. The span of each horizontal and vertical line represents the length of a given gene.

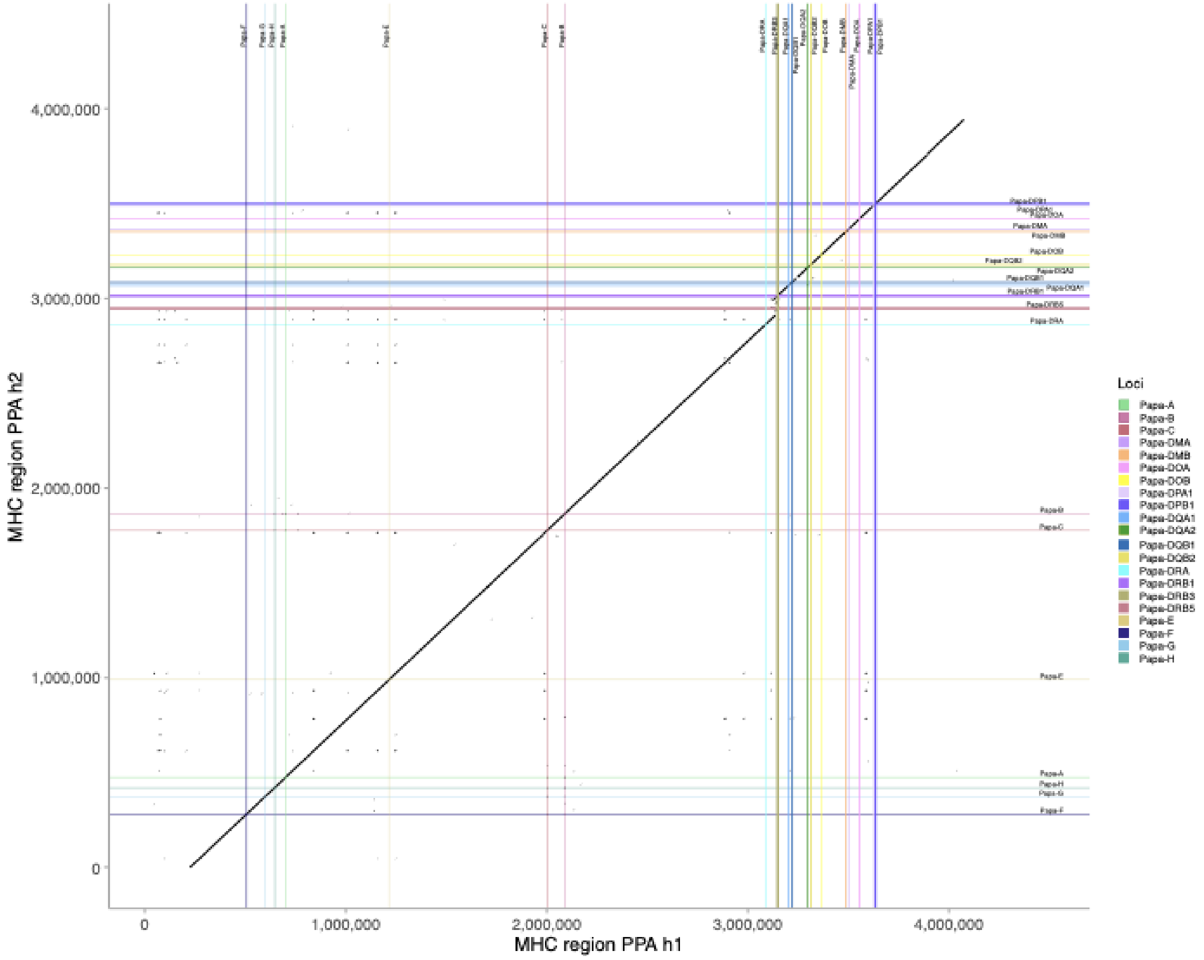


**Figure MHC.S4. Dot plot of PPA.h1/h2.** Dot plots were made for the two MHC region haplotypes of *Pan paniscus*. Locations of MHC coding genes and pseudogenes (labeled with *) for haplotypes 1 and 2 are labeled on the dot plot and also represented by horizontal lines (haplotype 1) and vertical (haplotype 2) lines. Unique colors representing each locus match those shown in **Fig. 3b-c** for a given MHC locus. The span of each horizontal and vertical line represents the length of a given gene.

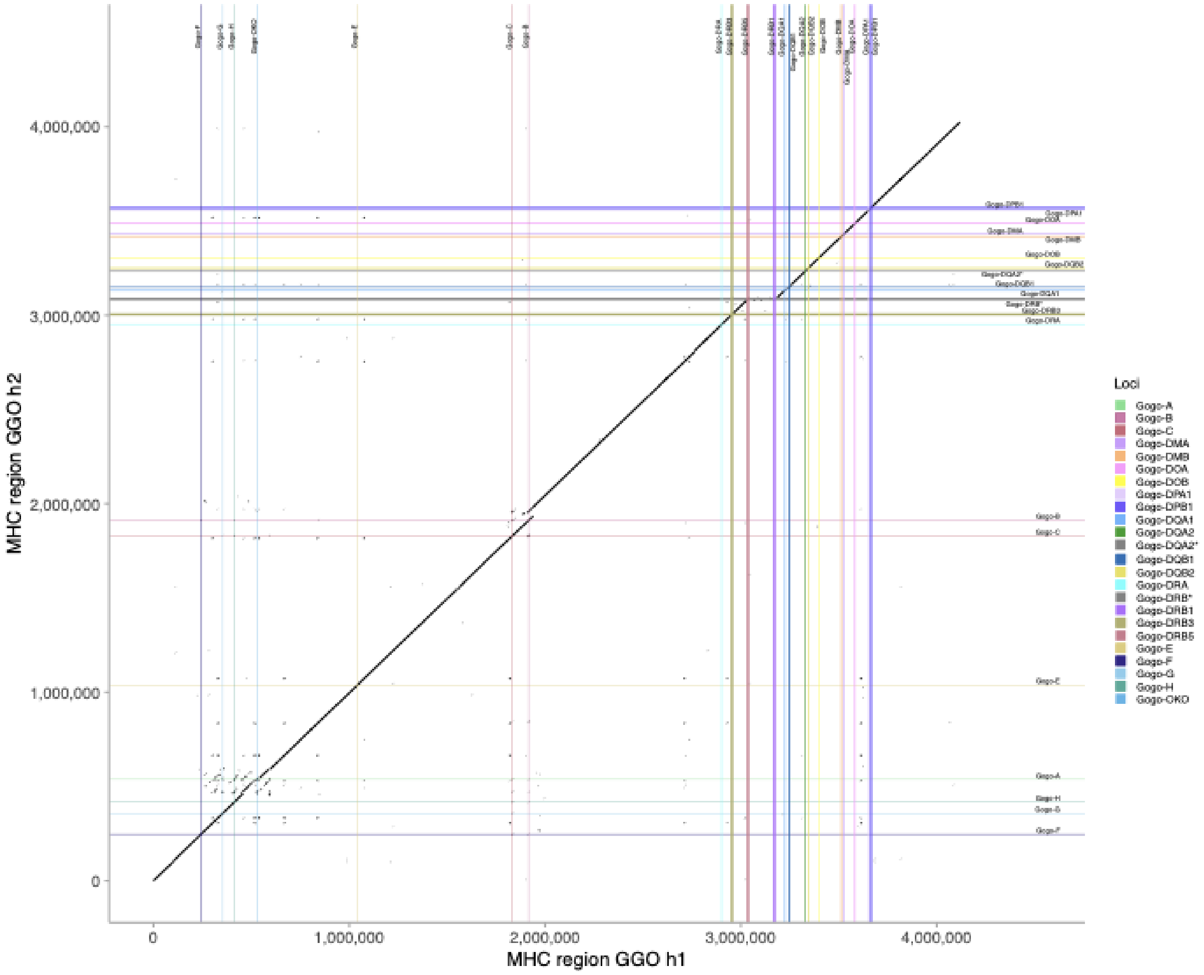


**Figure MHC.S5. Dot plot of GGO.h1/h2.** Dot plots were made for the two MHC region haplotypes of *Gorilla gorilla*. Locations of MHC coding genes and pseudogenes (labeled with *) for haplotypes 1 and 2 are labeled on the dot plot and also represented by horizontal lines (haplotype 1) and vertical (haplotype 2) lines. Unique colors representing each locus match those shown in **Fig. 3b-c** for a given MHC locus. The span of each horizontal and vertical line represents the length of a given gene.

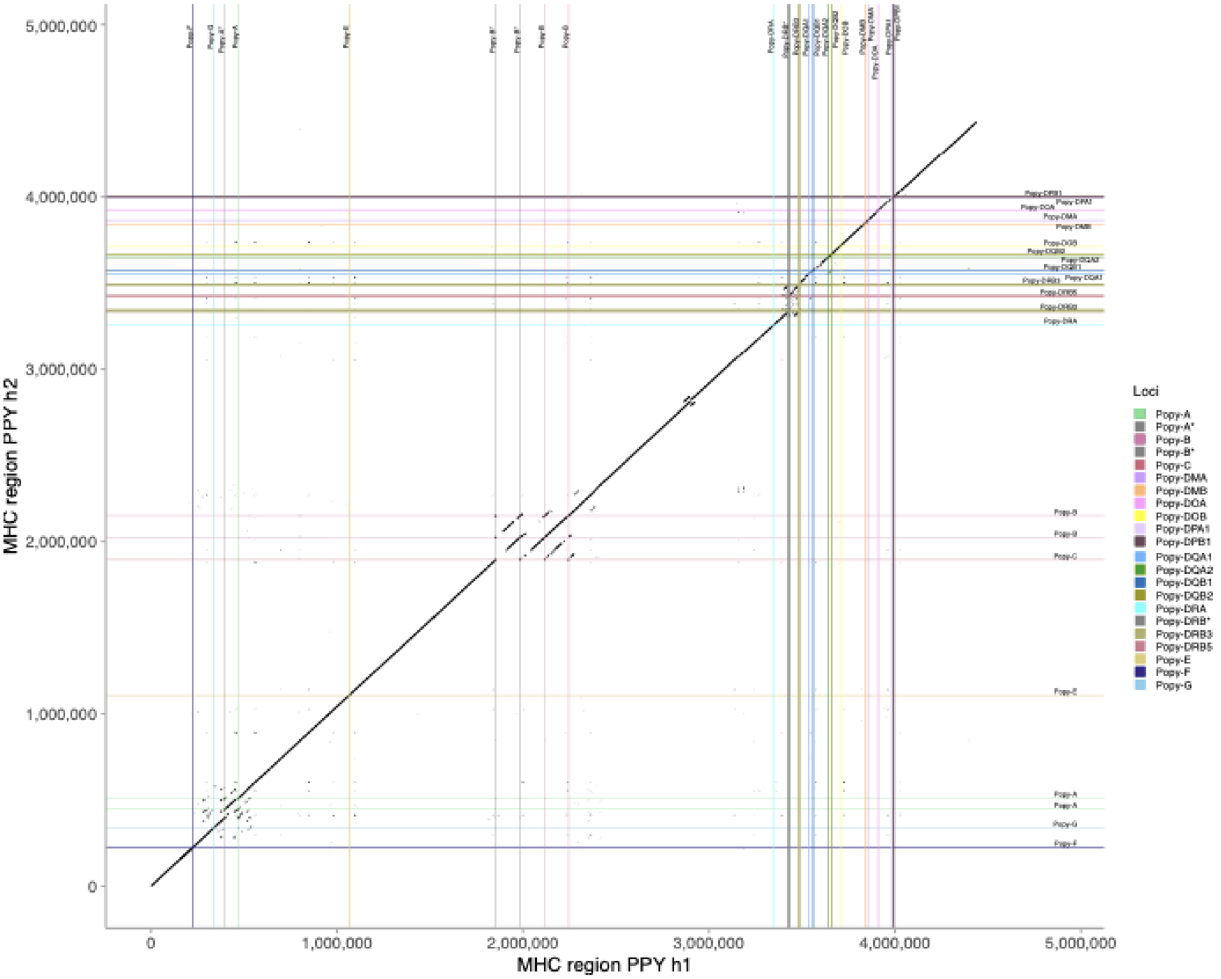


**Figure MHC.S6. Dot plot of PPY.h1/h2.** Dot plots were made for the two MHC region haplotypes of *Pongo pygmaeus*. Locations of MHC coding genes and pseudogenes (labeled with *) for haplotypes 1 and 2 are labeled on the dot plot and also represented by horizontal lines (haplotype 1) and vertical (haplotype 2) lines. Unique colors representing each locus match those shown in **Fig. 3b-c** for a given MHC locus. The span of each horizontal and vertical line represents the length of a given gene.

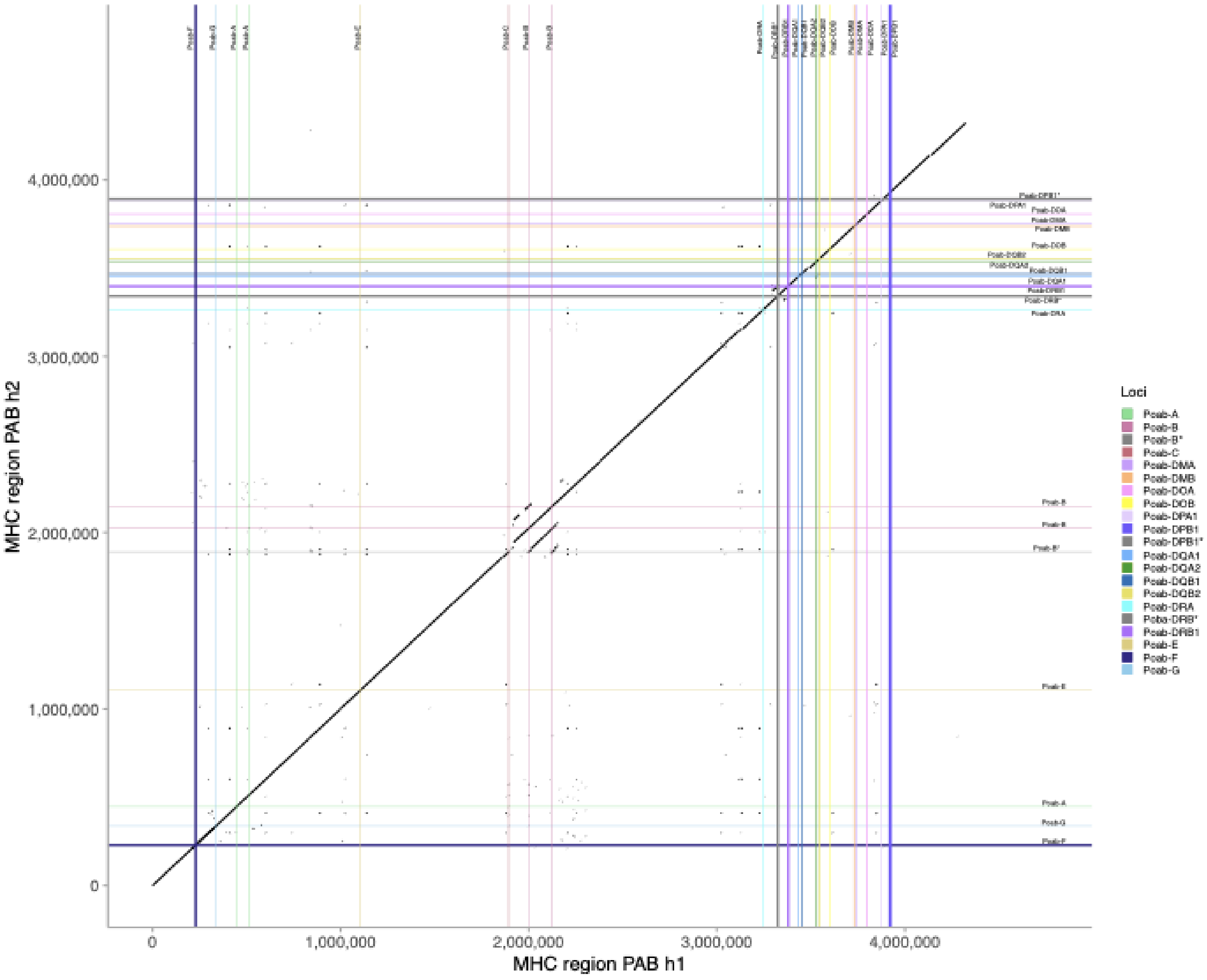


**Figure MHC.S7. Dot plot of PAB.h1/h2.** Dot plots were made for the two MHC region haplotypes of *Pongo abelii*. Locations of MHC coding genes and pseudogenes (labeled with *) for haplotypes 1 and 2 are labeled on the dot plot and also represented by horizontal lines (haplotype 1) and vertical (haplotype 2) lines. Unique colors representing each locus match those shown in **Fig. 3b-c** for a given MHC locus. The span of each horizontal and vertical line represents the length of a given gene.

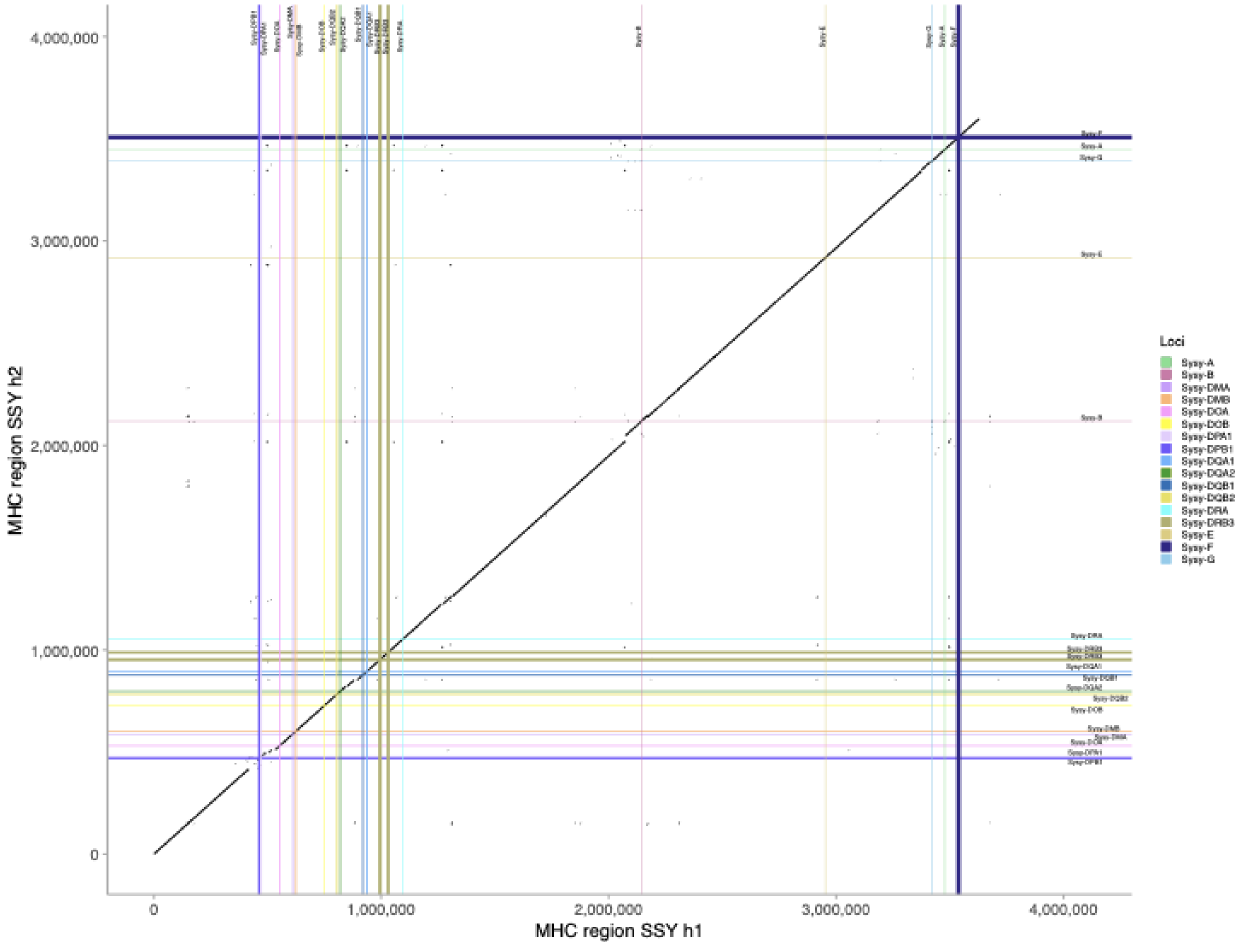


**Figure MHC.S8. Dot plot of SSY.h1/2.** Dot plots were made for the two MHC region haplotypes of *Symphalangus syndactylus*. Locations of MHC coding genes and pseudogenes (labeled with *) for haplotypes 1 and 2 are labeled on the dot plot and also represented by horizontal lines (haplotype 1) and vertical (haplotype 2) lines. Unique colors representing each locus match those shown in **Fig. 3b-c** for a given MHC locus. The span of each horizontal and vertical line represents the length of a given gene.

**Table MHC.S1. Manual annotation results of ape MHC class I coding genes and pseudogenes.** (See accompanying Excel file for table.) This table shows locations of annotated MHC class I coding and pseudogenes on haplotype 1 and 2 of each ape T2T assembly. The “MHC Class” column indicates these results are for MHC class I genes, the “Species” column indicates the common and scientific names of specific species, the “Assembly Name” column indicates names of specific genomic assemblies, the “Locus Name” column indicates specific MHC class I loci, the “Locus Biotype” column indicates whether the locus is a coding gene or pseudogene, the “Haplotype” column indicates the first or second haplotype in question, the

“Putative Gene Start Position” column indicates the putative start codon of each MHC class I locus, the “Putative Gene End Position” column indicates the putative end codon of each MHC class I locus, and the “Direction of Transcription” column indicates whether the is transcribed from the positive or negative DNA strand.

**Table MHC.S2. Manual annotation results of ape MHC class II coding genes and pseudogenes.** (See accompanying Excel file for table.) This table shows locations of annotated MHC class II coding and pseudogenes on haplotype 1 and 2 of each ape T2T assembly. The “MHC Class” column indicates these results are for MHC class II genes, the “Species” column indicates the common and scientific names of specific species, the “Assembly Name” column indicates names of specific genomic assemblies, the “Locus Name” column indicates specific MHC class II loci, the “Locus Biotype” column indicates whether the locus is a coding gene or pseudogene, the “Haplotype” column indicates the first or second haplotype in question, the “Putative Gene Start Position” column indicates the putative start codon of each MHC class II locus, the “Putative Gene End Position” column indicates the putative end codon of each MHC class II locus, and the “Direction of Transcription” column indicates whether the is transcribed from the positive or negative DNA strand.

**Phylogenetic tree reconstruction of MHC region**

We created 500 bp nonoverlapping bins from the 5.5 Mbp MHC locus from the human T2T sequence. These human 500 bp bins were pairwise aligned to each haplotype from the sequenced ape species using minimap2[27] to extract the homologous region from all the haplotypes. We also used the HG002 human sequence and did not include siamang in this analysis. We extracted homologous sequences in 500 bp increments and optimized local multiple sequence alignments (8259 bins for 500 bp) using MAFFT and then concatenated to generate 5 kbp regions (953 bins). Maximum likelihood phylogenetic trees were constructed with IQ-TREE2[121] applying an optimized base substitution model and 1000 bootstrap replicates. To compare tree topologies we applied the Robinson-Foulds method, as implemented in the python ete3 package (compare() function)[122]. Each haplotype from the diploid assembly was considered separately, by selectively removing the external nodes pertaining to another haplotype in a tree. We performed hierarchical clustering taking trees and their pairwise distances into account to identify distinct topologies, through hclust and cuttree functions in R.

Following are the 19 topologies constructed during the analysis:

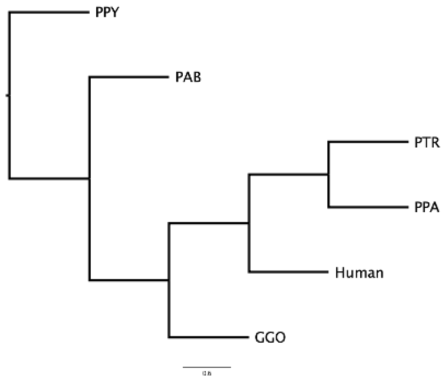


1

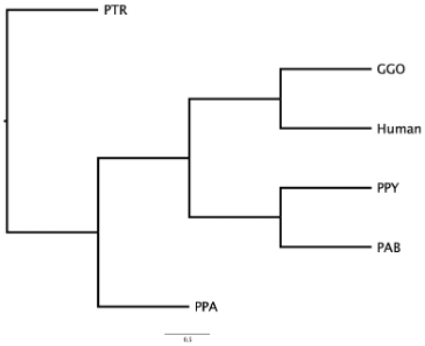


2

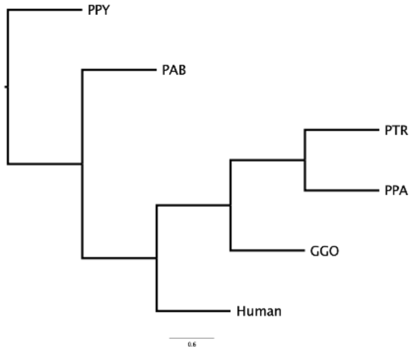


3

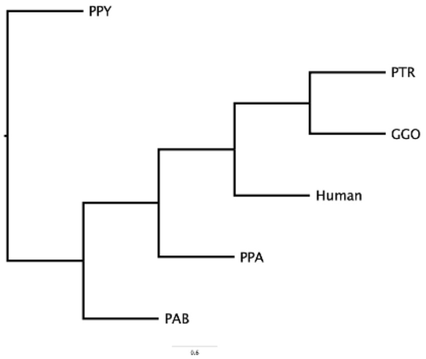


4

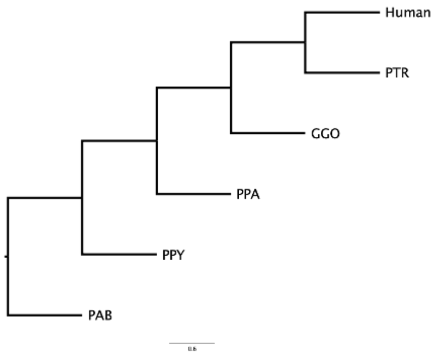


5

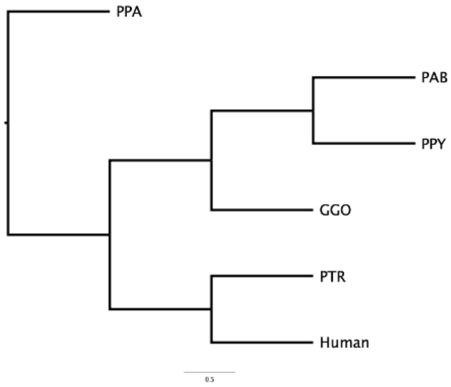


6

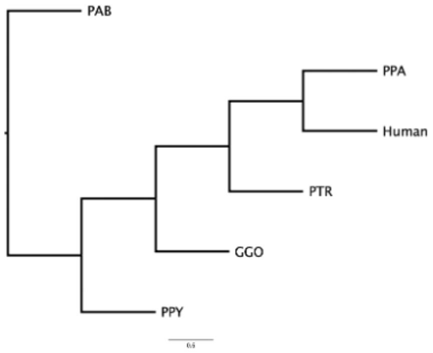


7

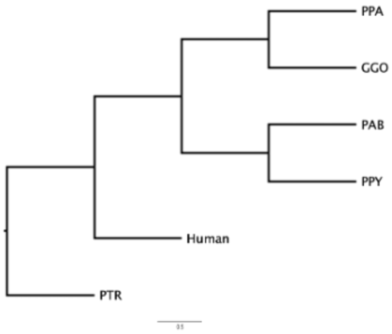


8

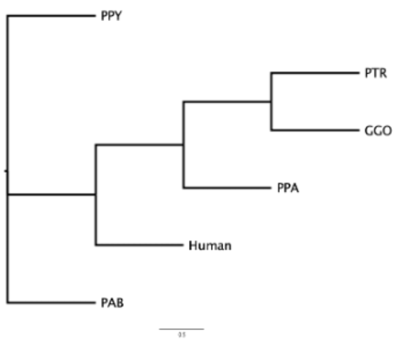


9

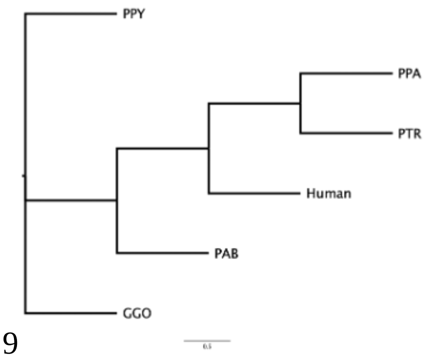


10

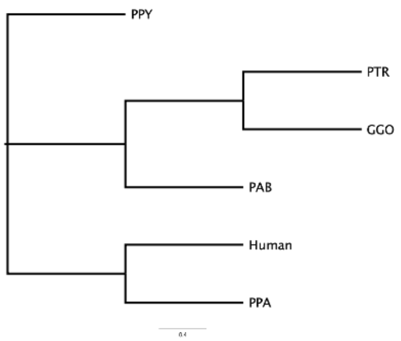


11

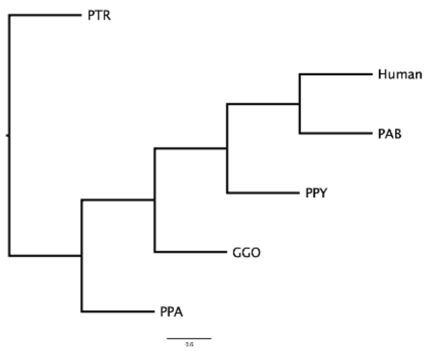


12

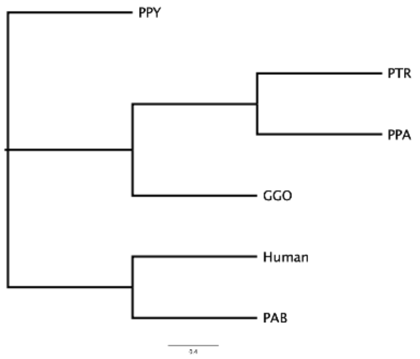


13

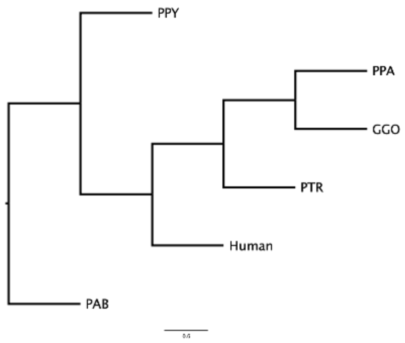


14

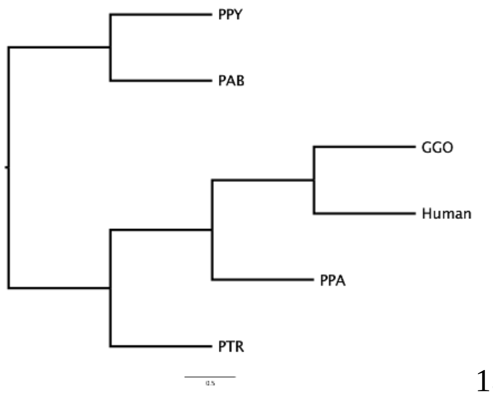


15

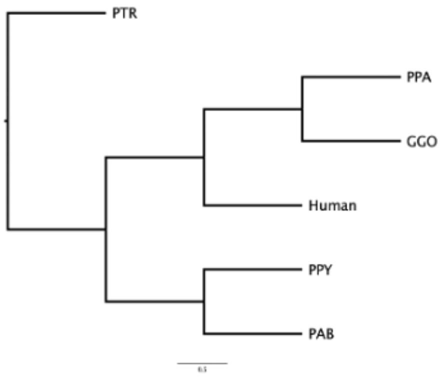


16

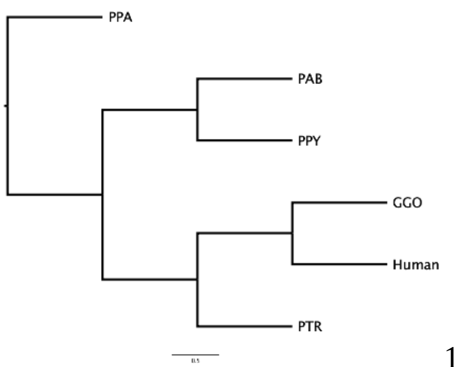


17

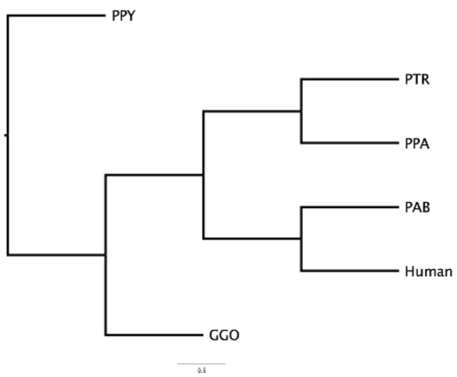


18

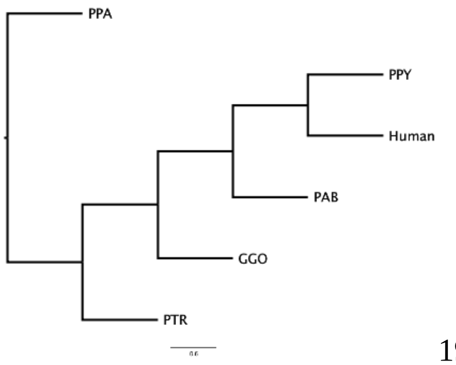


19

**Summary of results**

We also completely assembled and annotated the 4-5 Mbp region corresponding to 12 ape haplotypes for the MHC loci—a highly polymorphic region among mammals[123] encoding diverse cell surface proteins crucial for antigen presentation, adaptive immunity[124], and in humans disproportionately implicated in disease via genome-wide association[125]. Comparative sequence analyses confirm extraordinary sequence divergence and structural variation, including expansions and contractions associated with MHC genes (**Fig. 3b-d**). Overall, MHC class I genes reveal more extensive structural variation within and among the apes than MHC class II genes (**Fig. 3b,c**, **Fig. MHC.S1,2**). For instance, siamang carries a distinct Sysy-B locus but lacks a distinct Sysy-C locus (**Fig. 3b,c**). Furthermore, an inversion occurred between the MHC-G and MHC-A loci after the divergence of the great apes and humans from the siamang. The MHC I gene content and organization is identical across human, bonobo, and chimpanzee, but we observe relatively high levels of interspecific and intraspecific variability in the other species, including additional genes (e.g., Gogo-OKO (**Fig. 3b-d**), which is related but distinct from Gogo-A[123]). Furthermore, in orangutans, we observed expansion of MHC-A and MHC-B in both the Bornean and Sumatran lineages (**Fig 3d**). Unlike human, chimpanzee, gorilla, and bonobo, MHC-B is duplicated in both haplotypes of the two orangutan species. Both the Sumatran and

Bornean orangutan species possessed a duplicated MHC-A locus on one haplotype and a single MHC-A locus on the other haplotype. Human and the other great ape species possessed a single MHC-A locus on both haplotypes. Surprisingly, both orangutan species lack the MHC-C locus on one haplotype and have the MHC-C locus on the other haplotype, thus revealing another case of copy number variation of an MHC class I gene in these species. All ape and human species possessed the same identical set of MHC II loci, but there was copy number variation at the interspecific and intra-individual level in the DRB locus among all studied species (**Fig. 3b,c**). We also observed two cases where an MHC locus was present as a functional gene on one haplotype and as a pseudogene on the other haplotype, as exemplified by the Gogo-DQA2 locus in gorilla and the Poab-DPB locus in Sumatran orangutan.

### XII. Methylation

Contributing authors:

Dongmin R. Son, Yong Hwee Eddie Loh, Soojin V. Yi

#### Methods

##### DNA methylation analysis

To generate CpG methylation calls, ONT data were base-called using Guppy v6.3.7 with the model "dna_r9.4.1_450bps_modbases_5hmc_5mc_cg_sup_prom.cfg". Counts of modified bases at each cytosine position were generated using modbam2bed v0.10.0 after Minimap2 v2.26 mapping to the genome. Given that 5mC in the CpG context is symmetrical, read counts of two cytosines in the CpG context were combined to represent a single CpG site. The fractional methylation level at each CpG site was calculated for CpG sites, which had at least five read counts.

To generate a list of orthologous promoters (**Table MET.S1-2**), we first defined human promoter regions as 1000 bp upstream to 500 bp downstream of the transcription start site of protein-coding genes as annotated by the CuratedRefSeqv5.1 annotation track on the human HG002v1.0.1 paternal genome. We used halLliftOver against a 16-way Cactus alignment to identify orthologous promoters in NHP primary genomes. Additionally, we restricted our analysis to genes that are detected as single-copy and intact according to TOGA (where the middle 80% of the CDS was present and exhibited no gene-inactivating mutations) based on the TOGA hg38 gene annotations of NHPs. We calculated the distance between the center of orthologous promoters and the 5' end of genes using the RefSeq annotation track for each NHP, filtering out genes with distances over 2.5 kbp. Out of these 8,256 confident orthologous protein-coding promoters between humans and six NHPs, 8,177 promoters had at least six CpGs, and 8,174 also had available expression levels. We calculated the mean 5mC values per promoter. We normalized these values within each species using the z-score transformation to scale between 0 and 1, using scikit-learn MinMaxScaler{Pedregosa, 2011 #299}. Promoters were defined as consistently methylated if the maximum difference in 5mC levels between species was less than 0.15. This analysis was performed separately for lymphoblastoid and fibroblast cell lines. Species-specific 5mC gain promoters were defined as those with 5mC values at least 0.25 greater than the maximum 5mC values in other species. Conversely, species-specific loss of 5mC were defined as 5mC values more than 0.25 lower than the minimum 5mC values in other species.

We also computed how the rank of each gene’s expression level (ranging from 0 for the lowest expressed genes to 8,174 for the highest expressed genes) for the corresponding genes of those promoters changed compared to the rank of that gene in other species (**Table MET.S1-2**). RNA-seq data from matching NHP samples were downloaded from SRA (under BioProject accession no. PRJNA101639). The sequenced reads were then directly quantified using Salmon v1.10.0 under the mapping-based mode, where a Salmon index for each NHP was first built using a decoy-aware transcriptome generated from the UCSC Genome Browser TOGA_hg38 annotation track. The transcript-level expression transcripts per million (TPM) generated by Salmon was aggregated into gene-level TPMs. For Human (hg002), preliminary RNA-seq data was obtained from Genome in a Bottle Consortium, and processed similarly using Salmon against transcripts annotated under UCSC Genome Browser's CuratedRefSeqv5.1 annotation track.

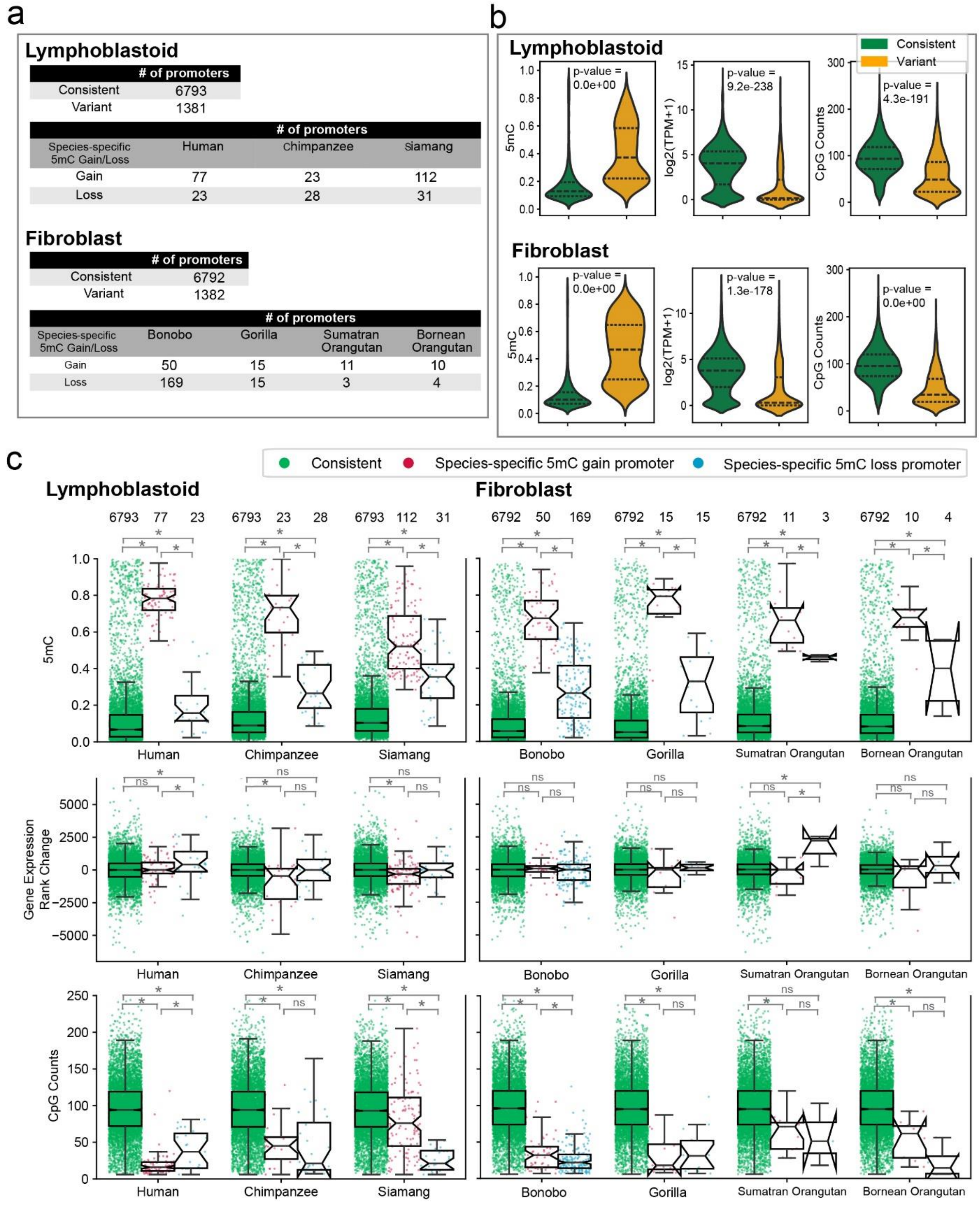


**Figure MET.S1. DNA methylation analysis of 8,174 orthologous promoters between humans and NHPs.** We analyzed data from lymphoblastoid and fibroblast cell lines separately.

(a) The number of consistently or variantly methylated promoters among species and the number of promoters that show significant 5mC gain or loss in specific species. The majority (~83%) of promoters are consistently methylated between NHP species (<15% variation), indicating that promoter methylation tends to be conserved during the evolution of apes.

(b) Consistently methylated promoters tend to be lowly methylated, more highly expressed, and have a higher density of CpG sites compared to variantly methylated promoters. P-values based on the two-sided Mann–Whitney U test are shown, and all are significant.

(c) Comparisons of promoters exhibiting species-specific 5mC gain and loss versus those consistently methylated across NHP species. Each dot represents a promoter. P-values are determined using the two-sided Mann–Whitney U test ($p<0.05$: '*', $p<0.01$: '**', or 'ns'). **The top panel** shows 5mC levels of those promoters for each species. Promoters that change DNA methylation between species tend to be more highly methylated than consistently methylated promoters, regardless of whether they gain or lose DNA methylation. **In the middle panel**, we computed the ranks of each gene’s expression (ranging from 0 for the lowest expressed genes to 8,174 for the highest expressed genes within each species) and calculated how gene ranks changed in a species compared to the ranks of the genes in other species. We examined how they are distributed compared to in other species for consistently methylated promoters and those that changed DNA methylation. While consistently methylated promoters show a distribution around 0 (meaning that their expression ranks remained constant), promoters with species-specific 5mC gain tend to show negative values below 0 in gene expression rank changes, indicating that their expression rank decreased compared to in other species. In contrast, those with 5mC loss tend to show positive values above 0 in gene expression rank changes, indicating a trend of higher expression in the species compared to in other species. Nevertheless, these trends were not always significant, as the numbers of 5mC gain or loss promoters were relatively small. **In the bottom panel,** we show that promoters that change DNA methylation between species tend to have fewer CpG sites compared to consistently methylated promoters.

### XIII. Replication timing

Contributing authors:

Jian Ma, Muyu Yang, Yang Zhang, David Gilbert, Takayo Sasaki, Gabrielle A. Hartley, Emry Brannan, & Rachel J. O'Neill

#### Methods

##### Cross-species replication timing profiling and evolutionary patterns identification

Replication timing profiling of primate species followed the same procedure described previously[126]. Repli-seq was processed based on a published workflow[127]. Cutadapt (version 4.2) removed the remaining adapters on the reads with parameters "-q 0 -O 1 -m 0", using the adapter sequence AGATCGGAAGAGCACACGTCTG. Reads were mapped to the human (T2T-CHM13v2.0 and NHP assemblies with bwa mem (version 0.7.17)[128] with default parameters. PCR duplicates were removed using SAMtools rmdup tool.

The genomes were segmented into nonoverlapping 5 kbp bins, and mapped reads were counted and normalized to RPKM. Bins with less than 0.1 RPKM across both early and late fractions were removed. The log2 ratio of early to late fractions was calculated and normalized using interquartile range (IQR) normalization, i.e., (value – median)/IQR. Ratios were smoothed with the loess method with 300 kbp window size.

For cross-species comparison, reciprocal liftover converted NHP replication timing profiles to the human T2T genome (chm13, version 2.0). The human genome was segmented into nonoverlapping 5 kbp bins and mapped to NHPs using liftover[129], retaining only reciprocally mapped bins. Next, Phylo-HMGP[126], a continuous-trait probabilistic model, inferred evolutionary states for functional genomic signals, identifying 20 states representing distinct evolutionary patterns of replication timing profiles across primate species.

#### Summary of results

##### Evolutionary patterns of replication timing

To characterize changes in replication timing across the primate species, we applied Phylo-HMGP[126], focusing on lymphoblastoid cell lines from human, chimpanzee, Bornean orangutan, and siamang. Phylo-HMGP identified 20 states with distinct evolutionary patterns (**Fig. RT.S1**A). These states were classified into five categories based on the replication timing signal values and the variability across the species: early (E), weakly conserved early (WE), late (L), weakly conserved late (WL), and non-conserved (NC). We observed that 31.5% of the genome is in conserved early state and 21.5% in the conserved late state. Lineage-specific states were mainly reflected by weakly conserved and NC states. For example, State 13 has a unique early replication timing specific to chimpanzees, while State 4 shows a distinct late replication timing unique to Bornean orangutans. The distribution of replication timing signals for each state, organized by category, is displayed in **Fig. RT.S1**B. **Fig. RT.S1**C shows an example of replication timing profiles and Phylo-HMGP states on the genome browser.

**Correlation between SDs and replication timing evolutionary patterns**

We analyzed the relationship between SDs and replication timing, focusing on lineage-specific, nonhomologous sequence elements. The genome was divided into 5 kbp bins, and replication timing signals were calculated for SDs and the genome-wide background. A Wilcoxon rank-sum test assessed the differences between SDs and the background. We found that ancient SDs shared across many species (e.g., BCHGO and BCHGOS) tend to replicate early (p-value < 1e-20), while more recent SDs (e.g., O and BC) and species-specific SDs tend to replicate late (p-value < 1e-50). Examples of the replication timing distribution are shown in **Fig. RT.S2**.

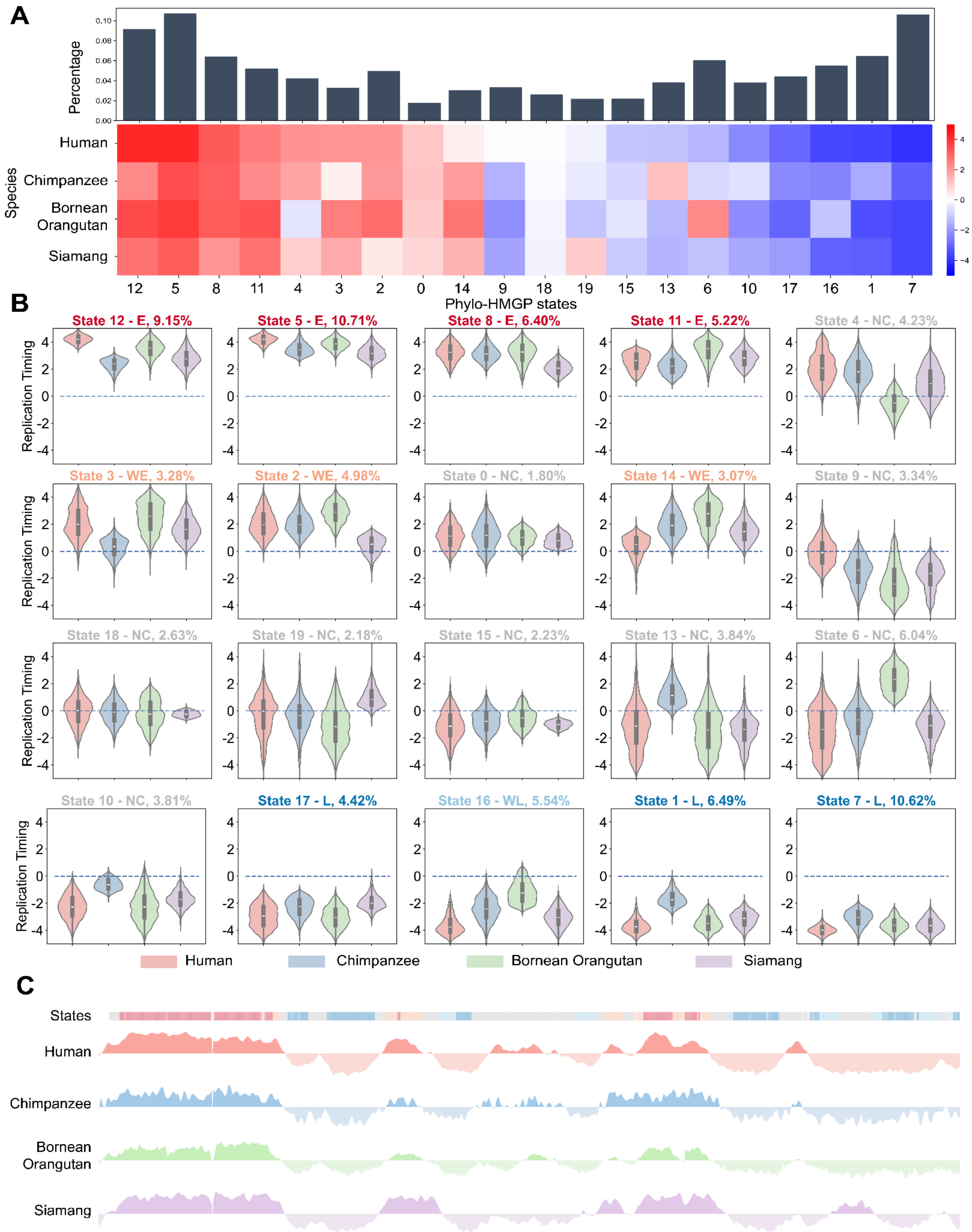


**Figure RT.S1. Phylo-HMGP identifies 20 states with distinct evolutionary patterns of replication timing in lymphoblastoid cells from four primary species. (A)** The top panel shows the proportion of each state across the entire genome. The bottom panel illustrates the average replication timing signals in each state, with columns representing different states ordered by the average human replication timing signals. **(B)** The 20 Phylo-HMGP states are categorized into five groups: early (denoted as E), weakly conserved early (WE), late (L), weakly conserved late (WL), and non-conserved (NC). Violin and box plots show the replication timing distributions for each state, organized by category. **(C)** Visualization of replication timing patterns and state annotations in the genome browser (chr1:50,000,000-75,000,000 in human T2T genome version 2.0).

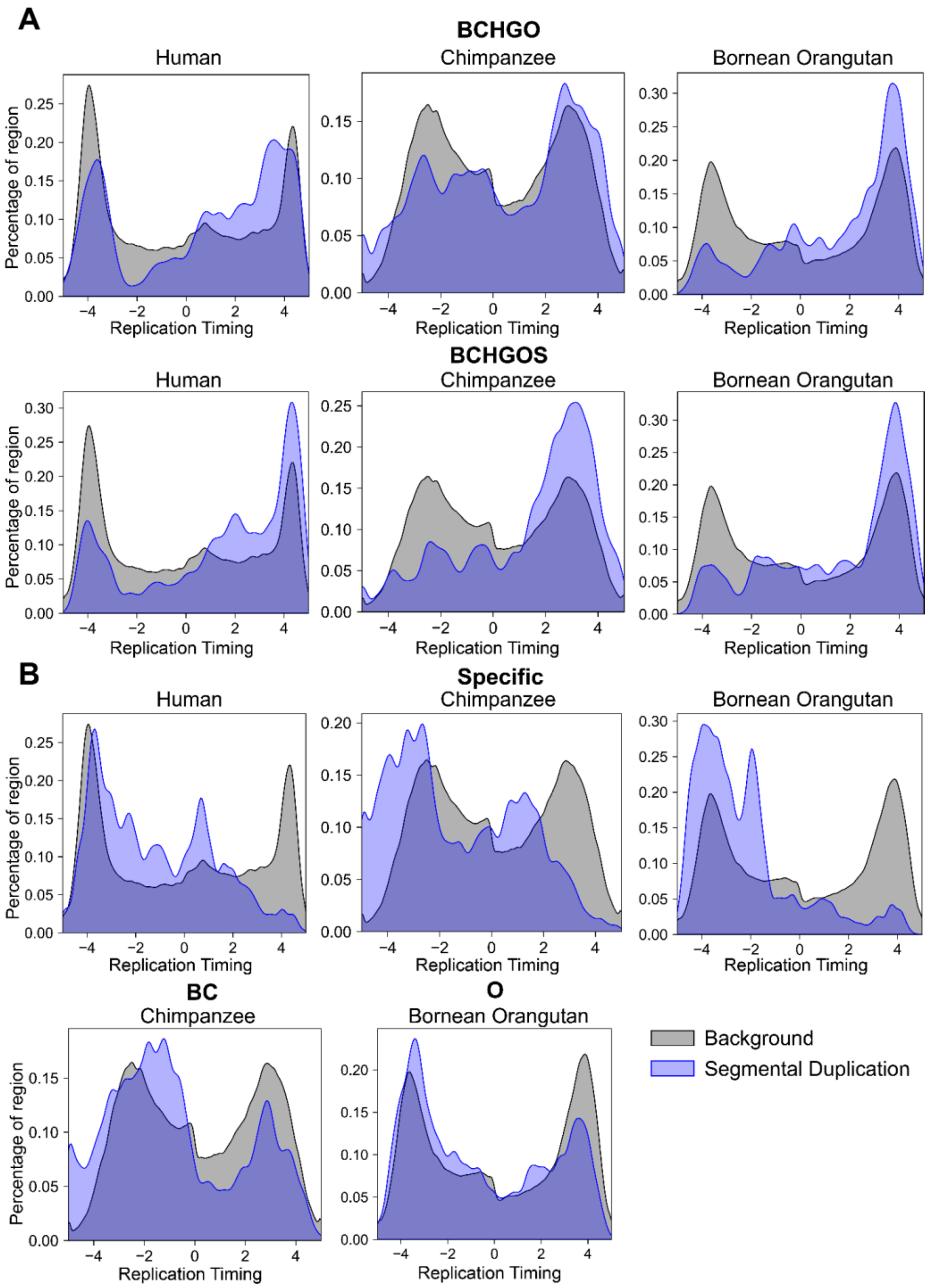


**Figure RT.S2. The correlation between segmental duplications (SDs) and replication timing (RT).** The blue density curves represent the distribution of RT signals in SD regions, and the gray density curves represent the background RT signals. **(A)** Distribution of RT signals in more ancient SDs, including BCHGO (shared among bonobo, chimpanzee, human, gorilla, and orangutan) and BCHGOS (shared among bonobo, chimpanzee, human, gorilla, orangutan, and siamang). **(B)** Distribution of RT signals in more recent SDs, including Specific (species-specific), BC (shared between bonobo and chimpanzee), and O (shared between two types of orangutans).

### XIV. Evolutionary rearrangements and inversion characterization

Contributing authors:

Francesca Antonacci, DongAhn Yoo, Luciana de Gennaro, David Porubsky, Mario Ventura

#### Methods

##### SV calling

We used the SYRI (v1.6.3)[130] and PAV (v2.3.2)[54] pipelines to identify SVs (>50 bp), including inversions, in six diploid genomes. This was done using the human T2T genome (T2T-CHM13v2.0) and Bornean orangutan's primary haplotype as the reference to check for effect of reference bias. SYRI inversions in general represented larger inversions, and the PAV-called inversions were added after merging inversion calls that share the same breakpoints (reciprocal coverage of 80%). Alignment artefacts (less than 50% supported by alignment or 1 Mbp of alignment support-minimap2) were further filtered to the merged set of inversion calls by SYRI and PAV. For the comparison to previous studies, inversions with size exceeding 10 kbp were used.

##### Validation against previous studies

We rigorously compared our identified inversions against an array of previously characterized large-scale inversions from cytogenetic studies and smaller inversions validated through single-cell strand-seq, fluorescence in situ hybridization (FISH), BAC clone sequencing, BAC-end mapping, and PCR[22,35,58,131-151]. This was done by comparing the inversion breakpoints liftovered onto hg38.

#### Summary of results

Near-T2T ape genome assemblies provide a more comprehensive map of inversions across the ape phylogenetic lineage. We identified a total of 1,175 inversion variants larger than 10 kbp as well as one large-scale chromosome fusion and one translocation (**Fig. INV.S1**) across six species; the breakdown of identified inversions is as follows: 188 in bonobo, 171 in chimpanzee, 199 in gorilla, 180 in Bornean orangutan, 180 in Sumatran orangutan, and 257 in siamang. Compared with the previously documented inversions, we find 558 represent novel discoveries. Focusing on the larger ones (>1 Mbp), we find that 160 were previously documented inversions and 29 are novel. Further observing 27 erroneous calls, 23 of which coming from siamang. We resolved 40 breakpoints of the Yunis and Prakash large chromosome inversions; all breakpoints differ by a maximum of ~700 kbp from previous cytogenetic mapping (**Table INV.S1**). In six instances, boundaries involving the centromere or telomere have been resolved for the first time, and in two instances the T2T genome assemblies revealed a more complex organization than reported by cytogenetics (orangutan chromosomes 2 and 19).

Filtering out the false inversion calls of 1,175 inversions larger than 10 kbp, 1,140 inversion calls remain. Among these, 617 are previously known and 522 are novel. Additionally, our complete assemblies allowed us to refine the breakpoints of 85/617 known inversions and revealed that one event (Jim_GGO.hap1.INVTR_4), which was previously classified as an inversion in gorilla{Porubsky, 2020 #85} is an inverted transposition. The transposition was confirmed with two FISH experiments on metaphase chromosomes in one gorilla individual (**Fig. 4b**). Out of the 1,140 inversions, 632 are homozygous (found in both haplotypes of the same individual), while 508 are heterozygous and potentially polymorphic in the population. Looking for genes mapping at the breakpoints, we found that 416 inversions have genes mapping at least at one breakpoint, while 724 events are completely devoid of genes at their breakpoints.

Of the 1,140 inversions, 529 have SDs at both breakpoints (46%), with 78% (412 out of 529) having SDs in inverted orientation. Additionally, 195 inversions (17%) have SDs at only one breakpoint, and 416 inversions are devoid of SDs at both breakpoints.

Focusing on just the 632 homozygous inversions, 197 are novel and 435 are previously known. The 197 novel homozygous inversions have an average size of 223 kbp and mostly map within pericentromeric regions and/or SDs. In particular, 72% (141/197) have annotated human SDs at one or both ends of the inversions, versus 43% (186/435) of the known homozygous inversions, which could explain why they were not detected in previous studies. Of the novel homozygous inversions, 78 have annotated genes mapping at least one of the breakpoints with 119 devoid of protein-coding genes (**Table INV.S2**).

Investigating 100 kbp flanking regions of inversions, we find that inversion breakpoints are typically enriched for T2T human SDs ($p<0.001$) compared to random genomic regions by 4.6-fold (flanking of the randomly shuffled inversions; **Fig. INV.S2**). We also observed that the enrichment was stronger for inverted duplications ($p<0.001$) with 6.2-fold more overlap compared to null genomic regions. Stronger enrichment was observed restricting the test to larger inversions (>50 kbp).

Of the inversions without SDs, 289 have other types of repeat elements mapping at both breakpoints and 35 have repeats at only one breakpoint, while 92 are completely devoid of any repeats.

Among the 523 novel inversions, we identified 78 in chimpanzee, 99 in bonobo, 96 in gorilla, 71 in Sumatran orangutan, 68 in Bornean orangutan, and 111 in siamang. All the novel inversions are smaller than 5 Mbp, just below the limit of cytogenetic resolution. Nearly half of these novel inversions (227 out of 523) have SDs at one or both breakpoints when mapped to the human reference genome. This complexity may have contributed to these inversions being overlooked in previous studies. Additionally, 325 out of the 523 novel inversions are heterozygous and potentially polymorphic, which may account for their absence in earlier analyses performed on different individuals (e.g., Porubsky et al.[131]). For 160 novel inversions, genes mapped at one or both breakpoints, while no genes were detected at the breakpoints of the remaining 363 inversions.

Based on the parsimony of homozygous inversions among the apes (comparing the breakpoints with 80% reciprocal overlap), we assigned 64-90% of inversion events (64% in bonobo and up to 90% in siamang) onto phylogeny (**Fig. INV.S3**) and predict the remaining inversions to be recurrent. The number of inversions were particularly less in certain lineages, including HSA which showed fivefold less than expected number considering the branch length. Among the phylogeny-classified inversions, we find a total of 133 new inversions, with the highest number gained from siamang lineage (n=44). We also find the number of inversion events correlated with phylogeny (R2 = 0.77).

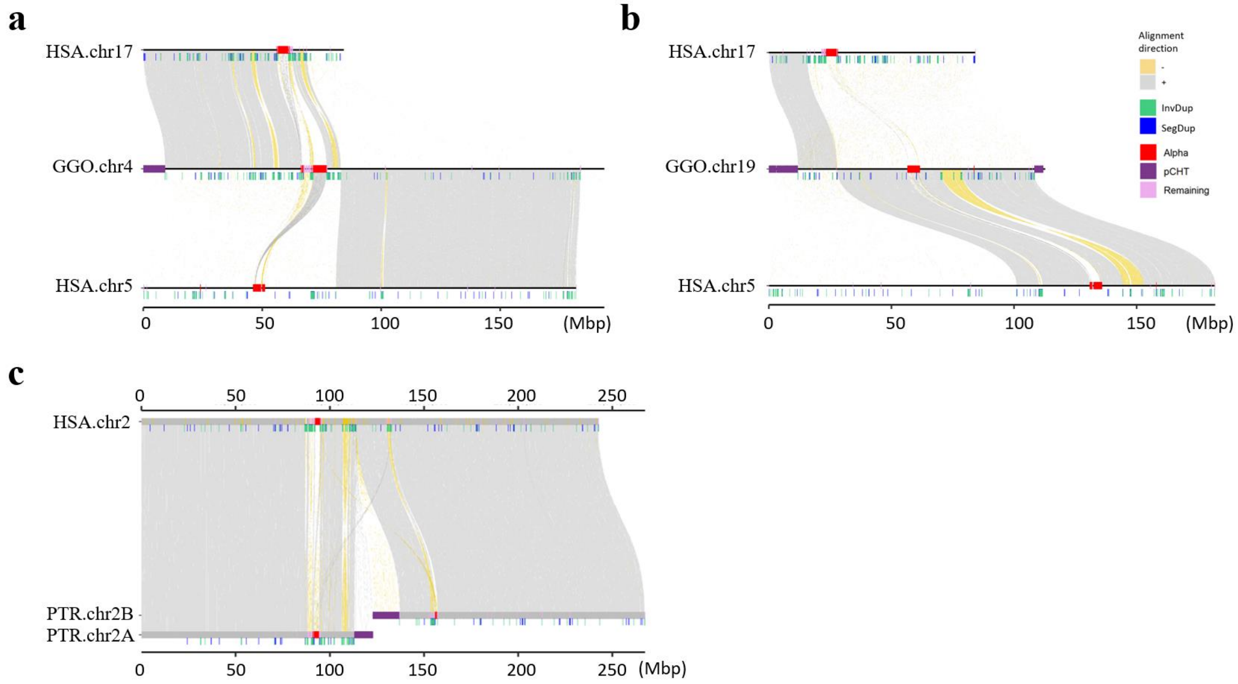


**Fig.INV.S1. Chr4/19 translocation in gorilla and chr2 fusion in human.** Alignment view of gorilla chr4 **(a)** and chr19 **(b)**. **c)** Alignment of human chr2 fusion.

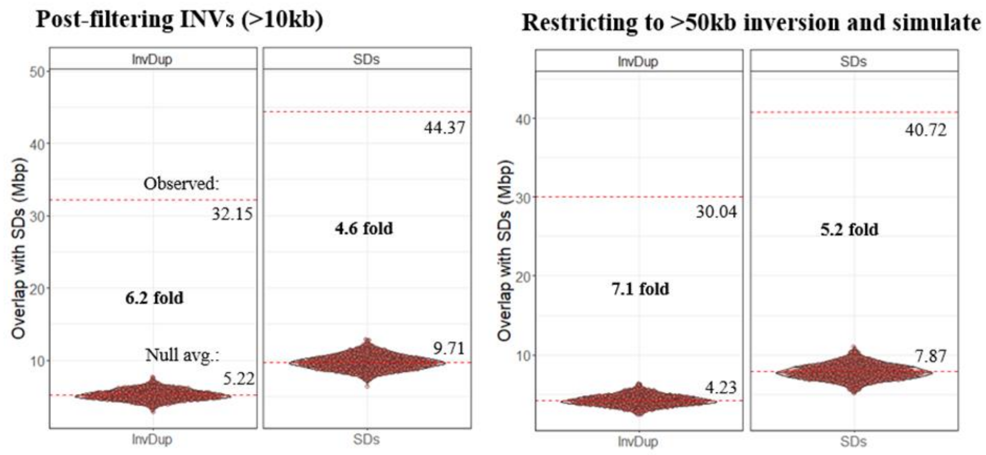


**Figure INV.S2. Enrichment of segmental duplications at the breakpoints (100kbp flanking) of inversions.** Left shows the enrichment for the inversions (INVs) in size >10kbp, while the right shows the enrichment for >50kbp inversions.

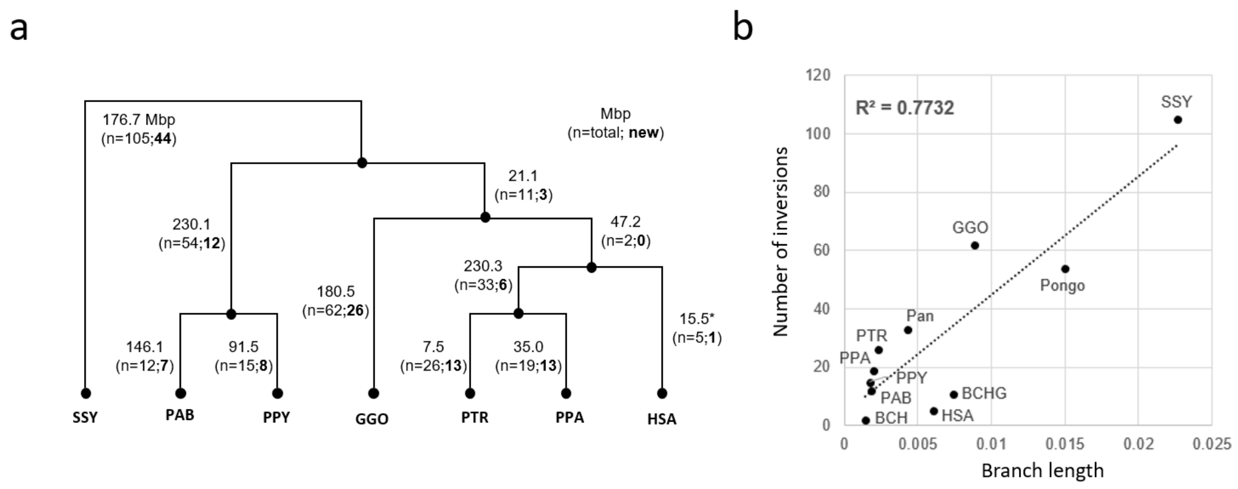


**Figure INV.S3. Inversion assignment into phylogeny and correlation of inversions with phylogenetic branch lengths.** Indicated by "*" are the human-specific inversion calls which are found in inverted orientation in all other apes compared to human.

### XV. AQER

Contributing authors:

Yanting Luo, Manolis Kellis, Riley J. Mangan, Sarah Zhao, Chul Lee, Youngho Lee, Erich D. Jarvis, Craig B. Lowe

#### Methods

We defined highly divergent regions in four great ape lineages by calculating the number of mutations that likely occurred in 500 bp regions between inferred ancestral nodes and extant genomes. The four lineages we analyzed were the inferred human-chimpanzee ancestor to the human (hs1), chimpanzee (primary haplotype), and bonobo (primary haplotype) genomes, as well as the inferred human-gorilla ancestor to the gorilla (primary haplotype) genome. We term these regions Ancestor Quickly Evolved Regions (AQERs), with the individual sets being (human) HAQERs, chimp-AQERs, bonobo-AQERs, and gorilla-AQERs. The identification of AQERs is not limited to conserved regions, but rather screens the entire genome, including elements that descended from previously neutral regions. This distinguishes HAQERs from many other searches for the genetic underpinnings of uniquely human traits, which have focused on human-specific divergence in conserved genomic regions and reflects modifications of existing functional elements. AQERs were identified in three steps. First, both haplotypes from the nearly complete great ape assemblies (https://github.com/marbl/Primates) were aligned to the T2T human assembly (hs1). Alignment fragments scoring greater than 60,000 (approximately 20 kbp) were retained. For each of the four lineages being analyzed for highly divergent regions, we inferred the sequence of the ancestral node in a conservative fashion so that uncertain positions (e.g., no base with a probability greater than 0.8) were assigned the value observed in the extant species. Finally, the number of mutations that separate the inferred ancestral node and the extant genome were counted by sliding windows of 500 bp along the hs1 assembly. Windows with significantly more divergences than expected (>29 mutations in a 500 bp window) were identified as HAQERs, chimp-AQERs, bonobo-AQERs, and gorilla-AQERs. Additionally, we ensured that chimp-AQERs, bonobo-AQERs, and gorilla-AQERs could be mapped to continuous sections of their reference genomes.

For *ADCYAP1*, to assemble methylation profiles of six ape species for downstream analyses of vocal-learning related genes, public raw PacBio long reads generated with 5-base HiFi sequencing with kinetics and methylation tags were downloaded from Human Pangenome Reference Consortium (HPRC, https://humanpangenome.org/data/) for human and GenomeArk (https://www.genomeark.org/t2t-all/) for chimpanzee, gorilla, Bornean orangutan, Sumatran orangutan, and siamang gibbon. First, all reads were merged into a single file for each species by SAMtools version 1.16.1 using SAMtools merge. Then, the reads were aligned to T2T genome assemblies for each of the respective species by pbmm2 version 1.10.0 using pbmm2 align. Aligned reads were sorted and indexed by SAMtools using SAMtools sort and SAMtools index, respectively. Pre-included methylation tags of the raw reads were called using align_bam_to_cpg_scores.py provided pb-CpG-tools version 2.3.2 (https://github.com/PacificBiosciences/pb-CpG-tools). Then, modification probability of all CpGs of genomes was calculated based on tags of reads mapped to the corresponding sites and adjusted based on overall distributions of modification scores using a machine-learning model pileup_calling_model.v1.tflite provided by pb-CpG-tools. After pileup of the modification probabilities, we classified CpGs with scores over 75% as hypermethylated, between 25-75% as heteromethylated, and under 25% as hypomethylated.

**Summary of results**

Nearly complete great ape assemblies resolve multi-scale divergence landscapes across human and great ape genomes

We investigated how the availability of nearly complete genome assemblies for humans and great apes improves the identification of highly divergent regions within the great apes. Previous searches for highly divergent regions on the human lineage in conserved regions have revealed functional modifications in regulatory elements[160] and divergent elements in unconstrained sequence are associated with *de novo* functional elements[161].

We identified 14,210 AQERs across four great ape lineages (3,268 HAQERs, 4,001 chimp-AQERs, 4,231 bonobo-AQERs, and 2,710 gorilla-AQERs) (**Fig. AQER.S1a, Methods**). Particular classes of repetitive elements are enriched or depleted for these highly diverged sequences across all four lineages. All four AQER sets show enrichments for simple repeats, low-complexity DNA, and satellites; however, all the lineages also show depletions for MEI, including the SINEs, LINEs, and LTRs. There are some particular MEI families that show enrichments, such as SVAs due to their internal VNTR region. There are also shared features among the four sets in terms of their location in relation to gene structures. All show a strong depletion for overlapping protein-coding exons, and a weaker depletion for transcribed regions, yet an enrichment for occurring in promoter sequences on all four lineages.

Having nearly complete great ape genomes has greatly expanded the original set of 1,581 HAQERs derived from gapped assemblies to 3,268. These T2T HAQERs are more highly enriched in SDs, simple repeats, and centromeric satellite regions, which were previously difficult to assemble (**Fig. AQER.S1**b). These new regions from T2T genomes also show evidence of gene regulatory function based on location in the genome and their overlap with epigenomic annotations in the EpiMap dataset[162]. Of these new HAQERs, 464 show evidence of promoter function through being located within 2 kbp upstream or 500 bp downstream of a transcription start site, or overlapping epigenetic modifications associated with promoters, and 684 new HAQERs show evidence of enhancer function based on their overlap with chromatin states associated with enhancer function.

We investigated whether rapid sequence divergence is associated with particular classes of functional elements. While there is limited functional genomic data for chimpanzees, bonobos, and gorillas, we were able to analyze the HAQERs in the context of epigenomes from the Roadmap Epigenomics Consortium and EpiMap[163]. Both T2T and Gapped HAQERs exhibit enrichments for bivalent promoter and enhancer chromatin states across diverse tissues (**Fig. AQER.S1c**). Bivalent domains are thought to contain gene regulatory elements that exhibit precise spatiotemporal activity patterns in the context of development and environmental response[164]. Notably, we observed more pronounced enrichments in the union set of Gapped and T2T HAQERs. We propose two possible explanations for this observation. First, the union set of HAQERs may refine the Gapped HAQER set by removing elements misidentified as fast-evolving due to ortholog misalignments or other errors in the gapped alignments, which have been resolved by T2T great ape assemblies. Second, by analyzing HAQERs in assemblies from different individuals within each species, this union set may partially control for intraspecific variation, thereby biasing the union set towards regions with fixed differences. Bivalent chromatin state enrichments were not observed in fast-evolving regions from other great apes, which may reflect limited cross-species transferability of epigenomic annotations in these regions, potentially due to functional divergence in these regions of elevated sequence divergence.

HAQERs were initially identified by scanning the genome for regions of elevated local divergence in 500 bp windows. To determine if functional annotation enrichments in HAQERs were dependent on this ascertainment window size, we generated HAQER sets at a range of sliding window sizes, from 25 bp to 20 kbp, to create multi-scale divergence landscapes of the human genome (**Fig. AQER.S1d**). This analysis revealed some rapidly evolving elements are only detectable at specific window sizes (**Fig. AQER.S1d**, left and right), and that many larger-scale rapidly evolving elements are composites of multiple, constituent regions at smaller scales (**Fig. AQER.S1d**, middle). While bivalent gene regulatory and heterochromatin state enrichments for HAQERs are heavily dependent on the ascertainment window size, HAQERs were consistently depleted from transcribed regions across window sizes (**Fig. AQER.S1e**).

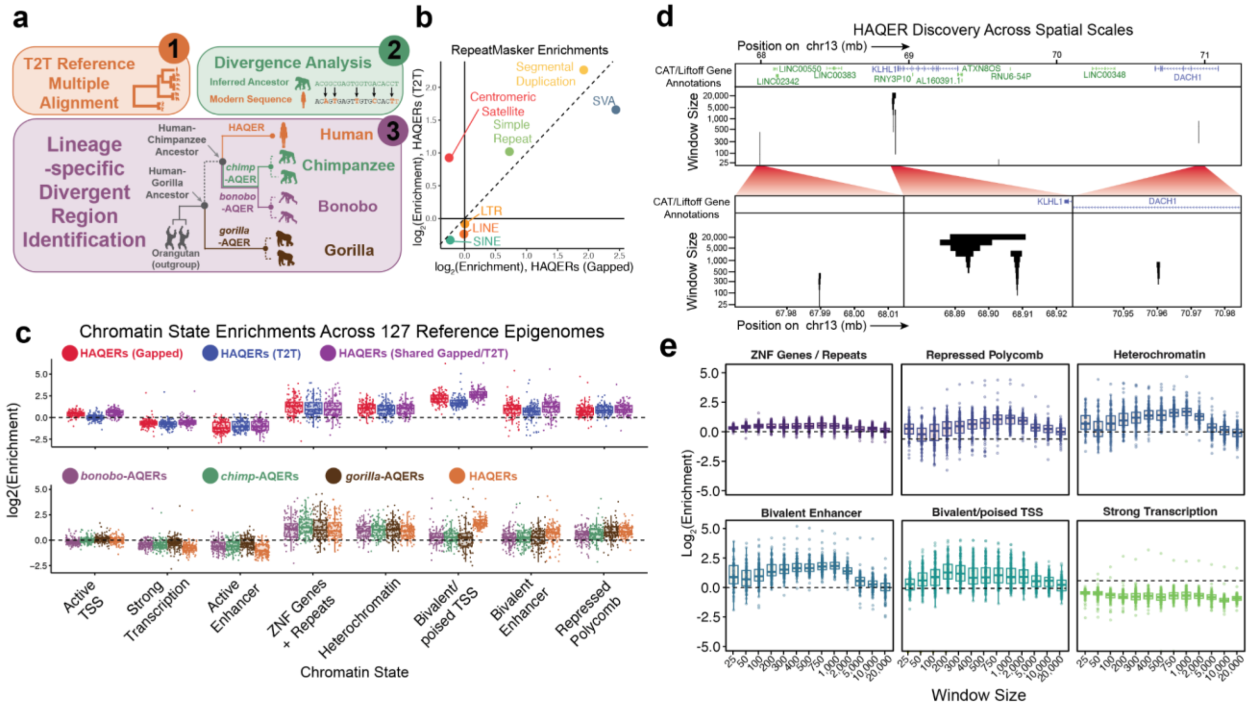


**Figure AQER.S1. Great ape T2T assemblies resolve highly divergent genomic regions.** (a) We ascertained HAQERs (human ancestor quickly evolved regions), and similarly fast-evolving regions in other great ape species, by identifying regions of elevated sequence divergence in 500 bp windows. (b) The newly added HAQERs tend to be located in centromeric sequences, simple repeats, and SDs. (c) The HAQER sets based on both gapped and T2T assemblies show enrichments for bivalent gene regulatory elements across 127 cell types and tissues. The set of HAQERs shared between these two sets shows an even stronger enrichment for this functional state. The tendency for HAQERs to occur in bivalent regulatory elements is not present in the sets of bonobo-AQERs, chimpanzee-AQERs, or gorilla-AQERs. (d) HAQER genomic locations, observed across spatial scales, in a 4 Mbp region of chr13. Bottom: 50 kbp zoomed-in inserts. (e) The influence of divergence scale on chromatin state enrichment

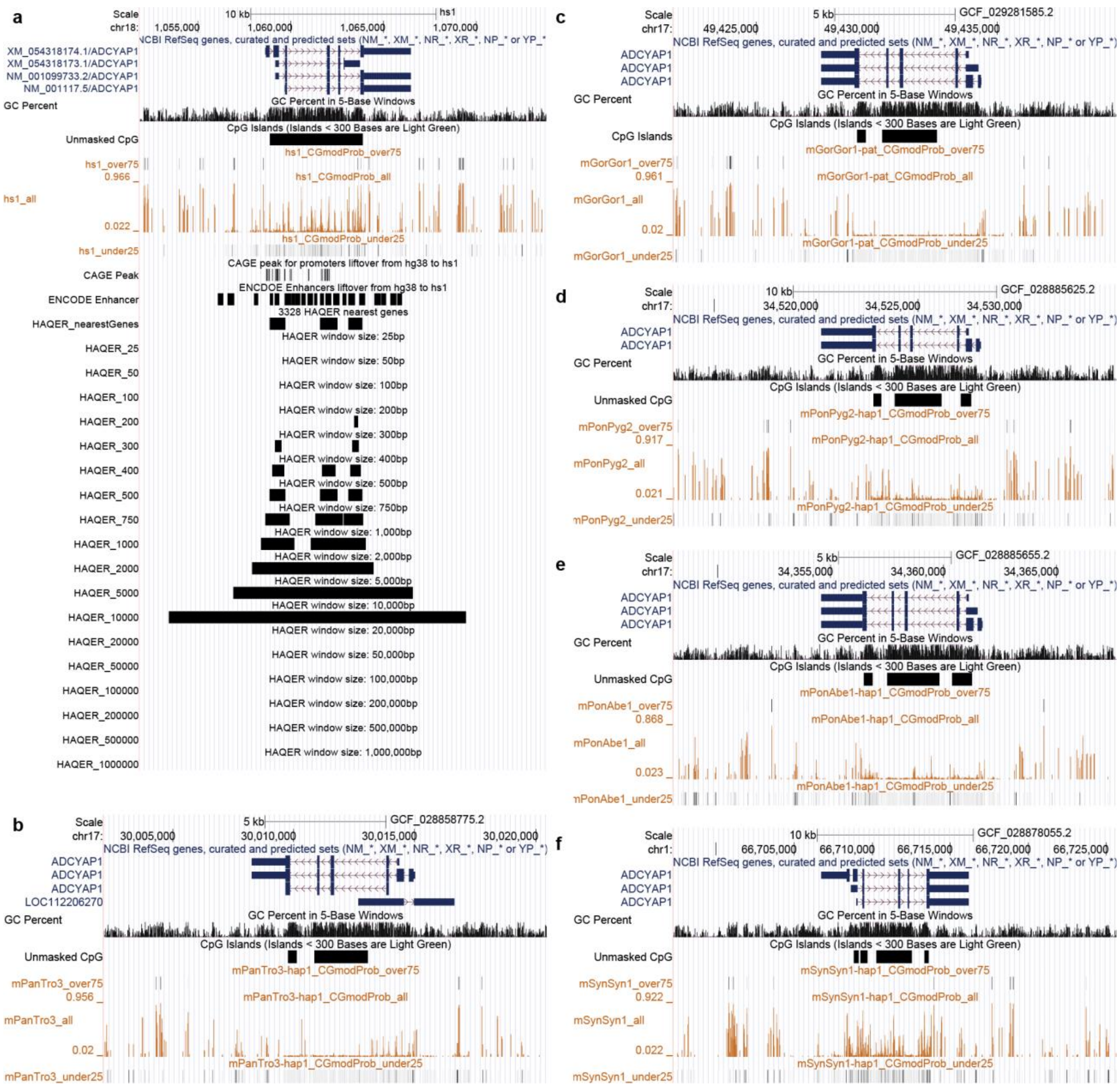

**Figure AQER.S2. DNA hypo-methylation pattern on *ADCYAP1* gene bodies of apes.** (a) Human, (b) chimpanzee, (c) gorilla, (d) Bornean orangutan, (e) Sumatran orangutan, and (f) siamang gibbon. All panels show NCBI annotation, GC percent, CpG islands, DNA hyper-methylated loci (probability > 75%), DNA methylation probabilities on CpG, and DNA hypo-methylated loci (probability < 25%). The methylation probabilities estimated from PacBio HiFi reads. CAGE peaks and ENCODE enhancers were lifted over from hg38 to hs1. HAQERs were identified in different windows (25 bp-10 Mbp) and HAQER nearest genes were classified by considering transcription start sites of each gene.

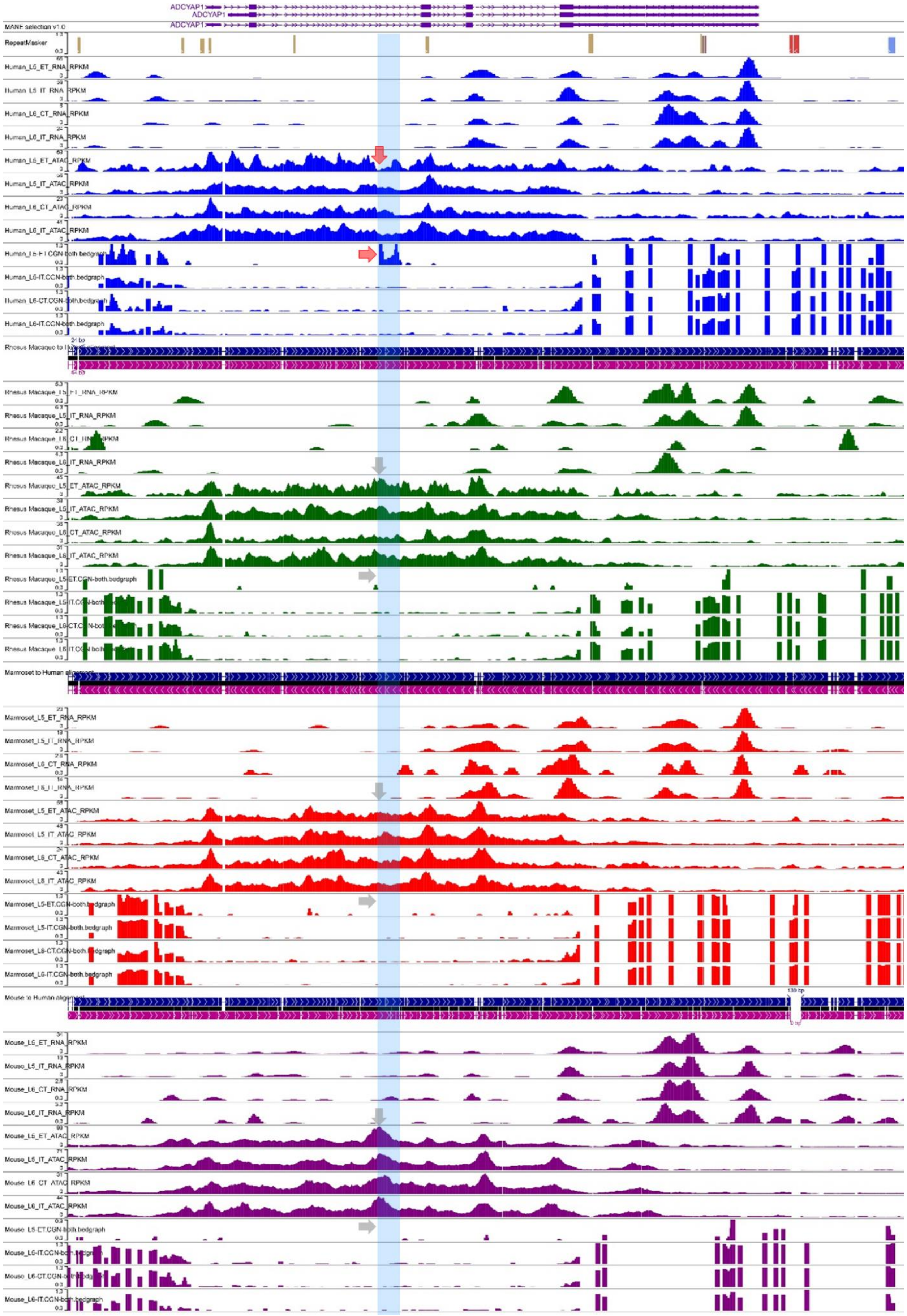
ADCYAP1
ADCYAP1
ADCYAP1
MANE selection v1.0
RepeatMasker
Human_L5_ET_RNA_RPKM
Human_L5_IT_RNA_RPKM
Human_L6_CT_RNA_RPKM
Human_L6_IT_RNA_RPKM
Human_L5_ET_ATAC_RPKM
Human_L5_IT_ATAC_RPKM
Human_L6_CT_ATAC_RPKM
Human_L6_IT_ATAC_RPKM
Rhesus Macaque_L5_ET_RNA_RPKM
Rhesus Macaque_L5_IT_RNA_RPKM
Rhesus Macaque_L6_IT_RNA_RPKM
Rhesus Macaque_L5_ET_ATAC_RPKM
Rhesus Macaque_L5_IT_ATAC_RPKM
Rhesus Macaque_L6_CT_ATAC_RPKM
Rhesus Macaque_L6_IT_ATAC_RPKM
Rhesus Macaque_L5-ET.CGN-both.bedgraph
Marmoset to Human alignment
Marmoset_L5_ET_RNA_RPKM
Marmoset_L5_IT_RNA_RPKM
Marmoset_L6_CT_RNA_RPKM
Marmoset_L6_IT_RNA_RPKM
Marmoset_L5_ET_ATAC_RPKM
Marmoset_L5_IT_ATAC_RPKM
Marmoset_L6_CT_ATAC_RPKM
Marmoset_L6_IT_ATAC_RPKM
Marmoset_L5-ET.CGN-both.bedgraph
Marmoset_L5-IT.CGN-both.bedgraph
Marmoset_L6-CT.CGN-both.bedgraph
Marmoset_L6-IT.CGN-both.bedgraph
Mouse to Human alignment
Mouse_L5_ET_RNA_RPKM
Mouse_L5_IT_RNA_RPKM
Mouse_L6_CT_RNA_RPKM
Mouse_L6_IT_RNA_RPKM
Mouse_L5_ET_ATAC_RPKM
Mouse_L5_IT_ATAC_RPKM
Mouse_L6_CT_ATAC_RPKM
Mouse_L6_IT_ATAC_RPKM
Mouse_L5-ET.CGN-both.bedgraph
Mouse_L6-IT.CGN-both.bedgraph

**Figure AQER.S3. Human-unique epigenetic patterns different from macaque, marmoset, and mouse.** Each four rows show scRNA seq read depths, scATAC seq read depths, and DNA methylation probabilities of layer 5 ET / IT and layer 6 CT / IT neurons of primary motor cortexes of human, macaque, marmoset, and mouse. Modified from Comparative Epigenomics browser (https://epigenome.wustl.edu/BrainComparativeEpigenome/).

### XVI. Structurally divergent regions

Contributing authors:

Jiadong Lin, Junmin Han, Shilong Zhang, Yafei Mao

#### Methods

The all-vs-all alignment created to detect lineage-specific SDs was used to identify lineage-specific structurally divergent region (SDRs) on each haplotype. For primate lineages (i.e., PTR, PAB, GGO, PAB, PPY, SSY), sequences that were not aligned or aligned of identity <85% were considered as divergent regions specifically to each species on the leaf node of the ape phylogeny. The human lineage-specific (HSA) SDRs were those conserved between human haplotypes (i.e., CHM13 and HG002) but divergent from other primate haplotypes. For the Pan lineage, we first obtained regions on PTR that are not aligned to other species except for PAB. We then subtract regions that are specific to PTR from the regions obtained in the previous step. We used PAB and applied the same approach to obtain Pongo lineage-specific SDRs. The SDRs were further annotated with centromere, acrocentric, subterminal, secondary constriction (qh) regions, and euchromatin. To count the SDR bases by genomic content, we classified SDRs in the order of centromere, acrocentric, subterminal, sex chromosome, and others. For the others category, we further examined whether it overlapped qh region and those non-overlapped parts were classified as euchromatin. Note that for centromere and subterminal, we also considered bases that are not in the centromere or subterminal as euchromatin.

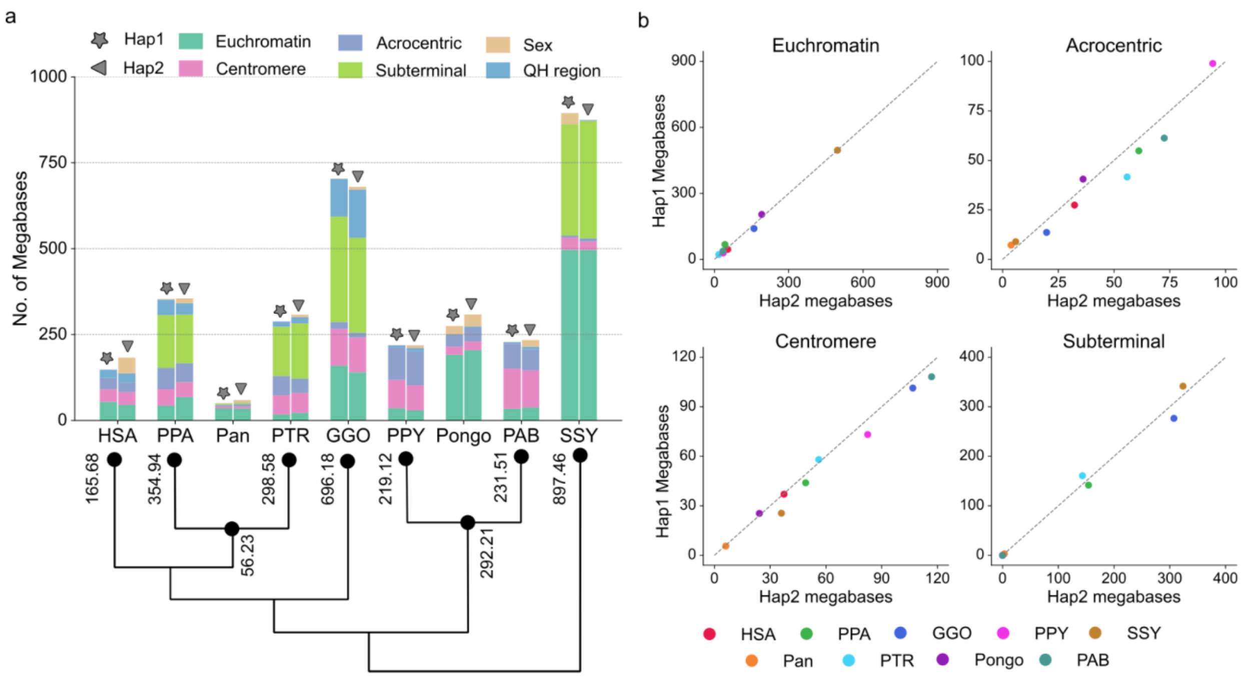


**Figure SDR.S1. Lineage-specific structurally divergent regions (SDRs) on each haplotype.** **(a)** The total megabases of SDRs detected on each haplotype. The average megabases of the two haplotypes were assigned to the phylogenetic tree. We used PTR and PAB to represent the ancestral node Pan and Pongo, respectively. **(b)** The comparison of SDR total megabases separated by euchromatin, acrocentric, centromere, and subterminal.

### XVII. TOGA analysis

Contributing authors:

Agnes Chan, Michael Hiller, Nicholas J. Schork

#### Method

TOGA (Tool to infer Orthologs from Genome Alignments) inference resource[152] was used to identify human gene loci that were reported as "absent" across more than 40 primate species. The pipeline was applied to the current six ape T2T assemblies to identify and refine genes lost during ape evolution.

#### Summary of results

##### TOGA human-specific genes

We identified six candidate genes that are likely unique to the human lineage and 19,148 (out of 19,244, 99.5%) protein-coding genes that are present in two or more of the six ape T2T assemblies. TOGA predictions for the primate assemblies (group 1) and the ape pre-T2T assemblies (group 2) as of 2023 were collected from Kirilenko et al.[152] The existing ape assemblies from 2023 included silvery gibbon, northern white-cheeked gibbon, Sumatran orangutan, western lowland gorilla, pygmy chimpanzee, and chimpanzee. New TOGA predictions were generated in this study for six ape T2T assemblies. Human-specific gene candidate sequences were identified based on TOGA predictions that reported gene absence across over 80% of the assemblies analyzed. A summary of human genes selected from each assembly group based on the "over 80% absence" criteria are shown in bold in **Table TOGA.S1**. Additional evidence was collected from the T2T-CHM13 UCSC Genome Browser, including whole-genome alignments of the ape T2T assemblies against T2T-CHM13, and SD predictions from this study. For one of the candidate genes, *FOXO3B*, we carried out in-depth validations using nucleotide comparison including flanking sequences and confirmed its absence across five of the ape T2T primate genomes except gorilla, and its unique genome architecture embedded in a clinically relevant locus in the human genome.

We describe an example of human-specific gene sequences related to the major longevity gene *FOXO3*, which encodes a transcription factor with prevalent functional roles in regulating apoptosis, autophagy, and metabolism[153]. The association of *FOXO3* with longevity was first reported in a Japanese-American Hawaiian cohort[153] and subsequently replicated in multiple European cohorts (e.g., Flachsbart et al.[154]). Proposed mechanisms for the longevity phenotype of *FOXO3* included a *FOXO3* haplotype-induced chromatin hub spanning multiple transcription factor binding sites, and its transcript isoforms[155,156]. Here, we leveraged the TOGA ortholog inference resource to identify human gene loci that were reported as "absent" across more than

40 primate species included in the TOGA analysis (**Fig. TOGA S1a**). *FOXO3B* was one of the top-ranked genes that were not detected across almost all primate species analyzed. In terms of gene structure, the annotated *FOXO3B* gene model shares 5' exons with a zinc finger protein, carries multiple but incomplete merged exons from *FOXO3*, and therefore has the properties of a processed gene. *FOXO3B* mRNA expression by RT-qPCR has been reported across multiple tissues, including the cerebellum, fetal brain, and neural progenitor cells and cell lines[157].

Using high-quality, long-read derived genome assemblies from the ape T2T project, we confirmed the unique presence of *FOXO3B* sequences in the human genome. Across the ape T2T species, *FOXO3B* sequence was not found in chimpanzee, bonobo, orangutan, or gibbon. *FOXO3B* was also not found in the New World monkey marmoset. Of note, the *FOXO3B* sequence was detected in the gorilla genome, though with a different genomic architecture than the human reference assemblies (i.e., T2T-CHM13, GRCh38) (**Fig. TOGA S1b**). CHM13 Iso-Seq data supports expression of *FOXO3B* locus.

We investigated the genomic context of the human *FOXO3B* locus and revealed a complex structure involving five local low-copy repeats or SDs of ~200 kbp at chromosome 17 near the centromere. We mapped the location of the *FOXO3B* locus to one of the low-copy repeats embedded within a 4 Mbp region at 17p11.2 (**Fig. TOGA S1c**). Previous studies have shown that microdeletion or microduplication of this region is associated with Smith-Magenis syndrome (SMS) and Potocki-Lupski syndrome (PLS), respectively[158]. The clinical impacts of SMS and PLS include developmental delay and cognitive impairment. The causal gene of SMS and PLS is believed to be *RAI1* located within 17p11.2, but whether *FOXO3B* could be involved in cognitive impairment requires further analysis.

It is worth noting that complex co-regulation networks of a parent gene and its pseudogene counterpart involving coding and noncoding mechanisms have been reported in transcription factors with significant roles in tumor suppression (e.g., *PTEN* and the pseudogene *PTENP1*), and embryonic and stem cell development (e.g., *POU5F1/OCT4* and five pseudogenes *POUR5F1P1* to *POUR5F1P5*)[159].

In summary, we discovered the human-specific property of the *FOXO3B* gene locus through comparative analysis of human and primate assemblies and ortholog annotation. Many questions remain, such as: the functional significance of *FOXO3B* in longevity, apoptosis, etc., whether *FOXO3* and *FOXO3B* could be subjected to co-regulation networks, and, if so, in which tissues/cell types and developmental stages, and the ancestral origin and evolution of the *FOXO3B* locus across humans and the great apes.

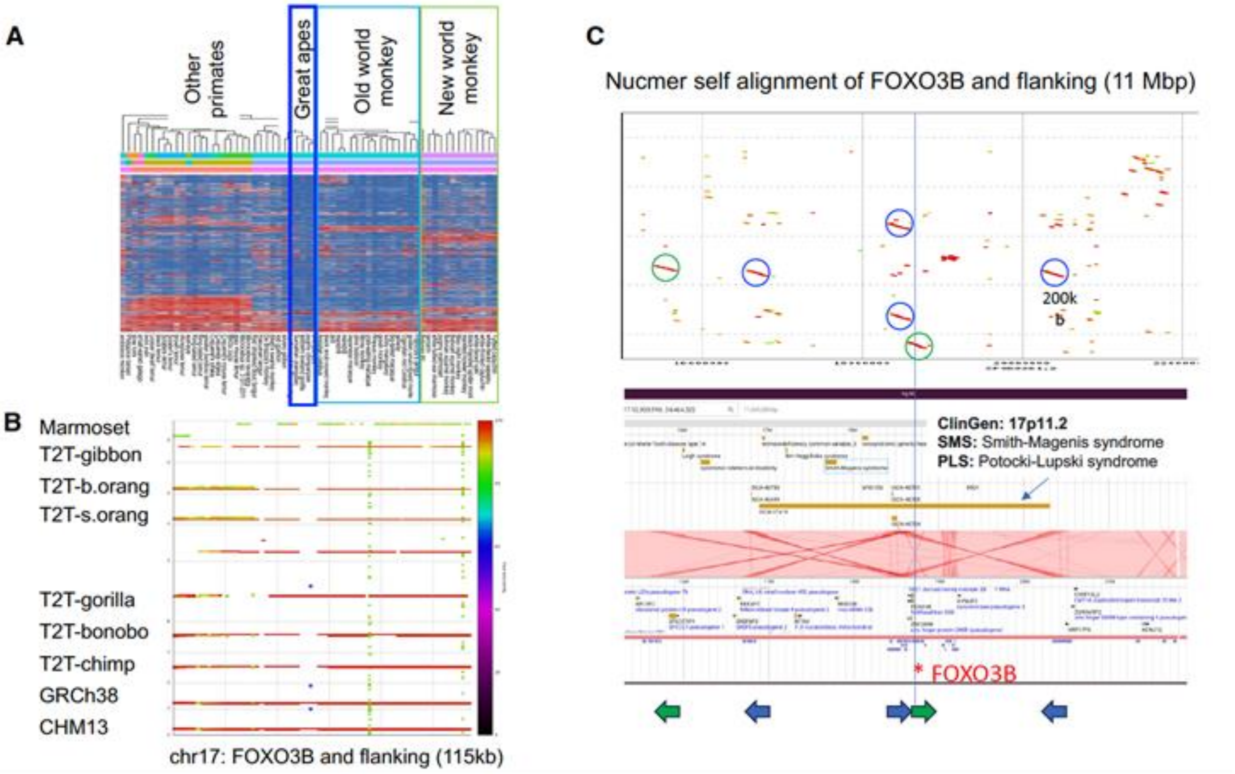


**Figure TOGA S1. Human-specific gene sequences encoding FOXO3-like sequences.**
A) Predicted gene loss (red) across over 40 primate species as reported by TOGA. B) The 7 kbp *FOXO3B* (asterisk) was only detected in the human (CHM13, GRCh38) and the T2T gorilla genome assemblies, yet absent from other T2T primate genomes. C) *FOXO3B* resides with a 4 Mbp clinically significant locus known to link to cognitive impairment.

### XVIII. Acrocentric region analysis

Contributing authors:

Steven J. Solar, Alexander P. Sweeten, Graciela Monfort Anez, Matthew Borchers, Tamara Potapova, Jennifer L. Gerton, Adam M. Phillippy

##### Methods

To identify rDNA-containing chromosomes and array orientations, a reference human 45S unit (GenBank accession KY962518) was mapped against all primate genomes using mashmap v3.1.1:

```
mashmap -r $primate_ref -q 45S.fa -t $cpus --pi 90 -s 13332 --filter_mode none -o $primate.45S.mashmap
```

The resulting mappings were manually validated, filtering out lower identity pseudogenes that were not part of intact arrays, and all chromosomes containing rDNA arrays were noted. These results were further confirmed by FISH.

##### Chromosome spreads and FISH

For the preparation of chromosome spreads, cells were blocked in mitosis by the addition of Karyomax colcemid solution (0.1 µg/ml, Life Technologies) for 6-7h and collected by trypsinization. Collected cells were incubated in hypotonic 0.4% KCl solution for 12 min and prefixed by addition of methanol:acetic acid (3:1) fixative solution (1% total volume). Pre-fixed cells were collected by centrifugation and then fixed in Methanol:Acetic acid (3:1). Spreads were dropped on a glass slide and incubated at 65°C overnight. Before hybridization, slides were treated with 0.1mg/ml RNAse A (Qiagen) in 2xSSC for 45 minutes at 37°C and dehydrated in a 70%, 80%, and 100% ethanol series for 2 minutes each. Slides were denatured in 70% formamide/2X SSC solution pre-heated to 72°C for 1.5 min. Denaturation was stopped by immersing slides in 70%, 80%, and 100% ethanol series chilled to -20°C. Labeled DNA probes were denatured separately in a hybridization buffer by heating to 80°C for 10 minutes before applying to denatured slides. Fluorescently labeled probe for human rDNA (BAC clone RP11-450E20) was obtained from Empire Genomics. Fluorescently labeled whole chromosome paints for chromosomes 2, 9, 13, 14, 15, 18, 21, and 22 were obtained from Applied Spectral Imaging. Human CenSat probe for D14Z1/D22Z1 was obtained from Cytocell. The probe for labeling distal junction (DJ) regions was prepared from the BAC CH251-351B7 (Eichler lab) and labeled with Biotin-16-dUTP using the nick translation kit (Enzo Life Sciences). Specimens were hybridized to the probes under a glass coverslip or HybriSlip hybridization cover (GRACE Biolabs) sealed with the rubber cement or Cytobond (SciGene) in a humidified chamber at 37°C for 48-72hours. After hybridization, slides were washed in 50% formamide/2X SSC 3 times for 5 minutes per wash at 45°C, then in 1x SSC solution at 45°C for 5 minutes twice, and at room temperature once. For biotin detection, slides were incubated with streptavidin conjugated to Cy5 (Thermo) for 2-3 hours in PBS containing 0.5% Triton X-100 and 5% bovine serum albumin (BSA), and then washed 3 times for 5 minutes with PBS/0.5% Triton X-100. Slides were mounted in Vectashield containing DAPI (Vector Laboratories). Wide-field images were acquired on the Nikon TiE microscope equipped with 100x objective NA 1.4 and Prime 95B sCMOS camera (Photometrics). Z-stack images were acquired on the Nikon TiE microscope equipped with 100x objective NA 1.45, Yokogawa CSU-W1 spinning disk, and Flash 4.0 sCMOS camera (Hamamatsu).

**Estimating rDNA copy number from FISH images**

Image processing was performed in FIJI and Python. Primate rDNA-containing chromosomes were identified as homo sapiens (HSA) homologs based on the labeling with human chromosome paints and morphological features. For chimpanzee and bonobo, painting hsa 14 and hsa 21 was sufficient to identify all rDNA-containing chromosomes, and for gorilla, hsa 22 paint alone was sufficient. For Sumatran and Bornean orangutans, all rDNA-containing chromosomes were painted on separate slides, and these data were aggregated across all slides. For human HG002 spreads, labeling centromeric satellite 14/22 was sufficient to identify all rDNA-containing chromosomes. Chromosome Y was identified by morphology.

For manual image quantifications performed for chromosome spreads from chimpanzee, bonobo and gorilla cells, sum intensity projections of confocal Z-planes were generated, and individual rDNA arrays were segmented based on threshold applied to the entire image. The fluorescence intensity of the regions of the same chromosomes that did not contain the rDNA was used to subtract the local background. The background-subtracted integrated intensity was measured for each array. For semi-automated quantification performed for chromosome spreads from Sumatran orangutan, Bornean orangutan, and human HG002 cells, wide-field single Z-plane images were used. rDNA-containing chromosomes were segmented using a Cellpose model trained on 2-channel images, including the DAPI and rDNA signals. rDNA regions were also segmented using a trained Cellpose model. The chromosome segmentations were examined and, if necessary, curated manually in Napari. rDNA intensities for each array were measured after subtracting the fluorescence background for the respective chromosomes. Custom Python and scripts are deposited in GitHub: https://github.com/jouyun/2024_Primate_rDNA.

The sum of all intensities of all rDNA loci represented the total amount of rDNA per cell, and the fraction of this total signal was calculated for each rDNA array. The total rDNA copy number was estimated from Illumina sequencing data (see “Estimating rDNA copy number from *k*-mer coverage”). The fraction of the total rDNA fluorescence intensity was used as a proportion of the total rDNA copy number to determine the number of rDNA copies on specific chromosomes in each chromosome spread.

##### Estimating rDNA copy number from *k*-mer coverage

Genomic DNA from primate cell lines was isolated using QIAamp DNA Micro Kit (Qiagen) according to the manufacturer's protocol. PCR-free DNA-seq libraries were constructed using NEBNext Ultra II DNA Library Prep Kit and sequenced on AVITI System (Element Biosciences) outputting 150 bp paired-end reads. Data for HG002 was obtained from Baid et al.[165], using the PCR-free 40× coverage whole-genome sequencing sample. rDNA copy numbers were estimated from *k*-mer frequencies in the whole-genome sequencing data. A reference 18S sequence was set for each species by choosing a single representative unit from rDNA loci identified by mashmap. The 18S copy number served as a proxy for the greater 45S unit, as each unit contains a single 18S segment. A custom pipeline counted *k*-mers of size 31 from the 18S consensus in short-read Illumina sequencing data and normalized it to counts of 31mers from G/C matched windows elsewhere in the rDNA containing chromosomes. The matched windows were of similar size to the 18S, and 30 were randomly selected per rDNA-containing chromosome. Any *k*-mers that also occurred outside the matched windows were removed to ensure that counts were exclusively from the matched windows. *k*-mer sets were filtered to remove those with whole-genome sequencing counts greater than three standard deviations from the mean of the set, or those that were missing entirely, and counts were divided by their genomic multiplicity. Finally, the median count from the 18S *k*-mers was divided by the median count of the matched windows to yield a copy number approximation. Three replicates of this process were done for each primate, as the G/C matching step had a random component, and the integer-rounded mean of the three replicates was taken as the copy number. A pipeline referred to as CONKORD (version 7) was used for this process. A slightly different version of CONKORD was used for HG002, which did not filter *k*-mers at high standard deviations or zero counts (version 5). CONKORD contains a set of custom Python and Bash scripts, which are deposited in GitHub (https://github.com/borcherm/primate_rdna_cn).

To identify distal junction (DJ) locations and orientations in the primate assemblies, a reference DJ was extracted from T2T-CHM13 as defined by Sluis et al.[166] extending from the end of the CER block at T2T-CHM13v2.0 chr13:5,424,523 to the beginning of the rDNA array at chr13:5,770,548. This reference DJ was aligned to the primate assemblies using minimap2 v2.28 and filtered for hits ≥ 100 kbp:

```
minimap2 -x asm20 -t $cpus -c --MD $primate_ref DJ.fa | awk -v OFS='\t' '{ if ($11 >= 100000) print }' > DJ.$primate.paf
```

This process was repeated aligning the DJ just chrY to look for possible remnant DJs using the same commands.

The DJ palindrome was identified by dot plotting the reference DJ using mummer v4.0.0 and manually identifying the start and end of each palindromic arm:

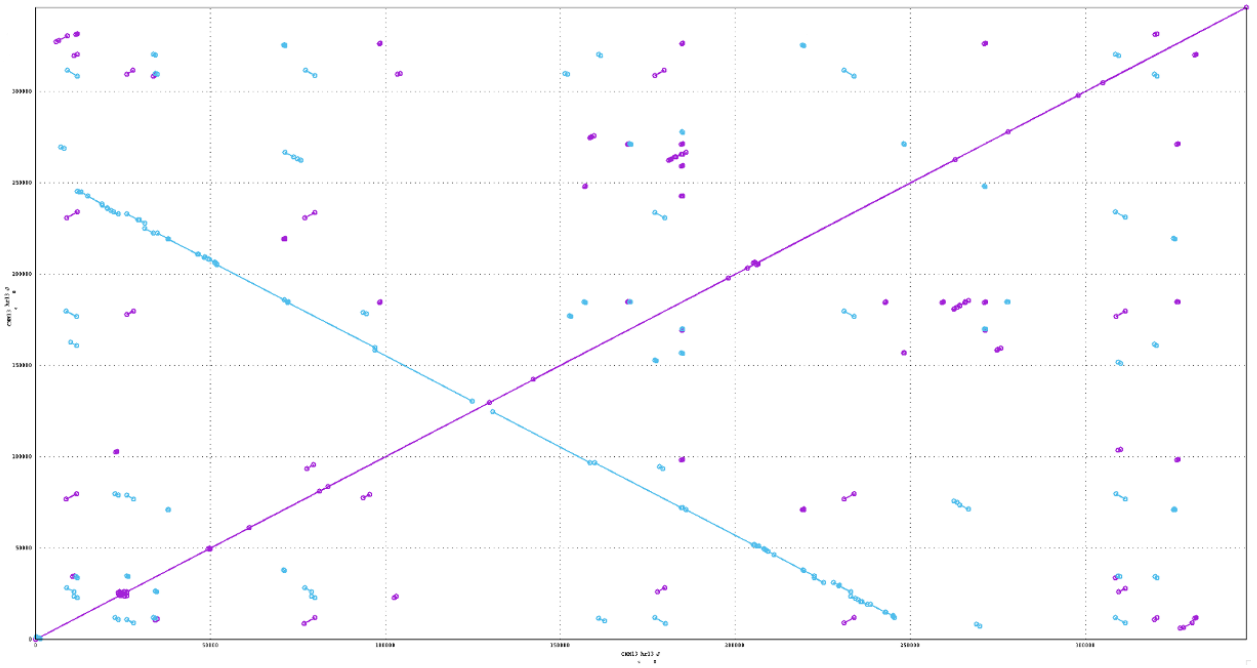

**Figure Acro.S1. Structure of the human distal junction (DJ).** A plot of maximal exact self-matches of at least 20 bp in the CHM13 chr13 DJ. Forward matches are indicated in purple and reverse matches in blue. The large X shape indicates the presence of an inverted repeat, in this case the characteristic DJ palindrome encoding the long ncRNA. The precise boundaries of this palindrome were extracted.

mummer -maxmatch DJ.fa DJ.fa > DJ.out

mummerplot DJ.out

chr13 5436542 5549309

chr13 5555214 5670026

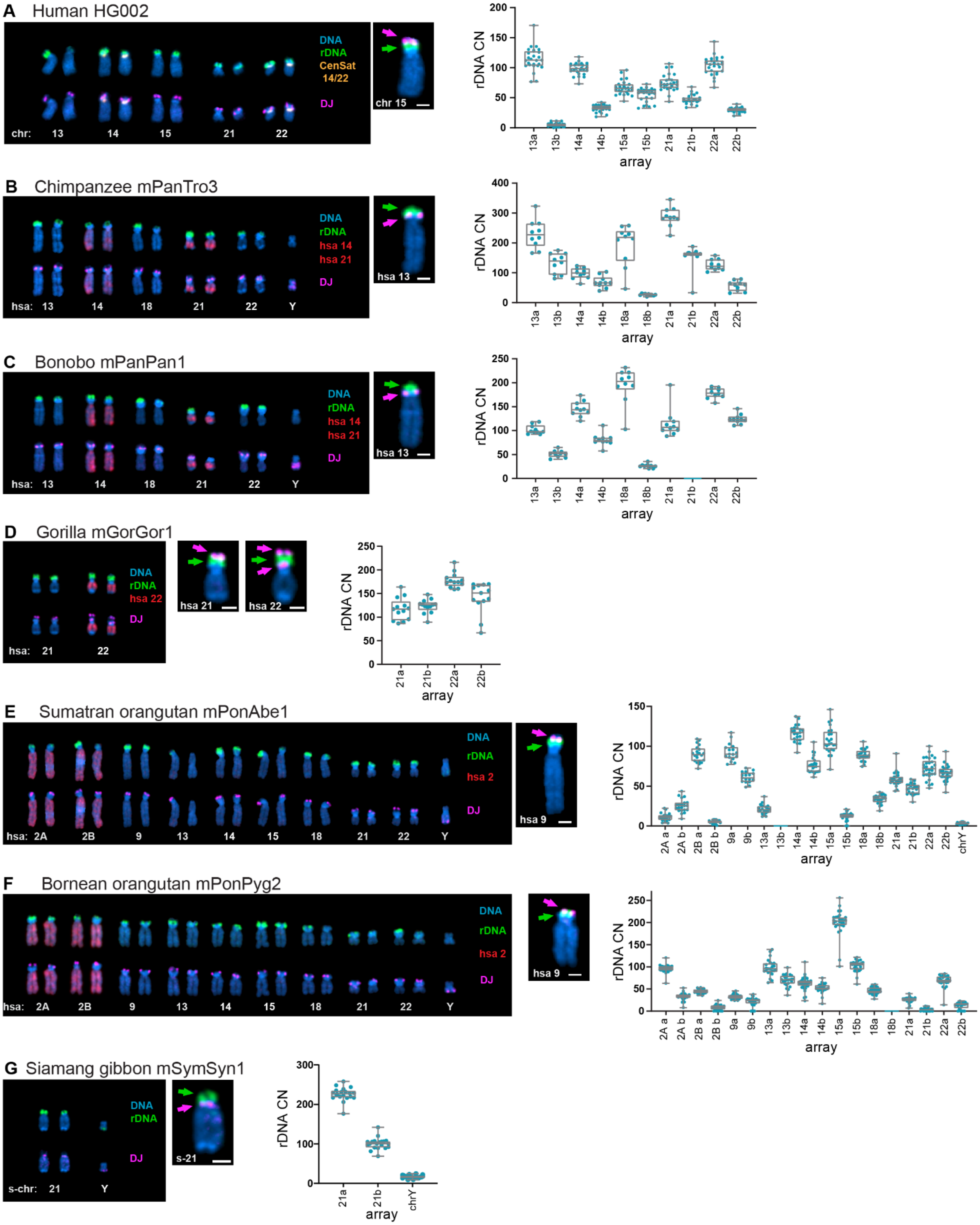


**Figure Acro.S2. Representative karyograms of NOR+ chromosomes in cells from human (A), chimpanzee (B), bonobo (C), western lowland gorilla (D), Sumatran orangutan (E), Bornean orangutan (F) and Siamang gibbon (G).** Primate chromosomes were identified as homo sapien (HSA) homologs except Siamang-specific rDNA-containing chromosome 21. The top chromosome rows show FISH labeling with the rDNA probe (green) and chromosome identification markers. For chromosome identification, human CenSat 14/22 probe (orange) or indicated human whole chromosome paints (red) were used. The bottom rows show labeling with the DJ-region probe (magenta). DNA was counter-stained with DAPI. Side panels show overlayed images of representative individual chromosomes. Corresponding quantifications of rDNA copy number are shown on the right. The boxes represent the interquartile range, with the edges indicating the upper and lower quartiles. The line inside the boxes indicates the median. Whiskers show the range from minimum to maximum values. Ten or more spreads were quantified for each specimen. All individual data points are shown. Estimated total numbers of rDNA units and rDNA units per haplotype for each species are listed in **Table Acro.S1**.

A second gorilla NOR on both haplotypes of HSA22 was identified by this DJ mapping using repeat-aware Winnowmap (meryl v1.4.1, Winnowmap v2.03, SAMtools v1.19):

```
meryl count k=15 output $primate.k15.DB $primate_ref

meryl print greater-than distinct=0.9998 $primate.k15.DB > $primate.repetitive_k15.txt

winnowmap -W $primate.repetitive_k15.txt -ax map-ont --MD ref.fa ont.fq.gz -y "MM,ML" | samtools sort -O BAM --write-index -o $primate.ONT.bam##idx##$primate.ONT.bam.bai > $primate.ONT.sam
```

This was confirmed by viewing the inverted duplication with ModDotPlot.

To assess rDNA copy number and activity in the second NOR, the mapped human 45S location was viewed in IGV along with ONT read alignments containing methylation calls.

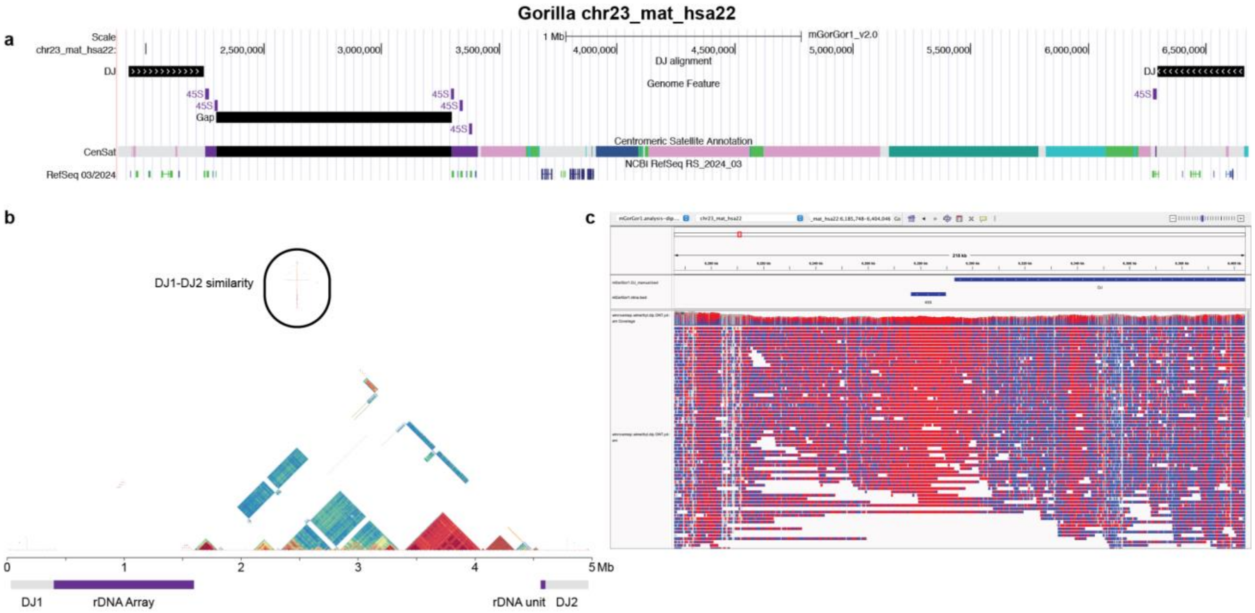


**Figure Acro S3. NOR structure of gorilla HSA22.** a) Viewing of alignments of the human DJ and 45S gene to gorilla's maternal HSA22 indicates the presence of two NORs. One faces toward the centromere (as in HSA21) and contains a full rDNA array; the other faces towards the telomere and contains a single rDNA unit. This is also the case in the paternal haplotype. b) A self-ModDotPlot of the same region shows high similarity between the two DJs (labeled DJ1 and DJ2). c) Whole-genome alignments of ONT reads show that the single rDNA unit associated with DJ2 is hypermethylated (red) in both haplotypes, which is known to indicate inactivity[1].

To assess rDNA conservation across species, representative units were extracted from the assemblies. The human 45S gene was minimapped to the assembly with the following command:

```
minimap2 -x asm20 -t $cpus -c --MD $primate_ref 45S.fa > DJ.$primate.paf
```

Then, the sequence from the start of one 45S to the start of the next was extracted using BEDTools (v2.29.0) in each primate and flipped when necessary to match the transcriptional direction, to serve as a representative unit, acknowledging that intra-species variation will exist. 45S and intergenic spacer (IGS) sequences were extracted using the coordinates from the original alignment. Multiple sequence alignments of rDNA units, 45S sequences, and the IGS were generated using Mafft-linsi v7.526:

```
mafft-linsi --thread 32 all.rdna.fa > msa.fa
```

This was used to compute pairwise gap-excluded percent identities for each region with a simple python script.

**Table Acro S2. Pairwise comparison of representative rDNA units.** To compare conservation of the entire rDNA unit (a), the 45S gene (b), and the intergenic spacer (c), pairwise gap-excluded identities were calculated from multiple sequence alignments of each region.

| **a) rDNA Identity** | | | | | | | |
|---|---|---|---|---|---|---|---|
| | HSA | PTR | PPA | GGO | PAB | PPY | SSY |
| HSA | 1 | 0.9563 | 0.9576 | 0.9491 | 0.8918 | 0.8914 | 0.8485 |
| PTR | 0.9563 | 1 | 0.9872 | 0.9484 | 0.893 | 0.8927 | 0.8668 |
| PPA | 0.9576 | 0.9872 | 1 | 0.9487 | 0.8932 | 0.8928 | 0.8693 |
| GGO | 0.9491 | 0.9484 | 0.9487 | 1 | 0.8973 | 0.8971 | 0.8794 |
| PAB | 0.8918 | 0.893 | 0.8932 | 0.8973 | 1 | 0.9934 | 0.8668 |
| PPY | 0.8914 | 0.8927 | 0.8928 | 0.8971 | 0.9934 | 1 | 0.8671 |
| SSY | 0.8485 | 0.8668 | 0.8693 | 0.8794 | 0.8668 | 0.8671 | 1 |

| **b) 45S Identity** | | | | | | | |
|---|---|---|---|---|---|---|---|
| | HSA | PTR | PPA | GGO | PAB | PPY | SSY |
| HSA | 1 | 0.9676 | 0.9696 | 0.9651 | 0.9395 | 0.9392 | 0.9363 |
| PTR | 0.9676 | 1 | 0.9889 | 0.9654 | 0.9406 | 0.9409 | 0.9355 |
| PPA | 0.9696 | 0.9889 | 1 | 0.9656 | 0.9407 | 0.9409 | 0.9352 |
| GGO | 0.9651 | 0.9654 | 0.9656 | 1 | 0.9414 | 0.9414 | 0.9351 |
| PAB | 0.9395 | 0.9406 | 0.9407 | 0.9414 | 1 | 0.9944 | 0.9324 |
| PPY | 0.9392 | 0.9409 | 0.9409 | 0.9414 | 0.9944 | 1 | 0.9331 |
| SSY | 0.9363 | 0.9355 | 0.9352 | 0.9351 | 0.9324 | 0.9331 | 1 |

| c) IGS Identity | | | | | | | |
|---|---|---|---|---|---|---|---|
| | HSA | PTR | PPA | GGO | PAB | PPY | SSY |
| HSA | 1 | 0.9511 | 0.9519 | 0.9432 | 0.8687 | 0.8678 | 0.806 |
| PTR | 0.9511 | 1 | 0.9867 | 0.9403 | 0.867 | 0.8659 | 0.8304 |
| PPA | 0.9519 | 0.9867 | 1 | 0.9405 | 0.8674 | 0.8661 | 0.834 |
| GGO | 0.9432 | 0.9403 | 0.9405 | 1 | 0.8749 | 0.874 | 0.849 |
| PAB | 0.8687 | 0.867 | 0.8674 | 0.8749 | 1 | 0.9928 | 0.8316 |
| PPY | 0.8678 | 0.8659 | 0.8661 | 0.874 | 0.9928 | 1 | 0.8302 |
| SSY | 0.8060 | 0.8304 | 0.834 | 0.849 | 0.8316 | 0.8302 | 1 |

rDNA units were dot plotted against each the human consensus sequence and themselves using ModDotPlot v0.8.4 with the following command:

```
moddotplot static --compare -r 100 -a 45000 -f $rdna_1 $rdna_2 -o
$species_1.$species_2
```

As expected, the 45S gene looks most highly conserved, with more divergence in the intergenic spacer.

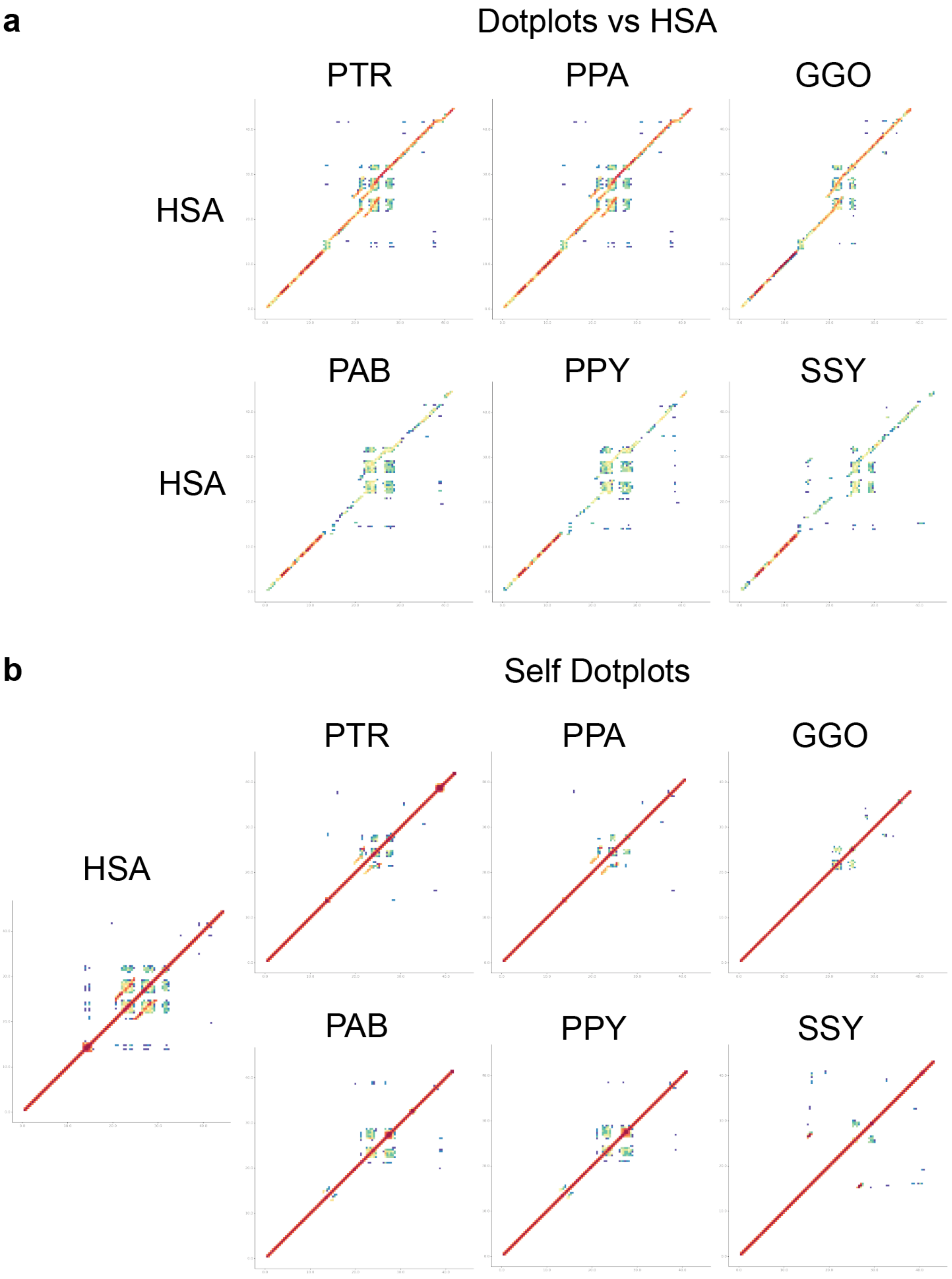
a
Dotplots vs HSA
PTR
PPA
GGO
HSA
PAB
PPY
SSY
HSA
b
Self Dotplots
PTR
PPA
GGO
HSA
PAB
PPY
SSY

**Figure Acro S4. Structure of representative rDNA units.** a) rDNA units from each primate were compared to the human reference rDNA unit KY962518 with ModDotPlot (https://www.ncbi.nlm.nih.gov/nuccore/KY962518) to identify similarities in structure. b) rDNA units from each primate were self-dot plotted to identify satellites and internal repeat structures.

Notably, only a single arm of this palindrome was identified in siamang, with the orientation of these sequence inverted between the two haplotypes.

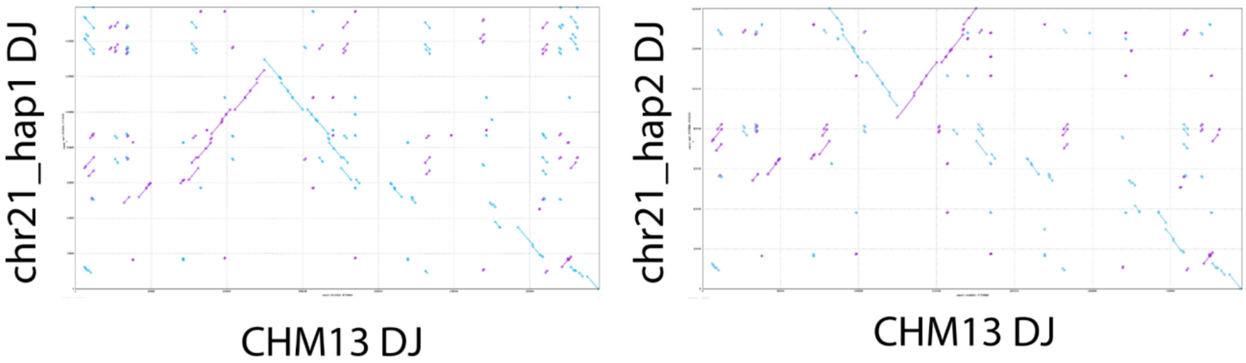


**Figure Acro S5. Comparison of two gibbon DJs to human reference.** The gibbon DJs on both haplotypes of chr21 were extracted. A plot of maximal exact matches to human of at least 20 bp indicates that gibbon has polymorphically lost one arm of the DJ palindrome. With human CHM13 chr13 DJ on the x-axis, the dot plots indicate that chr21_hap1 retained the first arm of the palindrome, while chr21_hap2 retained the second arm.

To dot plot the short arms, the short arms of all chromosomes seen containing an NOR in at least one haplotype were extracted:

```
samtools faidx $primate_ref:$chr:1-$cen_start > $primate.$chr.p_arm.fa
```

For the purposes of plotting, the 1 Mbp rDNA gap in each assembly was replaced with the human reference rDNA unit KY962518 duplicated as many times as indicated by the combined FISH and Illumina copy number quantification. These new sequences were then plotted with ModDotPlot v0.8.4 using scripts in the linked GitHub, where the axis limits were set to just shy of 55 Mbp based on the largest HSA22 short arm, which was mPanPan1 chr22_mat_hsa21.

To quantify satellite compositions, BED files were defined denoting the short arms of all the chromosomes where at least one haplotype contained an NOR in that primate, then used to filter the satellite annotation BED files to only contain records up to the start of the centromere. Then, for each satellite class total base pairs were counted from these files. rDNA bases were defined by multiplying the total copy number per species by 45 kbp, an estimated length of an rDNA unit. This is inexact, given rDNA units vary in size within and across species and individuals, and is meant to be taken as an estimate.

To quantify syntenic and non-syntenic bases on the acrocentric chromosomes, each T2T-CHM13 acrocentric (chr22 shown in the figure) was mashmapped to each of the primate haplotypes separately, and all hits within 1% of the best hit were retained.

```
mashmap -q chm13.$chr.fa -r $primate_ref -t $cpus -M --pi 80 -s 10000 --filter_mode none -o $primate.chm13_$chr.mashmap
```

Then colors were assigned according to what each CHM13 segment hit in the primates based on the key in the figure. Siamang was not included in this analysis due to its mosaic synteny relative to the human chromosomes. Both haplotype's mappings were combined to assess syntenic and non-syntenic bases on the short and long arms.

Hits to multiple haplotypes of the same chromosome were combined, and then each window was checked for all of its best hits. Next, colors were assigned according to the list of best hits based on the key in the figure. If the segment singly mapped to a chromosome that is acrocentric in humans, it was colored accordingly. A single-mapper to a non-acrocentric was colored black. Multimappers were colored light tan if all best hits were to acrocentric chromosomes, and brown if any hits were to non-acrocentric chromosomes. Siamang was not included as described above. Both haplotype's mappings were combined to assess syntenic and non-syntenic bases on the short and long arms.

rDNA conservation was assessed as described above

### XIX. Centromere analyses

Contributing authors:

Glennis A. Logsdon, Hailey Loucks, Karen Miga

#### Methods

##### Centromere identification and annotation

To identify the centromeric regions within each NHP genome, we first aligned the whole-genome assemblies to the T2T-CHM13v2.0 reference genome[167] using minimap2 (v2.24)[27] with the following parameters: -I 10G -a --eqx -x asm20 -s 5000. We filtered the alignments to only those regions that traversed each human centromere, from the p- to the q-arm, using SAMtools (v1.9)[99] and then ran RepeatMasker (v4.1.)[39] to identify regions containing α-satellite sequences, marked by “ALR/Alpha”. Once we identified the regions of the assemblies containing α-satellite repeats, we ran Hum-AS-HMMER (https://github.com/fedorrik/HumAS-HMMER_for_AnVIL) using the hmmer-run_SF.sh script and the AS-SFs-hmmer3.0.290621.hmm Hidden Markov Model. This generated a BED file with each α-satellite suprachromosomal family (SF) designation and its organization along the centromere. We used the α-satellite SF BED file to visualize the organization of the α-satellite higher-order repeat (HOR) arrays with R (v1.1.383)[168] and the ggplot2 package[120].

##### Validation of centromeric regions

We validated the construction of each centromeric region by first aligning native PacBio HiFi and ONT data from the same source genome to each whole-genome assembly using pbmm2 (v1.1.0) (for PacBio HiFi data; https://github.com/PacificBiosciences/pbmm2) or Winnowmap (v1.0) (for ONT data)[169]. We, then, assessed the assemblies for uniform read depth across the centromeric regions via IGV[170] and for collapses, duplications, and misjoins via NucFreq[171]. Centromeres that were found to have a misassembly were flagged and are indicated in the figures.

##### Estimation of α-satellite HOR array length

To estimate the length of the α-satellite HOR arrays of each centromere in the NHP genome assemblies, we first ran Hum-AS-HMMER (https://github.com/fedorrik/HumAS-HMMER_for_AnVIL) on the centromeric regions using the hmmer-run_SF.sh script and the AS-SFs-hmmer3.0.290621.hmm Hidden Markov Model. Then, we used the α-satellite SF BED file to calculate the length of the α-satellite HOR arrays by taking the minimum and maximum coordinate of continuous stretches of α-satellite SFs and plotting their lengths with Graphpad Prism (v9).

#### Pairwise sequence identity heatmaps

To generate pairwise sequence identity heatmaps of each centromeric region, we ran StainedGlass (v6.7.0)[172] with the following parameters: window=5000, mm_f=30000, and mm_s=1000. We normalized the color scale across the StainedGlass plots by binning the % sequence identities equally and recoloring the data points according to the binning.

#### CpG methylation analysis and CDR definition

To determine the CpG methylation status of each NHP centromere, we aligned ONT reads >30 kbp in length from the same source genome to the relevant whole-genome assembly via Winnowmap (v1.0) and then assessed the CpG methylation status of the centromeric regions with Epi2me modbam2bed (https://github.com/epi2me-labs/modbam2bed; v0.10.0) and the following parameters: -e -m 5mC --cpg. We converted the BED file to a bigWig using the bedGraphToBigWig tool (https://www.encodeproject.org/software/bedgraphtobigwig/) and then visualized the file in IGV. To determine the size of hypomethylated region (termed "centromere dip region", or CDR[173] in each centromere, we used CDR-Finder (https://github.com/arozanski97/CDR-Finder). This tool first bins the assembly into 5 kbp windows, computes the median CpG methylation frequency within windows containing α-satellite (as determined by RepeatMasker (v4.1.0)), selects bins that have a lower CpG methylation frequency than the median frequency in the region, merges consecutive bins into a larger bin, filters for merged bins that are >50 kbp, and reports the location of these bins.

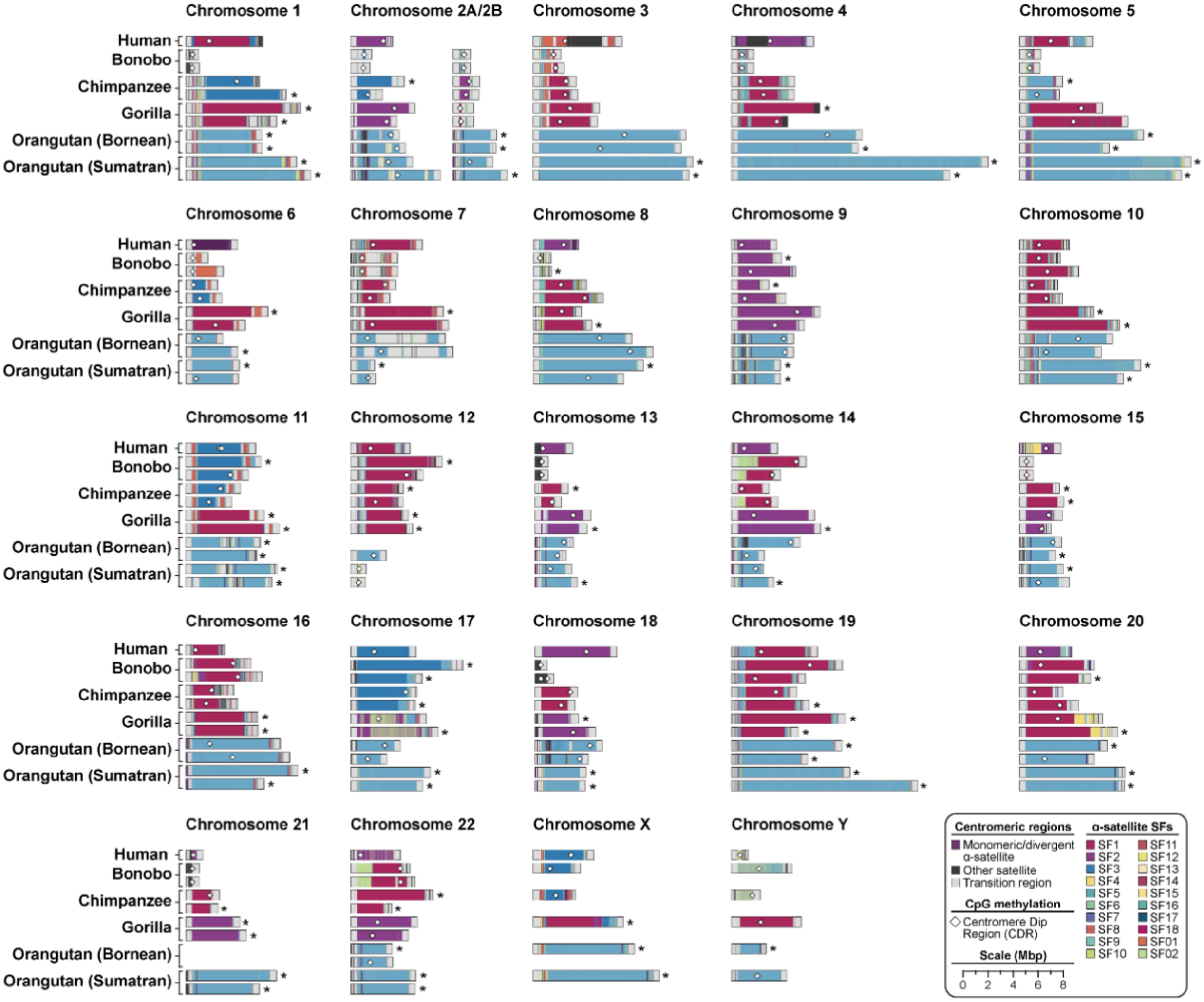


**Figure CEN.S1. Sequence and structure of 237 contiguous centromeres from five NHPs.** Maps of the active α-satellite HOR arrays from the human (CHM13), bonobo, chimpanzee, gorilla, Bornean orangutan, and Sumatran orangutan chromosome centromeres, with the α-satellite suprachromosomal family (SF), indicated for each centromere. Centromeres with an error in their assembly are indicated with an asterisk. Assemblies without any errors have the location of the centromere dip region (or CDR) indicated with a white diamond.

### XX. Subterminal satellite

Contributing authors:

DongAhn Yoo, Evan E. Eichler

#### Methods

Subterminal satellites or pCht repeats present in African great ape species (chimpanzee, bonobo and gorilla) were identified using BLASTN[55] with the consensus sequence (len = 32bp): "gatatttccatgtttatacagatagcggtgta". The blast hit with longer than 90% of the consensus (>28 bp) was recovered. The individual pCht unit was classified into different types based on the variants (small INS, DEL or substitution). The siamang genome, which contains subterminal α-satellite, was investigated using RepeatMasker (v4.1.5)[39].

In addition to the subterminal satellites, spacer SDs interrupting the satellites were investigated were investigated. This was done by subtracting the subterminal satellite arrays from the entire subterminal satellites regions obtained by "*bedtools merge -i [pCht/α-satellite] -d 1000000*". Size distribution was obtained from subtraction of satellites loci from the merged region blocks. Examining distribution of the spacer sequences and their size, 32 kbp highly conserved sequence was identified in Pan lineage and 34 kbp independently in gorilla; on the other hand, multiple modal lengths were identified for siamang spacer with 57 kbp spacers being the most abundant. Using a copy of spacer sequence, spacers were identified using BLASTN (n=793, 800 in chimpanzee and bonobo, and n=974 in gorilla). Methylation status of spacer and subterminal satellites were investigated via ONT long-read alignment against each of the ape genome assembly using Epi2me modbam2bed (https://github.com/epi2me-labs/modbam2bed; v0.10.0; "-*e -m 5mC --cpg*".

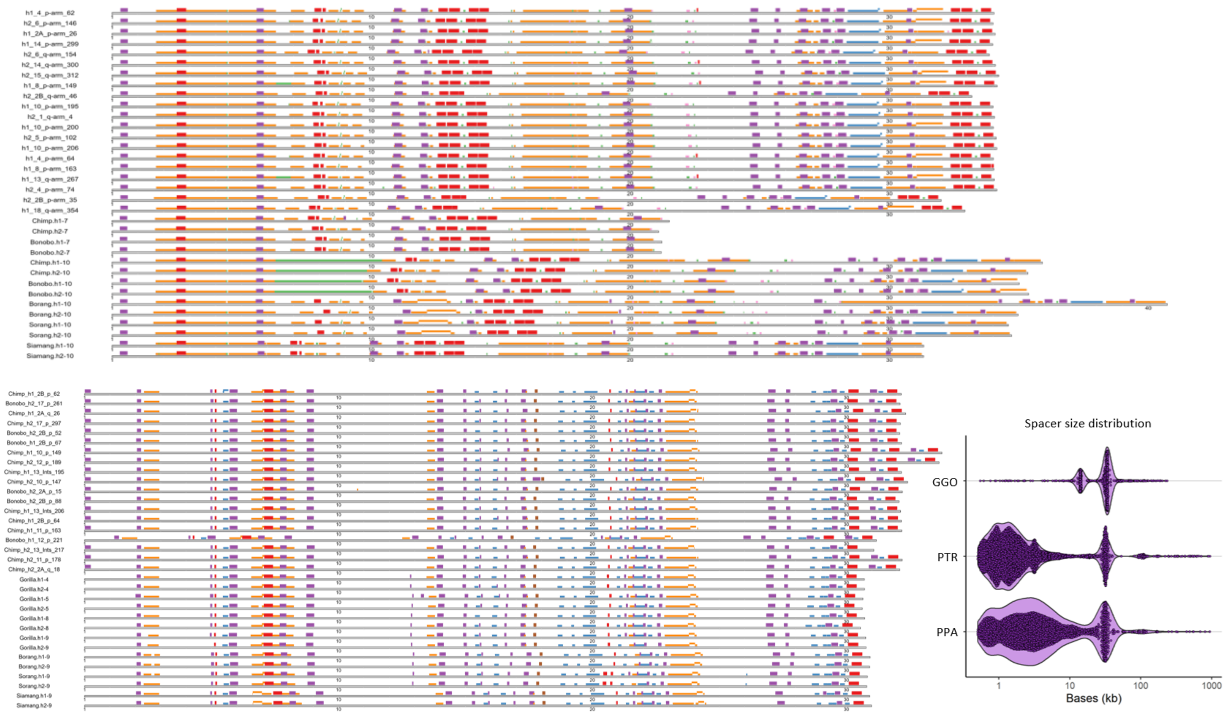


**Figure SuterminalS1. Sequence organization of the subterminal spacer SDs and their ortholog copies.**

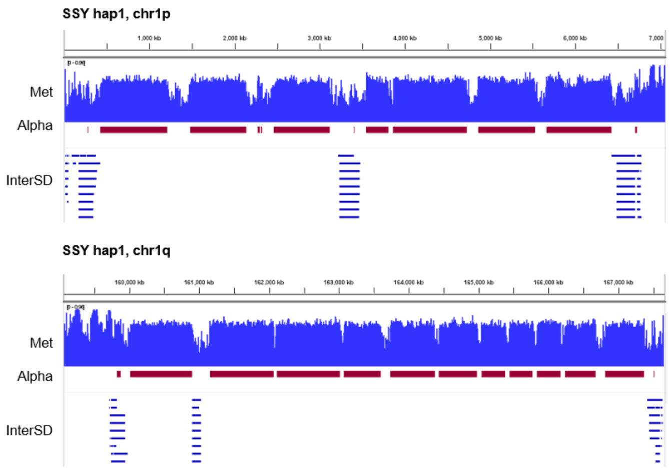


**Figure SuterminalS2. Epigenetic property of spacers, forming hypomethylated pockets, similar to African great ape spacers.**

### XXI. Segmental duplications

Contributing authors:

DongAhn Yoo, David Porubsky, Hyeonsoo Jeong, Evan E. Eichler

#### Methods

SDs were called via SEDEF (v1.1)[174] based on soft-masked genome assemblies - TRF v.4.1.0[41], RepeatMasker v.4.1.5 [39], and Windowmasker (v2.2.22)[175]. The SD calls with sequence identity >90%, length >1 kbp, and satellite content <70% were kept. Lineage-specific SDs were defined by chaining SDs that are within 100 kbp distance and comparing the putative homologous SD loci, defined as containing 100 kbp syntenic sequence flanking the SD. In addition, the SDs that are homologous by location were further checked for the contents using pairwise alignment (minimap 2.26)[27]. The SDs with sequence content (coverage >20%) changed were considered as specific, and SDs with expanded length (>2-fold) were identified. Thus, the SDs with 1) no homologous SDs of other species by position, 2) sequence content changed, and 3) expanded size were quantified. Homozygous and heterozygous genotypes were determined by comparing the two haplotypes. Homologous SDs shared by different apes were classified into phylogenetic branches based on maximum parsimony.

Candidate lineage-specific genes expanded for chr1 GGO double inversion and pongo chr16 expansions were identified by aligning of the gene copies using minimap2 (v2.26) (*-cx asm20 -f 5000 -k15 -w10 -p 0.05 -N 200 -m200 -s200 -z10000 --secondary=yes –eqx*) to find the mapping with >75% of query sequence coverage and >75% of percentage identity. We further screened for Iso-Seq support (indicating read count > 3) and assessment of ORF whether peptide sequence predicted by TransDecoder (https://github.com/TransDecoder/TransDecoder) covers at least >75% of the blastp best hit peptide sequence and percentage identity >80% (the longest transcript). Divergence time among the variable copies was estimated by aligning the complete gene sequence using MAFFT (v7.525)[114] and least square dating of IQTREE2 (v 2.1.2)[121] using chimpanzee-human divergence of 6.4 mya.

Visualization of the alignment of lineage-specific SDs in chr1 GGO double inversion and pongo chr16 expansions was done by alignment of sequences using a PAF file generated by minimap2 (v2.26; options *"-secondary=no -x asm20 -c –eqx*"), followed by breaking of the sequence alignment blocks containing insertions/deletions larger than 100 bp in size using Rustybam ("*rb break-paf -m 100*"), with the SVbyEye R package (https://github.com/daewoooo/SVbyEye) on the resulting PAF file.

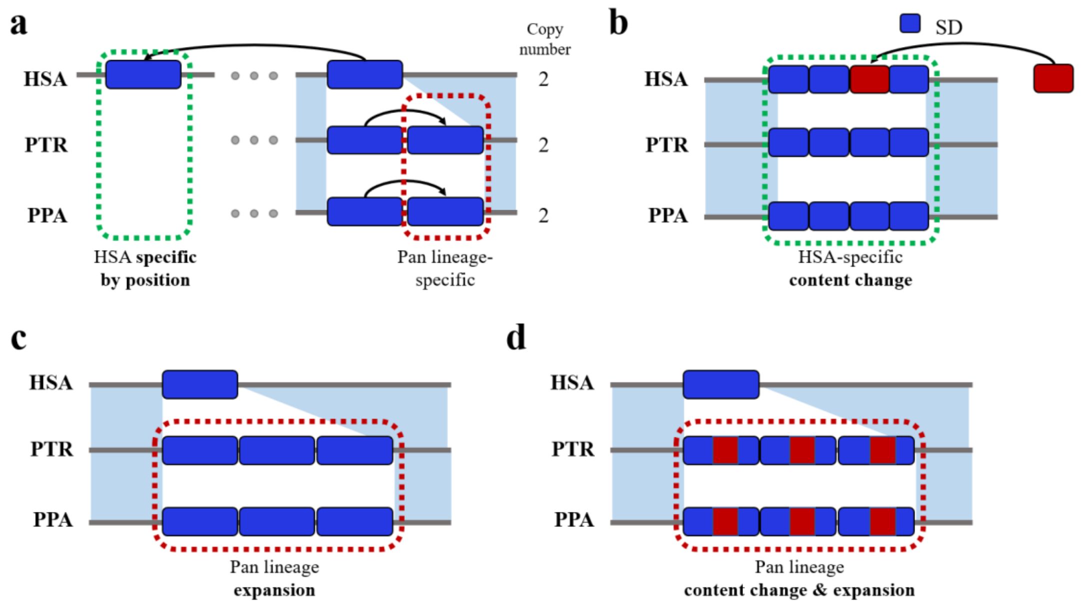


**Figure SD.S1. Examples of lineage-specific SDs. a)** location, **b)** content, **c)** length, and **d)** both content and length.

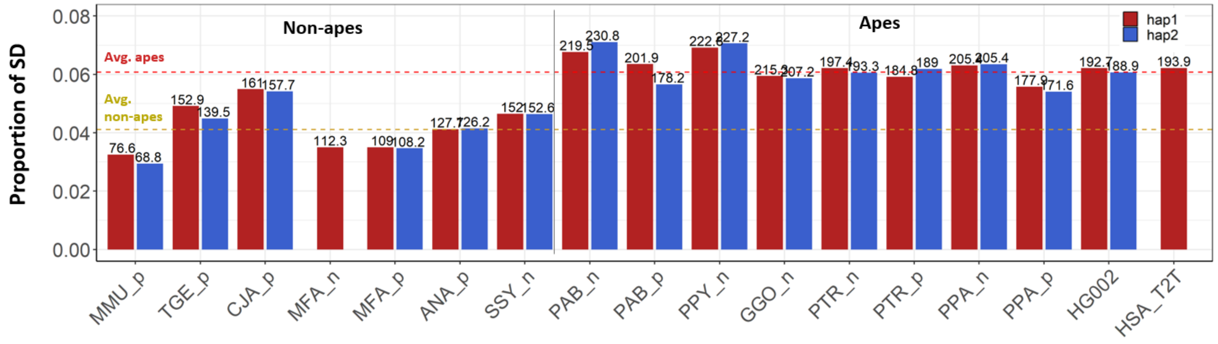


**Figure SD.S2. Total number of SD bases across apes and non-apes.**

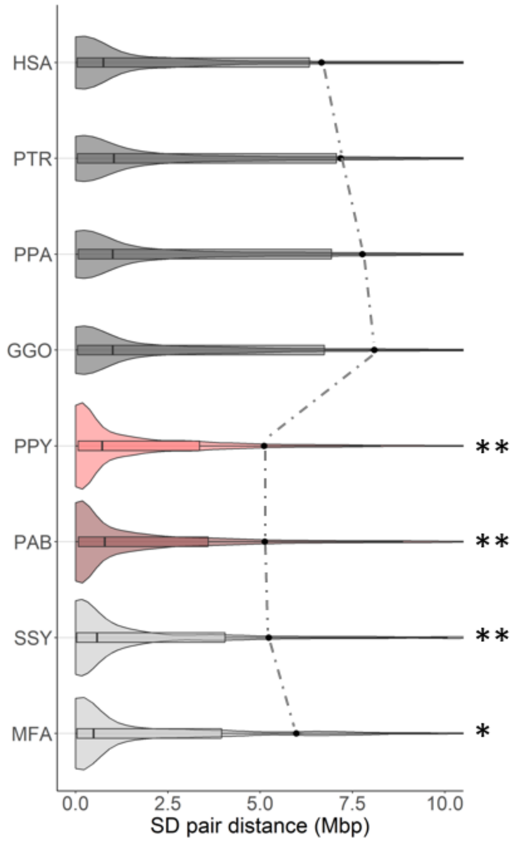


**Figure SD.S3. A violin plot distribution of pairwise SD distance to closest paralog.** The median (black line) and mean (dashed line) are compared for different apes

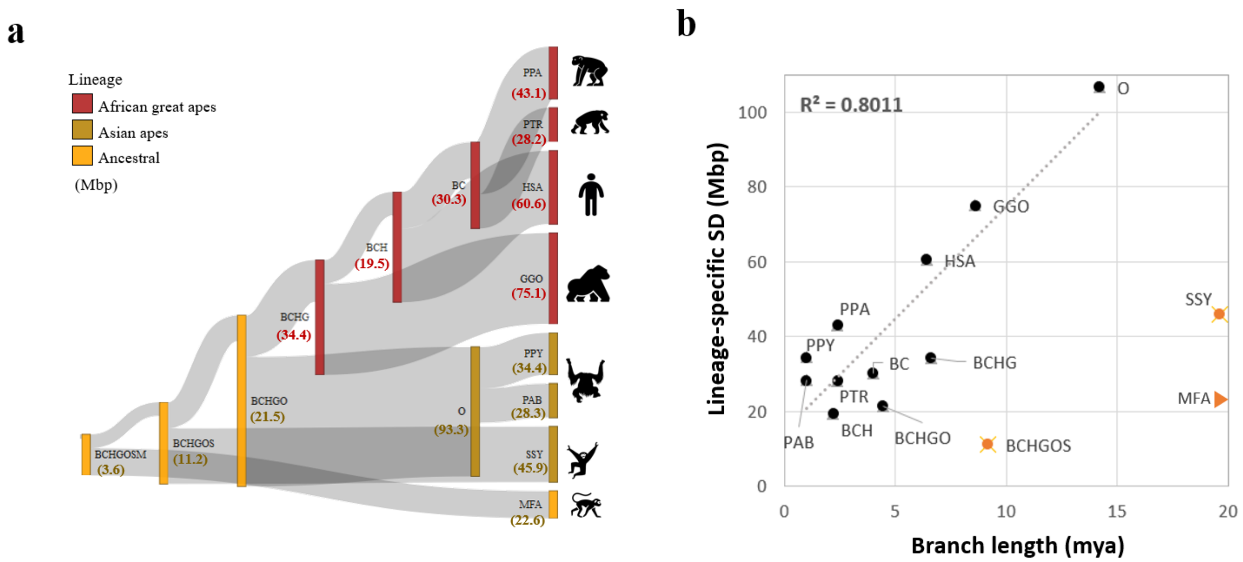


**Figure SD.S4. Identification of lineage-specific SDs.** a) Assignment of SDs (in Mbp) to ancestral and lineage-specific terminal branches based on content, location, and copy number differences (**Fig. SD1**). Asian (dark yellow) and African (red) apes are compared using macaque (MFA) (lighter yellow) as an outgroup. b) Estimated divergence time (based on SNVs) correlates with SD accumulation (r2=0.80) with notable outliers including siamang, macaque, and ancestral branches (e.g., BC= bonobo/chimpanzee, HBC= human/bonobo/chimpanzee ancestral node, etc.).

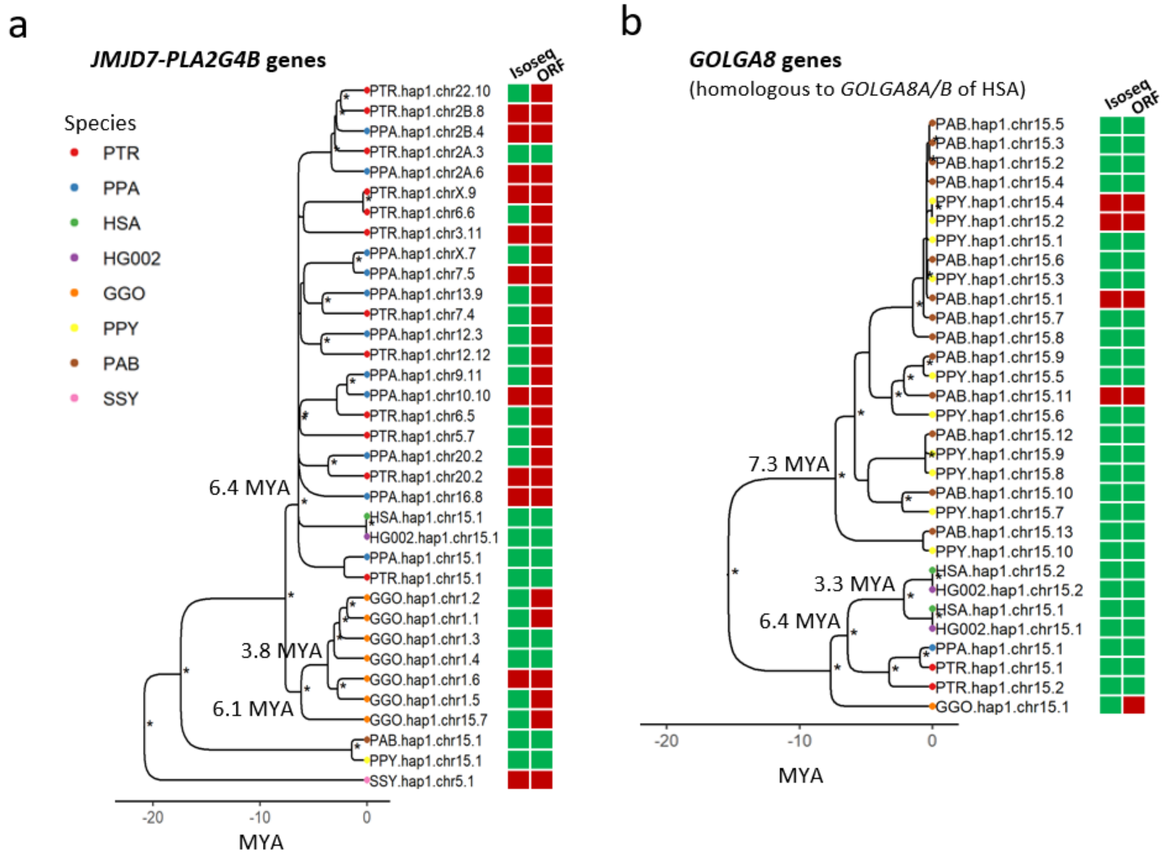


**Figure SD.S5. Phylogeny of the expanded genes.** a) *JMJD7-PLA2G4B* of GGO chr1 double inversion. b) *GOLGA8* expansion of pongo chr16q. Iso-Seq support and valid ORF (allowing for at least 70% of full-length peptide sequence) are indicated by green.

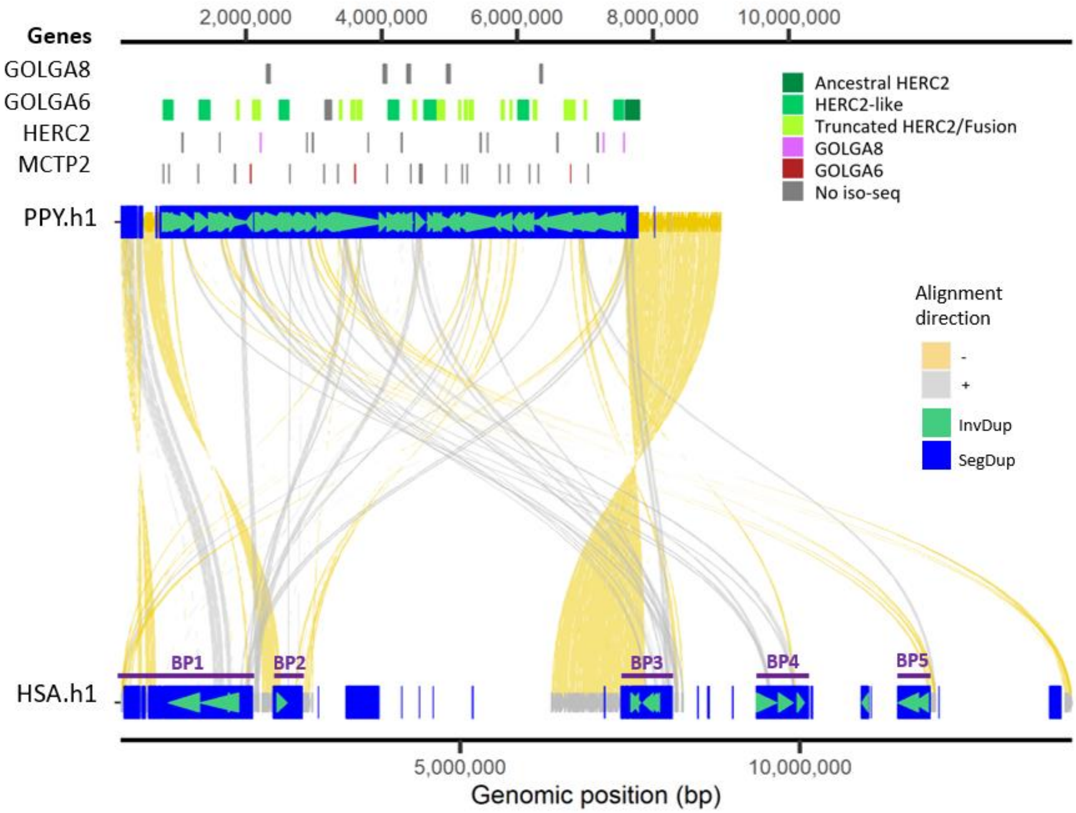


**Figure SD.S6. Zoomed-in view of the largest SD expansion in pongo chr16.**
